# Network Inference from Perturbation Time Course Data

**DOI:** 10.1101/341008

**Authors:** Deepraj Sarmah, Gregory R Smith, Mehdi Bouhaddou, Alan D. Stern, James Erskine, Marc R Birtwistle

**Affiliations:** Department of Chemical and Biomolecular Engineering, Clemson University, Clemson, SC; Department of Neurology, Center for Advanced Research on Diagnostic Assays, Icahn School of Medicine at Mount Sinai, New York, New York, USA; J. David Gladstone Institutes, San Francisco, CA 94158, USA; Department of Cellular and Molecular Pharmacology, University of California San Francisco, San Francisco, CA, 94158, USA; Department of Pharmacological Sciences, Icahn School of Medicine at Mount Sinai, New York, New York, USA; Department of Bioengineering, Clemson University, Clemson, SC

## Abstract

Networks underlie much of biology from subcellular to ecological scales. Yet, understanding what experimental data are needed and how to use them for unambiguously identifying the structure of even small networks remains a broad challenge. Here, we integrate a dynamic least squares framework into established modular response analysis (DL-MRA), that specifies sufficient experimental perturbation time course data to robustly infer arbitrary two and three node networks. DL-MRA considers important network properties that current methods often struggle to capture: (i) edge sign and directionality; (ii) cycles with feedback or feedforward loops including self-regulation; (iii) dynamic network behavior; (iv) edges external to the network; and (v) robust performance with experimental noise. We evaluate the performance of and the extent to which the approach applies to cell state transition networks, intracellular signaling networks, and gene regulatory networks. Although signaling networks are often an application of network reconstruction methods, the results suggest that only under quite restricted conditions can they be robustly inferred. For gene regulatory networks, the results suggest that incomplete knockdown is often more informative than full knockout perturbation, which may change experimental strategies for gene regulatory network reconstruction. Overall, the results give a rational basis to experimental data requirements for network reconstruction and can be applied to any such problem where perturbation time course experiments are possible.

## Introduction

Networks underlie much cellular and biological behavior, including transcriptional, protein-protein interaction, signaling, metabolic, cell-cell, endocrine, ecological, and social networks, among many others. As such, identifying and then representing their structure has been a focus of many for decades now. This is not just from experimental perspectives alone, but predominantly computational with a variety of statistical methodologies that integrate prior knowledge from interaction databases with new experimental data sets (Angulo et al., 2017a; Barabási and Albert, 1999a; Califano et al., 2012a; Calvano et al., 2005a; Dorel et al., 2018; Hackett et al., 2020; Hein et al., 2015a; Hill et al., 2017a, 2017b; Ideker et al., 2001a, 2002a; Liu et al., 2013a; Ma’ayan et al., 2005a; Margolin et al., 2006a; Mazloom et al., 2011a; Mehla et al., 2015a; Molinelli et al., 2013a; Nyman et al., 2020; Pe’er et al., 2001a; Pósfai et al., 2013a; Schraivogel et al., 2020; Shannon et al., 2003a; Stein et al., 2015a; Wynn et al., 2018a; Yuan et al., 2021). Alternatively, a variety of methods have investigated general ways to infer detailed reaction mechanisms—often a foundation of networks—from experimental data (Chevalier et al., 1993; Díaz-Sierra et al., 1999; Hoffmann et al., 2019; Kim et al., 2007; Schmidt et al., 2005). Such tasks may be considered a subset of network inference.

Network structure is usually represented as either an undirected or a directed graph, with edges between nodes specifying the system. There are five main areas where current approaches to reconstructing networks struggle to capture important features of biological networks. The first is directionality of edges (Hackett et al., 2020; Hill et al., 2017a; Morgan and Winship, 2014; Pearl, 2013). Commonly employed correlational methods predominantly generate undirected edges, which impedes causal and other mechanistic analyses. Second is cycles. Cycles such as feedback or feedforward loops are nearly ubiquitous in biological systems and central to their function (Mangan and Alon, 2003a; Reeves, 2019). This also includes an important type of cycle: self-regulation of a node, that is, an edge onto itself, which is rarely considered (Fournier et al., 2007). Third is that biological networks are often dynamic. Two notable examples are circadian and p53 oscillators (Bell-Pedersen et al., 2005a; Stewart-Ornstein et al., 2017a), where dynamics are key to biological function. Directionality and edge signs (i.e. positive or negative) dictate dynamics. Fourth is pinpointing how external variables impinge on network nodes. For example, is the effect of a growth factor on a network node direct, or though other nodes in the network? Fifth, the design and method employed should be robust to typical experimental noise levels. The experimental design and data requirements to uniquely identify the ***dynamic, directed and signed*** edge structures in biological networks containing all types of cycles and external stimuli remains a largely open but significant problem. Any such design should ideally be feasible to implement with current experimental technologies.

Modular Response Analysis (MRA) approaches, first pioneered by Kholodenko and colleagues in 2002 (Kholodenko et al., 2002a; Santra et al., 2018) inherently deal with cycles and directionality by prescribing systematic perturbation experiments followed by steady-state measurements. The premise for data requirements is to measure the entire system response to at least one perturbation for each node. Thus, an *n* node system requires *n* experiments, if the system response can be measured in a global fashion (i.e. all nodes measured at once). The original instantiations struggled with the impact of experimental noise, but total least squares MRA and Monte Carlo sampling helped to improve performance (Andrec et al., 2005a; Santos et al., 2007a; Thomaseth et al., 2018). Incomplete and prior knowledge can be handled as well using both maximum likelihood and Bayesian approaches (Gross and Blüthgen, 2020; Halasz et al., 2016a; Klinger et al., 2013a; Santra et al., 2013a). However, these approaches are based on steady-state data, or fixed time point data, limiting abilities to deal with dynamic systems. There is a formal requirement for small perturbations, which are experimentally problematic and introduce issues for estimation with noisy data. Subsequent approaches have recommended the use of large perturbations as a trade off in dealing with noisy data, but the theory still formally requires small perturbations (Thomaseth et al., 2018). Lastly, there are two classes of biologically relevant edges that MRA does not comprehensively address. First is self-regulation of a node, which is often normalized (to −1) causing it to not be uniquely identifiable. The other are the effects of stimuli external to the network (basally present or administered) on the modeled nodes.

In addition to perturbations, another experimental design feature that can inform directionality is a time-series. One MRA variant (Sontag et al., 2004a) uses time-series perturbation data to uniquely infer a signed, directed network that can predict dynamic network behavior. However, the data requirements are higher than MRA. In an *n* node open system (e.g. protein levels are not constant), multiple nodes need to be distinctly perturbed more than once, such as both production and degradation of a transcript, or phosphorylation and dephosphorylation of a protein. This can be experimentally challenging both in terms of scale and finding suitable distinct perturbations for a node. Moreover, as is often the case, noise in the experimental data severely limits inference accuracy (due to required estimation of 2^nd^ derivatives). A subsequent work on Sontag’s approach (Cho et al., 2005), recommends smaller perturbations and difference in timepoints but also does not address noisy data. Thus, there remains a need for methods that can infer signed, directed networks from feasible perturbation time course experiments that capture dynamics, can uniquely estimate edge properties related to self-regulation and external stimuli, and finally that function in the presence of typical experimental noise levels.

Here we describe a novel, MRA-inspired approach called Dynamic Least-squares MRA (DL-MRA). For an *n*-node system, *n* perturbation time courses are required, and thus experimental requirements scale linearly as the network size increases. The approach uses an underlying network model that captures dynamic, directional, and signed networks that include cycles, self-regulation, and external stimulus effects. We test DL-MRA using simulated timeseries perturbation data with known network topology under increasing levels of simulated noise. The approach has good accuracy and precision for identifying network structure in randomly generated two and three node networks that contain a wide variety of cycles. For the investigated cases, we find between 7 to 11 evenly distributed time points yielded reasonable results, although we expect this will strongly depend on time point placement. We apply the approach to models describing a cell state switching network (Gupta et al., 2011), a signal transduction network (Huang and Ferrell, 1996), and a gene regulatory network (Mangan and Alon, 2003b). Although signaling networks are often a focus in network biology, our analysis suggests they have unique properties that render them generally recalcitrant to reconstruction. Results from the gene regulatory network application suggest that incomplete perturbation (e.g. partial knockdown vs. knockout) is more informative than complete inhibition. While challenges remain for expanding to other and larger systems, the proposed algorithm robustly infers a wide range of networks with good specificity and sensitivity using feasible time course experiments, all while making progress on limitations of current inference approaches.

## Results

### Formulation of Sufficient Experimental Data Requirements for Network Reconstruction

Consider a 2-node network with four directed, weighted edges (Fig. 1a). An external stimulus may affect each of the two nodes differently and its effect is quantified by *S_1,ex_* and *S_2,ex_,* respectively (e.g. Methods, Eq. 15). We also allow for basal/constitutive production in each node (*S_i,b_*). Let *x_i_*(*k*) be the activity of node *i* at time point *t_k_*. The network dynamics can be cast as a system of ordinary differential equations (ODEs) as follows

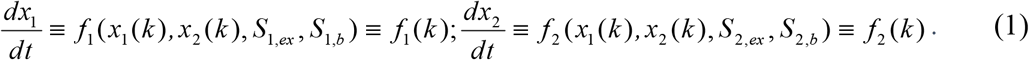

**Figure 1.**
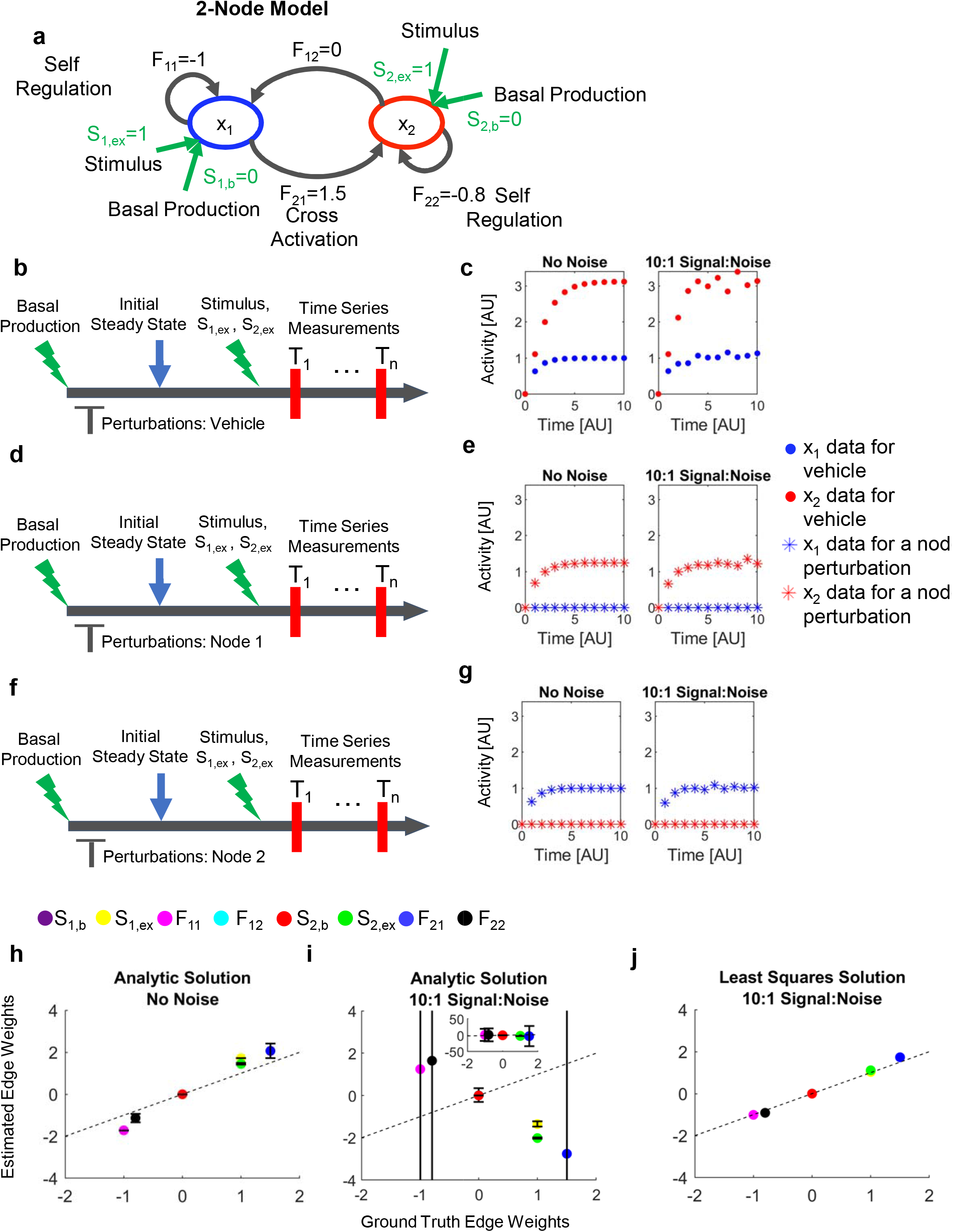
Overall DL-MRA Approach. **(a)** Two-node network with Jacobian elements labeled. Green arrows are stimuli and basal production terms; red arrows are network elements. **(b,d,f)** Time course experimental design with perturbations: vehicle (b), Node 1 (d), Node 2 (f). The vehicle may be the solvent like DMSO for inhibition with a drug, or a nontargeting si/shRNA for inhibition with si/shRNA. **(c,e,g)** Simulated time course data for Vehicle perturbation (c), Node 1 perturbation (e), Node 2 perturbation (g) from the network in (a). Left Column: no added noise; Right Column 10:1 signal-to-noise added. **(h-j)** Actual versus inferred model parameters (*S_1,b_, S_1,ex_, F_11_, F_12_, S_2,b_, S_2,ex_, F_21_, F_22_*) for direct solution of Eq. 3-4 in the absence (h) or presence (i) of noise, or with noise and the least-squares approach (j). In h-i, error bars are standard deviation across time points.

The network edges can be connected to the system dynamics through the Jacobian matrix **J** (Kholodenko et al., 2002a; Santra et al., 2018; Sontag et al., 2004a),

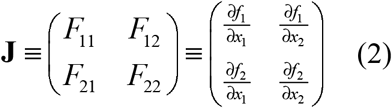

The network edge weights (*F_ij_*’s) describe how the activity of one node affects the dynamics of another node in a causal and direct sense, given the explicitly considered nodes (though not necessarily in a physical sense). In practice, however, causality can only be approached if every component of the system is included in the model, which is not typical (and even more so, there must be no model mismatch, which is almost impossible to guarantee)(Hackett et al., 2020; Höfler, 2005; Morgan and Winship, 2014; Pearl, 2013; Shipley, 2016). In MRA, these nodes may be individual species or “modules”. In order to simplify a complex network it may often be separated into “modules” comprising smaller networks of interconnected species with the assumption that each module is generally insulated from other modules except for information transfer through so-called communicating species (Kholodenko et al., 2002a). Cases where such modules may not be completely isolated are explored elsewhere (Lill et al., 2019).

What experimental data are sufficient to uniquely estimate the signed directionality of the network edges and thus infer the causal relationships within the system? Fundamentally, we know that perturbations and/or dynamics are important for inferring causality (Hackett et al., 2020; Kholodenko et al., 2002a; Lill et al., 2019; Shipley, 2016; Sontag et al., 2004a). Consider a simple setup of three time-course experiments that each measure *x_1_* and *x_2_* dynamics in response to a stimulus (Fig. 1b-g). One time course is in the presence of no perturbation (vehicle), one has a perturbation of Node 1, and one has a perturbation of Node 2. Consider further that the perturbations are reasonably specific, such that the perturbation of *x_1_* has negligible *direct* effects on *x_2_*, and vice versa, and that these perturbations may be large. Experimentally, this could be an shRNA or gRNA that is specific to a particular node, or that a small molecule inhibitor is used at low enough dose to predominantly inhibit the targeted node. A well-posed estimation problem can be formulated (see Methods) that, in principle, allows for unique estimation of the Jacobian elements as a function of time with the following set of linear algebra relations:

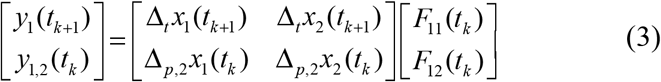

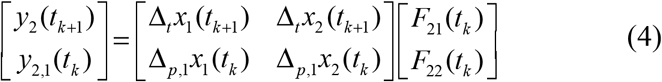

Here, *y_i,j_* refers to a measured first-time derivative of node *i* in the presence of node *j* perturbation (if used), and Δ to a difference with respect to perturbation (subscript *p*) or time (subscript *t*) (see Methods). Since we do not use data from the perturbation of node *i* for estimation of node *i* edges, we do not have to impose assumptions on how the perturbation functionally acts on the system dynamics (see Methods). Moreover, constraints on the perturbation strength can be relaxed, following recent recommendations (Thomaseth et al., 2018)(although accuracy of the underlying Taylor series approximation can affect estimation— see Methods). If these measurements with and without perturbations were each taken in their steady state as is done in MRA, the solution for *F_ij_* would be trivial. MRA gets around this by normalizing self-regulatory parameters *F_ii_* to −1. Using dynamic data allows unique estimation of self-regulatory parameters without such normalization. Estimation of the node-specific stimulus strengths or basal production rates (*S’s*) requires evaluation after specific functional assumptions, but in general these effects are knowable from the data to be generated (see Methods and below results).

Note that this formulation is generalizable to an *n* dimensional network. With *n^2^* unknown parameters in the Jacobian matrix, *n* equations originate from the vehicle perturbation and *n*-1 equations originate from each of the *n* perturbations (discarding equations from Node *i* with Perturbation *i*). This results in *n* + *n*\*(*n*–1) = *n* + *n*^2^ – *n* = *n^2^* independent equations.

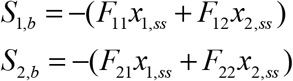

### Using Sufficient Simulated Data to Reconstruct a Network

As an initial test of the above formulation, we used a simple 2 node, single activator network where Node 1 activates Node 2, one node has first-order degradation (−1 diagonal elements), and the other has negative self-regulation (−0.8 diagonal) (Fig. 1a—see Methods for equations). A stimulus at t=0 (time-invariant; *S_,ex_* = 1) increases the activity of each node, which we sample with an evenly spaced 11-point time course. This simulation was done for no perturbation (i.e. vehicle) and for each perturbation (Node 1 and Node 2) to generate the necessary simulation data per the theoretical considerations above (Fig. 1c,e,g, left panel). Here, we modeled perturbations as complete inhibition; for example, a perturbation of Node 1 makes its value 0 at all times. Solving Eqs. 3-4 to infer the Jacobian elements at each time point yielded good agreement between the median estimates and the ground truth values (Fig. 1h, “Analytic Solution”, No Noise). Using the node activity data corresponding to the last time point in the time course and the median estimates of Jacobian elements, the external stimuli *S_1,ex_* and *S_2,ex_* were also determined (Eq.18-19) and reasonably agree with the ground truth values.

How does this approach fare when data are noisy? We performed the estimation with the same data but with a relatively small amount of simulated noise added (10:1 signal-to-noise— Fig. 1c,e,g). The resulting estimates are neither accurate nor precise, varying on a scale more than ten times greater than each parameter’s magnitude with median predictions both positive and negative regardless of the ground truth value (Fig. 1i). The stimulus strengths *S_1,ex_* and *S_2,ex_* are estimated to be negative, while the ground truth is positive.

Although the analytic equations suggest the sufficiency of the perturbation time course datasets to uniquely estimate the edge weights, in practice even small measurement noise corrupts estimates obtained from direct solution of these equations. Therefore, we considered an alternative representation by employing a least squares estimation approach rather than solving the linear equations directly. For a given set of guesses for edge weight and stimulus parameters, one can integrate to obtain a solution for the dynamic behavior of the resulting model, which can be directly compared to data in a least-squares sense. Least squares methods were shown to improve traditional MRA-based approaches(Andrec et al., 2005a; Santos et al., 2007a), but had never been formulated for such dynamic problems. Two hurdles were how to model the effect of a perturbation without (i) adding additional parameters to estimate or (ii) requiring strong functional assumptions regarding perturbation action. We solved these here by using the already-available experimental measurements within the context of the least-squares estimation (see Methods). We applied this approach to the single activator model, 10:1 signal-noise ratio case above where the analytic approach failed. This new estimation approach was able to infer the network structure accurately and precisely (Fig. 1j). We conclude that analytic formulations can be useful for suggesting experimental designs that should be sufficient for obtaining unique estimates for a network reconstruction exercise, but in practice directly applying those equations may not yield precise nor accurate estimates. Alternatively, using a least-squares formulation seems to work well for this application.

### Reconstruction of Random 2 and 3 Node Networks

To investigate the robustness of the least-squares estimation approach, we applied it to increasingly complex networks with larger amounts of measurement noise and smaller numbers of time points (Fig. 2). We focused on 2 and 3 node networks. We generated 50 randomized 2 and 3 node models, where each edge weight is randomly sampled from a uniform distribution over the interval [-2,2], and the basal and external strength from [0,2] (Fig. 2a, S1a, S2a). Each random network was screened for stability. Many networks (29/50 for 2 node and 3 node) displayed potential for oscillatory behavior (non-zero imaginary parts of eigenvalues of Jacobian matrix). However, since the real parts of the eigenvalues are non-zero and negative, these oscillations should dampen over time, and no sustained oscillatory behavior was analyzed. For each random model, we generated a simulated dataset based on the prescribed experimental design, using complete inhibition as the perturbation. We considered evenly-spaced sampling within the time interval of 0-10 AU (approximate time to reach steady-state—Fig. S1b, S2b) with different numbers of time points (3, 7, 11 and 21), and added 10:1 signal-to-noise, 5:1 signal-to-noise, and 2:1 signal-to-noise to the data. Non-uniform time point spacing may change inference results but that was not explored at these first investigations.

**Figure 2.**
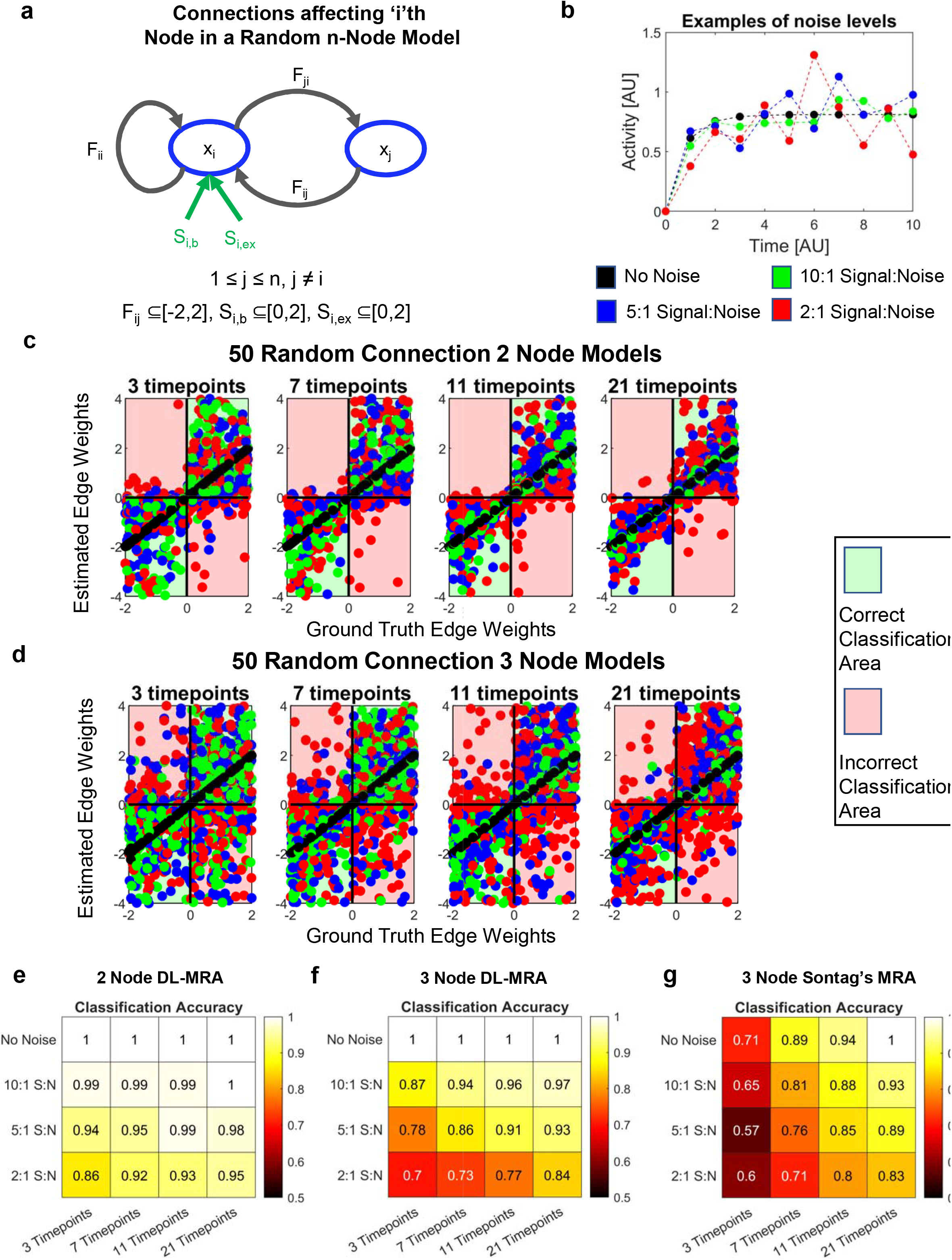
Application to Linear Two and Three Node Models. **(a)** Connections around a Node *i* in an n-Node Model. *S_i,b_* and *S_i,ex_* are the basal production and external stimulus terms acting on Node *i,* respectively. *F_ii_* is the self-regulation term; *Fij* the effect of Node *j* on Node *i* and *F_ji_* the effect of Node *i* on Node *j.* **(b)** Example of different signal-to-noise ratio effects on time course data. **(c,d)** Ground truth versus estimated edge weights across all 50 random networks and noise levels for data from four different total timepoints (3,7,11,21) for 2 node (c) and 3 node (d) networks. Quadrant shading indicates edge classification. **(e,f)** Fraction of network parameters correctly classified in 50 randomly generated 2 node networks (e) and 3 node networks (f) with different noise levels and total timepoints. (g) Fraction of network parameters correctly classified in 50 randomly generated 3 node networks with Sontag’s dynamic MRA using two sets of perturbation data.

For each random network model, number of time points, and noise level, we evaluated the fidelity of the proposed reconstruction approach in terms of signed directionality (Fig. 2c-f). We overall found reasonable agreement between inferred and ground truth values, even at the higher noise levels and low number of timepoints. Expectedly, the overall classification accuracy increases with more time points and decreases with higher noise levels. But, surprisingly, even in the worst case investigated of 3 timepoints and 2:1 signal-to-noise ratio, classification accuracy was above 85% for 2 node models and 70% for 3 node models. Increasing the number of nodes decreases performance, with 3-node reconstruction being slightly worse than 2-node reconstruction, other factors held constant.

We wondered whether the magnitude of an edge weight influenced its classification accuracy, since small edge weights may be more difficult to discriminate from noise. We found that edge weights with greater absolute values, which are expected to have a greater influence on the networks, were more likely to be classified correctly (Fig. S1c-f, S2c-f). Also, for models with damped oscillatory behavior, the classification accuracy is very similar to that of all 50 random models (Fig. S3a-b).

How does this method compare to similar network reconstruction methods? There are limited methods to compare to which also use dynamic data and sequential perturbations. MRA (Kholodenko et al., 2002a), from which this method was inspired, uses steady-state data. However, Sontag’s method (Sontag et al., 2004a) requires dynamic perturbation data as is used in our method, although it requires two perturbations per node for this set of models (double the data). To compare, we further generated another set of perturbation data with 50% perturbation (as opposed to 100%). We then used the two sets of perturbation data to estimate the network node edges with dynamic modular response analysis (Fig 2g). Even in absence of noise, for low to medium numbers of timepoints (3-11) the network is not always accurately inferred (Fig. 2g). In the presence of noise, DL-MRA performs better, although the difference between the two methods becomes lower at high number of timepoints. Thus, DL-MRA not only outperforms with half the data, but it also estimates 6 additional parameters-basal production and external stimulus for each node. Although Cho’s approach (Cho et al., 2005) builds upon Sontag’s method by recommending smaller time point intervals and smaller perturbations, for our purposes, the time intervals and perturbations are fixed and the results from Cho and Sontag’s approach would be similar. Moreover, subsequent work has recommended larger perturbations while dealing with noisy data (Thomaseth et al., 2018).

To explore a scenario where data from a node might be unavailable, we removed the data from one of the nodes in the 50 random 3 node models and used the remaining data to reconstruct a 2-node system (Fig. S4). Comparing with corresponding model parameters in the 3 node system, we find a good but expectedly reduced classification accuracy (No Noise-94.75%, 10:1 Signal: Noise-93.75%, 5:1 Signal: Noise-91.25%, 2:1 Signal: Noise-87).

A part of the inference process is performing parameter estimation using multiple starting guesses (i.e. multi-start), and we wanted to determine how robust the estimated parameters were across the multi-start processes. We looked at the distribution of coefficient of variation (CV) among the parameters in the multi-start results in the 50 random 3 node models where either the data generated from the estimated parameters had low sum of squared errors (SSE) compared to the original data (<10^-4^) or with SSE less than twice the minimum SSE. We find that the CVs peak around zero and generally have a small spread, especially for low noise scenarios (Fig. S5). This implies a good convergence of the parameter sets obtained through multi-start.

We conclude that the network parameters of 2 and 3 node systems can be robustly and uniquely estimated using DL-MRA. However, these were ideal conditions where there was no model mismatch that is expected in specific biological applications. How does DL-MRA perform when applied to data reflective of different biological use cases?

### Application to Cell State Networks

Cell state transitions are central to multi-cellular organism biology. They are commonly transcriptomic in nature and underlie development and tissue homeostasis and can also play roles in disease, such as drug resistance in cancer (Armond et al., 2014; Dirkse et al., 2019; Gupta et al., 2011; Hormoz et al., 2016a; Larsson et al., 2021; Neftel et al., 2019; Sha et al., 2020; Shen and Clairambault, 2020; Zarkoob et al., 2013). Could DL-MRA reconstruct cell state transition networks? As the application, we use previous data on SUM159 cells that transition between luminal, basal and stem-like cells (Gupta et al., 2011). Pure populations of luminal, basal and stem-like cells eventually grow to a stable final ratio amongst the three. The authors used a discrete time Markov transition probability model to describe the data and estimate a cell state transition network (Fig. 3a). Thus, we seek to compare DL-MRA to such a Markov model in this case.

**Figure 3.**
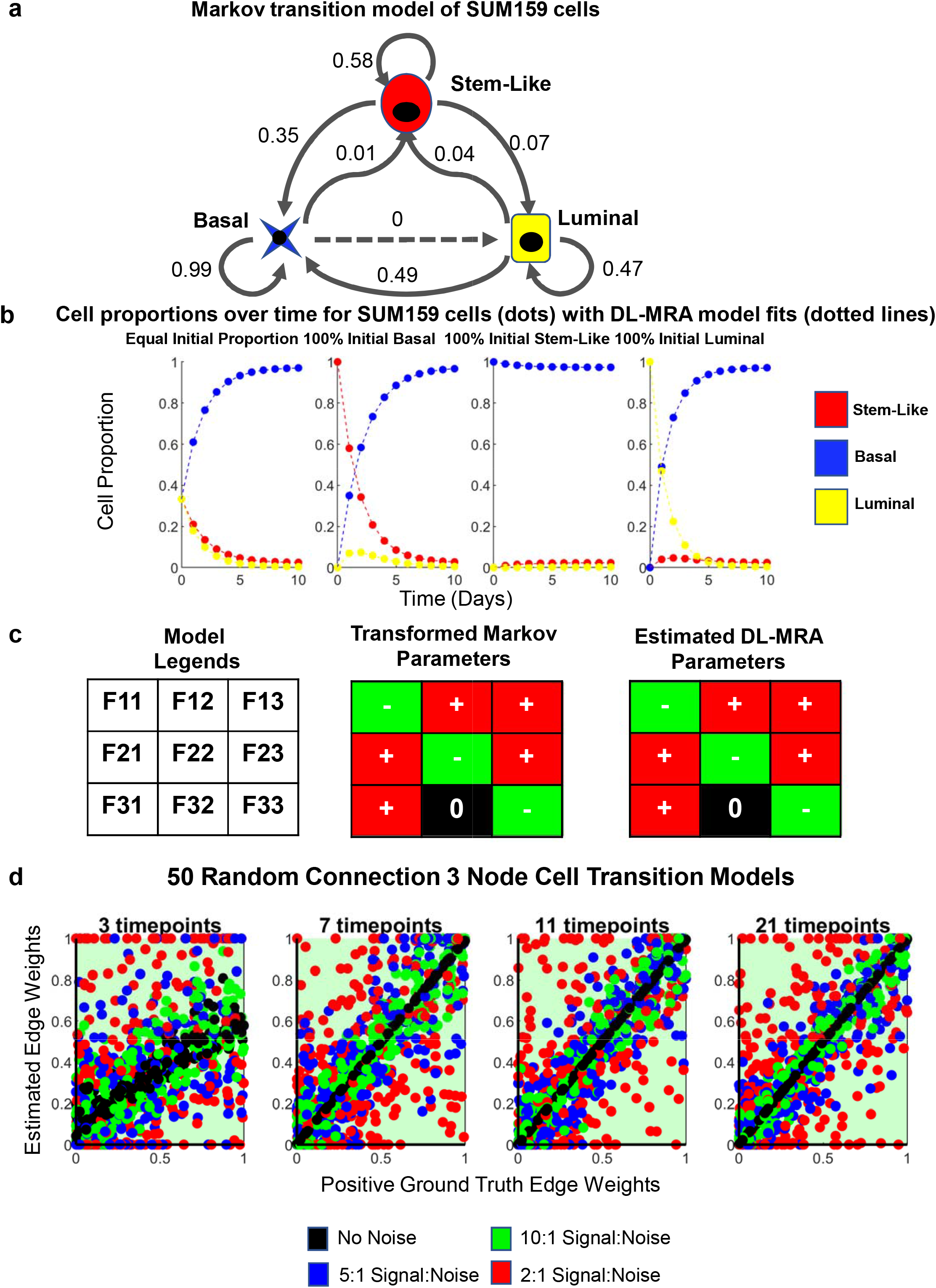
Application to Cell State Transition Networks. **(a)** Markov transition model of SUM159 cell states. **(b**) Cell proportions over time for SUM159 cells using Markov transition parameters (dots), starting at different initial proportions and respective DL-MRA model fits (lines). **(c)** Parameters from DL-MRA estimates of SUM159 data are similarly classified as transformed Markov parameters (See Methods, Eq. 29-30). **(d)** Ground truth versus estimated edge weights across 50 random cell transition networks and noise levels for data from four different total timepoints (3,7,11,21).

We hypothesized that perturbations to the system in this case, in contrast to above, did not have to change node activity (i.e. edges). Rather, we thought that perturbing the equilibrium cell state distribution could serve an equivalent purpose. Thus, the data for reconstruction consisted of observing the cell state proportions evolve over time from “pure” populations (Fig. 3b), in addition to equal proportions. DL-MRA is capable of explaining the data (Fig. 3b). Interpretation of the estimated network parameters to DL-MRA depends on the transformation of the original discrete time Markov probabilities to a continuous time formulation (see Methods—there are constraints on self-regulatory parameters), but DL-MRA correctly infers the cell state transition network as well (Fig 3c). Conveniently, DL-MRA is not constrained to 1-day time point spacing as is the original discrete time Markov model.

How does noise and the number of timepoints affect the reconstruction? As above, we generated data for 50 random cell state transition models with 3, 7, 11 and 21 timepoints within 5 days, as the models generally seemed to reach close to equilibrium within 5 days. Noise levels of 10:1, 5:1 and 2:1 were used. All parameters were classified accurately (Fig. 3d) (although additional constraints in the estimation—see Methods—facilitates this classification performance). With 3 timepoints, there was deviation from perfect fit even with no noise in the data. At 7 and higher number of timepoints, the estimates matched ground truth well, and noise expectedly reduced the accuracy (Fig. 3d). We conclude that DL-MRA can robustly infer cell state networks given perturbation data in the form of non-equilibrium proportions as initial conditions.

### Application to Intracellular Signaling Networks

How does the method perform for intracellular signaling networks? The Huang-Ferrell model (Huang and Ferrell, 1996) (Fig. 4a) is a well-known intracellular signaling pathway model and has been investigated by different reconstruction methods, including previous versions of MRA (Andrec et al., 2005b; Henriques et al., 2017; Kholodenko et al., 2002b; Sontag et al., 2004a; Thomaseth et al., 2018). It captures signal flux through a three-tiered MAPK cascade where the 2^nd^ and 3^rd^ tier contain two phosphorylation sites. An important aspect of the Huang-Ferrell model is that although the reaction scheme is a cascade and without obvious feedbacks, there may be hidden feedbacks due to sequestration effects and depending on how the perturbations were performed.

**Figure 4.**
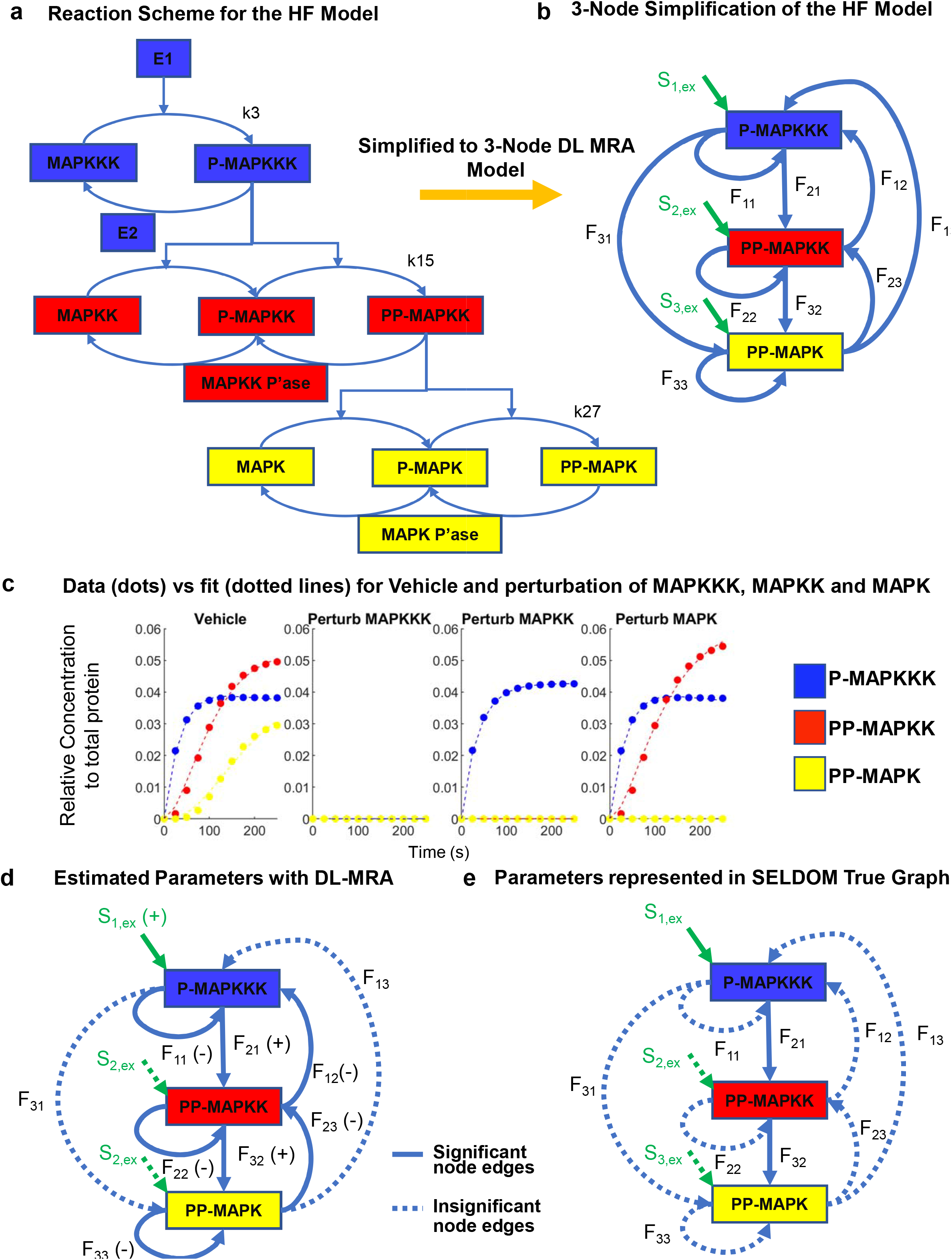
Application to a Signaling Network. **(a)** Full Reaction scheme for the Huang-Ferrell (HF) Model, depicting the parameters k3, k15 and k27 which were perturbed sequentially to generate the perturbation data. **(b)** Model coarse-graining to a 3-node network. **(c)** Data generated for each node with a small E1 stimulus (2.5×10^-6^ uM). **(d)** Model parameters estimated as significant (bold) and negligible (dotted lines). **(e)** SELDOM true graph values represented in the 3-node model with parameters considered (bold) and not considered (dotted lines).

In order to reconstruct the Huang-Ferrell MAPK network through DL-MRA, we first simplified it to a three-node model with p-MAPKKK, pp-MAPKK and pp-MAPK as observable nodes, as is typical for reconstruction efforts (Fig. 4b) (Andrec et al., 2005a; Henriques et al., 2017; Kholodenko et al., 2002b; Lill et al., 2019; Sontag et al., 2004a; Thomaseth et al., 2018). Second, to model perturbations, we sequentially perturbed the activation parameters of each of the observable species (k3, k15 and k27 respectively). Such perturbations, although hard to achieve experimentally, are important because modules must be “insulated” from one another and perturbations must be specific to the observables (Kholodenko et al., 2002b; Lill et al., 2019). Third, in the simplification of the reaction scheme, the observables are shown to influence each other but in the actual scheme, they conduct their effects through the unphosphorylated and semi-phosphorylated species. We sought to keep the levels of these two species relatively constant between different perturbations, so that they wouldn’t add to non-linearities in the estimation. Therefore, we used a stimulus which only activated the observables to a maximum of about 5% of the total forms of the protein (Lill et al., 2019)

Estimation with DL-MRA under the above conditions fits the data (Fig 4c) and predicts positive node edges down the reaction cascade (F_21_, F_32_), negligible direct relation between p-MAPKKK and pp-MAPK (F_13_, F_31_), negative self-regulation of each of the observables (F_11_, F_22_, F_33_) negative feedbacks from pp-MAPKK to p-MAPKKK (F_12_) and from pp-MAPK to pp-MAPKK (F_23_), and negligible external stimuli on pp-MAPK to pp-MAPKK (F_13_, F_31_). All these effects are consistent with the reaction scheme. The negative feedback effects, although not immediately obvious, are consistent with ground truth sequestration effects. For instance, pp-MAPK has an overall negative effect on pp-MAPKK as the existence of pp-MAPK lowers the amount of species MAPK and p-MAPK which sequester pp-MAPKK and makes it avoid deactivation by its phosphatase.

How do the estimation results for the Huang Ferrell model in our method compare with those obtained from other methods? Previous work using MRA also reported negative feedbacks from successive modules to the preceding ones (Kholodenko et al., 2002a; Lill et al., 2019; Sontag et al., 2004a). Similarly, self-regulation parameters in most preceding MRA based methods are also estimated to be negative but are fixed at −1(Andrec et al., 2005b; Kholodenko et al., 2002b; Lill et al., 2019).

Besides MRA inspired methods, SELDOM is another network reconstruction method which can also deal with dynamic data (Henriques et al., 2017). SELDOM is a data-driven method which uses ensembles of logic based dynamic models followed by training and model reduction steps to predict state trajectories under untested conditions. However, when dealing with the Huang-Ferrell network, the true value model of SELDOM does not map the effects of self-regulation, nor feedback effects between nodes (Fig 4e). This may be explained by the fact that although SELDOM uses an extensive number of models to test the data obtained from multiple different stimuli, perturbation data was not included to test the Huang-Ferrell Model. This implies that systematic perturbation of each of the nodes, as prescribed by MRA-based methods, are necessary in order to unearth feedbacks and self-regulation effects.

Although application of DL-MRA to the Huang-Ferrell model was able to unearth latent network structure, the simulation conditions were restrictive. First, the perturbation scheme chosen in this paper, although specifically targeted at the observable species, is hard to produce experimentally. The feedback effect observed could also depend on the perturbation scheme chosen-for instance knockdown of an entire module as a perturbation would likely have manifested as positive feedback to the preceding module. That is because such a knockdown would have reduced the effect of sequestration of the module on the preceding observable and would have made it more available for dephosphorylation. Second, we assumed a low stimulus to avoid effects from the unphosphorylated version of the proteins. A higher activation may increase non-linearities adding to the complexity of the model, whereas a lower stimulus may not activate enough proteins to be well detected in experiments. The degree of activation needed for an experiment may be hard to predict beforehand. Such specific perturbations and stimulus had to be done to reduce the effects arising from the non-observable species behavior. Hence application of DL-MRA to intracellular signaling networks with multiple physical interactions needs to be carefully considered before modeling or experiments.

### Application to Gene Regulatory Networks: Partial Perturbations are More Informative than Full Perturbations

Here, we applied DL-MRA further to a series of well-studied non-linear feed forward loop (FFL) gene regulatory network models that have time-varying Jacobian elements (Fig. 5a, Table 1) (Mangan and Alon, 2003a; Reeves, 2019). Such FFL motifs are strongly enriched in multiple organisms and are important for signaling functions such as integrative control, persistence detection, and fold-change responsiveness (Goentoro and Kirschner, 2009a; Goentoro et al., 2009a; Nakakuki et al., 2010a).

**Figure 5.**
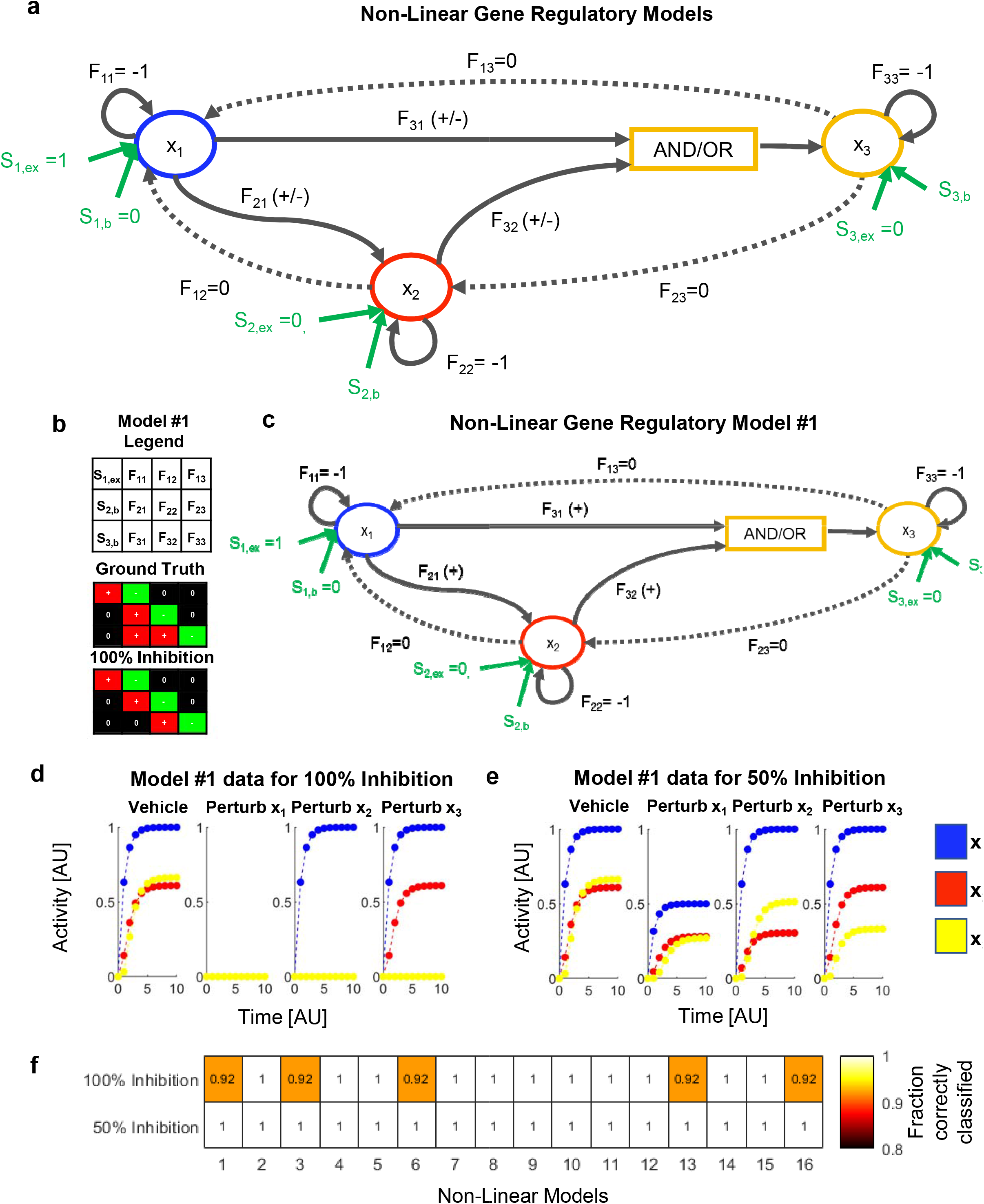
**(a)** Feedforward loop (FFL) network models. Across all 16 models (Table 1), F_11_, F_22_, and F_33_ values are fixed at −1 and F_12_, F_13_, and F_23_ values are fixed at 0. F_21_, F_31_, and F_32_ values can be positive or negative depending on the model. The combined effect of x_1_ and x_2_ on x_3_ is described by either an AND gate or an OR gate. There are 16 possible model structures (Table 1). **(b)** 100% inhibitory perturbations may not provide accurate classification even without noise. In Model #1, F_31_ is positive (ground truth) but is estimated as null. **(c)** Specific structure of Model #1. **(d)** Node activity simulation data for 100% inhibition in Model #1, implying that it is impossible to infer F_31_ from such data. **(e)** Node activity simulation data for 50% inhibition in Model #1, showing potential to infer F_31_. **(f)** Fraction of model parameters correctly classified in all the 16 non-linear models without noise, for 100% inhibition vs 50% inhibition.

**Table 1.** Structure of each of the 16 non-linear models.

| Model# | F<sub>21</sub> | Gate | F<sub>31</sub> | F<sub>32</sub> |
| --- | --- | --- | --- | --- |
| 1 | Activator | AND | Activator | Activator |
| 2 | Activator | AND | Activator | Repressor |
| 3 | Activator | AND | Repressor | Activator |
| 4 | Activator | AND | Repressor | Repressor |
| 5 | Activator | OR | Activator | Activator |
| 6 | Activator | OR | Activator | Repressor |
| 7 | Activator | OR | Repressor | Activator |
| 8 | Activator | OR | Repressor | Repressor |
| 9 | Repressor | AND | Activator | Activator |
| 10 | Repressor | AND | Activator | Repressor |
| 11 | Repressor | AND | Repressor | Activator |
| 12 | Repressor | AND | Repressor | Repressor |
| 13 | Repressor | OR | Activator | Activator |
| 14 | Repressor | OR | Activator | Repressor |
| 15 | Repressor | OR | Repressor | Activator |
| 16 | Repressor | OR | Repressor | Repressor |

The FFL network has three nodes (*x_1_*, *x_2_,* and *x_3_*), and the external stimulus acts on *x_1_* (*S_1,ex_*). There is no external stimulus on *x_2_* and *x_3_*; however, there may be basal production of *x_2_* (*S_2,b_*) and *x_3_* (*S_3,b_*),. Each node exhibits first-order decay (*F_ii_*=-1). The parameters *F_12_, F_13_,* and *F_23_* represent connections that do not exist in the model; we call these null edges, but we allow them to be estimated. The relationship between *x_1_* and *x_2_* (*F_21_*), between *x_1_* and *x_3_* (*F_31_*), or between *x_2_* and *x_3_* (*F_32_*) can be either activating or inhibitory. Furthermore, *x_1_* and *x_2_* can regulate *x_3_* through an “AND” gate (both needed) or an “OR” gate (either sufficient) (Fig. 5a). These permutations give rise to 16 different FFL structures (Table 1).

To generate simulated experimental data from these models, we first integrated the system of ODEs starting from a *zero* initial condition to find the steady state in the absence of stimulus. We then introduced the external stimulus and integrated the system of ODEs (see Methods) to generate time series perturbation data consistent with the proposed reconstruction algorithm, using full inhibitory perturbations. We used 11 evenly spaced timepoints for all 16 non-linear models, based on the random 3-node model analysis above, and also added noise as above.

We first noticed that even in the absence of added noise, a surprising number of inferences were incorrect (Fig. 5b, f). Model #1 (Table 1, Fig. 5b-c) is used as an example, where *F_21_, F_31_* and *F_32_* are activators with an AND gate, and *F_31_* is incorrectly predicted as null (Fig. 5b—compare ground truth to 100% inhibition). To understand the reason for the incorrect estimation, we looked at the node activity dynamics across the perturbation time courses (Fig. 5d). All three nodes start from an initial steady state of zero, but Node 3 is zero for all three perturbation cases. This is because of the following. Since x_1_ is required for the activation of x_2_ and x_3_, complete inhibition of x1 completely blocks both x_2_ and x_3_ activation. But, because both x_1_ and x_2_ are required for the activation of x_3_, completely inhibiting x_2_ activity also completely inhibits x_3_. Thus, given this experimental setup, it is impossible to discern if x_1_ directly influences x_3_ or if it acts solely through x_2_.

We thus reasoned that full inhibitory perturbation may suppress the information necessary to correctly reconstruct the network, but that a partial perturbation experiment may contain enough information available to make a correct estimate. If this were true, then upon applying partial perturbations (we chose 50% here), Node 3 dynamics should show differences across the perturbation time courses. Simulations showed that this is the case (Fig. 5e). Subsequently, we found that for partial perturbation data, *F_31_* is correctly identified as an activator. More broadly, we obtain perfect classification from noise-free data across all 16 FFL networks when partial perturbation data are used, as opposed to 5/16 networks having discrepancy with full perturbation data (Fig. 5f). The fits to simulated data from the reconstructed model align very closely, despite model mismatch (Fig. S6). We conclude that in these cases of non-linear networks, a partial inhibition is necessary to estimate all the network parameters accurately. Thus, moving forward, we instead applied 50% perturbation to all simulation data and proceeded with least squares estimation.

### Application to Gene Regulatory Networks: Performance

The above analysis prompted us to use a partial (50%) perturbation strategy, since it classified each edge for each model in the absence of noise correctly. What classification performance do we obtain in the presence of varying levels of experimental noise? We first devised the following strategy to assess classification performance. We generated 50 bootstrapped datasets for each network structure/signal-to-noise pair, and thus obtained 50 sets of network parameter estimates. To classify the network parameters, we used a symmetric cutoff of a percentile window around the median of these 50 estimates (Fig. 6a). We illustrate this approach with three different example edges and associated estimates, one being positive (Edge 1), one being negative (Edge 2), and one being null (Edge 3). Given the window of values defined by the percentile cutoff being chosen, if the estimates in this window are all positive, then the network parameter would be classified as positive. Similarly, if the estimates in this window are all negative, then the parameter would be classified as negative. Finally, if the estimates in the window cross zero (i.e. span both positive and negative terms), then it would be classified as null. First, consider the case that the percentile window is just set at the median with no percentile span. Then, the classifications for true positives and negatives are likely to be accurate while the null parameters are likely to be incorrectly categorized as either positive or negative (Fig. 6a). If we increase the percentile window span slightly (e.g. between the 40th and 60^th^ percentile, middle panel), we can categorize null edges better, while maintaining good classification accuracy of both true positive and negative edges. However, if we relax the percentile window too much, (e.g. between the 10th and 90^th^ percentile, far right panel) we may categorize most parameters as null, including the true positive and negatives. Thus, it is clear there will be an optimal percentile cutoff that maximizes true positives and minimizes false positives as the threshold is shifted from the median to the entire range.

**Figure 6.**
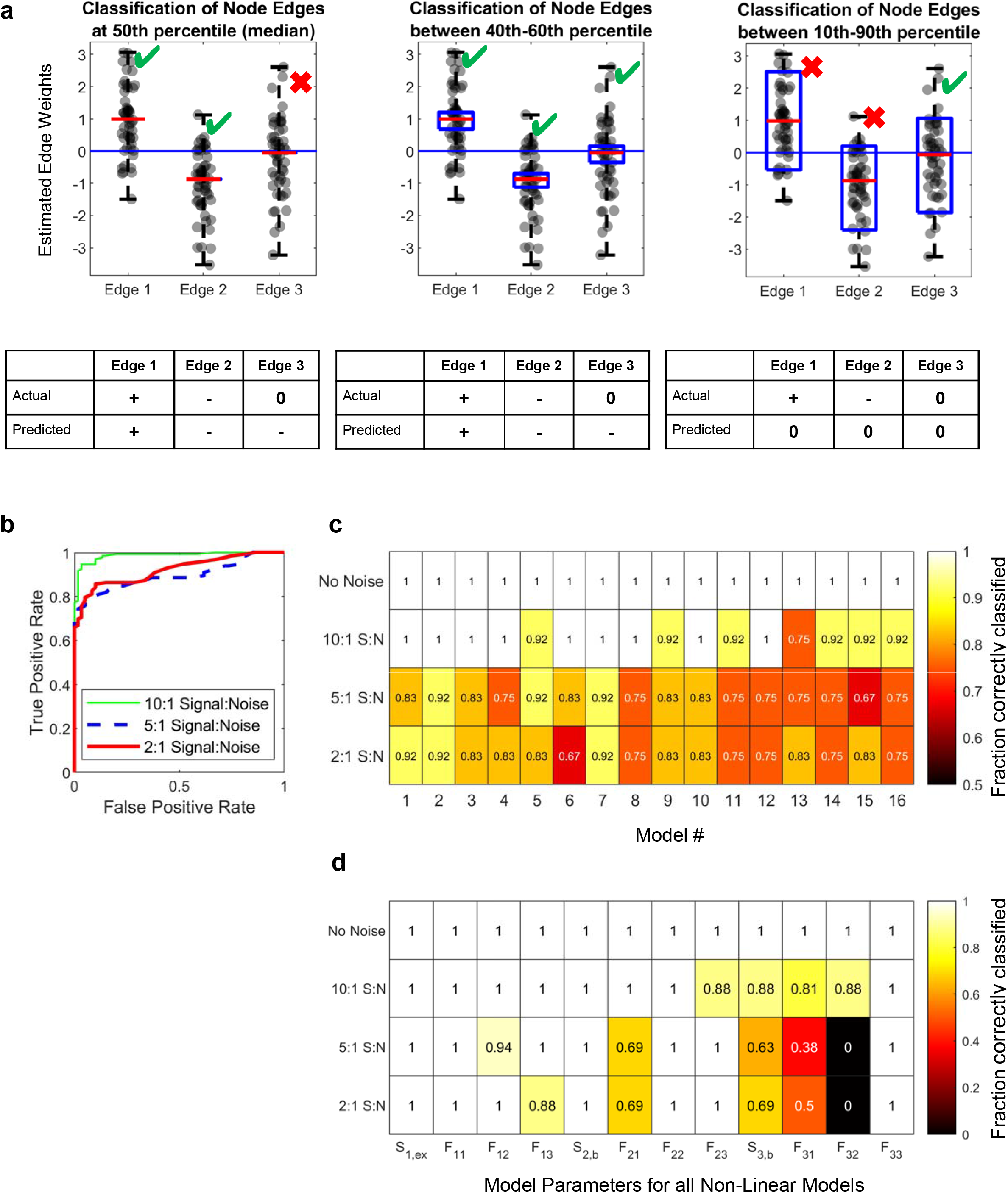
**(a)** Classification scheme for a distribution of parameter estimates. Going from left to right panels, the same parameter distribution with an actual (ground truth) value of positive (+), negative (-), or null (0), respectively, is estimated using different percentile windows centered on the median. The percentile “window” is the median value for the leftmost panel (rigorous classification), between 40th and 60^th^ percentile in the second panel, and between 10^th^ and 90^th^ percentile in the third panel (conservative classification). Going from rigorous to conservative (left to right), an intermediate between the two gives a good classification performance. **(b)** ROC curves across all parameters for all 16 FFL models. Different color lines are different noise levels. **(c)** Fraction of correctly classified model parameters for different noise levels broken down by FFL model type. **(d)** Fraction of each model parameter correctly classified for different noise levels broken down by parameter type.

Now, we applied this classification strategy to the 16 FFL model estimates from data with different noise levels. We varied the percentile window from the median only (50) to the entire range of estimated values (100) and calculated the true and false positive rates for all edges across all 16 FFL models, which allowed generation of receiver operator characteristic (ROC) curves (Fig. 6b). For each noise level, we chose the percentile window that yielded a 5% false positive rate (13-87 percentile for 10:1 Signal:Noise, 19-81 percentile for 5:1 Signal:Noise, and 21-79 percentile for 2:1 Signal:Noise). Using this simple cutoff classifier, we observed good classification performance across all noise levels according to traditional area under the ROC curve metrics (10:1 AUC=0.99, 5:1 AUC=0.9, 2:1 AUC=0.92).

How does classification accuracy break down by FFL model and edge type? To evaluate the performance for each of the 16 FFL cases, we calculated the fraction of the 12 links in each FFL model that was classified correctly as a function of signal-to-noise, given the percentile windows determined above (Fig. 6c). We also looked at the fraction of the 16 models where each of the 12 links were correctly classified (Fig. 6d). Perfect classification is a value of one, which is the case for no noise, and for many cases with 10:1 signal-to-noise.

In general, as noise level increases, prediction accuracy decreases, as expected. Although for some models and parameters, performance at 2:1 signal-to-noise is poor, in some cases it is surprisingly good. This suggests that the proposed method can yield information even in high noise cases; this information is particularly impactful for null, self-regulatory, and stimulus edges. High noise has strong effects on inference of edges that are either distinct across models, time variant or reliant on other node activities (*F_21_, F_31_, F_32_*) (Fig. 6c-d, S7). *F_21_,* which is reliant on activity of *x_1_*, is inferred better than *F_31_* and *F_32_*. This may be caused by the fact that *x_3_* dynamics depend on both *x_1_* and *x_2_,* whereas *x_2_* dynamics only depend on *x_1_.*

Comparing across models, we find that Models 1-8 are reconstructed slightly better than Models 9-16 (Fig. 6c) when noise is high. This performance gap is predominantly caused by *S_3,b_* misclassification—basal production of Node 3 (Fig. S7). What is the reason for the possible misclassification of *S_3,b_* in Models 9-16? We know that *S_3,b_* depends on the initial values of *x_1_, x_2_* and *x_3_* and the estimated values of *F_31_, F_32_* and *F_33_* (See Methods, Eq. 19). For Models 1-8, *x_1_(t=0)* and *x_2_(t=0)* are both zero and therefore *S_3,b_* is effectively only dependent on estimated value of *F_33_* and *x_3_(t=0)* (Fig. S6 and Methods). But for Models 9-16, *x_2_(t=0)* is non-zero and *S_3,b_* is dependent on the estimated values of both *F_32_* and *F_33_,* in addition to *x_2_(t=0)* and *x_3_(t=0)*, which increases the variability of *S_3,b_* estimates. Therefore with high levels of noise, *S_3,b_* is more likely to be mis-classified in Models 9-16, whereas this does not happen in Models 1-8 (Fig.6c,d, S7). In the future, including stimulus and basal production parameters in the least squares estimations themselves, rather than further deriving algebraic relations to estimate them, will likely help improve reliability.

We conclude that (i) when dealing with non-linear gene regulatory networks, complete perturbations such as genetic knockouts may fundamentally impede one’s ability to deduce network architecture and (ii) this class of non-linear networks can be reconstructed with reasonable performance using the proposed strategy employing partial perturbations.

## Discussion

Despite intensive research focus on network reconstruction, there is still room to improve discrimination between direct and indirect edges (towards causality), particularly when biologically-ubiquitous feedback and feedforward cycles are present that stymie many statistical or correlation-based methods, and given that experimental noise is inevitable. The presented DL-MRA method prescribes a realistic experimental design for inference of signed, directed edges when typical levels of noise are present. It allows estimation of self-regulation edges as well as those for basal production and external stimuli. For 2 and 3 node networks, the method can successfully handle random linear networks, cell state transition networks, and gene regulatory networks, and, under certain limiting conditions, signaling networks. Prediction accuracy improved with more timepoints, which in our case accounted for more relevant dynamic data. However, we would like to stress that here we did not explore time point placement, which likely underlies the performance increase rather than simply number of timepoints. Prediction accuracy was strong in many cases even with simulated noise that exceeds typical experimental variability (2:1 signal-to-noise). The method presented here is quite general and could be applied not only to cell and molecular biology, but also vastly different fields where perturbation time course experiments are possible, and where network structures are important to determine.

MRA and its subsequent methods allow for inference of direct edges by prescribing systematic perturbation of each node (Andrec et al., 2005b; Halasz et al., 2016a; Kholodenko et al., 2002b, 2002b; Santra et al., 2013a; Thomaseth et al., 2018) and the idea of directionality has been followed through in DL-MRA. Often, such edge directness is equated to causality, but this is not necessarily the case, especially when the entire system is not explicitly represented. In practice, the causality and strength of an edge may be dependent on how well the model represents the underlying phenomenon and might be affected by simplification of larger networks, non-linearities in the actual model and even by noise in the data. Secondly, in discussions about causal system inferences, consideration of the counterfactuals is important (Höfler, 2005; Morgan and Winship, 2014; Pearl, 2013; Shipley, 2016). For a network of nodes going through dynamics, the counterfactuals to intrinsic network edges causing the dynamics would be the environmental factors extrinsic to the network edges. In DL-MRA, by evaluating external stimuli and basal production as well as the network edges, we have mapped some counterfactuals to node dynamics, thus presenting a more complete map of the causal factors to the network dynamics compared to methods which only show network edges. This also allows for a concise mapping of the environmental contexts in which the network edges are reconstructed.

Application of DL-MRA could reconstruct cell state transition networks based on discrete time Markov transition models, with the added benefit of not being constrained to specific time intervals. It can also successfully handle noisy data. The additional constraints in DL-MRA in the context of cell state transitions (summations of transition rates—see Methods) implies that the underlying network may be estimated even with less data requirements than in other cases. This method can be a useful tool to model cell state transitions and predict cell state. Perturbations were modeled as a difference in initial states, and that worked well in this case, suggesting that such modeling of perturbations may work in other cell state transition or biological networks.

Although application of DL-MRA to an intracellular signaling network (Huang-Ferrell MAPK) was able to explain its ground truth, including feedback due to sequestration, the method was constrained to specific, difficult-to-implement perturbations and a low stimulus which may not always be feasible experimentally. In MRA, a larger reaction scheme is often simplified into modules with one species in the module representing the activity of the module. But often, the activity of the other species in the module is implicit and becomes significant in dictating how perturbations and stimulus affect the network dynamics. Moreover, the type of perturbation chosen also may yield different network inference results. Therefore, the use of MRA methods on simplified large intracellular signaling networks, especially while dealing with experiments, have significant caveats that should be carefully considered.

(Fuente et al., 2002)(Thomaseth et al., 2018)Although complete inhibition is often used for perturbation studies of gene regulatory networks (e.g. CRISPR-mediated gene knockout), we found that partial inhibition is important to fully reconstruct the considered non-linear gene regulatory networks. It is important to distinguish here, however, small perturbations vs. partial perturbations. Small perturbations are formally recommended for both MRA and other techniques (Fuente et al., 2002) where the effects of noise are not extensively explored. In practice however, there is a tradeoff between perturbation strength and feasibility, since the effects of small perturbations are masked by noise (Thomaseth et al., 2018). Partial perturbations, as considered in this work (~50%) are much larger than what are typically considered small perturbations. The theoretical formulation of DL-MRA reduces the impact of not having small perturbations, because perturbation data from a particular node is not used for inference of edges connected to that node. Yet, DL-MRA still uses linearizations of the Jacobian which are are always subject to greater inaccuracy the further away from reference points such perturbations take the system. Since many biological networks share the same types of non-linear features contained within the considered FFL models, this is not likely to be the only case when partial inhibition will be important. We are thus inclined to speculate that large partial perturbations may be a generally important experimental design criterion moving forward. Partial inhibition is often “built-in” to certain assay types, such as si/shRNA or pharmacological inhibition that are titratable to a certain extent.

One major remaining challenge is scaling to larger networks. Here, we limited our analysis to 2 and 3 node networks. Conveniently, the number of necessary perturbation time courses needed grows linearly (as opposed to exponentially) with the number of considered nodes. Furthermore, as long as system-wide or omics-scale assays are available, the experimental workload also grows linearly. This is routine for transcriptome analyses (Stark et al., 2019), and is becoming even more commonplace for proteomic assays (e.g. mass cytometry (Spitzer and Nolan, 2016), cyclic immunofluorescence (Lin et al., 2016a), mass spectrometry (Aksenov et al., 2017), RPPA (Akbani et al., 2014)) (Lin et al., 2016a). Thus, the method is arguably experimentally scalable to larger networks.

However, the computational scaling past 2 and 3 node models remains to be determined and is likely to require different approaches for parameter estimation. Increasing the network size will quadratically increase the number of unknown parameters. Reducing this parameter space and obtaining good initial guesses will be important. Imposing prior knowledge can also reduce the parametric space, such as in Bayesian Modular Response Analysis (Santra et al., 2013a), or with functional database information (Wu et al., 2010). As network size grows, the sparseness of the Jacobian will increase, so judicious allocation of non-zero elements will be important. Checking estimated Jacobians for emergent properties such as degree distributions for scale-free networks (Barabási and Albert, 1999a) can provide additional important constraints. The approach used in this paper is accommodative of such prior knowledge and in principle can be scaled up for larger network size. Lastly, large estimation problems may be broken into several smaller problems to be merged subsequently, which is likely to yield large computational speed up by allowing parallelization of much smaller tasks.

In conclusion, the proposed approach to network reconstruction is systematic and feasible, robustly operating in the presence of experimental noise and accepting data from large perturbations. It addresses important features of biological networks that current methods struggle to account for: causality/directionality/sign, cycles (including self-regulation), dynamic behavior and environmental stimuli. It does so while leveraging dynamic data of the network and only requires one perturbation per node for completeness. We expect this approach to be broadly useful not only for reconstruction of biological networks, but to enable using such networks to build more predictive models of disease and response to treatment, and more broadly, to other fields where such networks are important for system behavior.

## Acknowledgments

We would like to thank Clemson University and the CCIT team for the generous allotment of time and support in the Palmetto cluster for running the simulations in this paper.

## Data and Software

The code needed to reproduce the data and figures are included. Parallelization when necessary to generate data was run on Palmetto cluster (372 GB, 48 nodes) and MATLAB 2020a.

## Author Contributions

MRB, GRS, and DS conceived of the work. DS, GRS, MB, MRB, and JE performed analyses. DS, MB, and GRS made the figures. DS, GRS, MB, and MRB wrote the manuscript.

## Supplementary Figure Legends

**Figure S1:**
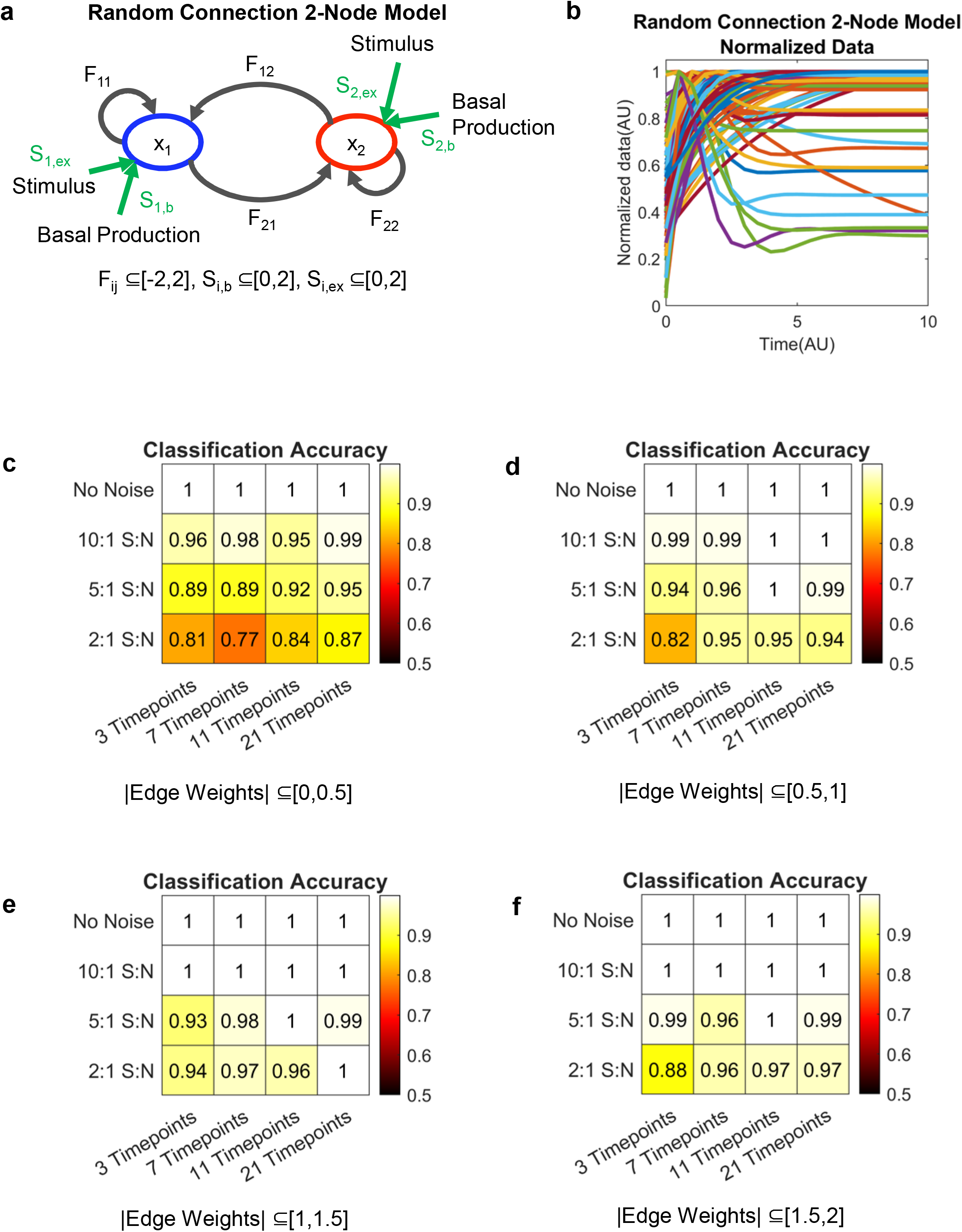
**(a)** Random 2 node network with Jacobian elements labeled. Green arrows are basal production and external stimulus terms. **(b)** Time courses for the 50 random 2 node networks, normalized by the maximum value **(c,d,e,f)** Fraction of network parameters correctly classified in 50 randomly generated 2 node networks with an absolute value between 0 to 0.5 (c), 0.5 to 1 (d), 1 to 1.5 (e) and 1.5 to 2 (f).

**Figure S2:**
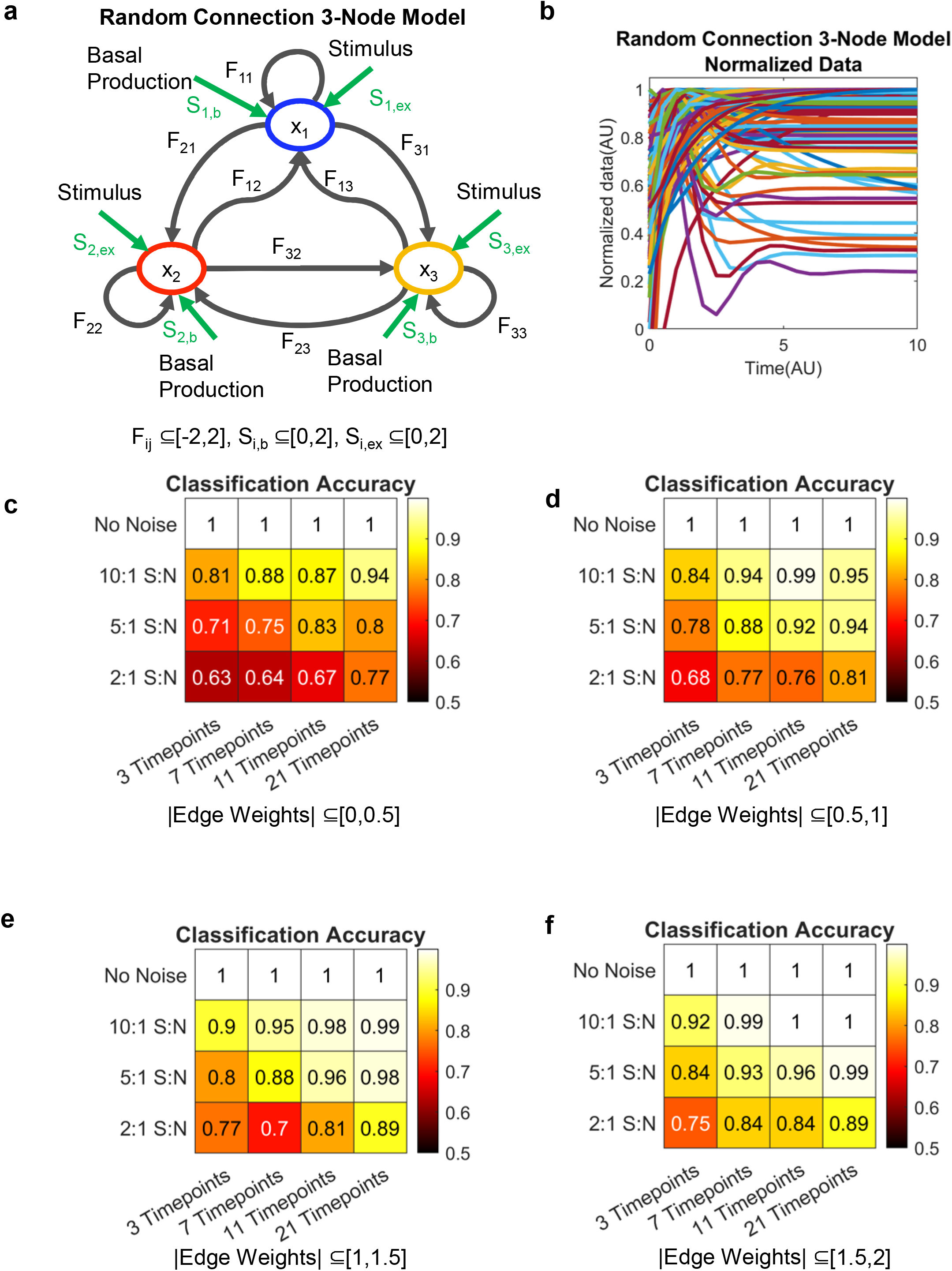
**(a)** Random 3 node network with Jacobian elements labeled. Green arrows are basal production and external stimulus terms. **(b)** Time courses for 50 random 3 node networks, normalized by the maximum value **(c,d,e,f)** Fraction of network parameters correctly classified in 50 randomly generated 3 node networks with an absolute value between 0 to 0.5 (c), 0.5 to 1 (d), 1 to 1.5 (e) and 1.5 to 2 (f).

**Figure S3:**
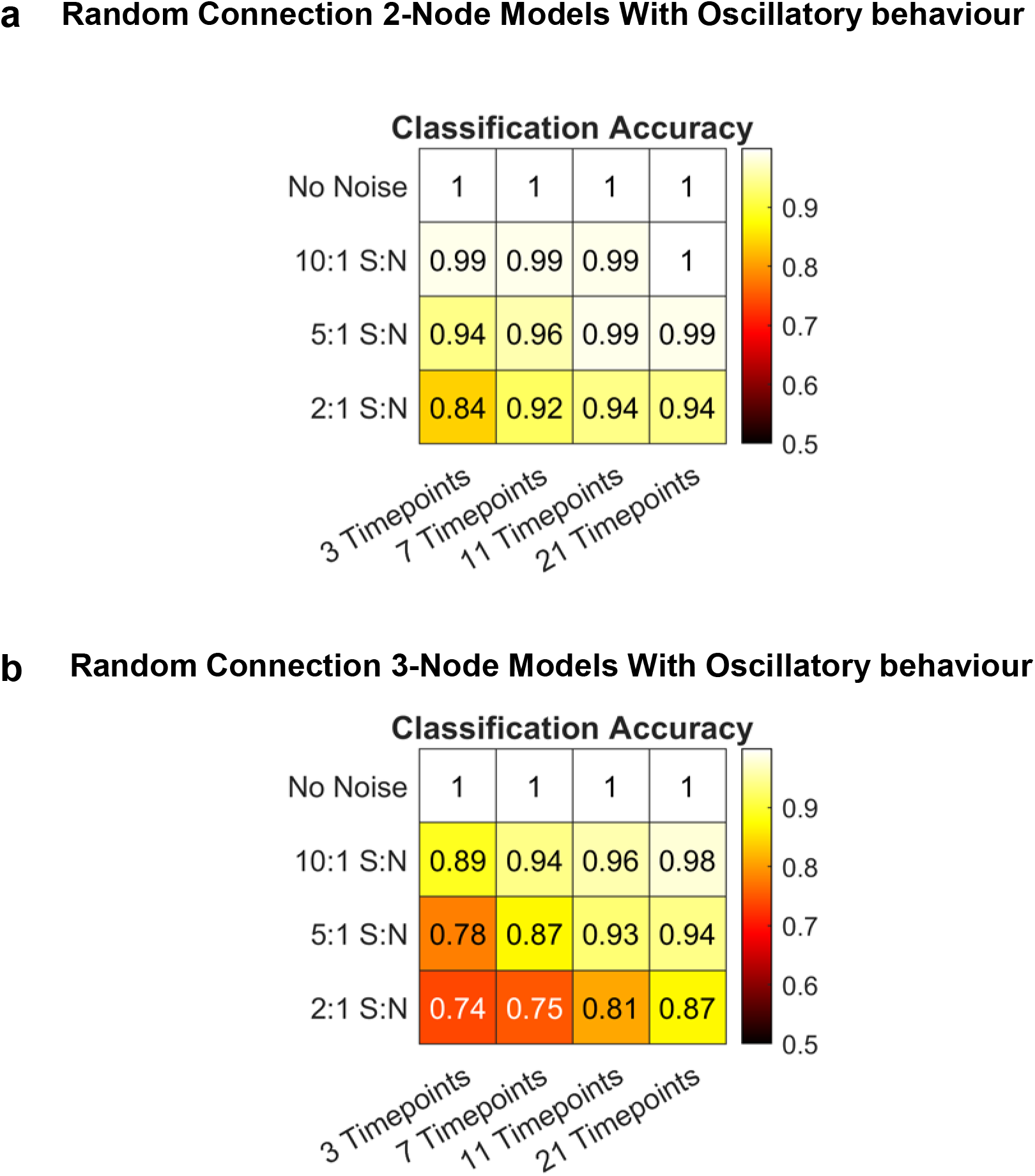
**(a,b)** Fraction of network parameters correctly classified in 29 two node networks (a) and 29 three node networks (b) with potential for oscillatory behavior (non-zero imaginary parts of eigenvalues of Jacobian elements).

**Figure S4:**
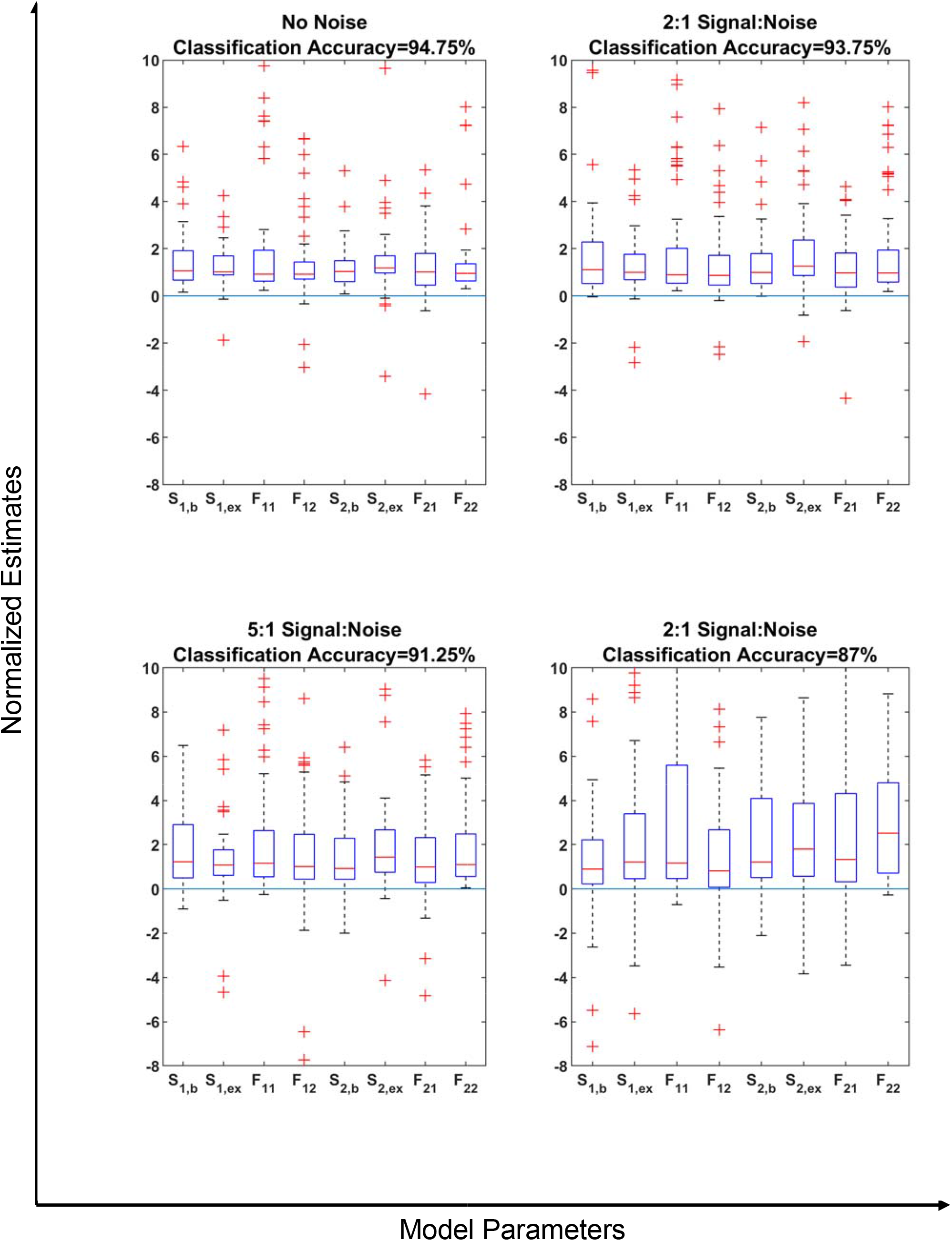
Distribution of estimated parameters in two node networks normalized to corresponding actual parameters in 50 random three node systems, when the data from only node 1 and node 2 is included to make the estimation. A value of 1 means the parameters estimate did not change.

**Figure S5:**
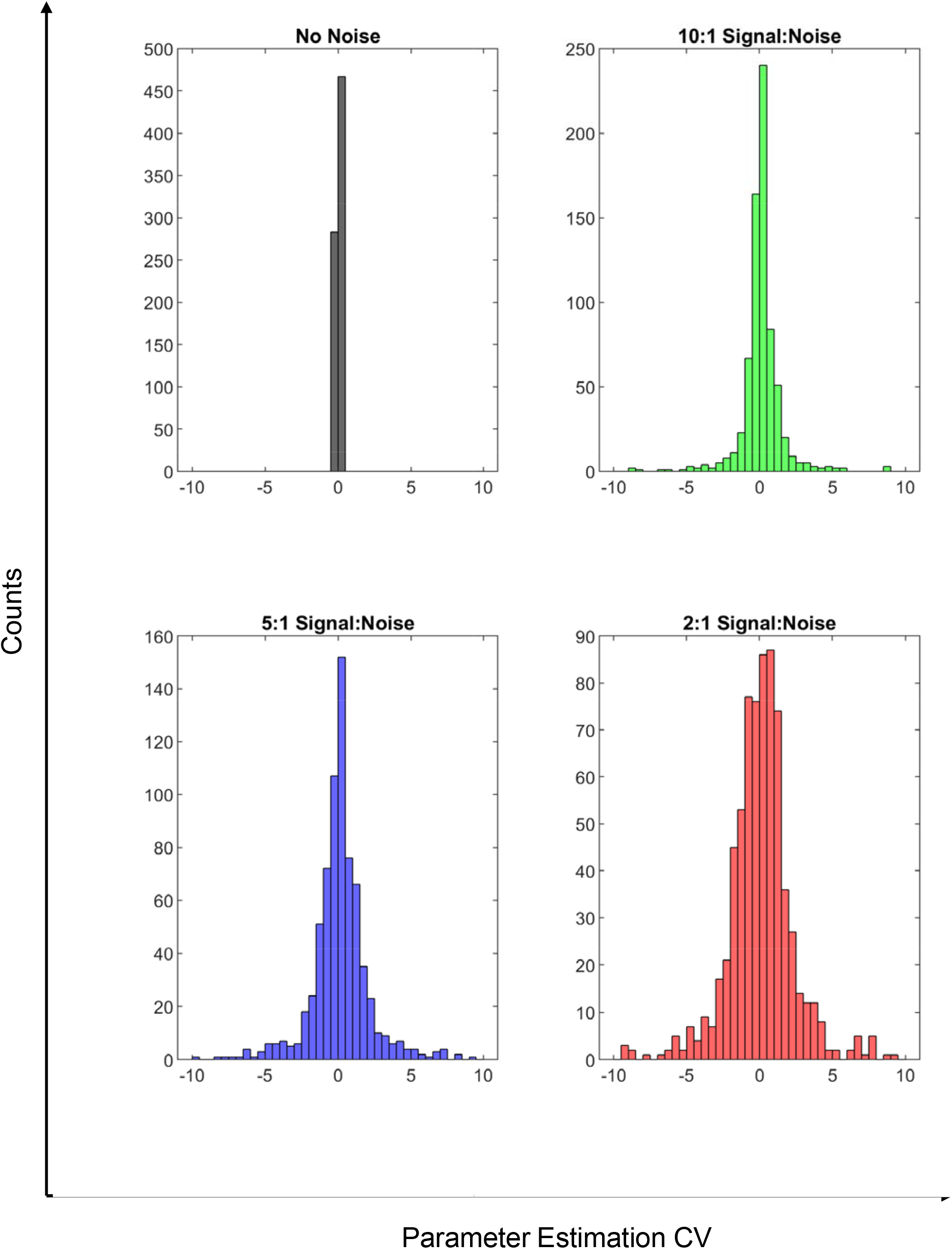
Histogram of coefficient of variation (CV) among the parameters from multi start results in the 50 random three node models. Only parameter sets with sum of squared errors (SSE) less than twice the minimum SSE were included as acceptable.

**Figure S6:**
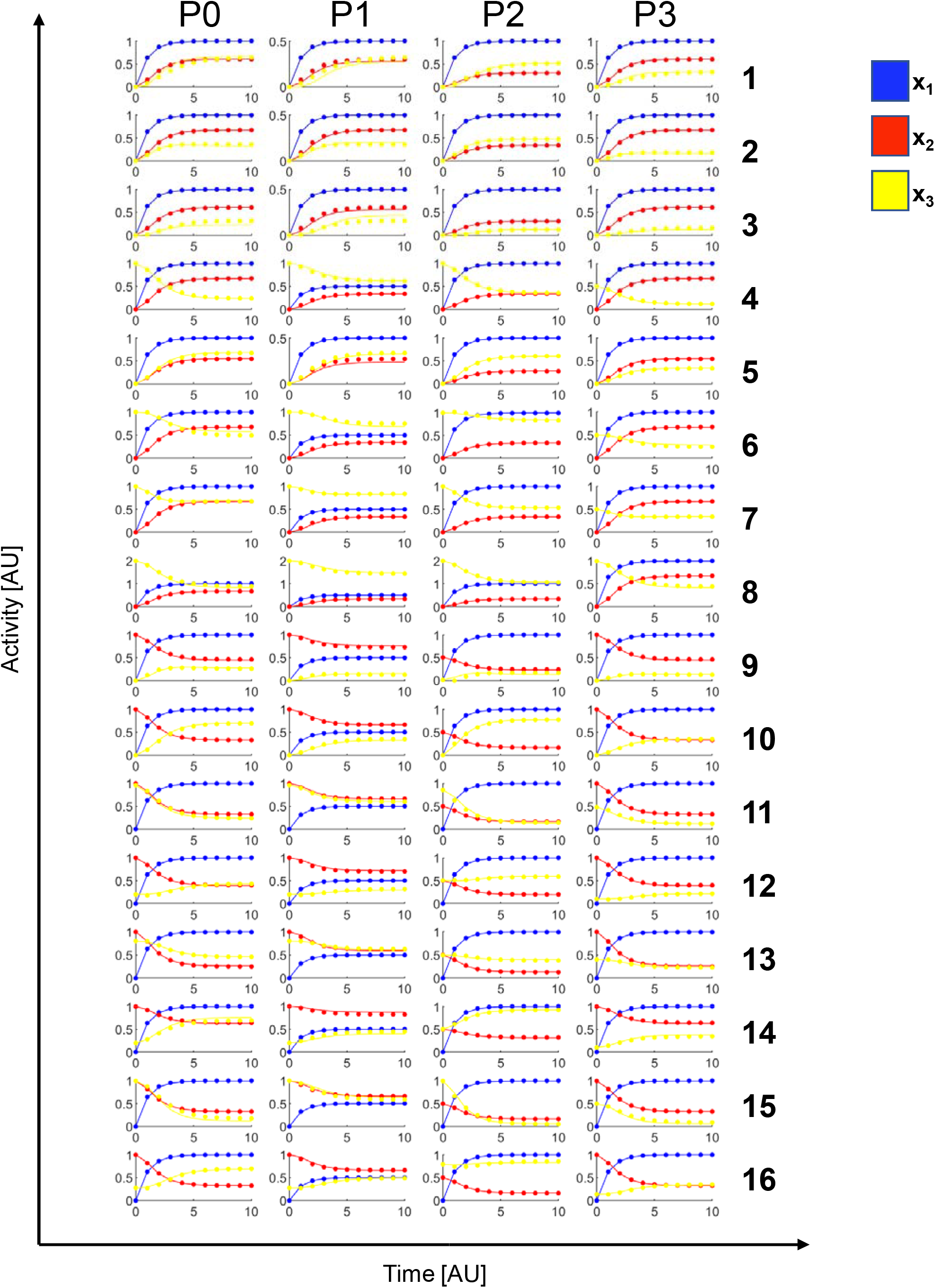
Simulated, noise-free experimental data (dots) and model-generated fits (lines) for each FFL model structure. Different perturbations—vehicle (P0), perturb x1 (P1), perturb x2 (P2), perturb x3 (P3)—are across the columns and different model structures (1-16) are down the rows. Each node (1-3) is a different color as indicated in the legend.

**Figure S7:**
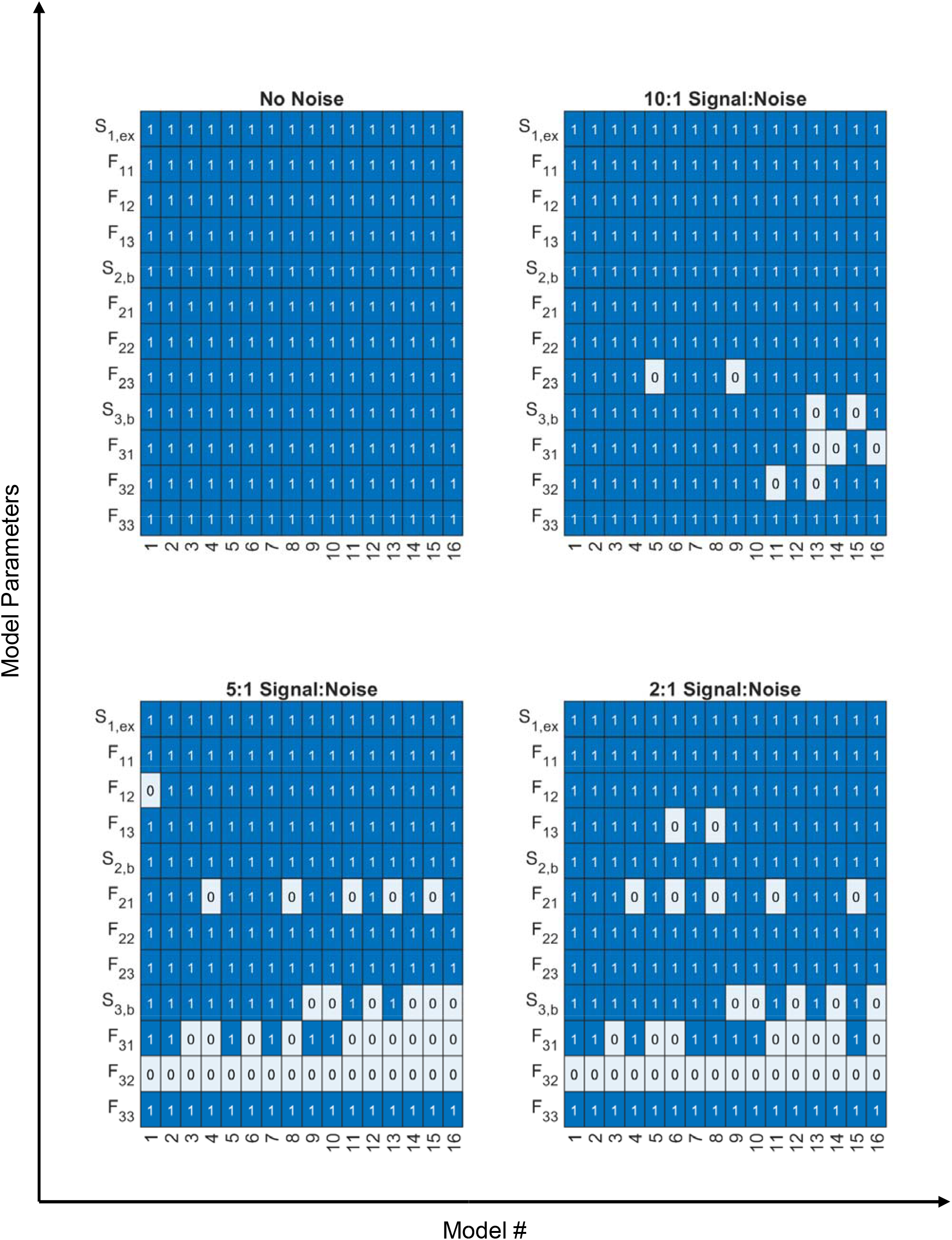
Detailed results from the FFL models depicting whether a model parameter was correctly (1) or incorrectly (0) predicted for each model structure (1-16) under each noise level. Correct predictions were classified based on the optimal percentile cutoffs identified for each noise level.

## Methods

### Deriving Sufficiency Conditions for Unique Estimation of Jacobian Elements

The first-order partial derivatives comprising **J** (Eq. 2) can be approximated by a first-order Taylor series expansion of Eq. 1 about a time point *k*

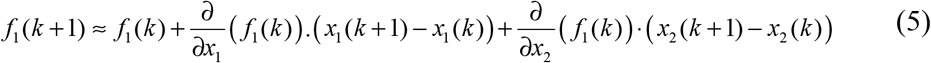

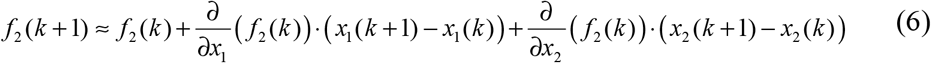

Eq. 5-6 may be written more succinctly as

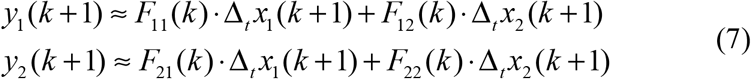

where

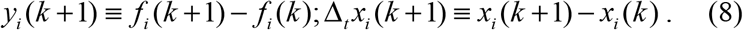

The approximation in Eq. 7 becomes more accurate as more time points are measured. Also, the edge weights are potentially time-dependent, although this is rarely considered when describing biological networks.

How do we estimate the edge weights *F* in Eq. 7 and thus reconstruct the network? Time series data can inform *x_i_*’s and *f_i_*’s as a function of time, following application of a stimulus. Given such stimulus-response data, however, for each time point there are only two equations for four unknowns, an underdetermined system for which more data are needed.

Consider now stimulus-response time course data in the presence of single perturbations. Let *p_i_* be a variable that reflects the strength and/or presence of different potential perturbations: *p_1_* represents perturbation of x_1_ and *p_2_* represents perturbation of x_2_. If *p_j_* is not explicitly written, its value is zero and/or it has no effect. Now, the ODEs become a function of the perturbation variables

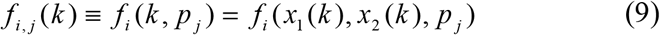

The 1^st^ order Taylor series expansions for cases with perturbations become

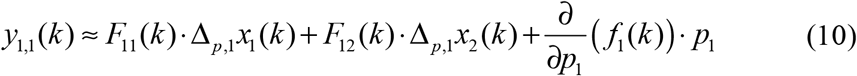

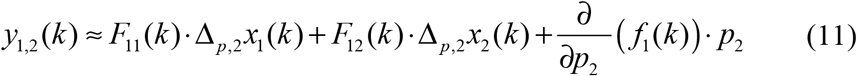

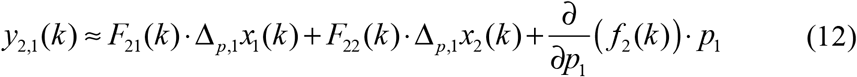

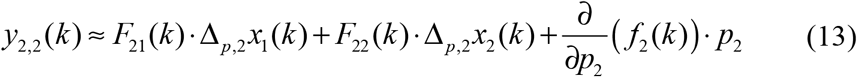

where

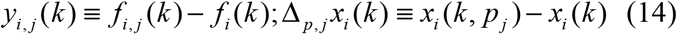

Here, we have expanded with respect to the perturbation, rather than with respect to time as previously. However, since the reference point is the same, the Jacobian elements remain identical in these equations. It is also interesting to note that the Jacobian elements, or network, may be affected by the perturbation, but we do not necessarily have to know those effects mathematically, since the reference point is the same. Now we have six potential equations with which to estimate the four Jacobian elements. However, we must make some determination as to how the perturbations *p_1_* and *p_2_* directly affect Node 1 and Node 2 dynamics *f_1_* and *f_2_* to account for the perturbation variable partial derivatives.

By design, the Node 1 perturbation has significant *direct* effects on Node 1 dynamics, and similarly for the Node 2 perturbation on Node 2 dynamics. Using equations including ∂*f*_1_/∂*p*_1_. and ∂*f*_2_/∂*p*_2_ require precise definition of perturbation strength and their effects on dynamics, which could be difficult to determine experimentally and implement in simulations. Therefore, we do not employ equations involving such terms. On the other hand, if the Node 1 perturbation has negligible *direct* effect on Node 2 dynamics, that is, the effects on Node 2 dynamics are through the network (i.e. *p_1_*) and not explicit in *f_2_*), and similarly the Node 2 perturbation has negligible *direct* effect on Node 1 dynamics, then ∂*f*_2_/∂*p*_1_ and ∂*f*_1_/∂*p*_2_ are approximately zero. This mild condition is often the case experimentally. The only determining factors for the suitability of the Taylor series truncation are the spacing of time points and the accuracy of the expansion about the perturbation difference. From this, the main set of linear equations presented in Eq. 3-4 are obtained.

### General Estimation Model Equations

We employ the following general model for a two-node network: -

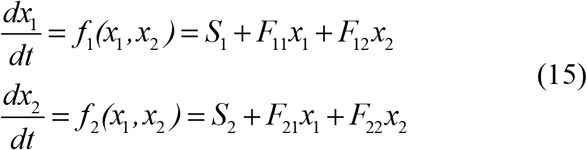

Here, *S_1_* and *S_2_* are the stimuli strengths on Node 1 and Node 2 respectively, and *F_11_, F_12_, F_21_* and *F_22_* are the network edge weights (Figure 1a). In many systems, there may be a basal or constitutive production driving the node activities, besides an external stimulus. For these cases, the Stimulus term (*S_i_*), may be considered as an addition of these two effects- the basal production term (*S_i,b_*) and the external stimulus (*S_i,ex_*). Then the two-node model can be represented by the following equations-

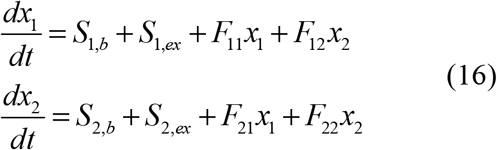

Or more generally,

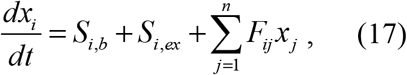

where *n* is the total number of nodes.

When a steady state exists, the *dx_i_/dt* terms become zero and it becomes easy to represent the stimulus terms as a function of the node activities (*x_i_*) and network edges (*F_ij_*).

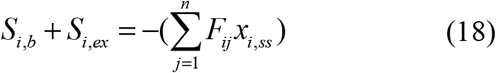

This is helpful to understand that the perturbation time course data also generally constrains not only the edge weights, but also the stimulus terms. For a system at a steady state without an external stimulus, for example at t=0:

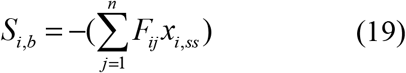

### The Two-node Single Activator model

The two-node single activator model (Fig. 1a, S1a) is described by

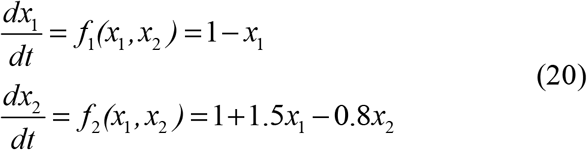

Here, *S_1,ex_=1, F_11_=-1, F_12_=0, S_2,ex_=1, F_21_=1.5, F_22_=-0.8*. The basal production terms are both zero, for simplicity, and the initial conditions for *x_1_(t=0)* and *x_2_(t=0)* are zero. The stimulus terms *S_i,ex_* are calculated through Eq. 18, using the median values of *F_ij_* and the *x_i_(t=10)*, when the system reaches near steady state.

### Random Two-node and Three-node models

The random 2 node network is described by

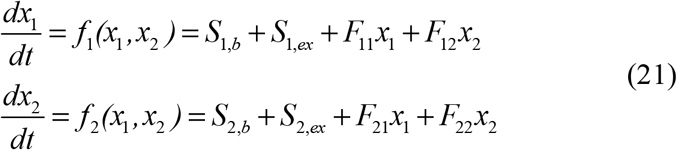

Values for *S_1,b_, S_2,b_, S_1,ex_* and *S_2,ex_* are sampled from a uniform distribution over the range [0,2] and values for F_11_, F_12_, F_21_, and F_22_ are sampled from a uniform distribution over the range [-2,2] using the MATLAB function rand. To capture basal activity, we use a two-step approach. First, starting from node activity values of zero, without the external stimulus on Node 1 and Node 2 (*S_1,ex_=S_2,ex_=0* in Eq 22) we simulate until the network reaches steady-state with just basal production driving the network behavior. Then, we introduce the external stimulus on Node 1 and Node 2, integrate the ODEs, and sample evenly spaced time-points using `ode15s` in MATLAB with default settings. We sample 3,7, 11, and 21 evenly spaced time points across a time course, from 0 to 10 arbitrary time units in all the cases.

The random 3 node networks use the same sampling rules as the 2 node networks with the following equations.

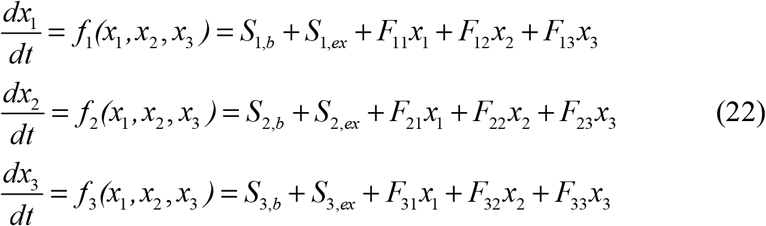

### Intracellular Signaling Networks

In the simplification of the Huang-Ferrell network to three nodes, p-MAPKKK, pp-MAPKK and pp-MAPK were taken as nodes. Since, in absence of external stimuli, the basal values of the nodes are zero, the basal production was estimated as zero beforehand and not considered in the estimation of the rest of the network. Aside from the basal production edges, a full 3 node network (Fig 4b) was estimated from the simulation data of each of the observables. After estimation, parameters with values less than 1/100th of the largest parameter, were considered negligible.

### Cell State Transition Models

The cell transition model from (Gupta et al., 2011) is a discrete time Markov probability model. Here, we show how this form is related to the ODE model used in DL-MRA. Starting at any initial value, each next step representing a time difference of one day follows from the previous time point as follows-

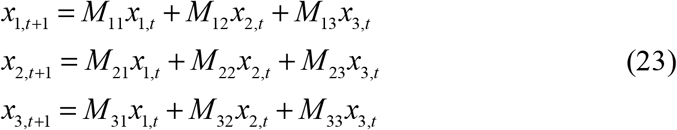

Where *M_ij_* denotes the Markov transition probabilities of species *j* into species *i*. In matrix form it may be represented as follows-

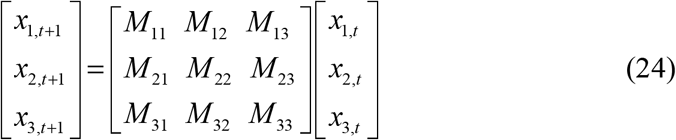

Representing the Markov parameter matrix as *M* and the species relative concentration variables as vector X, the equation becomes

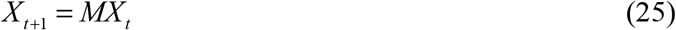

The Markov transition probabilities for a species must add up to 1. In experimental terms, a species can either transition to other species or stay the same and the sum of all those probabilities is 1.

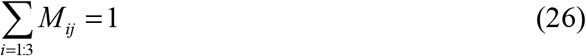

As a first step in relating these equations to the ODE form underlying DL-MRA, we put the variables in terms in terms of Δx (with respect to time),

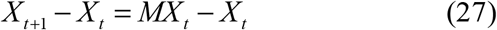

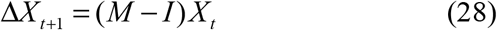

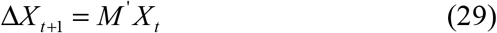

Where ***M’*** is ***M-I***, and ***I*** is the identity matrix. ***M’*** is ***M***, except that 1 is subtracted from all its diagonal elements. Hence Eq. 26 for *M’* becomes

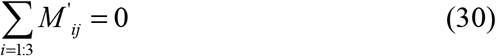

This also implies that the diagonal term for ***M’*** is negative of the sum of the other two terms in the same column. In experimental terms, the amount of reduction of a species is equal to how much it got converted to other species.

The above equations apply for the cases where Δt is 1. We can incorporate arbitrary time steps as

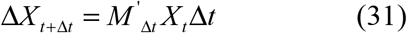

Where Δt is the scalar value of time difference and *M’*_Δ*t*_ is the matrix of the set of parameters, specific to the time difference chosen. For a case where Δt tends to 0, the equation becomes-

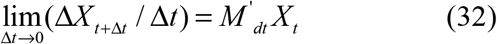

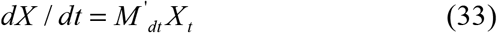

Where *M’_dt_* is the matrix of the set of parameters specific to the case where Δ*t* is infinitesimally small. Note that Eq. 33 is similar in form to Eq. 22, only without the extra stimulus terms and where *M’_dt_* is equivalent to the Jacobian matrix F with terms *F_ij_*. There would be an added constraint that the sum of the terms in the same column would add up to zero, or that the diagonal term is the negative of the sum of the other two terms in the same column.

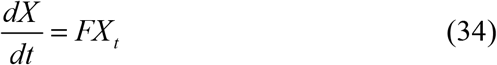

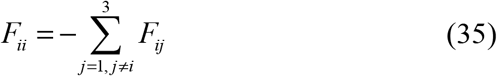

### Non-Linear Models

The non-linear feedforward loop models (Mangan and Alon, 2003a) are described by:

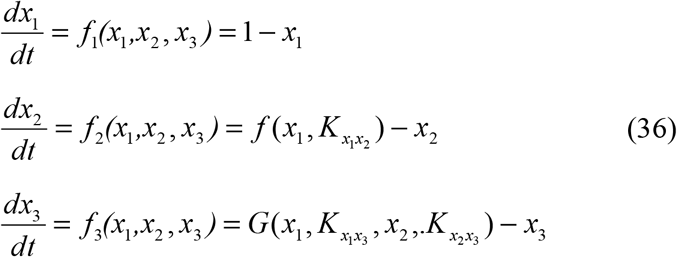

When an AND gate is present

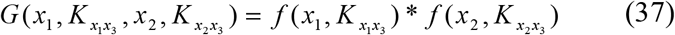

When an OR gate is present

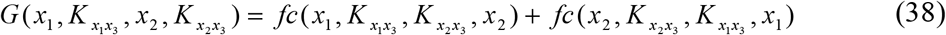

For a given u, v □ {x_1_, x_2_, x_3_} and K, K_u_, K_v_ □{*K*_*x*_1_*x*_2__, *K*_*x*_1_*x*_3__, *K*_*x*_2_*x*_3__ }:

If u activates its target, then:

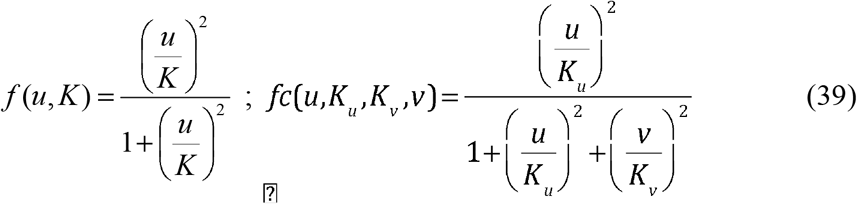

If u represses its target, then:

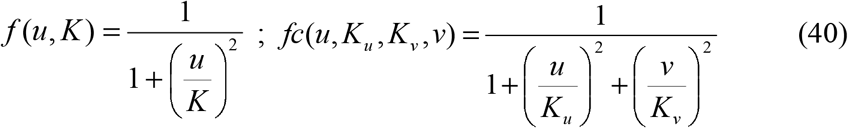

Effectively, an external stimulus of *S_1,ex_*=1’, acts on Node 1 at *t=0* and is propagated through the network. There is no external stimulus acting on Node 2 and Node 3. However, in many cases there is basal production in one or both of Node 2 and Node 3. This leads to a nonzero steady-state of the network before the external stimulus is introduced.

To capture basal activity, we use a two-step approach. First, starting from node activity values of zero, without the external stimulus on Node 1 (*S_1,ex_=0)*, we simulate until the network reaches steady-state. Then, we introduce the external stimulus on Node 1, integrate the ODEs, and sample 11 evenly spaced time-points using `ode15s` in MATLAB with default settings and steady-state node values without the external stimulus as the initial conditions. We chose 11 timepoints because it yields good classification accuracy for the above random 3 node model even in presence of noisy data. For each of the 16 non-linear models, the values of the parameters (*K, Ku, Kv*), were varied and chosen so that the resulting node activity data are responsive to the stimulus and perturbations (Fig. S6, See Supplementary Code for values).

### Modeling Perturbations

Precisely modeling perturbations can be a challenge, since experimentally, there may be several ways of causing a perturbation with different mechanisms such as siRNAs, competitive/non-competitive/uncompetitive inhibition, etc. It may be hard to quantify how much a perturbation is affecting a node, in terms of its dynamics (i.e. right-hand sides of the ODEs). Therefore, we employ the following approaches which circumvent the need to model how each perturbation mechanistically manifests in the ODEs during parameter estimation. There are two cases to consider: (i) when we have a perturbation of node *i* and we need to simulate node *i* dynamics; (ii) when we have a perturbation of node *i* and we need to simulate other node *j* dynamics. To illustrate the approach, we use the above-described 2 node model with an example of a Node 1 perturbation. Recall that

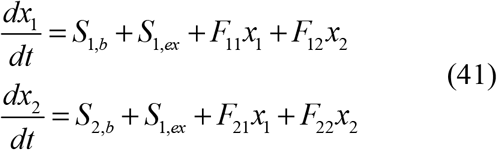

For case (i), we have to obtain values for *x_1_* under perturbation of Node 1. We refer to the perturbed time-course as *x_1,1_.* In experimental situations, *x_1,1_* would be measured directly. To obtain simulation data for *x_1,1_* we use the following:

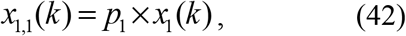

where *x_1_* is obtained from the simulations without perturbations, and recall that *k* refers to time point *k*. For a 50% inhibition, *p*=0.5 and for a complete inhibition, *p*=0.

For case (ii), we have to obtain the values for *x_2_* under perturbation of Node 1, which we refer to as *x_2,1_*. To do this, we have to integrate the ODE for *dx_2_/dt*, but using *x_1,1_* values, as follows

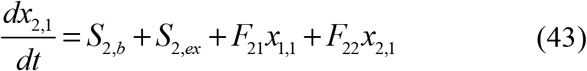

Here, *x_2_* has been replaced with *x_2,1_* to represent *x_2_* under perturbation of Node 1, for clarity. To solve this equation, we simply use the “measured” *x_1,1_* time course directly in the ODE.

When data are generated by simulations, there is little practical limit to temporal resolution, but with real data, to solve Eq. 43 one may need values for *x_1,1_* at multiple time points where measurements are not available, depending on the solver being used. We therefore fit *x_1,1_* data to a polynomial using polyfit in MATLAB, and use the polynomial to interpolate given a required time point. In this work, we have used an order of 5 to fit the data as well as avoid overfitting, but the functional form is quite malleable so long as it captures the data trends.

For modeling perturbations of the cell transition model, the initial value of the simulated data for the perturbed node was taken as zero during simulation. The estimation was performed in a similar way as a random 3 node network as described above.

For modeling perturbations for the Huang Ferrell model, the parameters k3, k15 and k27 were sequentially set as zero. The estimation was performed in a similar way as a random 3 node network as described above.

### Simulated Noise

Normally distributed white (zero mean) noise is added to simulated time courses pointwise with

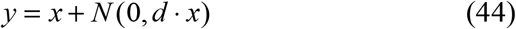

where *x* is the simulation data point, *y* is the noisy data point, and *d* represents the noise level. Signal-to-noise ratio of 10:1, 5:1 and 2:1 are, respectively *d* = 0.1, 0.2, and 0.5. Normally distributed samples are obtained using randn in MATLAB. While there are many different distributional options for modeling noise, we chose this for simplicity and to capture the effects generically of noisier data. We do not intend to answer questions related to whether specific distributional assumptions about the form of the noise have significant impact of the methods performance.

### Parameter Estimation

For the two-node model, the entire network, with and without perturbations, can be explained by the following system of equations

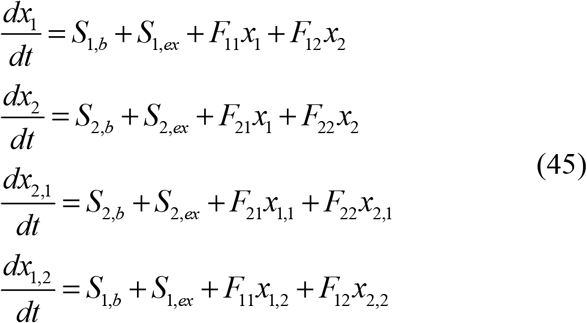

where *x_1,1_* and *x_2,2_* are the perturbed node values, from either simulated or experimental data. Eight parameters (*S_1,b_, S_1,ex_, F_11_, F_12_, S_2,b_, S_2,ex_, F_21_, F_22_*) need to be estimated to fully reconstruct this network. We seek a set of parameters that minimizes deviation between simulated and measured dynamics.

For an initial guess, the node edge parameters (*F_ij_*) are randomly sampled from a uniform distribution over the interval [-2,2] and the stimulus parameters (*S_i,ex_*) are sampled from a uniform distribution over the interval [0,2]. Using data at *t=0,* which corresponds to a steadystate without *S_i,ex_,* the *S_i,b_* can be estimated during each iteration of the estimation as follows-

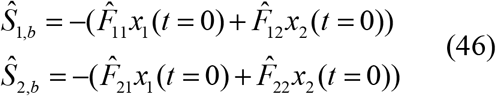

For an n-node model, this equation can be scaled accordingly to obtain each *Ŝ_i,b_.*

For these initial guesses we compute the activity data using Eq. 45. The perturbation data *x_k,k_* is used in the perturbation equations as detailed above (Eq. 43). Let 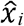, and 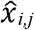 denote the predicted node activity values for non-perturbed and perturbed cases respectively. For a total of *n* nodes and *N_t_* timepoints, the objective function is the sum of squared errors Φ

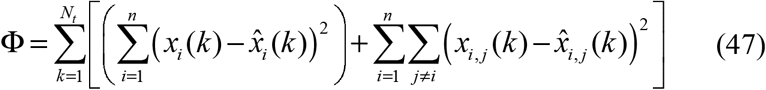

Note here that we do not use data from node *j*, when perturbation *j* was used (per the derivation). The MATLAB function fmincon is used to minimize Φ by changing edge weights and stimulus terms within the range [-10,10].

We employ “multi-start” by running the estimation 10 times, starting from different randomly generated initial starting points (Raue et al., 2013). The estimated parameters and their respective final sum of squared errors (Φ) are saved and the estimated parameter set corresponding to the minimum Φ is chosen as the final parameter set. Variability of parameter estimates across multi-start runs is explored in Supplementary Figure S5.

### Parameter Estimation for Non-Linear Models

For estimating the Non-Linear models, we start with a prior knowledge that *S_1,b_* is always *zero* and *S_2,ex_* and *S_3,ex_* are always zero as well, which is directly evident from *x_1_* initial conditions and *x_2_, x_3_* stimulus response in the presence of a complete Node 1 perturbation. The equations for the non-perturbation case become as follows

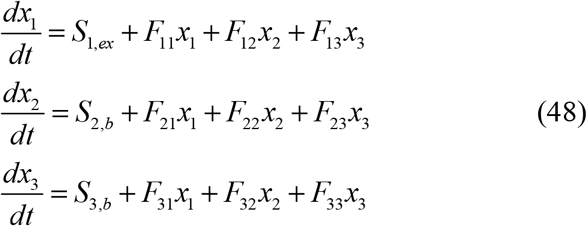

Since the system is at steady-state before the external stimulus, the basal production parameter can be estimated during each iteration of the estimation as-

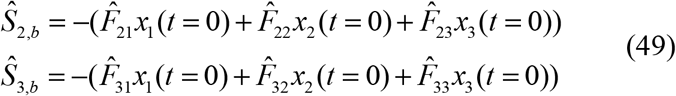

where 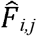 are the current model parameter estimates and *x_i_(t=0)* are the *x* values at the initial system steady state before the induction of external stimulus.

### Bootstrapping Simulated Data for the FFL Model Cases

To generate multiple parameter set estimates to classify edge weights for the FFL model cases, we employ a bootstrapping approach. In an experimental scenario, each data point will have a mean and a standard deviation, and upon a distributional assumption (e.g. normal), one can then resample datasets to obtain measures of estimation uncertainty. We use the simulated data as the mean, and then vary the standard deviation as described above to generate 50 bootstrapped datasets for each of the 16 considered models. Estimation is carried out for each of the 50 datasets using multi-start, which yields 50 best-fitting parameter sets for each model. Uncertainty analysis and classification error is based on these sets.

