## Supplementary Code for "Network Inference from Perturbation Time Course Data": Analytic2node.m

```
function fmin=Analytic2node(data,timeint)

Y0=data.Y;
YP1=data.YP1;
YP2=data.YP2;
T=data.T;
smoothmethod='moving';
span=1;

Y0smooth = zeros(length(T),2);
YP1smooth = zeros(length(T),2);
YP2smooth = zeros(length(T),2);
Y0smoothy = zeros(length(T),2);
YP1smoothy = zeros(length(T),2);
YP2smoothy = zeros(length(T),2);

% %Smooth the data 
% for j = 1:2
%    Y0smoothy(:,j) = smooth(Y(:,j),span,smoothmethod);    
% end
% 
% for j = 1:2
%    Y0smooth(:,j) = smooth(Y0smoothy(:,j),span,smoothmethod);    
% end
% 
% for j = 1:2
%    YP1smoothy(:,j) = smooth(YP1(:,j),span,smoothmethod);    
% end
% 
% for j = 1:2
%    YP1smooth(:,j) = smooth(YP1smoothy(:,j),span,smoothmethod);    
% end
% 
% for j = 1:2
%    YP2smoothy(:,j) = smooth(YP2(:,j),span,smoothmethod);    
% end
% 
% for j = 1:2
%    YP2smooth(:,j) = smooth(YP2smoothy(:,j),span,smoothmethod);    
% end

f1 = zeros(length(T)-2,1);
f2 = zeros(length(T)-2,1);
f1p = zeros(length(T)-2,1);
f2p = zeros(length(T)-2,1);

obstwojacdata = zeros(2,2,(length(T)-1));

for i = 2:(length(T)-1)
   
    %First, Node 1 using Perturbation 2
    
%     f1(i) = (1/timeint)*((Y0smooth(i+1,1) - Y0smooth(i,1))-(Y0smooth(i,1) - Y0smooth(i-1,1)));
%     f1p(i) = (1/timeint)*((YP2smooth(i+1,1) - YP2smooth(i,1))-(Y0smooth(i+1,1) - Y0smooth(i,1)));
%     
%     f = [f1(i); f1p(i)];
%     Y = [(Y0smooth(i+1,1)-Y0smooth(i,1)) (Y0smooth(i+1,2)-Y0smooth(i,2)); (YP2smooth(i,1)-Y0smooth(i,1)) (YP2smooth(i,2)-Y0smooth(i,2))];
%     
%     jacvector1 = lsqlin(Y,f);
%     
%     obstwojacdata(1,:,i-1) = jacvector1';
%     
%     
%     
%     %Now, Node 2 using Perturbation 1
%     
%     f2(i) = (1/timeint)*((Y0smooth(i+1,2) - Y0smooth(i,2))-(Y0smooth(i,2) - Y0smooth(i-1,2)));
%     f2p(i) = (1/timeint)*((YP1smooth(i+1,2) - YP1smooth(i,2))-(Y0smooth(i+1,2) - Y0smooth(i,2)));
%     
%     f = [f2(i); f2p(i)];
%     Y = [(Y0smooth(i+1,1)-Y0smooth(i,1)) (Y0smooth(i+1,2)-Y0smooth(i,2)); (YP1smooth(i,1)-Y0smooth(i,1)) (YP1smooth(i,2)-Y0smooth(i,2))];
        %First, Node 1 using Perturbation 2

    f1(i) = (1/timeint)*((Y0(i+1,1) - Y0(i,1))-(Y0(i,1) - Y0(i-1,1)));
    f1p(i) = (1/timeint)*((YP2(i+1,1) - YP2(i,1))-(Y0(i+1,1) - Y0(i,1)));
    
    f = [f1(i); f1p(i)];
    Y = [(Y0(i+1,1)-Y0(i,1)) (Y0(i+1,2)-Y0(i,2)); (YP2(i,1)-Y0(i,1)) (YP2(i,2)-Y0(i,2))];
    
    jacvector1 = lsqlin(Y,f);
    
    obstwojacdata(1,:,i-1) = jacvector1';
    
    
    
    %Now, Node 2 using Perturbation 1
    
    f2(i) = (1/timeint)*((Y0(i+1,2) - Y0(i,2))-(Y0(i,2) - Y0(i-1,2)));
    f2p(i) = (1/timeint)*((YP1(i+1,2) - YP1(i,2))-(Y0(i+1,2) - Y0(i,2)));
    
    f = [f2(i); f2p(i)];
    Y = [(Y0(i+1,1)-Y0(i,1)) (Y0(i+1,2)-Y0(i,2)); (YP1(i,1)-Y0(i,1)) (YP1(i,2)-Y0(i,2))];

    jacvector2 = lsqlin(Y,f);
    
    obstwojacdata(2,:,i-1) = jacvector2';
    
end

for i = 1:(length(T)-2)
    fmin(i,1) = obstwojacdata(1,1,i);
    fmin(i,2) = obstwojacdata(1,2,i);
    fmin(i,3) = obstwojacdata(2,1,i);
    fmin(i,4) = obstwojacdata(2,2,i);
end
```
