## Supplementary Code for "Network Inference from Perturbation Time Course Data": estimate2node.m

```
function [fmin,finSSE]=estimate2node (data,timeint)
tic
Y=data.Y;
YP1=data.YP1;
YP2=data.YP2;
t=data.T;

% Decide the order of polynomial function to fit the data
poly_order=length(t)-1;

poly_YP1=polyfit(t,YP1(:,1),poly_order);
poly_YP2=polyfit(t,YP2(:,2),poly_order);


         conoptions = optimoptions('fmincon','MaxFunctionEvaluations',10000);
% Set upper and lower bounds for edge weights
         lb = [-10 -10 -10 -10 -10 -10 -10 -10];
         ub = [10 10 10 10 10 10 10 10];

% Obtain LSE solution through fmincon starting from random initial guess. 
% Try/catch is used-if there is an error, it will restart from a different
% random initial guess
         try
            initial=[2*rand(1),2*rand(1),4*rand(1)-2,4*rand(1)-2,2*rand(1),2*rand(1),4*rand(1)-2,4*rand(1)-2];
            [fmin,finSSE]=fmincon(@nestedfun,initial,[],[],[],[],lb,ub,[],conoptions);
         catch
            try
                initial=[2*rand(1),2*rand(1),4*rand(1)-2,4*rand(1)-2,2*rand(1),2*rand(1),4*rand(1)-2,4*rand(1)-2];
                [fmin,finSSE]=fmincon(@nestedfun,initial,[],[],[],[],lb,ub,[],conoptions);
            catch
                try
                    initial=[2*rand(1),2*rand(1),4*rand(1)-2,4*rand(1)-2,2*rand(1),2*rand(1),4*rand(1)-2,4*rand(1)-2];
                    [fmin,finSSE]=fmincon(@nestedfun,initial,[],[],[],[],lb,ub,[],conoptions);
                catch
                    try
                        initial=[2*rand(1),2*rand(1),4*rand(1)-2,4*rand(1)-2,2*rand(1),2*rand(1),4*rand(1)-2,4*rand(1)-2];
                        [fmin,finSSE]=fmincon(@nestedfun,initial,[],[],[],[],lb,ub,[],conoptions);
                    catch
                    end
                end
            end      
         end
% Basal production estimates                    
fmin(1)=-(fmin(3)*Y(1,1)+fmin(4)*Y(1,2));
                
fmin(5)=-(fmin(7)*Y(1,1)+fmin(8)*Y(1,2));

        display(fmin);
        toc
        
function SSE = nestedfun(r2)
        
    % Solve the ODEs 
    S1=-(r2(3)*Y(1,1)+r2(4)*Y(1,2));
                
    S2=-(r2(7)*Y(1,1)+r2(8)*Y(1,2));       
    
[t,guessY] = ode15s(@odeqns,0:timeint:10,[Y(1,1);Y(1,2)],odeset('RelTol',1e-6),r2);
[t,guessYP1] = ode15s(@odep1,0:timeint:10,[YP1(1,2)],odeset('RelTol',1e-6),r2);
[t,guessYP2] = ode15s(@odep2,0:timeint:10,[YP2(1,1)],odeset('RelTol',1e-6),r2);

     SSE = (sum(sum((Y-guessY).^2)+sum((YP1(:,2)-guessYP1).^2)+sum((YP2(:,1)-guessYP2).^2))).^0.5;
    
    function dzdt1 = odeqns(t,z,r)
        dzdt1 = [
            (S1+r(2)+r(3)*z(1)+r(4)*z(2))
            (S2+r(6)+r(7)*z(1)+r(8)*z(2))
        ];
    end
    function dzdt1 = odep1(t,z,r2)
         Y_pert=polyval(poly_YP1,t);

        dzdt1 = [
            (S2+r2(6)+r2(7)*Y_pert+r2(8)*z)
                 ];
    end
       function dzdt1 = odep2(t,z,r2)
        Y_pert=polyval(poly_YP2,t);

        dzdt1 = [
            (S1+r2(2)+r2(3)*z+r2(4)*Y_pert)
        ];
       end
        
end
end
```
