## Supplementary Code for "Network Inference from Perturbation Time Course Data": Make_Figures_1.m

```
% This function reproduces all the MATLAB figures in Figure 1 of the
% manuscript. It is intended to be run in sections. The functions needed 
% to generate the simulation data as well as plot the figures are included.
% Sometimes,the code to generate the data for the models are provided but 
% they take a long time to run. Parallelization is recommended. 
% Alternatively, those Also smaller versions of the data are provided as 
% well which can be run.

%% Figure 1C,E,G

clear all;
% Chosen Parameters for this specific model
params = [0,1,-1,0,0,1,1.5,-0.8];
pert1 = 0;    
pert2 = 0;
timeint=1;
tmax=10;
y0 = [0;0];
display(params);  
noise=0;
data_nostim=eval_data_2node_polyval_nostim(timeint,100,params,y0,y0,y0,pert1,pert2,0);
        
y_0=[data_nostim.Y(101,1),data_nostim.Y(101,2)];
y_p1=[data_nostim.YP1(101,1),data_nostim.YP1(101,2)];
y_p2=[data_nostim.YP2(101,1),data_nostim.YP2(101,2)];
        
data=eval_data_2node_polyval(timeint,tmax,params,y_0,y_p1,y_p2,pert1,pert2,0);

colors=[0 0 1;1 0 0];
a=figure;
a1=subplot(1,2,1);
h1=plot(data.T,data.Y,'.','MarkerSize',12);
ylim([0 3.4]); 
set(h1,{'color'},num2cell(colors,2))
xlabel('Time [AU]','visible','on','FontSize',10);
title("No Noise",'FontSize',9)

%load("data_noise.mat") reproduces the data used in the paper. To generate your
%own random noise data, Uncomment the two lines below load
%("data_noise.mat") and run the section
load("data_noise.mat")
%noise=0.10;
%data=eval_data_2node_polyval(timeint,tmax,params,y0,pert1,pert2,noise);

a2=subplot(1,2,2);
h2=plot(data.T,data.Y,'.','MarkerSize',12);
ylim([0 3.4]); 
set(h2,{'color'},num2cell(colors,2))
xlabel('Time [AU]','visible','on','FontSize',10);
title("10:1 Signal:Noise",'FontSize',9)
res = 300;
set(gcf,'paperunits','inches','paperposition',[0 0 2.5 1.55]);
print('1C.tiff','-dtiff',['-r' num2str(res)]);

b=figure;
data=eval_data_2node_polyval(timeint,tmax,params,y_0,y_p1,y_p2,pert1,pert2,0);
b1=subplot(1,2,1);
h1=plot(data.T,data.YP1,'*','MarkerSize',6);
ylim([0 3.4]);
set(h1,{'color'},num2cell(colors,2))
xlabel('Time [AU]','visible','on','FontSize',10);
title("No Noise",'FontSize',9)

%load("data_noise.mat") reproduces the data used in the paper. To generate your
%own random noise data, Uncomment the two lines below load
%("data_noise.mat") and run the section
load("data_noise.mat")
%noise=0.10;
%data=eval_data_2node_polyval(timeint,tmax,params,y0,pert1,pert2,noise);

b2=subplot(1,2,2);
h2=plot(data.T,data.YP1,'*','MarkerSize',6);
ylim([0 3.4]); 
set(h2,{'color'},num2cell(colors,2));
xlabel('Time [AU]','visible','on','FontSize',10);
title("10:1 Signal:Noise",'FontSize',9)
res = 300;
set(gcf,'paperunits','inches','paperposition',[0 0 2.5 1.55]);
print('1E.tiff','-dtiff',['-r' num2str(res)]);

c=figure;
data=eval_data_2node_polyval(timeint,tmax,params,y_0,y_p1,y_p2,pert1,pert2,0);
c1=subplot(1,2,1);
h1=plot(data.T,data.YP2,'*','MarkerSize',6);
ylim([0 3.4]); 
set(h1,{'color'},num2cell(colors,2))
xlabel('Time [AU]','visible','on','FontSize',10);
title("No Noise",'FontSize',9)

%load("data_noise.mat") reproduces the data used in the paper. To generate your
%own random noise data, Uncomment the two lines below load
%("data_noise.mat") and run the section
load("data_noise.mat")
%noise=0.10;
%data=eval_data_2node_polyval(timeint,tmax,params,y0,pert1,pert2,noise);

c2=subplot(1,2,2);
h2=plot(data.T,data.YP2,'*','MarkerSize',6);
ylim([0 3.4]); 
set(h2,{'color'},num2cell(colors,2));
xlabel('Time [AU]','visible','on','FontSize',10);
title("10:1 Signal:Noise",'FontSize',9)
%sgtitle("Perturbation x_2",'FontSize',12)
res = 300;
set(gcf,'paperunits','inches','paperposition',[0 0 2.5 1.55]);
print('1G.tiff','-dtiff',['-r' num2str(res)]);


%% Figure 1H
% Analytic Solution with no noise
clear all;
params = [0,1,-1,0,0,1,1.5,-0.8];
pert1 = 0;    
pert2 = 0;
timeint=1;
tmax=10;
y0 = [0;0];
display(params);  
noise=0;

data_nostim=eval_data_2node_polyval_nostim(timeint,100,params,y0,y0,y0,pert1,pert2,0);        
y_0=[data_nostim.Y(101,1),data_nostim.Y(101,2)];
y_p1=[data_nostim.YP1(101,1),data_nostim.YP1(101,2)];
y_p2=[data_nostim.YP2(101,1),data_nostim.YP2(101,2)];
data=eval_data_2node_polyval(timeint,tmax,params,y_0,y_p1,y_p2,pert1,pert2,0);

estimateAnalytical=Analytic2node(data,timeint);
c=figure;
    hold on;
    x=0;
    S1b=[-(median(estimateAnalytical(1:9,1))*data.Y(1,1)+median(estimateAnalytical(1:9,2))*data.Y(1,2))
        -(median(estimateAnalytical(1:9,1))*data.YP2(1,1)+median(estimateAnalytical(1:9,2))*data.YP2(1,2))];
    plot(x,median(S1b),'.','Color',[113 18 145]/255,'MarkerSize',20)
    errorbar(x,median(S1b),std(S1b),'k','LineWidth', 1)
    
    x=1;
    S1ex=[-(median(estimateAnalytical(1:9,1))*data.Y(11,1)+median(estimateAnalytical(1:9,2))*data.Y(11,2))
        -(median(estimateAnalytical(1:9,1))*data.YP2(11,1)+median(estimateAnalytical(1:9,2))*data.YP2(11,2))];
    plot(x,median(S1ex),'y.','MarkerSize',20)
    errorbar(x,median(S1ex),std(S1ex),'k','LineWidth', 1)

    x=-1;
    y=estimateAnalytical(1:9,1);
    plot(x,median(y),'m.','MarkerSize',20)
    errorbar(x,median(y),std(y),'k','LineWidth', 1)

    x=0;
    y=estimateAnalytical(1:9,2);    
    plot(x,median(y),'c.','MarkerSize',20)
    errorbar(x,median(y),std(y),'k','LineWidth', 1)
    
    x=0;
    S2b=[-(median(estimateAnalytical(1:9,3))*data.Y(1,1)+median(estimateAnalytical(1:9,4))*data.Y(1,2))
        -(median(estimateAnalytical(1:9,3))*data.YP1(1,1)+median(estimateAnalytical(1:9,4))*data.YP1(1,2))];    
    plot(x,median(S2b),'r.','MarkerSize',20)
    errorbar(x,median(S2b),std(S2b),'k','LineWidth', 1)
    
    x=1;
    S2ex=[-(median(estimateAnalytical(1:9,3))*data.Y(11,1)+median(estimateAnalytical(1:9,4))*data.Y(11,2))
        -(median(estimateAnalytical(1:9,3))*data.YP1(11,1)+median(estimateAnalytical(1:9,4))*data.YP1(11,2))];    
    plot(x,median(S2ex),'g.','MarkerSize',20)
    errorbar(x,median(S2ex),std(S2ex),'k','LineWidth', 1)
    x=1.5;
    y=estimateAnalytical(1:9,3); 
    plot(x,median(y),'b.','MarkerSize',20)
    errorbar(x,median(y),std(y),'k','LineWidth', 1)
    x=-0.8;
    y=estimateAnalytical(1:9,4);    
    plot(x,median(y),'k.','MarkerSize',20)
    errorbar(x,median(y),std(y),'k','LineWidth', 1)
   
%     xlabel('Actual', 'FontSize',10)
%     ylabel('Predicted', 'FontSize',10)  

    ax = gca;
    ax.FontSize = 10;
    ylim([-4 4]); xlim([-2 2]);
    r=refline(1,0);
    r.LineStyle='--';
    r.Color=[0 0 0]; 
    title({"Analytic Solution", "No Noise"}, 'FontSize',10)
    res = 300;
    set(gcf,'paperunits','inches','paperposition',[0 0 2.5 2]);
    print('1H.tiff','-dtiff',['-r' num2str(res)]);

%% %% Figure 1I and inset
% Analytic Solution with 10% noise
clear all;
params = [0,1,-1,0,0,1,1.5,-0.8];
pert1 = 0;    
pert2 = 0;
timeint=1;
tmax=10;
y0 = [0;0];
%load("data_noise.mat") reproduces the data used in the paper. To generate your
%own random noise data, Uncomment the two lines below load
%("data_noise.mat") and run the section

load("data_noise.mat")
%noise=0.10;
%data=eval_data_2node_polyval(timeint,tmax,params,y0,pert1,pert2,noise);
estimateAnalytical=Analytic2node(data,timeint);
c=figure;
hold on;
    x=0;
    S1b=[-(median(estimateAnalytical(1:9,1))*data.Y(1,1)+median(estimateAnalytical(1:9,2))*data.Y(1,2))
        -(median(estimateAnalytical(1:9,1))*data.YP2(1,1)+median(estimateAnalytical(1:9,2))*data.YP2(1,2))];
    plot(x,median(S1b),'.','Color',[113 18 145]/255,'MarkerSize',20)
    errorbar(x,median(S1b),std(S1b),'k','LineWidth', 1)
    
    x=1;
    S1ex=[-(median(estimateAnalytical(1:9,1))*data.Y(11,1)+median(estimateAnalytical(1:9,2))*data.Y(11,2))
        -(median(estimateAnalytical(1:9,1))*data.YP2(11,1)+median(estimateAnalytical(1:9,2))*data.YP2(11,2))];
    plot(x,median(S1ex),'y.','MarkerSize',20)
    errorbar(x,median(S1ex),std(S1ex),'k','LineWidth', 1)

    x=-1;
    y=estimateAnalytical(1:9,1);
    plot(x,median(y),'m.','MarkerSize',20)
    errorbar(x,median(y),std(y),'k','LineWidth', 1)

    x=0;
    y=estimateAnalytical(1:9,2);    
    plot(x,median(y),'c.','MarkerSize',20)
    errorbar(x,median(y),std(y),'k','LineWidth', 1)
    
    x=0;
    S2b=[-(median(estimateAnalytical(1:9,3))*data.Y(1,1)+median(estimateAnalytical(1:9,4))*data.Y(1,2))
        -(median(estimateAnalytical(1:9,3))*data.YP1(1,1)+median(estimateAnalytical(1:9,4))*data.YP1(1,2))];    
    plot(x,median(S2b),'r.','MarkerSize',20)
    errorbar(x,median(S2b),std(S2b),'k','LineWidth', 1)
    
    x=1;
    S2ex=[-(median(estimateAnalytical(1:9,3))*data.Y(11,1)+median(estimateAnalytical(1:9,4))*data.Y(11,2))
        -(median(estimateAnalytical(1:9,3))*data.YP1(11,1)+median(estimateAnalytical(1:9,4))*data.YP1(11,2))];    
    plot(x,median(S2ex),'g.','MarkerSize',20)
    errorbar(x,median(S2ex),std(S2ex),'k','LineWidth', 1)
    x=1.5;
    y=estimateAnalytical(1:9,3); 
    plot(x,median(y),'b.','MarkerSize',20)
    errorbar(x,median(y),std(y),'k','LineWidth', 1)
    x=-0.8;
    y=estimateAnalytical(1:9,4);    
    plot(x,median(y),'k.','MarkerSize',20)
    errorbar(x,median(y),std(y),'k','LineWidth', 1)
%     ax.FontSize = 10;
    ylim([-4 4]); xlim([-2 2]);
%     xlabel('Actual', 'FontSize',10)
%     ylabel('Predicted', 'FontSize',10)
    r=refline(1,0);
    r.LineStyle='--';
    r.Color=[0 0 0]; 
    res = 300;

% For 1i,have the next three lines uncommented. Comment them for inset
% title({"Analytic Solution", "10:1 Signal:Noise"}, 'FontSize',10)
% set(gcf,'paperunits','inches','paperposition',[0 0 2.5 2]);
% print('1I.tiff','-dtiff',['-r' num2str(res)]);
% For inset, comment the previous three lines and uncomment the following four lines and run 
    ylim([-50 50]); xlim([-2 2]);
    size = [0.85 0.50];
    set(gcf,'paperunits','inches','paperposition',[0 0 size]);
    print('1I_inset.tiff','-dtiff',['-r' num2str(res)]);


%% Figure 1J
% Least Squares Solution with 10% noise

clear all;

params = [0,1,-1,0,0,1,1.5,-0.8];
pert1 = 0;    
pert2 = 0;
timeint=1;
tmax=10;
y0 = [0;0];
%load("data_noise.mat") reproduces the data used in the paper. To generate your
%own random noise data, Uncomment the two lines below load
%("data_noise.mat") and run the section

load("data_noise.mat")
% noise=0.10;
% data=eval_data_2node_polyval(timeint,tmax,params,y0,y0,y0,pert1,pert2,noise);
estimateLS=estimate2node(data,timeint);
c=figure;
hold on;
    x=0;
    y=estimateLS(1,1);
    plot(x,y,'.','Color',[113 18 145]/255,'MarkerSize',20)
    x=1;
    y=estimateLS(1,2);
    plot(x,y,'y.','MarkerSize',20)
    x=-1;
    y=estimateLS(1,3);
    plot(x,y,'m.','MarkerSize',20)
    x=0;
    y=estimateLS(1,4);
    plot(x,y,'c.','MarkerSize',20)
    x=0;
    y=estimateLS(1,5);
    plot(x,y,'r.','MarkerSize',20)
    x=1;
    y=estimateLS(1,6);
    plot(x,y,'g.','MarkerSize',20)
    x=1.5;
    y=estimateLS(1,7);
    plot(x,y,'b.','MarkerSize',20)
    x=-0.8;
    y=estimateLS(1,8);
    plot(x,y,'k.','MarkerSize',20)
    ax = gca;
    ax.FontSize = 10;
    ylim([-4 4]); xlim([-2 2]);
    res = 300;

    r=refline(1,0);
    r.LineStyle='--';
    r.Color=[0 0 0]; 
    title({"Least Squares Solution", "10:1 Signal:Noise"}, 'FontSize',10)
    set(gcf,'paperunits','inches','paperposition',[0 0 2.5 2]);
    print('1J','-dtiff',['-r' num2str(res)]);

    
```
