## Supplementary Code for "Network Inference from Perturbation Time Course Data": Readme.rtf

To reproduce analysis and figures, run the script “Make_Figures_1.m”. We recommend that you run the file in sections, as delineated in the file. Some segments that take a long time to run are either commented out or delineated as a new section. To simplify this, we provide all data needed to recreate the figures as mat files. 
