## Supplementary Code for "Network Inference from Perturbation Time Course Data": estimate3node.m

```
function [fmin,finSSE]=estimate3node (data,timeint)
tic
Y=data.Y;
YP1=data.YP1;
YP2=data.YP2;
YP3=data.YP3;
t=data.T;
% Decide the order of polynomial function to fit the data
    poly_order=length(t)-1;


poly_YP1=polyfit(t,YP1(:,1),poly_order);
poly_YP2=polyfit(t,YP2(:,2),poly_order);
poly_YP3=polyfit(t,YP3(:,3),poly_order);
        conoptions = optimoptions('fmincon','MaxFunctionEvaluations',10000);
        % Set upper and lower bounds for edge weights
        lb = [-10 -10 -10 -10 -10 -10 -10 -10 -10 -10 -10 -10 -10 -10 -10];
        ub = [10 10 10 10 10 10 10 10 10 10 10 10 10 10 10];
% Obtain LSE solution through fmincon starting from random initial guess. 
% Try/catch is used-if there is an error, it will restart from a different
% random initial guess
try
        initial=[2*rand(1),2*rand(1),4*rand(1)-2,4*rand(1)-2,4*rand(1)-2,2*rand(1),2*rand(1),4*rand(1)-2,4*rand(1)-2,4*rand(1)-2,2*rand(1),2*rand(1),4*rand(1)-2,4*rand(1)-2,4*rand(1)-2];
        [fmin,finSSE]=fmincon(@nestedfun,initial,[],[],[],[],lb,ub,[],conoptions);
catch
    try
        initial=[2*rand(1),2*rand(1),4*rand(1)-2,4*rand(1)-2,4*rand(1)-2,2*rand(1),2*rand(1),4*rand(1)-2,4*rand(1)-2,4*rand(1)-2,2*rand(1),2*rand(1),4*rand(1)-2,4*rand(1)-2,4*rand(1)-2];
        [fmin,finSSE]=fmincon(@nestedfun,initial,[],[],[],[],lb,ub,[],conoptions);
    catch
        try
        initial=[2*rand(1),2*rand(1),4*rand(1)-2,4*rand(1)-2,4*rand(1)-2,2*rand(1),2*rand(1),4*rand(1)-2,4*rand(1)-2,4*rand(1)-2,2*rand(1),2*rand(1),4*rand(1)-2,4*rand(1)-2,4*rand(1)-2];
            [fmin,finSSE]=fmincon(@nestedfun,initial,[],[],[],[],lb,ub,[],conoptions);
        catch
            try
        initial=[2*rand(1),2*rand(1),4*rand(1)-2,4*rand(1)-2,4*rand(1)-2,2*rand(1),2*rand(1),4*rand(1)-2,4*rand(1)-2,4*rand(1)-2,2*rand(1),2*rand(1),4*rand(1)-2,4*rand(1)-2,4*rand(1)-2];
                [fmin,finSSE]=fmincon(@nestedfun,initial,[],[],[],[],lb,ub,[],conoptions);
            catch
            end
        end
    end
end
fmin(1)=-(fmin(3)*Y(1,1)+fmin(4)*Y(1,2)+fmin(5)*Y(1,3));           
fmin(6)=-(fmin(8)*Y(1,1)+fmin(9)*Y(1,2)+fmin(10)*Y(1,3));
fmin(11)=-(fmin(13)*Y(1,1)+fmin(14)*Y(1,2)+fmin(15)*Y(1,3));

        display(fmin);
        toc
        
function SSE = nestedfun(r2)
        
    S1=-(r2(3)*Y(1,1)+r2(4)*Y(1,2)+r2(5)*Y(1,3));           
    S2=-(r2(8)*Y(1,1)+r2(9)*Y(1,2)+r2(10)*Y(1,3));
    S3=-(r2(13)*Y(1,1)+r2(14)*Y(1,2)+r2(15)*Y(1,3));
    % Solve the ODEs 
           
[t,guessY] = ode15s(@odeqns,0:timeint:10,[Y(1,1);Y(1,2);Y(1,3)],odeset('RelTol',1e-6),r2);
[t,guessYP1] = ode15s(@odep1,0:timeint:10,[YP1(1,2);YP1(1,3)],odeset('RelTol',1e-6),r2);
[t,guessYP2] = ode15s(@odep2,0:timeint:10,[YP2(1,1);YP2(1,3)],odeset('RelTol',1e-6),r2);
[t,guessYP3] = ode15s(@odep3,0:timeint:10,[YP3(1,1);YP3(1,2)],odeset('RelTol',1e-6),r2);

    SSE = (sum(sum((Y-guessY).^2)+sum((YP1(:,2)-guessYP1(:,1)).^2)+sum((YP1(:,3)-guessYP1(:,2)).^2)+sum((YP2(:,1)-guessYP2(:,1)).^2)+sum((YP2(:,3)-guessYP2(:,2)).^2)+sum((YP3(:,1)-guessYP3(:,1)).^2)+sum((YP3(:,2)-guessYP3(:,2)).^2))).^0.5;
    
    function dzdt1 = odeqns(t,z,r)
        dzdt1 = [
            (S1+r(2)+r(3)*z(1)+r(4)*z(2)+r(5)*z(3))
            (S2+r(7)+r(8)*z(1)+r(9)*z(2)+r(10)*z(3))
            (S3+r(12)+r(13)*z(1)+r(14)*z(2)+r(15)*z(3))
        ];
    end
    function dzdt1 = odep1(t,z,r)
        Y_pert=polyval(poly_YP1,t);   
        dzdt1 = [
            (S2+r(6)+r(7)*Y_pert+r(9)*z(1)+r(10)*z(2))
            (S3+r(12)+r(13)*Y_pert+r(14)*z(1)+r(15)*z(2))
                 ];
    end
       function dzdt1 = odep2(t,z,r)
        Y_pert=polyval(poly_YP2,t);
        dzdt1 = [
            (S1+r(2)+r(3)*z(1)+r(4)*Y_pert+r(5)*z(2))
            (S3+r(12)+r(13)*z(1)+r(14)*Y_pert+r(15)*z(2))
        ];
       end
       function dzdt1 = odep3(t,z,r)
        Y_pert=polyval(poly_YP3,t);
        dzdt1 = [
            (S1+r(2)+r(3)*z(1)+r(4)*z(2)+r(5)*Y_pert)
            (S2+r(7)+r(8)*z(1)+r(9)*z(2)+r(10)*Y_pert)
        ];
       end
        
end
end
```
