## Supplementary Code for "Network Inference from Perturbation Time Course Data": eval_data_3node_polyval.m

```
function data= eval_data_3node_polyval(timeint,tmax,params,y0,yp1,yp2,yp3,pert1,pert2,pert3,noise)

% Generate data without perturbation
[t,Y] = ode15s(@odeqns,0:timeint:tmax,y0,odeset('RelTol',1e-6),params);

% Decide the order of polynomial function to fit the data
    poly_order=length(t)-1;


YP1=zeros((1+(tmax/timeint)),3);
YP2=zeros((1+(tmax/timeint)),3);
YP3=zeros((1+(tmax/timeint)),3);

% Generate perturbation data of the perturbed nodes
YP1(:,1)=pert1*Y(:,1);
YP2(:,2)=pert2*Y(:,2);
YP3(:,3)=pert3*Y(:,3);

poly_YP1=polyfit(t,YP1(:,1),poly_order);
poly_YP2=polyfit(t,YP2(:,2),poly_order);
poly_YP3=polyfit(t,YP3(:,3),poly_order);

% Generate perturbation data of the non-perturbed nodes
[t,YP1_data] = ode15s(@odep1,0:timeint:tmax,[yp1(2);yp1(3)],odeset('RelTol',1e-6),params);
[t,YP2_data] = ode15s(@odep2,0:timeint:tmax,[yp2(1);yp2(3)],odeset('RelTol',1e-6),params);
[t,YP3_data] = ode15s(@odep3,0:timeint:tmax,[yp3(1);yp1(2)],odeset('RelTol',1e-6),params);

YP1(:,2)=YP1_data(:,1);
YP1(:,3)=YP1_data(:,2);
YP2(:,1)=YP2_data(:,1);
YP2(:,3)=YP2_data(:,2);
YP3(:,1)=YP3_data(:,1);
YP3(:,2)=YP3_data(:,2);

% Add random noise from normal distribution
D=zeros(1,3);
for i = 1:length(t)
    for j = 1:3
        for k = 1:3
        randomdata = normrnd(0,noise*(abs(Y(i,j))),1,1);
        D(1,k) = Y(i,j) + randomdata;
        end
        Y(i,j)=median(D(1,:));
    end
end

for i = 1:length(t)
    for j = 1:3
        for k = 1:3
        randomdata = normrnd(0,noise*(abs(YP1(i,j))),1,1);
        D(1,k) = YP1(i,j) + randomdata;
        end
        YP1(i,j)=median(D(1,:));
    end
end
        

for i = 1:length(t)
    for j = 1:3
        for k = 1:3
        randomdata = normrnd(0,noise*(abs(YP2(i,j))),1,1);
        D(1,k) = YP2(i,j) + randomdata;
        end
        YP2(i,j)=median(D(1,:));
    end
end


for i = 1:length(t)
    for j = 1:3
        for k = 1:3
        randomdata = normrnd(0,noise*(abs(YP3(i,j))),1,1);
        D(1,k) = YP3(i,j) + randomdata;
        end
        YP3(i,j)=median(D(1,:));
    end
end

data.Y=Y;
data.YP1=YP1;
data.YP2=YP2;
data.YP3=YP3;
data.T=t;
    
    function dzdt1 = odeqns(t,z,r)
        dzdt1 = [
            (r(1)+r(2)+r(3)*z(1)+r(4)*z(2)+r(5)*z(3))
            (r(6)+r(7)+r(8)*z(1)+r(9)*z(2)+r(10)*z(3))
            (r(11)+r(12)+r(13)*z(1)+r(14)*z(2)+r(15)*z(3))
        ];
    end
    function dzdt1 = odep1(t,z,r)
        Y_pert=polyval(poly_YP1,t);   
        dzdt1 = [
            (r(6)+r(7)+r(8)*Y_pert+r(9)*z(1)+r(10)*z(2))
            (r(11)+r(12)+r(13)*Y_pert+r(14)*z(1)+r(15)*z(2))
                 ];
    end
       function dzdt1 = odep2(t,z,r)
        Y_pert=polyval(poly_YP2,t);
        dzdt1 = [
            (r(1)+r(2)+r(3)*z(1)+r(4)*Y_pert+r(5)*z(2))
            (r(11)+r(12)+r(13)*z(1)+r(14)*Y_pert+r(15)*z(2))
        ];
       end
   
        function dzdt1 = odep3(t,z,r)
        Y_pert=polyval(poly_YP3,t);
        dzdt1 = [
            (r(1)+r(2)+r(3)*z(1)+r(4)*z(2)+r(5)*Y_pert)
            (r(6)+r(7)+r(8)*z(1)+r(9)*z(2)+r(10)*Y_pert)
        ];
       end
end
```
