## Supplementary Code for "Network Inference from Perturbation Time Course Data": Make_Figures_2.m

```
% This function reproduces all the MATLAB figures in the manuscript. It is
% intendend to be run in sections. The functions needed to generate the 
% simulation data as well as plot the figures are included. Sometimes,
% the code to generate the data for the models are provided but they take 
% a long time to run. Parallelization is recommended. Alternatively, those 
% sections can be skipped for pre-run mat files which are also provided. 
% Also smaller versions of the data are provided as well which can be run.

    %% Generate full data for Figure 2C,2E
%This section generates 50 random parameter sets for 2 node network and 
%their data for 3,7,11 and 21 timepoints. For each of these cases it 
%generates No Noise, 10:1 S:N, 5:1 S:N and 2:1 S:N and estimates their 
%respective parameters using 10 random multistarts

%This may take a few days to run in a PC without parallelization, so try
%running it using parallelization on multiple nodes.

%In order to run data for only one random parameter and one multistart
%estimate, run the next section

count=50; %Number of random parameter sets
multistart=10; %Number of estimates for every data using a random initial point
pert1 = 0; %perturbation of node 1-100%
pert2 = 0; %perturbation of node 2-100%
tmax=10;
y0 = [0.0;0.0];
[twonoderesults]=twonodewrapper_parfor(count,tmax,y0,pert1,pert2,multistart); 


%% Figure 2C, 2E  (Only one random parameter set and one estimate)
% Run this section to generate data for only one random parameter set and 
% without 10 different initial starts.
clear all; 

count=1; %Number of random parameter sets
multistart=1; %Number of estimates for every data using a random initial point
pert1 = 0; %perturbation of node 1-100%
pert2 = 0; %perturbation of node 2-100%
tmax=10;
y0 = [0.0;0.0];
[twonoderesults]=twonodewrapper(count,tmax,y0,pert1,pert2,multistart);     

count=1;
figure;
ax_x=[-2,2];
ax_y=[-4,4];
subplot(1,4,1)
hold on;
area([0,2],[4,4],'FaceColor','g');
alpha(0.2)
area([0,-2],[-4,-4],'FaceColor','g');
alpha(0.2)
area([0,2],[-4,-4],'FaceColor','r');
alpha(0.2)
area([0,-2],[4,4],'FaceColor','r');
alpha(0.2)

[min3,sum0_3,sum1_3,sum2_3,sum5_3]=fig3func(twonoderesults.actual_param,twonoderesults.predicted_param_3tp,twonoderesults.SSEs_3tp,count);
for i=1:count
hold on;
plot(min3.paramtheor(i,:),min3.parampred5(i,:),"r.",'MarkerSize',20);
plot(min3.paramtheor(i,:),min3.parampred2(i,:),"b.",'MarkerSize',20);
plot(min3.paramtheor(i,:),min3.parampred1(i,:),"g.",'MarkerSize',20);
plot(min3.paramtheor(i,:),min3.parampred0(i,:),"k.",'MarkerSize',20);
xlim([-2 2]);
ylim([-4 4]);
end
plot([0,0],ax_y,'k-','LineWidth',1.5)
plot(ax_y,[0,0],'k-','LineWidth',1.5)
title("3 timepoints",'FontSize',12);

subplot(1,4,2)
hold on;
area([0,2],[4,4],'FaceColor','g');
alpha(0.2)
area([0,-2],[-4,-4],'FaceColor','g');
alpha(0.2)
area([0,2],[-4,-4],'FaceColor','r');
alpha(0.2)
area([0,-2],[4,4],'FaceColor','r');
alpha(0.2)

[min7,sum0_7,sum1_7,sum2_7,sum5_7]=fig3func(twonoderesults.actual_param,twonoderesults.predicted_param_7tp,twonoderesults.SSEs_7tp,count);
for i=1:count
hold on;
plot(min7.paramtheor(i,:),min7.parampred5(i,:),"r.",'MarkerSize',20);
plot(min7.paramtheor(i,:),min7.parampred2(i,:),"b.",'MarkerSize',20);
plot(min7.paramtheor(i,:),min7.parampred1(i,:),"g.",'MarkerSize',20);
plot(min7.paramtheor(i,:),min7.parampred0(i,:),"k.",'MarkerSize',20);
xlim([-2 2]);
ylim([-4 4]);
end
plot([0,0],ax_y,'k-','LineWidth',1.5)
plot(ax_y,[0,0],'k-','LineWidth',1.5)
title("7 timepoints",'FontSize',12);

subplot(1,4,3)
hold on;
area([0,2],[4,4],'FaceColor','g');
alpha(0.2)
area([0,-2],[-4,-4],'FaceColor','g');
alpha(0.2)
area([0,2],[-4,-4],'FaceColor','r');
alpha(0.2)
area([0,-2],[4,4],'FaceColor','r');
alpha(0.2)

[min11,sum0_11,sum1_11,sum2_11,sum5_11]=fig3func(twonoderesults.actual_param,twonoderesults.predicted_param_11tp,twonoderesults.SSEs_11tp,count);
for i=1:count
hold on;
plot(min11.paramtheor(i,:),min11.parampred5(i,:),"r.",'MarkerSize',20);
plot(min11.paramtheor(i,:),min11.parampred2(i,:),"b.",'MarkerSize',20);
plot(min11.paramtheor(i,:),min11.parampred1(i,:),"g.",'MarkerSize',20);
plot(min11.paramtheor(i,:),min11.parampred0(i,:),"k.",'MarkerSize',20);
xlim([-2 2]);
ylim([-4 4]);
end
plot([0,0],ax_y,'k-','LineWidth',1.5)
plot(ax_y,[0,0],'k-','LineWidth',1.5)
title("11 timepoints",'FontSize',12);

subplot(1,4,4)
hold on;
area([0,2],[4,4],'FaceColor','g');
alpha(0.2)
area([0,-2],[-4,-4],'FaceColor','g');
alpha(0.2)
area([0,2],[-4,-4],'FaceColor','r');
alpha(0.2)
area([0,-2],[4,4],'FaceColor','r');
alpha(0.2)

[min21,sum0_21,sum1_21,sum2_21,sum5_21]=fig3func(twonoderesults.actual_param,twonoderesults.predicted_param_21tp,twonoderesults.SSEs_21tp,count);
for i=1:count
hold on;
plot(min21.paramtheor(i,:),min21.parampred5(i,:),"r.",'MarkerSize',20);
plot(min21.paramtheor(i,:),min21.parampred2(i,:),"b.",'MarkerSize',20);
plot(min21.paramtheor(i,:),min21.parampred1(i,:),"g.",'MarkerSize',20);
plot(min21.paramtheor(i,:),min21.parampred0(i,:),"k.",'MarkerSize',20);
xlim([-2 2]);
ylim([-4 4]);
end
plot([0,0],ax_y,'k-','LineWidth',1.5)
plot(ax_y,[0,0],'k-','LineWidth',1.5)
title("21 timepoints",'FontSize',12);

size = [8.5 2.125];
res = 300;
set(gcf,'paperunits','inches','paperposition',[0 0 size]);
print('2node_1param_all.tiff','-dtiff',['-r' num2str(res)]);

d=figure; 
in=roundn([sum0_3,sum1_3,sum2_3,sum5_3;sum0_7,sum1_7,sum2_7,sum5_7;sum0_11,sum1_11,sum2_11,sum5_11;sum0_21,sum1_21,sum2_21,sum5_21],-2);
hm=heatmap(in');
caxis([0.8 1])
hm.Colormap=hot;
hm.YData = ["No Noise" "10:1 S:N" "5:1 S:N" "2:1 S:N"];
hm.XData = ["3 Timepoints" "7 Timepoints" "11 Timepoints" "21 Timepoints"];
hm.Title="Classification Accuracy";
hm.FontSize=11;
res = 300;
set(gcf,'paperunits','inches','PaperPosition',[0 0 3.5 3]);
print('2node_1param_hm.tiff','-dtiff',['-r' num2str(res)]);
%% Figure 2C
clear all;
load('param2wrapper_paper.mat')
%Uncomment the next line and run if you generated the full data. Otherwise
%just run the section to generate paper figures
%load('param2wrapper.mat')

count=50;
figure;
ax_x=[-2,2];
ax_y=[-4,4];
subplot(1,4,1)
hold on;
area([0,2],[4,4],'FaceColor','g');
alpha(0.2)
area([0,-2],[-4,-4],'FaceColor','g');
alpha(0.2)
area([0,2],[-4,-4],'FaceColor','r');
alpha(0.2)
area([0,-2],[4,4],'FaceColor','r');
alpha(0.2)

[min3,sum0_3,sum1_3,sum2_3,sum5_3]=fig3func(twonoderesults.actual_param,twonoderesults.predicted_param_3tp,twonoderesults.SSEs_3tp,count);
for i=1:count
hold on;
plot(min3.paramtheor(i,:),min3.parampred5(i,:),'r.','MarkerSize',20);
plot(min3.paramtheor(i,:),min3.parampred2(i,:),'b.','MarkerSize',20);
plot(min3.paramtheor(i,:),min3.parampred1(i,:),'g.','MarkerSize',20);
plot(min3.paramtheor(i,:),min3.parampred0(i,:),'k.','MarkerSize',20);
xlim([-2 2]);
ylim([-4 4]);
end
plot([0,0],ax_y,'k-','LineWidth',1.5)
plot(ax_y,[0,0],'k-','LineWidth',1.5)
title("3 timepoints",'FontSize',12);

subplot(1,4,2)
hold on;
area([0,2],[4,4],'FaceColor','g');
alpha(0.2)
area([0,-2],[-4,-4],'FaceColor','g');
alpha(0.2)
area([0,2],[-4,-4],'FaceColor','r');
alpha(0.2)
area([0,-2],[4,4],'FaceColor','r');
alpha(0.2)

[min7,sum0_7,sum1_7,sum2_7,sum5_7]=fig3func(twonoderesults.actual_param,twonoderesults.predicted_param_7tp,twonoderesults.SSEs_7tp,count);
for i=1:count
hold on;
plot(min7.paramtheor(i,:),min7.parampred5(i,:),'r.','MarkerSize',20);
plot(min7.paramtheor(i,:),min7.parampred2(i,:),'b.','MarkerSize',20);
plot(min7.paramtheor(i,:),min7.parampred1(i,:),'g.','MarkerSize',20);
plot(min7.paramtheor(i,:),min7.parampred0(i,:),'k.','MarkerSize',20);
xlim([-2 2]);
ylim([-4 4]);
end
plot([0,0],ax_y,'k-','LineWidth',1.5)
plot(ax_y,[0,0],'k-','LineWidth',1.5)
title("7 timepoints",'FontSize',12);

subplot(1,4,3)
hold on;
area([0,2],[4,4],'FaceColor','g');
alpha(0.2)
area([0,-2],[-4,-4],'FaceColor','g');
alpha(0.2)
area([0,2],[-4,-4],'FaceColor','r');
alpha(0.2)
area([0,-2],[4,4],'FaceColor','r');
alpha(0.2)

[min11,sum0_11,sum1_11,sum2_11,sum5_11]=fig3func(twonoderesults.actual_param,twonoderesults.predicted_param_11tp,twonoderesults.SSEs_11tp,count);
for i=1:count
hold on;
plot(min11.paramtheor(i,:),min11.parampred5(i,:),'r.','MarkerSize',20);
plot(min11.paramtheor(i,:),min11.parampred2(i,:),'b.','MarkerSize',20);
plot(min11.paramtheor(i,:),min11.parampred1(i,:),'g.','MarkerSize',20);
plot(min11.paramtheor(i,:),min11.parampred0(i,:),'k.','MarkerSize',20);
xlim([-2 2]);
ylim([-4 4]);
end

  plot([0,0],ax_y,'k-','LineWidth',1.5)
  plot(ax_y,[0,0],'k-','LineWidth',1.5)
title("11 timepoints",'FontSize',12);

subplot(1,4,4)
hold on;
area([0,2],[4,4],'FaceColor','g');
alpha(0.2)
area([0,-2],[-4,-4],'FaceColor','g');
alpha(0.2)
area([0,2],[-4,-4],'FaceColor','r');
alpha(0.2)
area([0,-2],[4,4],'FaceColor','r');
alpha(0.2)

[min21,sum0_21,sum1_21,sum2_21,sum5_21]=fig3func(twonoderesults.actual_param,twonoderesults.predicted_param_21tp,twonoderesults.SSEs_21tp,count);
for i=1:count
hold on;
plot(min21.paramtheor(i,:),min21.parampred5(i,:),'r.','MarkerSize',20);
plot(min21.paramtheor(i,:),min21.parampred2(i,:),'b.','MarkerSize',20);
plot(min21.paramtheor(i,:),min21.parampred1(i,:),'g.','MarkerSize',20);
plot(min21.paramtheor(i,:),min21.parampred0(i,:),'k.','MarkerSize',20);
xlim([-2 2]);
ylim([-4 4]);
end
  plot([0,0],ax_y,'k-','LineWidth',1.5)
  plot(ax_y,[0,0],'k-','LineWidth',1.5)
title("21 timepoints",'FontSize',12);

size = [6.5 2.125];
res = 300;
set(gcf,'paperunits','inches','paperposition',[0 0 size]);
print('2C.tiff','-dtiff',['-r' num2str(res)]);


%% Figure 2E
clear all;
load('param2wrapper_paper.mat')
%Uncomment the next line and run if you generated the full data. Otherwise
%just run the section to generate paper figures
%load('param2wrapper.mat')

count=50;
[min3,sum0_3,sum1_3,sum2_3,sum5_3]=fig3func(twonoderesults.actual_param,twonoderesults.predicted_param_3tp,twonoderesults.SSEs_3tp,count);
[min7,sum0_7,sum1_7,sum2_7,sum5_7]=fig3func(twonoderesults.actual_param,twonoderesults.predicted_param_7tp,twonoderesults.SSEs_7tp,count);
[min11,sum0_11,sum1_11,sum2_11,sum5_11]=fig3func(twonoderesults.actual_param,twonoderesults.predicted_param_11tp,twonoderesults.SSEs_11tp,count);
[min21,sum0_21,sum1_21,sum2_21,sum5_21]=fig3func(twonoderesults.actual_param,twonoderesults.predicted_param_21tp,twonoderesults.SSEs_21tp,count);

d=figure; 
in=roundn([sum0_3,sum1_3,sum2_3,sum5_3;sum0_7,sum1_7,sum2_7,sum5_7;sum0_11,sum1_11,sum2_11,sum5_11;sum0_21,sum1_21,sum2_21,sum5_21],-2);
hm=heatmap(in');
caxis([0.5 1])
hm.Colormap=hot;
hm.YData = ["No Noise" "10:1 S:N" "5:1 S:N" "2:1 S:N"];
hm.XData = ["3 Timepoints" "7 Timepoints" "11 Timepoints" "21 Timepoints"];
hm.Title="Classification Accuracy";
hm.FontSize=11;
res = 300;
set(gcf,'paperunits','inches','PaperPosition',[0 0 3.5 3]);
print('2E.tiff','-dtiff',['-r' num2str(res)]);
%% Number of oscillating networks and positive feedbacks
clear;
load('param2wrapper_paper.mat')
params=zeros(1,6);
j=0;
k=0;
for i=1:50
        param=twonoderesults.actual_param{1,i};
        Jacob=[param(3),param(4);param(7),param(8)];
        eigen=eigs(Jacob);
        imag_eigen=imag(eigen(1));
        feedback=[param(3);param(8)];
        if abs(imag_eigen)>0
            j=j+1;
        end
        if feedback(1)>0||feedback(2)>0
            k=k+1;
        end
end    
%j=networks with oscillating properties 
%k=networks with a positive self regulation
%% Generate full data for Figure 2D, 2F
%This section generates 50 random parameter sets and their data for 3,7,11 and
%21 timepoints. For each of these cases it generates No Noise, 10:1 S:N, 
%5:1 S:N and 2:1 S:N and estimates their respective parameters using
%10 random multistarts

%This may take a few days to run in a PC without parallelization, so try
%running it using parallelization on multiple nodes.

%In order to run data for only one random parameter and one multistart
%estimate, run the next section
clear all;
count=50;
multistart=10;
pert1 = 0;    
pert2 = 0;
pert3 = 0;
tmax=10;
y0 = [0.0;0.0;0.0];
[threenoderesults]=threenodewrapper_parfor(count,tmax,y0,pert1,pert2,pert3,multistart);

%% Figure 2D, 2F (Only one random parameter set and one estimate)

%Run this section to generate data for only one parameter set and without
%10 different initial starts.
clear all;
count=1;
multistart=1;
pert1 = 0;    
pert2 = 0;
pert3 = 0;
tmax=10;
y0 = [0.0;0.0;0.0];
[threenoderesults]=threenodewrapper(count,tmax,y0,pert1,pert2,pert3,multistart);

figure;
ax_x=[-2,2];
ax_y=[-4,4];
subplot(1,4,1)
hold on;
area([0,2],[4,4],'FaceColor','g');
alpha(0.2)
area([0,-2],[-4,-4],'FaceColor','g');
alpha(0.2)
area([0,2],[-4,-4],'FaceColor','r');
alpha(0.2)
area([0,-2],[4,4],'FaceColor','r');
alpha(0.2)
[min3,sum0_3,sum1_3,sum2_3,sum5_3]=fig3func(threenoderesults.actual_param,threenoderesults.predicted_param_3tp,threenoderesults.SSEs_3tp,count);
for i=1:count
hold on;
plot(min3.paramtheor(i,:),min3.parampred5(i,:),"r.",'MarkerSize',20);
plot(min3.paramtheor(i,:),min3.parampred2(i,:),"b.",'MarkerSize',20);
plot(min3.paramtheor(i,:),min3.parampred1(i,:),"g.",'MarkerSize',20);
plot(min3.paramtheor(i,:),min3.parampred0(i,:),"k.",'MarkerSize',20);
xlim([-2 2]);
ylim([-4 4]);
end
plot([0,0],ax_y,'k-','LineWidth',1.5)
plot(ax_y,[0,0],'k-','LineWidth',1.5)
title("3 timepoints",'FontSize',12);

subplot(1,4,2)
area([0,2],[4,4],'FaceColor','g');
alpha(0.2)
area([0,-2],[-4,-4],'FaceColor','g');
alpha(0.2)
area([0,2],[-4,-4],'FaceColor','r');
alpha(0.2)
area([0,-2],[4,4],'FaceColor','r');
alpha(0.2)
[min7,sum0_7,sum1_7,sum2_7,sum5_7]=fig3func(threenoderesults.actual_param,threenoderesults.predicted_param_7tp,threenoderesults.SSEs_7tp,count);
for i=1:count
hold on;
plot(min7.paramtheor(i,:),min7.parampred5(i,:),"r.",'MarkerSize',20);
plot(min7.paramtheor(i,:),min7.parampred2(i,:),"b.",'MarkerSize',20);
plot(min7.paramtheor(i,:),min7.parampred1(i,:),"g.",'MarkerSize',20);
plot(min7.paramtheor(i,:),min7.parampred0(i,:),"k.",'MarkerSize',20);
xlim([-2 2]);
ylim([-4 4]);
end
plot([0,0],ax_y,'k-','LineWidth',1.5)
plot(ax_y,[0,0],'k-','LineWidth',1.5)
title("7 timepoints",'FontSize',12);

subplot(1,4,3)
area([0,2],[4,4],'FaceColor','g');
alpha(0.2)
area([0,-2],[-4,-4],'FaceColor','g');
alpha(0.2)
area([0,2],[-4,-4],'FaceColor','r');
alpha(0.2)
area([0,-2],[4,4],'FaceColor','r');
alpha(0.2)
[min11,sum0_11,sum1_11,sum2_11,sum5_11]=fig3func(threenoderesults.actual_param,threenoderesults.predicted_param_11tp,threenoderesults.SSEs_11tp,count);
for i=1:count
hold on;
plot(min11.paramtheor(i,:),min11.parampred5(i,:),"r.",'MarkerSize',20);
plot(min11.paramtheor(i,:),min11.parampred2(i,:),"b.",'MarkerSize',20);
plot(min11.paramtheor(i,:),min11.parampred1(i,:),"g.",'MarkerSize',20);
plot(min11.paramtheor(i,:),min11.parampred0(i,:),"k.",'MarkerSize',20);
xlim([-2 2]);
ylim([-4 4]);
end
plot([0,0],ax_y,'k-','LineWidth',1.5)
plot(ax_y,[0,0],'k-','LineWidth',1.5)
title("11 timepoints",'FontSize',12);

subplot(1,4,4)
area([0,2],[4,4],'FaceColor','g');
alpha(0.2)
area([0,-2],[-4,-4],'FaceColor','g');
alpha(0.2)
area([0,2],[-4,-4],'FaceColor','r');
alpha(0.2)
area([0,-2],[4,4],'FaceColor','r');
alpha(0.2)
[min21,sum0_21,sum1_21,sum2_21,sum5_21]=fig3func(threenoderesults.actual_param,threenoderesults.predicted_param_21tp,threenoderesults.SSEs_21tp,count);
for i=1:count
hold on;
plot(min21.paramtheor(i,:),min21.parampred5(i,:),"r.",'MarkerSize',20);
plot(min21.paramtheor(i,:),min21.parampred2(i,:),"b.",'MarkerSize',20);
plot(min21.paramtheor(i,:),min21.parampred1(i,:),"g.",'MarkerSize',20);
plot(min21.paramtheor(i,:),min21.parampred0(i,:),"k.",'MarkerSize',20);
xlim([-2 2]);
ylim([-4 4]);
end
plot([0,0],ax_y,'k-','LineWidth',1.5)
plot(ax_y,[0,0],'k-','LineWidth',1.5)
title("21 timepoints",'FontSize',12);

size = [8.5 2.125];
res = 300;
set(gcf,'paperunits','inches','paperposition',[0 0 size]);
print('3node_1param_all.tiff','-dtiff',['-r' num2str(res)]);

d=figure; 
in=roundn([sum0_3,sum1_3,sum2_3,sum5_3;sum0_7,sum1_7,sum2_7,sum5_7;sum0_11,sum1_11,sum2_11,sum5_11;sum0_21,sum1_21,sum2_21,sum5_21],-2);
hm=heatmap(in');
caxis([0.8 1])
hm.Colormap=hot;
hm.YData = ["No Noise" "10:1 S:N" "5:1 S:N" "2:1 S:N"];
hm.XData = ["3 Timepoints" "7 Timepoints" "11 Timepoints" "21 Timepoints"];
hm.Title="Classification Accuracy";
hm.FontSize=11;
res = 300;
set(gcf,'paperunits','inches','PaperPosition',[0 0 3.5 3]);
print('3node_1param_hm.tiff','-dtiff',['-r' num2str(res)]);

%% Figure 2D
clear all;
load('param3wrapper_paper.mat')
count=50;
ax_x=[-2,2];
ax_y=[-4,4];
figure;
subplot(1,4,1)
hold on;
area([0,2],[4,4],'FaceColor','g');
alpha(0.2)
area([0,-2],[-4,-4],'FaceColor','g');
alpha(0.2)
area([0,2],[-4,-4],'FaceColor','r');
alpha(0.2)
area([0,-2],[4,4],'FaceColor','r');
alpha(0.2)

[min3,sum0_3,sum1_3,sum2_3,sum5_3]=fig3func(threenoderesults.actual_param,threenoderesults.predicted_param_3tp,threenoderesults.SSEs_3tp,count);
for i=1:count
hold on;
plot(min3.paramtheor(i,:),min3.parampred5(i,:),"r.",'MarkerSize',20);
plot(min3.paramtheor(i,:),min3.parampred2(i,:),"b.",'MarkerSize',20);
plot(min3.paramtheor(i,:),min3.parampred1(i,:),"g.",'MarkerSize',20);
plot(min3.paramtheor(i,:),min3.parampred0(i,:),"k.",'MarkerSize',20);
xlim([-2 2]);
ylim([-4 4]);
end
  plot([0,0],ax_y,'k-','LineWidth',1.5)
  plot(ax_y,[0,0],'k-','LineWidth',1.5)
title("3 timepoints",'FontSize',12);

subplot(1,4,2)
hold on;
area([0,2],[4,4],'FaceColor','g');
alpha(0.2)
area([0,-2],[-4,-4],'FaceColor','g');
alpha(0.2)
area([0,2],[-4,-4],'FaceColor','r');
alpha(0.2)
area([0,-2],[4,4],'FaceColor','r');
alpha(0.2)

[min7,sum0_7,sum1_7,sum2_7,sum5_7]=fig3func(threenoderesults.actual_param,threenoderesults.predicted_param_7tp,threenoderesults.SSEs_7tp,count);
for i=1:count
hold on;
plot(min7.paramtheor(i,:),min7.parampred5(i,:),"r.",'MarkerSize',20);
plot(min7.paramtheor(i,:),min7.parampred2(i,:),"b.",'MarkerSize',20);
plot(min7.paramtheor(i,:),min7.parampred1(i,:),"g.",'MarkerSize',20);
plot(min7.paramtheor(i,:),min7.parampred0(i,:),"k.",'MarkerSize',20);
xlim([-2 2]);
ylim([-4 4]);
end
  plot([0,0],ax_y,'k-','LineWidth',1.5)
  plot(ax_y,[0,0],'k-','LineWidth',1.5)
title("7 timepoints",'FontSize',12);

subplot(1,4,3)
hold on;
area([0,2],[4,4],'FaceColor','g');
alpha(0.2)
area([0,-2],[-4,-4],'FaceColor','g');
alpha(0.2)
area([0,2],[-4,-4],'FaceColor','r');
alpha(0.2)
area([0,-2],[4,4],'FaceColor','r');
alpha(0.2)

[min11,sum0_11,sum1_11,sum2_11,sum5_11]=fig3func(threenoderesults.actual_param,threenoderesults.predicted_param_11tp,threenoderesults.SSEs_11tp,count);
for i=1:count
hold on;
plot(min11.paramtheor(i,:),min11.parampred5(i,:),"r.",'MarkerSize',20);
plot(min11.paramtheor(i,:),min11.parampred2(i,:),"b.",'MarkerSize',20);
plot(min11.paramtheor(i,:),min11.parampred1(i,:),"g.",'MarkerSize',20);
plot(min11.paramtheor(i,:),min11.parampred0(i,:),"k.",'MarkerSize',20);
xlim([-2 2]);
ylim([-4 4]);
end
  plot([0,0],ax_y,'k-','LineWidth',1.5)
  plot(ax_y,[0,0],'k-','LineWidth',1.5)
title("11 timepoints",'FontSize',12);

subplot(1,4,4)
hold on;
area([0,2],[4,4],'FaceColor','g');
alpha(0.2)
area([0,-2],[-4,-4],'FaceColor','g');
alpha(0.2)
area([0,2],[-4,-4],'FaceColor','r');
alpha(0.2)
area([0,-2],[4,4],'FaceColor','r');
alpha(0.2)

[min21,sum0_21,sum1_21,sum2_21,sum5_21]=fig3func(threenoderesults.actual_param,threenoderesults.predicted_param_21tp,threenoderesults.SSEs_21tp,count);
for i=1:count
hold on;
plot(min21.paramtheor(i,:),min21.parampred5(i,:),"r.",'MarkerSize',20);
plot(min21.paramtheor(i,:),min21.parampred2(i,:),"b.",'MarkerSize',20);
plot(min21.paramtheor(i,:),min21.parampred1(i,:),"g.",'MarkerSize',20);
plot(min21.paramtheor(i,:),min21.parampred0(i,:),"k.",'MarkerSize',20);
xlim([-2 2]);
ylim([-4 4]);
end
  plot([0,0],ax_y,'k-','LineWidth',1.5)
  plot(ax_y,[0,0],'k-','LineWidth',1.5)
title("21 timepoints",'FontSize',12);

size = [6.5 2.125];
res = 300;
set(gcf,'paperunits','inches','paperposition',[0 0 size]);
print('2D.tiff','-dtiff',['-r' num2str(res)]);
%% Figure 2F
clear all;
load('param3wrapper_paper.mat')
%Uncomment the next line and run if you generated the full data. Otherwise
%just run the section to generate paper figures
%load('param3wrapper.mat')
count=50;
actual_param=threenoderesults.actual_param;
[min3,sum0_3,sum1_3,sum2_3,sum5_3]=fig3func(threenoderesults.actual_param,threenoderesults.predicted_param_3tp,threenoderesults.SSEs_3tp,count);
[min7,sum0_7,sum1_7,sum2_7,sum5_7]=fig3func(threenoderesults.actual_param,threenoderesults.predicted_param_7tp,threenoderesults.SSEs_7tp,count);
[min11,sum0_11,sum1_11,sum2_11,sum5_11]=fig3func(threenoderesults.actual_param,threenoderesults.predicted_param_11tp,threenoderesults.SSEs_11tp,count);
[min21,sum0_21,sum1_21,sum2_21,sum5_21]=fig3func(threenoderesults.actual_param,threenoderesults.predicted_param_21tp,threenoderesults.SSEs_21tp,count);

d=figure; 
in=roundn([sum0_3,sum1_3,sum2_3,sum5_3;sum0_7,sum1_7,sum2_7,sum5_7;sum0_11,sum1_11,sum2_11,sum5_11;sum0_21,sum1_21,sum2_21,sum5_21],-2);
hm=heatmap(in');
caxis([0.5 1])
hm.Colormap=hot;
hm.YData = ["No Noise" "10:1 S:N" "5:1 S:N" "2:1 S:N"];
hm.XData = ["3 Timepoints" "7 Timepoints" "11 Timepoints" "21 Timepoints"];
hm.Title="Classification Accuracy";
hm.FontSize=11;
res = 300;
set(gcf,'paperunits','inches','PaperPosition',[0 0 3.5 3]);
print('2F.tiff','-dtiff',['-r' num2str(res)]);

%% Oscillating and positive feedbacks for 3 node network
load('param3wrapper_paper.mat')
params=zeros(1,12);
j=0;
k=0;
for i=1:50
        param=threenoderesults.actual_param{1,i};
        Jacob=[param(3),param(4),param(5);param(8),param(9),param(10);param(13),param(14),param(15)];
        eigen=eigs(Jacob);
        imag_eigen=imag(eigen(1));
        feedback=[param(3);param(9);param(15)];
        if abs(imag_eigen)>0
            j=j+1;
        end
        if feedback(1)>0||feedback(2)>0||feedback(3)>0
            k=k+1;
        end
end
%j=networks with oscillating properties 
%k=networks with a positive self regulation
%% Supplement 1B
load ("param2wrapper_paper.mat")

for i=1:50
    plot(twonoderesults.datas_21tp{1,i}.T,twonoderesults.datas_21tp{1,i}.Y(:,1)./max(twonoderesults.datas_21tp{1,i}.Y(:,1)),'LineWidth',2)
    plot(twonoderesults.datas_21tp{1,i}.T,twonoderesults.datas_21tp{1,i}.Y(:,2)./max(twonoderesults.datas_21tp{1,i}.Y(:,2)),'LineWidth',2)

%     plot(twonoderesults.datas_21tp{1,i}.T,twonoderesults.datas_21tp{1,i}.YP1/max(twonoderesults.datas_21tp{1,i}.YP1))
%     plot(twonoderesults.datas_21tp{1,i}.T,twonoderesults.datas_21tp{1,i}.YP2/max(twonoderesults.datas_21tp{1,i}.YP2))
    hold on
end
xlabel('Time(AU)')
ylabel('Normalized data(AU)')
title({"Random Connection 2-Node Model", "Normalized Data"}, 'FontSize',11)
ylim([0 1.05])
res = 300;
set(gcf,'paperunits','inches','PaperPosition',[0 0 2.7 2.7]);
print('S1B.tiff','-dtiff',['-r' num2str(res)]);
%% Supplement 1C
clear;
load('param2wrapper_paper.mat')
count=50;
bot=0;
top=0.5;
actual_param=twonoderesults.actual_param;
[min3,sum0_3,sum1_3,sum2_3,sum5_3]=fig3func_ranges(actual_param,twonoderesults.predicted_param_3tp,twonoderesults.SSEs_3tp,count,bot,top);
[min7,sum0_7,sum1_7,sum2_7,sum5_7]=fig3func_ranges(actual_param,twonoderesults.predicted_param_7tp,twonoderesults.SSEs_7tp,count,bot,top);
[min11,sum0_11,sum1_11,sum2_11,sum5_11]=fig3func_ranges(actual_param,twonoderesults.predicted_param_11tp,twonoderesults.SSEs_11tp,count,bot,top);
[min21,sum0_21,sum1_21,sum2_21,sum5_21]=fig3func_ranges(actual_param,twonoderesults.predicted_param_21tp,twonoderesults.SSEs_21tp,count,bot,top);
d=figure; 
in=roundn([sum0_3,sum1_3,sum2_3,sum5_3;sum0_7,sum1_7,sum2_7,sum5_7;sum0_11,sum1_11,sum2_11,sum5_11;sum0_21,sum1_21,sum2_21,sum5_21],-2);
hm=heatmap(in');
caxis([0.5 1])
hm.Colormap=hot;
hm.YData = ["No Noise" "10:1 S:N" "5:1 S:N" "2:1 S:N"];
hm.XData = ["3 Timepoints" "7 Timepoints" "11 Timepoints" "21 Timepoints"];
hm.Title="Classification Accuracy";
hm.FontSize=11;
res = 300;
set(gcf,'paperunits','inches','PaperPosition',[0 0 2.8 2.5]);
print('S1C.tiff','-dtiff',['-r' num2str(res)]);
%% Supplement 1D
clear;
load('param2wrapper_paper.mat')
count=50;
bot=0.5;
top=1;
actual_param=twonoderesults.actual_param;
[min3,sum0_3,sum1_3,sum2_3,sum5_3]=fig3func_ranges(actual_param,twonoderesults.predicted_param_3tp,twonoderesults.SSEs_3tp,count,bot,top);
[min7,sum0_7,sum1_7,sum2_7,sum5_7]=fig3func_ranges(actual_param,twonoderesults.predicted_param_7tp,twonoderesults.SSEs_7tp,count,bot,top);
[min11,sum0_11,sum1_11,sum2_11,sum5_11]=fig3func_ranges(actual_param,twonoderesults.predicted_param_11tp,twonoderesults.SSEs_11tp,count,bot,top);
[min21,sum0_21,sum1_21,sum2_21,sum5_21]=fig3func_ranges(actual_param,twonoderesults.predicted_param_21tp,twonoderesults.SSEs_21tp,count,bot,top);
d=figure; 
in=roundn([sum0_3,sum1_3,sum2_3,sum5_3;sum0_7,sum1_7,sum2_7,sum5_7;sum0_11,sum1_11,sum2_11,sum5_11;sum0_21,sum1_21,sum2_21,sum5_21],-2);
hm=heatmap(in');
caxis([0.5 1])
hm.Colormap=hot;
hm.YData = ["No Noise" "10:1 S:N" "5:1 S:N" "2:1 S:N"];
hm.XData = ["3 Timepoints" "7 Timepoints" "11 Timepoints" "21 Timepoints"];
hm.Title="Classification Accuracy";
hm.FontSize=11;
res = 300;
set(gcf,'paperunits','inches','PaperPosition',[0 0 2.8 2.5]);
print('S1D.tiff','-dtiff',['-r' num2str(res)]);
%% Supplement 1E
clear;
load('param2wrapper_paper.mat')
count=50;
bot=1;
top=1.5;
actual_param=twonoderesults.actual_param;
[min3,sum0_3,sum1_3,sum2_3,sum5_3]=fig3func_ranges(actual_param,twonoderesults.predicted_param_3tp,twonoderesults.SSEs_3tp,count,bot,top);
[min7,sum0_7,sum1_7,sum2_7,sum5_7]=fig3func_ranges(actual_param,twonoderesults.predicted_param_7tp,twonoderesults.SSEs_7tp,count,bot,top);
[min11,sum0_11,sum1_11,sum2_11,sum5_11]=fig3func_ranges(actual_param,twonoderesults.predicted_param_11tp,twonoderesults.SSEs_11tp,count,bot,top);
[min21,sum0_21,sum1_21,sum2_21,sum5_21]=fig3func_ranges(actual_param,twonoderesults.predicted_param_21tp,twonoderesults.SSEs_21tp,count,bot,top);
d=figure; 
in=roundn([sum0_3,sum1_3,sum2_3,sum5_3;sum0_7,sum1_7,sum2_7,sum5_7;sum0_11,sum1_11,sum2_11,sum5_11;sum0_21,sum1_21,sum2_21,sum5_21],-2);
hm=heatmap(in');
caxis([0.5 1])
hm.Colormap=hot;
hm.YData = ["No Noise" "10:1 S:N" "5:1 S:N" "2:1 S:N"];
hm.XData = ["3 Timepoints" "7 Timepoints" "11 Timepoints" "21 Timepoints"];
hm.Title="Classification Accuracy";
hm.FontSize=11;
res = 300;
set(gcf,'paperunits','inches','PaperPosition',[0 0 2.8 2.5]);
print('S1E.tiff','-dtiff',['-r' num2str(res)]);
%% Supplement 1F
clear;
load('param2wrapper_paper.mat')
count=50;
bot=1.5;
top=2;
actual_param=twonoderesults.actual_param;
[min3,sum0_3,sum1_3,sum2_3,sum5_3]=fig3func_ranges(actual_param,twonoderesults.predicted_param_3tp,twonoderesults.SSEs_3tp,count,bot,top);
[min7,sum0_7,sum1_7,sum2_7,sum5_7]=fig3func_ranges(actual_param,twonoderesults.predicted_param_7tp,twonoderesults.SSEs_7tp,count,bot,top);
[min11,sum0_11,sum1_11,sum2_11,sum5_11]=fig3func_ranges(actual_param,twonoderesults.predicted_param_11tp,twonoderesults.SSEs_11tp,count,bot,top);
[min21,sum0_21,sum1_21,sum2_21,sum5_21]=fig3func_ranges(actual_param,twonoderesults.predicted_param_21tp,twonoderesults.SSEs_21tp,count,bot,top);
d=figure; 
in=roundn([sum0_3,sum1_3,sum2_3,sum5_3;sum0_7,sum1_7,sum2_7,sum5_7;sum0_11,sum1_11,sum2_11,sum5_11;sum0_21,sum1_21,sum2_21,sum5_21],-2);
hm=heatmap(in');
caxis([0.5 1])
hm.Colormap=hot;
hm.YData = ["No Noise" "10:1 S:N" "5:1 S:N" "2:1 S:N"];
hm.XData = ["3 Timepoints" "7 Timepoints" "11 Timepoints" "21 Timepoints"];
hm.Title="Classification Accuracy";
hm.FontSize=11;
res = 300;
set(gcf,'paperunits','inches','PaperPosition',[0 0 2.8 2.5]);
print('S1F.tiff','-dtiff',['-r' num2str(res)]);

%% Supplement @B
clear;
load ("param3wrapper_paper.mat")
figure
for i=1:50
    plot(threenoderesults.datas_21tp{1,i}.T,threenoderesults.datas_21tp{1,i}.Y(:,1)./max(threenoderesults.datas_21tp{1,i}.Y(:,1)),'LineWidth',2)
    plot(threenoderesults.datas_21tp{1,i}.T,threenoderesults.datas_21tp{1,i}.Y(:,2)./max(threenoderesults.datas_21tp{1,i}.Y(:,2)),'LineWidth',2)
    plot(threenoderesults.datas_21tp{1,i}.T,threenoderesults.datas_21tp{1,i}.Y(:,3)./max(threenoderesults.datas_21tp{1,i}.Y(:,3)),'LineWidth',2)

%     plot(threenoderesults.datas_21tp{1,i}.T,threenoderesults.datas_21tp{1,i}.YP1/max(threenoderesults.datas_21tp{1,i}.YP1))
%     plot(threenoderesults.datas_21tp{1,i}.T,threenoderesults.datas_21tp{1,i}.YP2/max(threenoderesults.datas_21tp{1,i}.YP2))
hold on
end
xlabel('Time(AU)')
ylabel('Normalized data(AU)')
title({"Random Connection 3-Node Model", "Normalized Data"}, 'FontSize',11)
ylim([0 1.05])
res = 300;
set(gcf,'paperunits','inches','PaperPosition',[0 0 2.7 2.7]);
print('S2B.tiff','-dtiff',['-r' num2str(res)]);

%% Supplement 2C
clear;
load('param3wrapper_paper.mat')
count=50;
bot=0;
top=0.5;
actual_param=threenoderesults.actual_param;
[min3,sum0_3,sum1_3,sum2_3,sum5_3]=fig3func_ranges(actual_param,threenoderesults.predicted_param_3tp,threenoderesults.SSEs_3tp,count,bot,top);
[min7,sum0_7,sum1_7,sum2_7,sum5_7]=fig3func_ranges(actual_param,threenoderesults.predicted_param_7tp,threenoderesults.SSEs_7tp,count,bot,top);
[min11,sum0_11,sum1_11,sum2_11,sum5_11]=fig3func_ranges(actual_param,threenoderesults.predicted_param_11tp,threenoderesults.SSEs_11tp,count,bot,top);
[min21,sum0_21,sum1_21,sum2_21,sum5_21]=fig3func_ranges(actual_param,threenoderesults.predicted_param_21tp,threenoderesults.SSEs_21tp,count,bot,top);
d=figure; 
in=roundn([sum0_3,sum1_3,sum2_3,sum5_3;sum0_7,sum1_7,sum2_7,sum5_7;sum0_11,sum1_11,sum2_11,sum5_11;sum0_21,sum1_21,sum2_21,sum5_21],-2);
hm=heatmap(in');
caxis([0.5 1])
hm.Colormap=hot;
hm.YData = ["No Noise" "10:1 S:N" "5:1 S:N" "2:1 S:N"];
hm.XData = ["3 Timepoints" "7 Timepoints" "11 Timepoints" "21 Timepoints"];
hm.Title="Classification Accuracy";
hm.FontSize=11;
res = 300;
set(gcf,'paperunits','inches','PaperPosition',[0 0 2.8 2.5]);
print('S2C.tiff','-dtiff',['-r' num2str(res)]);
%% Supplement 2D
clear;
load('param3wrapper_paper.mat')
count=50;
bot=0.5;
top=1;
actual_param=threenoderesults.actual_param;
[min3,sum0_3,sum1_3,sum2_3,sum5_3]=fig3func_ranges(actual_param,threenoderesults.predicted_param_3tp,threenoderesults.SSEs_3tp,count,bot,top);
[min7,sum0_7,sum1_7,sum2_7,sum5_7]=fig3func_ranges(actual_param,threenoderesults.predicted_param_7tp,threenoderesults.SSEs_7tp,count,bot,top);
[min11,sum0_11,sum1_11,sum2_11,sum5_11]=fig3func_ranges(actual_param,threenoderesults.predicted_param_11tp,threenoderesults.SSEs_11tp,count,bot,top);
[min21,sum0_21,sum1_21,sum2_21,sum5_21]=fig3func_ranges(actual_param,threenoderesults.predicted_param_21tp,threenoderesults.SSEs_21tp,count,bot,top);
d=figure; 
in=roundn([sum0_3,sum1_3,sum2_3,sum5_3;sum0_7,sum1_7,sum2_7,sum5_7;sum0_11,sum1_11,sum2_11,sum5_11;sum0_21,sum1_21,sum2_21,sum5_21],-2);
hm=heatmap(in');
caxis([0.5 1])
hm.Colormap=hot;
hm.YData = ["No Noise" "10:1 S:N" "5:1 S:N" "2:1 S:N"];
hm.XData = ["3 Timepoints" "7 Timepoints" "11 Timepoints" "21 Timepoints"];
hm.Title="Classification Accuracy";
hm.FontSize=11;
res = 300;
set(gcf,'paperunits','inches','PaperPosition',[0 0 2.8 2.5]);
print('S2D.tiff','-dtiff',['-r' num2str(res)]);
%% Supplement 2E
clear;
load('param3wrapper_paper.mat')
count=50;
bot=1;
top=1.5;
actual_param=threenoderesults.actual_param;
[min3,sum0_3,sum1_3,sum2_3,sum5_3]=fig3func_ranges(actual_param,threenoderesults.predicted_param_3tp,threenoderesults.SSEs_3tp,count,bot,top);
[min7,sum0_7,sum1_7,sum2_7,sum5_7]=fig3func_ranges(actual_param,threenoderesults.predicted_param_7tp,threenoderesults.SSEs_7tp,count,bot,top);
[min11,sum0_11,sum1_11,sum2_11,sum5_11]=fig3func_ranges(actual_param,threenoderesults.predicted_param_11tp,threenoderesults.SSEs_11tp,count,bot,top);
[min21,sum0_21,sum1_21,sum2_21,sum5_21]=fig3func_ranges(actual_param,threenoderesults.predicted_param_21tp,threenoderesults.SSEs_21tp,count,bot,top);
d=figure; 
in=roundn([sum0_3,sum1_3,sum2_3,sum5_3;sum0_7,sum1_7,sum2_7,sum5_7;sum0_11,sum1_11,sum2_11,sum5_11;sum0_21,sum1_21,sum2_21,sum5_21],-2);
hm=heatmap(in');
caxis([0.5 1])
hm.Colormap=hot;
hm.YData = ["No Noise" "10:1 S:N" "5:1 S:N" "2:1 S:N"];
hm.XData = ["3 Timepoints" "7 Timepoints" "11 Timepoints" "21 Timepoints"];
hm.Title="Classification Accuracy";
hm.FontSize=11;
res = 300;
set(gcf,'paperunits','inches','PaperPosition',[0 0 2.8 2.5]);
print('S2E.tiff','-dtiff',['-r' num2str(res)]);
%% Supplement 2F
clear;
load('param3wrapper_paper.mat')
count=50;
bot=1.5;
top=2;
actual_param=threenoderesults.actual_param;
[min3,sum0_3,sum1_3,sum2_3,sum5_3]=fig3func_ranges(actual_param,threenoderesults.predicted_param_3tp,threenoderesults.SSEs_3tp,count,bot,top);
[min7,sum0_7,sum1_7,sum2_7,sum5_7]=fig3func_ranges(actual_param,threenoderesults.predicted_param_7tp,threenoderesults.SSEs_7tp,count,bot,top);
[min11,sum0_11,sum1_11,sum2_11,sum5_11]=fig3func_ranges(actual_param,threenoderesults.predicted_param_11tp,threenoderesults.SSEs_11tp,count,bot,top);
[min21,sum0_21,sum1_21,sum2_21,sum5_21]=fig3func_ranges(actual_param,threenoderesults.predicted_param_21tp,threenoderesults.SSEs_21tp,count,bot,top);
d=figure; 
in=roundn([sum0_3,sum1_3,sum2_3,sum5_3;sum0_7,sum1_7,sum2_7,sum5_7;sum0_11,sum1_11,sum2_11,sum5_11;sum0_21,sum1_21,sum2_21,sum5_21],-2);
hm=heatmap(in');
caxis([0.5 1])
hm.Colormap=hot;
hm.YData = ["No Noise" "10:1 S:N" "5:1 S:N" "2:1 S:N"];
hm.XData = ["3 Timepoints" "7 Timepoints" "11 Timepoints" "21 Timepoints"];
hm.Title="Classification Accuracy";
hm.FontSize=11;
res = 300;
set(gcf,'paperunits','inches','PaperPosition',[0 0 2.8 2.5]);
print('S2F.tiff','-dtiff',['-r' num2str(res)]);
%% Supplement 3A
clear;
load('param2wrapper_paper.mat')
params=zeros(1,8);
j=0;
k=0;
for i=1:50
        param=twonoderesults.actual_param{1,i};
        Jacob=[param(3),param(4);param(7),param(8)];
        eigen=eigs(Jacob);
        imag_eigen=imag(eigen(1));
        if abs(imag_eigen)>0
            j=j+1;
            actual_param_os{1,j}=twonoderesults.actual_param{1,i};
            for k=1:4
            predicted_param_3tp_os{k,j}=twonoderesults.predicted_param_3tp{k,i};
            predicted_param_7tp_os{k,j}=twonoderesults.predicted_param_7tp{k,i};
            predicted_param_11tp_os{k,j}=twonoderesults.predicted_param_11tp{k,i};
            predicted_param_21tp_os{k,j}=twonoderesults.predicted_param_21tp{k,i};
            SSEs_3tp_os{k,j}=twonoderesults.SSEs_3tp{k,i};
            SSEs_7tp_os{k,j}=twonoderesults.SSEs_7tp{k,i};
            SSEs_11tp_os{k,j}=twonoderesults.SSEs_11tp{k,i};
            SSEs_21tp_os{k,j}=twonoderesults.SSEs_21tp{k,i};
            end
        end
end 

count=j;
bot=0;
top=2;
[min3,sum0_3,sum1_3,sum2_3,sum5_3]=fig3func_ranges(actual_param_os,predicted_param_3tp_os,SSEs_3tp_os,count,bot,top);
[min7,sum0_7,sum1_7,sum2_7,sum5_7]=fig3func_ranges(actual_param_os,predicted_param_7tp_os,SSEs_7tp_os,count,bot,top);
[min11,sum0_11,sum1_11,sum2_11,sum5_11]=fig3func_ranges(actual_param_os,predicted_param_11tp_os,SSEs_11tp_os,count,bot,top);
[min21,sum0_21,sum1_21,sum2_21,sum5_21]=fig3func_ranges(actual_param_os,predicted_param_21tp_os,SSEs_21tp_os,count,bot,top);

d=figure; 
in=roundn([sum0_3,sum1_3,sum2_3,sum5_3;sum0_7,sum1_7,sum2_7,sum5_7;sum0_11,sum1_11,sum2_11,sum5_11;sum0_21,sum1_21,sum2_21,sum5_21],-2);
hm=heatmap(in');
caxis([0.5 1])
hm.Colormap=hot;
hm.YData = ["No Noise" "10:1 S:N" "5:1 S:N" "2:1 S:N"];
hm.XData = ["3 Timepoints" "7 Timepoints" "11 Timepoints" "21 Timepoints"];
hm.Title="Classification Accuracy";
hm.FontSize=11;
res = 300;
set(gcf,'paperunits','inches','PaperPosition',[0 0 2.8 2.5]);
print('S3A.tiff','-dtiff',['-r' num2str(res)]);

%% Supplement 3B
load('param3wrapper_paper.mat')
params=zeros(1,15);
j=0;
k=0;
for i=1:50
        param=threenoderesults.actual_param{1,i};
        Jacob=[param(3),param(4),param(5);param(8),param(9),param(10);param(13),param(14),param(15)];
        eigen=eigs(Jacob);
        imag_eigen=imag(eigen(1));
        if abs(imag_eigen)>0
            j=j+1;
            actual_param_os{1,j}=threenoderesults.actual_param{1,i};
            for k=1:4
            predicted_param_3tp_os{k,j}=threenoderesults.predicted_param_3tp{k,i};
            predicted_param_7tp_os{k,j}=threenoderesults.predicted_param_7tp{k,i};
            predicted_param_11tp_os{k,j}=threenoderesults.predicted_param_11tp{k,i};
            predicted_param_21tp_os{k,j}=threenoderesults.predicted_param_21tp{k,i};
            SSEs_3tp_os{k,j}=threenoderesults.SSEs_3tp{k,i};
            SSEs_7tp_os{k,j}=threenoderesults.SSEs_7tp{k,i};
            SSEs_11tp_os{k,j}=threenoderesults.SSEs_11tp{k,i};
            SSEs_21tp_os{k,j}=threenoderesults.SSEs_21tp{k,i};
            end
        end
end 

count=j;
bot=0;
top=2;
[min3,sum0_3,sum1_3,sum2_3,sum5_3]=fig3func_ranges(actual_param_os,predicted_param_3tp_os,SSEs_3tp_os,count,bot,top);
[min7,sum0_7,sum1_7,sum2_7,sum5_7]=fig3func_ranges(actual_param_os,predicted_param_7tp_os,SSEs_7tp_os,count,bot,top);
[min11,sum0_11,sum1_11,sum2_11,sum5_11]=fig3func_ranges(actual_param_os,predicted_param_11tp_os,SSEs_11tp_os,count,bot,top);
[min21,sum0_21,sum1_21,sum2_21,sum5_21]=fig3func_ranges(actual_param_os,predicted_param_21tp_os,SSEs_21tp_os,count,bot,top);

d=figure; 
in=roundn([sum0_3,sum1_3,sum2_3,sum5_3;sum0_7,sum1_7,sum2_7,sum5_7;sum0_11,sum1_11,sum2_11,sum5_11;sum0_21,sum1_21,sum2_21,sum5_21],-2);
hm=heatmap(in');
caxis([0.5 1])
hm.Colormap=hot;
hm.YData = ["No Noise" "10:1 S:N" "5:1 S:N" "2:1 S:N"];
hm.XData = ["3 Timepoints" "7 Timepoints" "11 Timepoints" "21 Timepoints"];
hm.Title="Classification Accuracy";
hm.FontSize=11;
res = 300;
set(gcf,'paperunits','inches','PaperPosition',[0 0 2.8 2.5]);
print('S3B.tiff','-dtiff',['-r' num2str(res)]);
%% Generate data for Supplement 4
% Skip to next section to use generated data
count=50;
load("param3wrapper_paper.mat");
multistart=10;
for j=1:4
for i=1:count
timeint=1;
data_2node.Y=threenoderesults.datas_11tp{j,i}.Y(:,1:2);
data_2node.YP1=threenoderesults.datas_11tp{j,i}.YP1(:,1:2);
data_2node.YP2=threenoderesults.datas_11tp{j,i}.YP2(:,1:2);
data_2node.T=threenoderesults.datas_11tp{j,i}.T;
for k=1:multistart
[predicted_param_2node{j,i}(k,:),SSEs_2node{j,i}(k,1)]=estimate2node(data_2node,timeint);
end
paramtheor(i,1:4)=threenoderesults.actual_param{1,i}(1,1:4);
paramtheor(i,5:8)=threenoderesults.actual_param{1,i}(1,6:9);
end
end
for j=1:4
    for i=1:count
        [M,I] = min(SSEs_2node{j,i});
predicted_param_2node_best{j,1}(i,:)=predicted_param_2node{j,i}(I,:);
    end
normalized_params{j,1}=predicted_param_2node_best{j,1}./paramtheor;
end
%% Figure for Supplement 4
figure
subplot(2,2,1)
boxplot(normalized_params{1,1});
set(gca,'TickLabelInterpreter', 'tex');
set(gca, 'Fontsize', 8, 'Fontweight', 'bold');
set(gca,'xticklabel',{'S_1_,_b',"S_1_,_e_x","F_1_1","F_1_2","S_2_,_b","S_2_,_e_x","F_2_1","F_2_2"})
title({"No Noise", "Classification Accuracy=94.75%"}, 'FontSize',10)
ylim([-8 10])
xlim=get(gca,'xlim');
hold on
plot(xlim,[0 0])

subplot(2,2,2)
boxplot(normalized_params{2,1});
set(gca,'TickLabelInterpreter', 'tex');
set(gca, 'Fontsize', 8, 'Fontweight', 'bold');
set(gca,'xticklabel',{'S_1_,_b',"S_1_,_e_x","F_1_1","F_1_2","S_2_,_b","S_2_,_e_x","F_2_1","F_2_2"})
title({"2:1 Signal:Noise", "Classification Accuracy=93.75%"}, 'FontSize',10)
ylim([-8 10])
xlim=get(gca,'xlim');
hold on
plot(xlim,[0 0])

subplot(2,2,3)
boxplot(normalized_params{3,1});
set(gca,'TickLabelInterpreter', 'tex');
set(gca, 'Fontsize', 8, 'Fontweight', 'bold');
set(gca,'xticklabel',{'S_1_,_b',"S_1_,_e_x","F_1_1","F_1_2","S_2_,_b","S_2_,_e_x","F_2_1","F_2_2"})
title({"5:1 Signal:Noise", "Classification Accuracy=91.25%"}, 'FontSize',10)
ylim([-8 10])
xlim=get(gca,'xlim');
hold on
plot(xlim,[0 0])


subplot(2,2,4)
boxplot(normalized_params{4,1});
set(gca,'TickLabelInterpreter', 'tex');
set(gca, 'Fontsize', 8, 'Fontweight', 'bold');
set(gca,'xticklabel',{'S_1_,_b',"S_1_,_e_x","F_1_1","F_1_2","S_2_,_b","S_2_,_e_x","F_2_1","F_2_2"})
title({"2:1 Signal:Noise", "Classification Accuracy=87%"}, 'FontSize',10)
ylim([-8 10])
xlim=get(gca,'xlim');
hold on
plot(xlim,[0 0])

size = [6.75 8.75];
res = 300;
set(gcf,'paperunits','inches','paperposition',[0 0 size]);
print('S4.tiff','-dtiff',['-r' num2str(res)]);

%% count accurately mapped parameters
load("normalized_params.mat")
count=50;
for j=1:4
    s=sign(normalized_params{j,1});
    % look at ipositif for mapping accuracy for each noise level
    ipositif{j,1}=sum(s(:)==1)/4;
    
end
%% Supplement S5
load ("param3wrapper_paper.mat")
count=50;

for j=1:4
    k=1;
    for i=1:count
        [M,I] = min(threenoderesults.SSEs_11tp{j,i});
        all_SSEs_diff{j,1}(k:k+10-1,1)= (threenoderesults.SSEs_11tp{j,i}-threenoderesults.SSEs_11tp{j,i}(I,:));
        k=k+10;
    end
    all_SSEs_diff_sorted{j,1}=sort(all_SSEs_diff{j,1});
end
for j=1:4
        for i=1:count
            [M,I] = min(threenoderesults.SSEs_11tp{j,i});
            k=1;
            for m=1:10
%           if SSEs_3node_norm{j,1}(m,i)<2
                if threenoderesults.SSEs_11tp{j,i}(m,1)<2*threenoderesults.SSEs_11tp{j,i}(I,1)||threenoderesults.SSEs_11tp{j,i}(m,1)<0.0001
                    best_params{j,i}(k,1:15)=threenoderesults.predicted_param_11tp{j,i}(m,:);
                    k=k+1;
                end
            end
        end
end
for j=1:4
    k=1;
    for i=1:count
          [M,I] = min(threenoderesults.SSEs_11tp{j,i});
          X=(threenoderesults.predicted_param_11tp{j,i}(I,:))';
          cov{j,1}(i,:)=std(best_params{j,i})./(mean(best_params{j,i}));
    end
    cov_col{j,1}=reshape(cov{j,1},[],1);
    hold on
end
edges = linspace(-10, 10, 41); % Create 10 bins.
subplot(2,2,1)
h1=histogram(cov{1,1},'BinEdges',edges,'EdgeColor', 'black', 'FaceColor',  'black');
title('No Noise','Fontsize',10);
subplot(2,2,2)
h1=histogram(cov{2,1},'BinEdges',edges,'EdgeColor', 'black', 'FaceColor',  'green');
title('10:1 Signal:Noise','Fontsize',10);

subplot(2,2,3)
h1=histogram(cov{3,1},'BinEdges',edges,'EdgeColor', 'black', 'FaceColor',  'blue');
title('5:1 Signal:Noise','Fontsize',10);

subplot(2,2,4)
h1=histogram(cov{4,1},'BinEdges',edges,'EdgeColor', 'black', 'FaceColor',  'red');
title('2:1 Signal:Noise','Fontsize',10);

size = [6.75 8.75];
res = 300;
set(gcf,'paperunits','inches','paperposition',[0 0 size]);
print('S5.tiff','-dtiff',['-r' num2str(res)]);
```
