## Supplementary Code for "Network Inference from Perturbation Time Course Data": min_LS2node.m

```
function fmin=min_LS2node(data,timeint)
finSSE=zeros(1,100);
tic
Y=data.Y;
YP1=data.YP1;
YP2=data.YP2;
t=data.T;

if length(t)>10
    poly_order=9;
else
    poly_order=length(t)-1;
end

poly_YP1=polyfit(t,YP1(:,1),poly_order);
poly_YP2=polyfit(t,YP2(:,2),poly_order);

        initial = [1 0 0 1 0 0];
        options = optimoptions('fminunc','MaxFunEval',6000);
        conoptions = optimoptions('fmincon','MaxFunctionEvaluations',10000);
        [fmin,finSSE(1,1)] = fminunc(@nestedfun,initial,options);
        display(fmin);
        toc
        
function SSE = nestedfun(r2)
        
    % Solve the ODEs 
           
[t,guessY] = ode15s(@odeqns,0:timeint:10,[0;0],odeset('RelTol',1e-6),r2);
[t,guessYP1] = ode15s(@odep1,0:timeint:10,[0],odeset('RelTol',1e-6),r2);
[t,guessYP2] = ode15s(@odep2,0:timeint:10,[0],odeset('RelTol',1e-6),r2);

    SSE = (sum(sum((Y-guessY).^2)+sum((YP1(:,2)-guessYP1).^2)+sum((YP2(:,1)-guessYP2).^2))).^0.5;

%     SSE = (sum(sum((Y-guessY).^2))).^0.5;
    
    function dzdt1 = odeqns(t,z,r)
        dzdt1 = [
            (r(1)+r(2)*z(1)+r(3)*z(2))
            (r(4)+r(5)*z(1)+r(6)*z(2))
        ];
    end
    function dzdt1 = odep1(t,z,r2)
         Y_pert=polyval(poly_YP1,t);
%         Y_pert=polyval([f3_YP1.p1,f3_YP1.p2,f3_YP1.p3,f3_YP1.p4,f3_YP1.p5,f3_YP1.p6,f3_YP1.p7,f3_YP1.p8,f3_YP1.p9,f3_YP1.p10],t);
%         Y_pert=polyval([f3_YP1.p1,f3_YP1.p2,f3_YP1.p3],t);

        dzdt1 = [
            (r2(4)+r2(5)*Y_pert+r2(6)*z)
                 ];
    end
       function dzdt1 = odep2(t,z,r2)
        Y_pert=polyval(poly_YP2,t);
%         Y_pert=polyval([f3_YP2.p1,f3_YP2.p2,f3_YP2.p3,f3_YP2.p4,f3_YP2.p5,f3_YP2.p6,f3_YP2.p7,f3_YP2.p8,f3_YP2.p9,f3_YP2.p10],t);
%           Y_pert=polyval([f3_YP2.p1,f3_YP2.p2,f3_YP2.p3],t);

        dzdt1 = [
            (r2(1)+r2(2)*z+r2(3)*Y_pert)
        ];
       end
        
end
end
```
