## Supplementary Code for "Network Inference from Perturbation Time Course Data": twonodewrapper.m

```
function [twonoderesults] = twonodewrapper_parfor(count,tmax,y0,pert1,pert2,multistart)

tic
for i = 1:count
    
    timeint=1;
    counter = 1;
    disp(i);
    good = 0;
    
    param = zeros(1,6);
    initact = zeros(1,2);
    
    while(~good)
        
        initact(1) = 0;
        initact(2) = 0;
        
        param(1) = 2*rand(1);
        param(2) = 2*rand(1);
        param(3) = 4*rand(1)-2;
        param(4) = 4*rand(1)-2;
        param(5) = 2*rand(1);
        param(6) = 2*rand(1);
        param(7) = 4*rand(1)-2;
        param(8) = 4*rand(1)-2;
        
        data_nostim=eval_data_2node_polyval_nostim(timeint,100,param,y0,y0,y0,pert1,pert2,0);
        
        y_0=[data_nostim.Y(101,1),data_nostim.Y(101,2)];
        y_p1=[data_nostim.YP1(101,1),data_nostim.YP1(101,2)];
        y_p2=[data_nostim.YP2(101,1),data_nostim.YP2(101,2)];
        
        data=eval_data_2node_polyval(timeint,tmax,param,y_0,y_p1,y_p2,pert1,pert2,0);

        topval = max([max(abs(data.Y)) max(abs(data.YP1)) max(abs(data.YP2))]);
        
        Jacob=[param(3),param(4);param(7),param(8)];
        eigen=eigs(Jacob);    
        
        if eigen <= 0 & topval <= 10 
            good = 1;
        end           
        
        counter = counter+1;
        
    end 


display(param);  
actual_param{1,i}=param;

timeint=5;
data0_3tp=eval_data_2node_polyval(timeint,tmax,param,y_0,y_p1,y_p2,pert1,pert2,0);
data10_3tp=eval_data_2node_polyval(timeint,tmax,param,y_0,y_p1,y_p2,pert1,pert2,0.10);
data20_3tp=eval_data_2node_polyval(timeint,tmax,param,y_0,y_p1,y_p2,pert1,pert2,0.20);
data50_3tp=eval_data_2node_polyval(timeint,tmax,param,y_0,y_p1,y_p2,pert1,pert2,0.50);

timeint=5/3;
data0_7tp=eval_data_2node_polyval(timeint,tmax,param,y_0,y_p1,y_p2,pert1,pert2,0);
data10_7tp=eval_data_2node_polyval(timeint,tmax,param,y_0,y_p1,y_p2,pert1,pert2,0.10);
data20_7tp=eval_data_2node_polyval(timeint,tmax,param,y_0,y_p1,y_p2,pert1,pert2,0.20);
data50_7tp=eval_data_2node_polyval(timeint,tmax,param,y_0,y_p1,y_p2,pert1,pert2,0.50);

timeint=1;
data0_11tp=eval_data_2node_polyval(timeint,tmax,param,y_0,y_p1,y_p2,pert1,pert2,0);
data10_11tp=eval_data_2node_polyval(timeint,tmax,param,y_0,y_p1,y_p2,pert1,pert2,0.10);
data20_11tp=eval_data_2node_polyval(timeint,tmax,param,y_0,y_p1,y_p2,pert1,pert2,0.20);
data50_11tp=eval_data_2node_polyval(timeint,tmax,param,y_0,y_p1,y_p2,pert1,pert2,0.50);

timeint=0.5;
data0_21tp=eval_data_2node_polyval(timeint,tmax,param,y_0,y_p1,y_p2,pert1,pert2,0);
data10_21tp=eval_data_2node_polyval(timeint,tmax,param,y_0,y_p1,y_p2,pert1,pert2,0.10);
data20_21tp=eval_data_2node_polyval(timeint,tmax,param,y_0,y_p1,y_p2,pert1,pert2,0.20);
data50_21tp=eval_data_2node_polyval(timeint,tmax,param,y_0,y_p1,y_p2,pert1,pert2,0.50);

datas_3tp{1,i}=data0_3tp;
datas_3tp{2,i}=data10_3tp;
datas_3tp{3,i}=data20_3tp;
datas_3tp{4,i}=data50_3tp;

datas_7tp{1,i}=data0_7tp;
datas_7tp{2,i}=data10_7tp;
datas_7tp{3,i}=data20_7tp;
datas_7tp{4,i}=data50_7tp;

datas_11tp{1,i}=data0_11tp;
datas_11tp{2,i}=data10_11tp;
datas_11tp{3,i}=data20_11tp;
datas_11tp{4,i}=data50_11tp;

datas_21tp{1,i}=data0_21tp;
datas_21tp{2,i}=data10_21tp;
datas_21tp{3,i}=data20_21tp;
datas_21tp{4,i}=data50_21tp;

end

for i=1:count
% 3 timepoints
for j=1:4
        for k=1:multistart
            timeint=5;
            [predicted_param_3tp{j,i}(k,:),SSEs_3tp{j,i}(k,1)]=estimate2node (datas_3tp{j,i},timeint);
        
    
% 7 timepoints
timeint=5/3;
    
            [predicted_param_7tp{j,i}(k,:),SSEs_7tp{j,i}(k,1)]=estimate2node (datas_7tp{j,i},timeint);
     
% 11 timepoints
timeint=1;
    
            [predicted_param_11tp{j,i}(k,:),SSEs_11tp{j,i}(k,1)]=estimate2node (datas_11tp{j,i},timeint);
            
% 21 timepoints
timeint=0.5;
            [predicted_param_21tp{j,i}(k,:),SSEs_21tp{j,i}(k,1)]=estimate2node (datas_21tp{j,i},timeint);
        end
    end
    
end

twonoderesults.actual_param=actual_param;

twonoderesults.datas_3tp=datas_3tp;
twonoderesults.predicted_param_3tp=predicted_param_3tp;
twonoderesults.SSEs_3tp=SSEs_3tp;

twonoderesults.datas_7tp=datas_7tp;
twonoderesults.predicted_param_7tp=predicted_param_7tp;
twonoderesults.SSEs_7tp=SSEs_7tp;

twonoderesults.datas_11tp=datas_11tp;
twonoderesults.predicted_param_11tp=predicted_param_11tp;
twonoderesults.SSEs_11tp=SSEs_11tp;

twonoderesults.datas_21tp=datas_21tp;
twonoderesults.predicted_param_21tp=predicted_param_21tp;
twonoderesults.SSEs_21tp=SSEs_21tp;


    %save ('param2wrapper.mat', 'twonoderesults')
    toc
end
```
