## Supplementary Code for "Network Inference from Perturbation Time Course Data": Untitled.m

```
%% plot all unperturbed data
for i=1:50
    plot(twonoderesults.datas_11tp{1,i}.T,twonoderesults.datas_11tp{1,i}.Y/max(twonoderesults.datas_11tp{1,i}.Y))
hold on
end
xlabel('Time(AU)')
ylabel('Normalized data(AU)')


figure
for i=1:50
    plot(threenoderesults.datas_11tp{1,i}.T,threenoderesults.datas_11tp{1,i}.Y/max(threenoderesults.datas_11tp{1,i}.Y))
hold on
end
xlabel('Time(AU)')
ylabel('Normalized data(AU)')

%% plot oscillatory data
%% Supplement 3A
clear;
load('param2wrapper_paper.mat')
params=zeros(1,8);
j=0;
k=0;
for i=1:50
        param=twonoderesults.actual_param{1,i};
        Jacob=[param(3),param(4);param(7),param(8)];
        eigen=eigs(Jacob);
        imag_eigen=imag(eigen(1));
        if abs(imag_eigen)>0
            j=j+1;
            plot(twonoderesults.datas_21tp{1,i}.T,twonoderesults.datas_21tp{1,i}.Y/max(twonoderesults.datas_21tp{1,i}.Y))
        hold on
        end
end

%% Supplement 3A
clear;
load('param3wrapper_paper.mat')
params=zeros(1,8);
j=0;
k=0;
for i=1:50
        param=threenoderesults.actual_param{1,i};
        Jacob=[param(3),param(4),param(5);param(8),param(9),param(10);param(13),param(14),param(15)];
        eigen=eigs(Jacob);
        imag_eigen=imag(eigen(1));
        if abs(imag_eigen)>0
            j=j+1;
        plot(threenoderesults.datas_11tp{1,i}.T,threenoderesults.datas_11tp{1,i}.Y/max(threenoderesults.datas_11tp{1,i}.Y))
        hold on
        end
end
```
