## Supplementary Code for "Network Inference from Perturbation Time Course Data": ~WRL0725.tmp

To reproduce analysis and figures, run the script “Make_Figures.m”. We recommend that you run the file in sections, as delineated in the file. Some segments that take a long time to run are either commented out or delineated as a new section. To simplify this, we provide all data needed to recreate the figures as mat files. 

For figure 5-6, models each of the 16 models is assigned a binary code. There are four separate options that distinguish the FFL structures and they can be classified as a four-digit binary code (Table 1): [C_1_ C_2_ C_3_ C_4_], C_i_ ϵ {1,2}. C_1_ specifies if x_1_ activates (C_1_ = 1) or inhibits (C_1_ = 2) x_2_. C_2_ determines if the combined effects of x_1_ and x_2_ on x_3_ act via an AND gate (C_2_ = 1) or an OR gate (C_2_ = 2). C_3_ specifies if x_1_ activates (C_3_ = 1) or inhibits (C_3_ = 2) x_3_. Lastly, C_4_ specifies if x_2_ activates (C_4_ = 1) or inhibits (C_4_ = 2) x_3_. An AND gate dictates that the effects of x_1_ and x_2_ on x_3_ are coupled: any effect on x_3_ requires both x_1_ and x_2_ to be present. An OR gate implies these effects are not coupled, although competitive inhibition still plays a role in the modeling of each effect.  The parameters F_12_, F_13_, and F_23_ remain fixed at zero in all models; however, in estimation we allow them to be non-zero should the algorithm make such incorrect inference.
