## Supplementary Code for "Network Inference from Perturbation Time Course Data": dmrathreeanalyzer.m

```
function [theorthreejacdata,obsthreejacdata,ssjac] = dmrathreeanalyzer(perturb,noise,timeint,inputdata,inputdata50pert)

opts.TRANSA=true;

theorthreejacdata = [-1 0 0; 0 -1 0; 0 0 -1];
%theorthreejacdata = calcthreecomplexJacobian(timeint);

%{
[T,Y0] = ode45(@threeODE,0:timeint:10, [0 0 0 0 0 0]);

for i = 1:length(T)
    for j = 1:3
        randomdata = normrnd(0,noise,1,1);
        Y0(i,j) = Y0(i,j) + randomdata;
    end
end

[T,YP1] = ode45(@threeODE,0:timeint:10, [0 0 0 10*perturb 0 0]);

for i = 1:length(T)
    for j = 1:3
        randomdata = normrnd(0,noise,1,1);
        YP1(i,j) = YP1(i,j) + randomdata;
    end
end

[T,YP2] = ode45(@threeODE,0:timeint:10, [0 0 0 0 10*perturb 0]);

for i = 1:length(T)
    for j = 1:3
        randomdata = normrnd(0,noise,1,1);
        YP2(i,j) = YP2(i,j) + randomdata;
    end
end


[T,YP3] = ode45(@threeODE,0:timeint:10, [0 0 0 0 0 10*perturb]);

for i = 1:length(T)
    for j = 1:3
        randomdata = normrnd(0,noise,1,1);
        YP3(i,j) = YP3(i,j) + randomdata;
    end
end
%}

Y0 = inputdata.Y;
YP1 = inputdata.YP1;
YP2 = inputdata.YP2;
YP3 = inputdata.YP3;

Y0pert50 = inputdata50pert.Y;
YP1pert50 = inputdata50pert.YP1;
YP2pert50 = inputdata50pert.YP2;
YP3pert50 = inputdata50pert.YP3;


% T = 0:(size(inputdata.Y,1)-1);
T = 0:timeint:10;

R111 = YP2pert50(:,1) - Y0(:,1);
R112 = YP2(:,1) - Y0(:,1);
R113 = YP3(:,1) - Y0(:,1);
R121 = YP2pert50(:,2) - Y0(:,2);
R122 = YP2(:,2) - Y0(:,2);
R123 = YP3(:,2) - Y0(:,2);
R131 = YP2pert50(:,3) - Y0(:,3);
R132 = YP2(:,3) - Y0(:,3);
R133 = YP3(:,3) - Y0(:,3);

for i = 1:(length(T)-1)
    p11(i) = (R111(i+1)-R111(i))/(T(i+1)-T(i));
    p12(i) = (R112(i+1)-R112(i))/(T(i+1)-T(i));
    p13(i) = (R113(i+1)-R113(i))/(T(i+1)-T(i));    
end

obsthreejacdata = zeros(3,3,(length(T)-1));

for i = 1:(length(T)-1)
    p1 = [p11(i) p12(i) p13(i)];
    R1 = [R111(i) R112(i) R113(i); R121(i) R122(i) R123(i); R131(i) R132(i) R133(i)];
    
    %p1 = [p11(i) p12(i) p14(i)];
    %R1 = [R111(i) R112(i) R114(i); R121(i) R122(i) R124(i); R141(i) R142(i) R144(i)];
    
    %jacvector = p1/R1;
    jacvector = linsolve(R1,p1',opts);
    %jacdata(1,:) = jacvector';
    %obsmrnajacdata(1,:,i) = jacdata(1,:);
    obsthreejacdata(1,:,i) = jacvector';
    %obsmrnajacdata(1,1,i) = jacvector(1);
    %obsmrnajacdata(1,2,i) = jacvector(2);
    %obsmrnajacdata(1,3,i) = 0;
    %obsmrnajacdata(1,4,i) = jacvector(3);
end

R211 = YP1(:,1) - Y0(:,1);
R212 = YP3pert50(:,1) - Y0(:,1);
R213 = YP3(:,1) - Y0(:,1);
R221 = YP1(:,2) - Y0(:,2);
R222 = YP3pert50(:,2) - Y0(:,2);
R223 = YP3(:,2) - Y0(:,2);
R231 = YP1(:,3) - Y0(:,3);
R232 = YP3pert50(:,3) - Y0(:,3);
R233 = YP3(:,3) - Y0(:,3);

for i = 1:(length(T)-1)
    p21(i) = (R221(i+1)-R221(i))/(T(i+1)-T(i));
    p22(i) = (R222(i+1)-R222(i))/(T(i+1)-T(i));
    p23(i) = (R223(i+1)-R223(i))/(T(i+1)-T(i));
end

for i = 1:(length(T)-1)
    p2 = [p21(i) p22(i) p23(i)];
    R2 = [R211(i) R212(i) R213(i); R221(i) R222(i) R223(i); R231(i) R232(i) R233(i)];
    %p2 = [p22(i) p24(i)];
    %R2 = [R222(i) R224(i); R242(i) R244(i)];    
    %jacvector = p2/R2;
    jacvector = linsolve(R2,p2',opts);
    %jacdata(2,:) = jacvector';
    %obsmrnajacdata(2,:,i) = jacdata(2,:);
    obsthreejacdata(2,:,i) = jacvector';
    %obsmrnajacdata(2,1,i) = 0;
    %obsmrnajacdata(2,2,i) = jacvector(1);
    %obsmrnajacdata(2,3,i) = 0;
    %obsmrnajacdata(2,4,i) = jacvector(2);
end


R311 = YP1(:,1) - Y0(:,1);
R312 = YP2(:,1) - Y0(:,1);
R313 = YP1pert50(:,1) - Y0(:,1);
R321 = YP1(:,2) - Y0(:,2);
R322 = YP2(:,2) - Y0(:,2);
R323 = YP1pert50(:,2) - Y0(:,2);
R331 = YP1(:,3) - Y0(:,3);
R332 = YP2(:,3) - Y0(:,3);
R333 = YP1pert50(:,3) - Y0(:,3);

for i = 1:(length(T)-1)
    p31(i) = (R331(i+1)-R331(i))/(T(i+1)-T(i));
    p32(i) = (R332(i+1)-R332(i))/(T(i+1)-T(i));
    p33(i) = (R333(i+1)-R333(i))/(T(i+1)-T(i));
end


for i = 1:(length(T)-1)
    p3 = [p31(i) p32(i) p33(i)];
    R3 = [R311(i) R312(i) R313(i); R321(i) R322(i) R323(i); R331(i) R332(i) R333(i)];
    %p3 = [p31(i) p32(i) p33(i)];
    %R3 = [R311(i) R312(i) R313(i); R321(i) R322(i) R323(i); R331(i) R332(i) R333(i)];
    %jacvector = p3/R3;
    jacvector = linsolve(R3,p3',opts);
    %jacdata(3,:) = jacvector';
    %obsmrnajacdata(3,:,i) = jacdata(3,:);
    obsthreejacdata(3,:,i) = jacvector';
    %obsmrnajacdata(3,1,i) = jacvector(1);
    %obsmrnajacdata(3,2,i) = jacvector(2);
    %obsmrnajacdata(3,3,i) = jacvector(3);
    %obsmrnajacdata(3,4,i) = 0;
end

lasttimepoint = length(Y0);
ss(1) = Y0(lasttimepoint,1);
ss(2) = Y0(lasttimepoint,2);
ss(3) = Y0(lasttimepoint,3);

[T,YSS0] = ode45(@threeODE,0:timeint:max(T), [ss(1) ss(2) ss(3) 0 0 0]);
[T,YSS1] = ode45(@threeODE,0:timeint:max(T), [ss(1) ss(2) ss(3) 10*perturb 0 0]);
[T,YSS2] = ode45(@threeODE,0:timeint:max(T), [ss(1) ss(2) ss(3) 0 10*perturb 0]);
[T,YSS3] = ode45(@threeODE,0:timeint:max(T), [ss(1) ss(2) ss(3) 0 0 10*perturb]);

fc = zeros(3,3);
fc(:,1) = YSS1(lasttimepoint,1:3)';
fc(:,2) = YSS2(lasttimepoint,1:3)';
fc(:,3) = YSS3(lasttimepoint,1:3)';

Rp = zeros(3,3);

for i = 1:3
    for j = 1:3
        
        frac = fc(i,j)/ss(j);
        
        Rp(i,j) = 2*(frac-1)/(frac+1);
    
    end
end

Rpinv = inv(Rp);
%invdgRpinv = inv(diag(diag(Rpinv)));
dgRpinv = diag(diag(Rpinv));

ssjac = (-1)*dgRpinv\Rpinv;
end
%ssjac = (-1)*invdgRpinv*Rpinv;
```
