## Supplementary Code for "Network Inference from Perturbation Time Course Data": drop_node3_50.m

```
%%
count=50;
load("param3wrapper_paper.mat");
multistart=10;
for j=1:4
for i=1:count
timeint=1;
data_2node.Y=threenoderesults.datas_11tp{j,i}.Y(:,1:2);
data_2node.YP1=threenoderesults.datas_11tp{j,i}.YP1(:,1:2);
data_2node.YP2=threenoderesults.datas_11tp{j,i}.YP2(:,1:2);
data_2node.T=threenoderesults.datas_11tp{j,i}.T;
for k=1:multistart
[predicted_param_2node{j,i}(k,:),SSEs_2node{j,i}(k,1)]=estimate2node(data_2node,timeint);
end
paramtheor(i,1:4)=threenoderesults.actual_param{1,i}(1,1:4);
paramtheor(i,5:8)=threenoderesults.actual_param{1,i}(1,6:9);
end
end
for j=1:4
    for i=1:count
        [M,I] = min(SSEs_2node{j,i});
predicted_param_2node_best{j,1}(i,:)=predicted_param_2node{j,i}(I,:);
    end
normalized_params{j,1}=predicted_param_2node_best{j,1}./paramtheor;
end

%%
figure
subplot(2,2,1)
boxplot(normalized_params{1,1});
set(gca,'TickLabelInterpreter', 'tex');
set(gca, 'Fontsize', 8, 'Fontweight', 'bold');
set(gca,'xticklabel',{'S_1_,_b',"S_1_,_e_x","F_1_1","F_1_2","S_2_,_b","S_2_,_e_x","F_2_1","F_2_2"})
title({"No Noise", "Classification Accuracy=94.75%"}, 'FontSize',10)
ylim([-8 10])
xlim=get(gca,'xlim');
hold on
plot(xlim,[0 0])

subplot(2,2,2)
boxplot(normalized_params{2,1});
set(gca,'TickLabelInterpreter', 'tex');
set(gca, 'Fontsize', 8, 'Fontweight', 'bold');
set(gca,'xticklabel',{'S_1_,_b',"S_1_,_e_x","F_1_1","F_1_2","S_2_,_b","S_2_,_e_x","F_2_1","F_2_2"})
title({"2:1 Signal:Noise", "Classification Accuracy=93.75%"}, 'FontSize',10)
ylim([-8 10])
xlim=get(gca,'xlim');
hold on
plot(xlim,[0 0])

subplot(2,2,3)
boxplot(normalized_params{3,1});
set(gca,'TickLabelInterpreter', 'tex');
set(gca, 'Fontsize', 8, 'Fontweight', 'bold');
set(gca,'xticklabel',{'S_1_,_b',"S_1_,_e_x","F_1_1","F_1_2","S_2_,_b","S_2_,_e_x","F_2_1","F_2_2"})
title({"5:1 Signal:Noise", "Classification Accuracy=91.25%"}, 'FontSize',10)
ylim([-8 10])
xlim=get(gca,'xlim');
hold on
plot(xlim,[0 0])


subplot(2,2,4)
boxplot(normalized_params{4,1});
set(gca,'TickLabelInterpreter', 'tex');
set(gca, 'Fontsize', 8, 'Fontweight', 'bold');
set(gca,'xticklabel',{'S_1_,_b',"S_1_,_e_x","F_1_1","F_1_2","S_2_,_b","S_2_,_e_x","F_2_1","F_2_2"})
title({"2:1 Signal:Noise", "Classification Accuracy=87%"}, 'FontSize',10)
ylim([-8 10])
xlim=get(gca,'xlim');
hold on
plot(xlim,[0 0])

size = [6.75 8.75];
res = 300;
set(gcf,'paperunits','inches','paperposition',[0 0 size]);
print('S4.tiff','-dtiff',['-r' num2str(res)]);

%% best SSEs
count=50;
for j=1:4
    for i=1:count
        [M,I] = min(SSEs_2node{j,i});
SSEs_2node_best{j,1}(i,:)=SSEs_2node{j,i}(I,:);
        [M,I] = min(threenoderesults.SSEs_11tp{j,i});
SSEs_3node_best{j,1}(i,:)=threenoderesults.SSEs_11tp{j,i}(I,:);
    end
figure
edges = linspace(0, 25, 25); % Create 20 bins.
h1=histogram(SSEs_2node_best{j,1},'BinEdges',edges,'EdgeColor', 'blue', 'FaceColor',  'blue');
hold on
h2=histogram(SSEs_3node_best{j,1},'BinEdges',edges,'EdgeColor', 'red', 'FaceColor',  'red');
legend ('2 node SSE','3 node SSE')
end

%% count best SSEs
load("dropped3node.mat")
count=50;
for j=1:4
    s=sign(normalized_params{j,1});
    ipositif{j,1}=sum(s(:)==1)/4;
    
end
%% count best normalized
load ("param3wrapper_paper.mat")
count=50;
for j=1:4
    for i=1:count
        [M,I] = min(threenoderesults.SSEs_11tp{j,i});
SSEs_3node_norm{j,1}(1:10,i)=threenoderesults.SSEs_11tp{j,i}/threenoderesults.SSEs_11tp{j,i}(I,:);
    end
figure
end
%% count all SSEs
load ("param3wrapper_paper.mat")
count=50;
for j=1:4
    k=1;
    for i=1:count
        [M,I] = min(threenoderesults.SSEs_11tp{j,i});
        all_SSEs_diff{j,1}(k:k+10-1,1)= (threenoderesults.SSEs_11tp{j,i}-threenoderesults.SSEs_11tp{j,i}(I,:));
        k=k+10;
    end
    all_SSEs_diff_sorted=sort(all_SSEs_diff{j,1});
    
end
%%
load ("param3wrapper_positive.mat")
count=50;

for j=1:4
    k=1;
    for i=1:count
        [M,I] = min(threenoderesults.SSEs_11tp{j,i});
        all_SSEs_diff{j,1}(k:k+10-1,1)= (threenoderesults.SSEs_11tp{j,i}-threenoderesults.SSEs_11tp{j,i}(I,:));
        k=k+10;
    end
    all_SSEs_diff_sorted{j,1}=sort(all_SSEs_diff{j,1});
end
for j=1:4
        for i=1:count
            k=1;
            for m=1:10
%           if SSEs_3node_norm{j,1}(m,i)<2
                if threenoderesults.SSEs_11tp{j,i}(m,1)<threenoderesults.SSEs_11tp{j,i}(I,1)+all_SSEs_diff_sorted{j,1}(401,1)
                    best_params{j,i}(k,1:15)=threenoderesults.predicted_param_11tp{j,i}(m,:);
                    k=k+1;
                end
            end
        end
end
for j=1:4
    k=1;
    for i=1:count
          [M,I] = min(threenoderesults.SSEs_11tp{j,i});
          X=(threenoderesults.predicted_param_11tp{j,i}(I,:))';
          corre=corr(X,(best_params{j,i})');
%           corre=getCV((best_params{j,i})');
          l=length(corre);
          correl{j,1}(1,k:k+l-1)=corre;
          k=k+l;
    end
    
%     plot(correl{j,1},'.')
%     ylim([-1.2,1.2])
    hold on
end
edges = linspace(-1, 1, 21); % Create 10 bins.
subplot(2,2,1)
h1=histogram(correl{1,1},'BinEdges',edges,'EdgeColor', 'black', 'FaceColor',  'black');

subplot(2,2,2)
h1=histogram(correl{2,1},'BinEdges',edges,'EdgeColor', 'black', 'FaceColor',  'green');

subplot(2,2,3)
h1=histogram(correl{3,1},'BinEdges',edges,'EdgeColor', 'black', 'FaceColor',  'blue');

subplot(2,2,4)
h1=histogram(correl{4,1},'BinEdges',edges,'EdgeColor', 'black', 'FaceColor',  'red');

size = [6.75 8.75];
res = 300;
set(gcf,'paperunits','inches','paperposition',[0 0 size]);
print('S6.tiff','-dtiff',['-r' num2str(res)]);

%%
load ("param3wrapper_paper.mat")
count=50;

for j=1:4
    k=1;
    for i=1:count
        [M,I] = min(threenoderesults.SSEs_11tp{j,i});
        all_SSEs_diff{j,1}(k:k+10-1,1)= (threenoderesults.SSEs_11tp{j,i}-threenoderesults.SSEs_11tp{j,i}(I,:));
        k=k+10;
    end
    all_SSEs_diff_sorted{j,1}=sort(all_SSEs_diff{j,1});
end
for j=1:4
        for i=1:count
            [M,I] = min(threenoderesults.SSEs_11tp{j,i});
            k=1;
            for m=1:10
%           if SSEs_3node_norm{j,1}(m,i)<2
                if threenoderesults.SSEs_11tp{j,i}(m,1)<2*threenoderesults.SSEs_11tp{j,i}(I,1)||threenoderesults.SSEs_11tp{j,i}(m,1)<0.0001
                    best_params{j,i}(k,1:15)=threenoderesults.predicted_param_11tp{j,i}(m,:);
                    k=k+1;
                end
            end
        end
end
for j=1:4
    k=1;
    for i=1:count
          [M,I] = min(threenoderesults.SSEs_11tp{j,i});
          X=(threenoderesults.predicted_param_11tp{j,i}(I,:))';
          cov{j,1}(i,:)=std(best_params{j,i})./(mean(best_params{j,i}));
    end
    cov_col{j,1}=reshape(cov{j,1},[],1);
    hold on
end
edges = linspace(-10, 10, 41); % Create 10 bins.
subplot(2,2,1)
h1=histogram(cov{1,1},'BinEdges',edges,'EdgeColor', 'black', 'FaceColor',  'black');
title('No Noise','Fontsize',10);
subplot(2,2,2)
h1=histogram(cov{2,1},'BinEdges',edges,'EdgeColor', 'black', 'FaceColor',  'green');
title('10:1 Signal:Noise','Fontsize',10);

subplot(2,2,3)
h1=histogram(cov{3,1},'BinEdges',edges,'EdgeColor', 'black', 'FaceColor',  'blue');
title('5:1 Signal:Noise','Fontsize',10);

subplot(2,2,4)
h1=histogram(cov{4,1},'BinEdges',edges,'EdgeColor', 'black', 'FaceColor',  'red');
title('2:1 Signal:Noise','Fontsize',10);

size = [6.75 8.75];
res = 300;
set(gcf,'paperunits','inches','paperposition',[0 0 size]);
print('S7.tiff','-dtiff',['-r' num2str(res)]);


%% Data for 50% perturbation
load("param3wrapper.mat");
count=50;
y0=[0;0;0];
pert1=0.5;
pert2=0.5;
pert3=0.5;
for i = 1:count
param = threenoderesults.actual_param{1,i};
clc;
display(i);  
display(param);  
actual_param{1,i}=param;
timeint=1;
tmax=100;
        data_nostim=eval_data_3node_polyval_nostim(timeint,tmax,param,y0,y0,y0,y0,pert1,pert2,pert3,0);
   
        y_0=[data_nostim.Y(tmax+1,1),data_nostim.Y(tmax+1,2),data_nostim.Y(tmax+1,3)];
        y_p1=[data_nostim.YP1(tmax+1,1),data_nostim.YP1(tmax+1,2),data_nostim.YP1(tmax+1,3)];
        y_p2=[data_nostim.YP2(tmax+1,1),data_nostim.YP2(tmax+1,2),data_nostim.YP2(tmax+1,3)];
        y_p3=[data_nostim.YP3(tmax+1,1),data_nostim.YP3(tmax+1,2),data_nostim.YP3(tmax+1,3)];
tmax=10;
timeint=5;
data0_3tp=eval_data_3node_polyval(timeint,tmax,param,y_0,y_p1,y_p2,y_p3,pert1,pert2,pert3,0);
data10_3tp=eval_data_3node_polyval(timeint,tmax,param,y_0,y_p1,y_p2,y_p3,pert1,pert2,pert3,0.10);
data20_3tp=eval_data_3node_polyval(timeint,tmax,param,y_0,y_p1,y_p2,y_p3,pert1,pert2,pert3,0.20);
data50_3tp=eval_data_3node_polyval(timeint,tmax,param,y_0,y_p1,y_p2,y_p3,pert1,pert2,pert3,0.50);

timeint=5/3;
data0_7tp=eval_data_3node_polyval(timeint,tmax,param,y_0,y_p1,y_p2,y_p3,pert1,pert2,pert3,0);
data10_7tp=eval_data_3node_polyval(timeint,tmax,param,y_0,y_p1,y_p2,y_p3,pert1,pert2,pert3,0.10);
data20_7tp=eval_data_3node_polyval(timeint,tmax,param,y_0,y_p1,y_p2,y_p3,pert1,pert2,pert3,0.20);
data50_7tp=eval_data_3node_polyval(timeint,tmax,param,y_0,y_p1,y_p2,y_p3,pert1,pert2,pert3,0.50);

timeint=1;
data0_11tp=eval_data_3node_polyval(timeint,tmax,param,y_0,y_p1,y_p2,y_p3,pert1,pert2,pert3,0);
data10_11tp=eval_data_3node_polyval(timeint,tmax,param,y_0,y_p1,y_p2,y_p3,pert1,pert2,pert3,0.10);
data20_11tp=eval_data_3node_polyval(timeint,tmax,param,y_0,y_p1,y_p2,y_p3,pert1,pert2,pert3,0.20);
data50_11tp=eval_data_3node_polyval(timeint,tmax,param,y_0,y_p1,y_p2,y_p3,pert1,pert2,pert3,0.50);

timeint=0.5;
data0_21tp=eval_data_3node_polyval(timeint,tmax,param,y_0,y_p1,y_p2,y_p3,pert1,pert2,pert3,0);
data10_21tp=eval_data_3node_polyval(timeint,tmax,param,y_0,y_p1,y_p2,y_p3,pert1,pert2,pert3,0.10);
data20_21tp=eval_data_3node_polyval(timeint,tmax,param,y_0,y_p1,y_p2,y_p3,pert1,pert2,pert3,0.20);
data50_21tp=eval_data_3node_polyval(timeint,tmax,param,y_0,y_p1,y_p2,y_p3,pert1,pert2,pert3,0.50);

datas_3tp{1,i}=data0_3tp;
datas_3tp{2,i}=data10_3tp;
datas_3tp{3,i}=data20_3tp;
datas_3tp{4,i}=data50_3tp;

datas_7tp{1,i}=data0_7tp;
datas_7tp{2,i}=data10_7tp;
datas_7tp{3,i}=data20_7tp;
datas_7tp{4,i}=data50_7tp;

datas_11tp{1,i}=data0_11tp;
datas_11tp{2,i}=data10_11tp;
datas_11tp{3,i}=data20_11tp;
datas_11tp{4,i}=data50_11tp;

datas_21tp{1,i}=data0_21tp;
datas_21tp{2,i}=data10_21tp;
datas_21tp{3,i}=data20_21tp;
datas_21tp{4,i}=data50_21tp;

end
threenoderesults_50pert.actual_param=actual_param;

threenoderesults_50pert.datas_3tp=datas_3tp;
threenoderesults_50pert.datas_7tp=datas_7tp;
threenoderesults_50pert.datas_11tp=datas_11tp;
threenoderesults_50pert.datas_21tp=datas_21tp;

save ('50pertdata.mat', 'threenoderesults_50pert')
```
