## Supplementary Code for "Network Inference from Perturbation Time Course Data": fig3func.m

```
function [minparams,sum0,sum1,sum2,sum5] = fig3func(actual_param,predicted_param,SSE,count)
len=length(SSE);
for i=1:count
    paramtheor(i,:)=actual_param{1,i};
    [M,I] = min(SSE{1,i});
    parampred0(i,:)=predicted_param{1,i}(I,:);
    [M,I] = min(SSE{2,i});
    parampred1(i,:)=predicted_param{2,i}(I,:);
    [M,I] = min(SSE{3,i});
    parampred2(i,:)=predicted_param{3,i}(I,:);
    [M,I] = min(SSE{4,i});
    parampred5(i,:)=predicted_param{4,i}(I,:);
end
    minparams.paramtheor=paramtheor;
    minparams.parampred0=parampred0;
    minparams.parampred1=parampred1;
    minparams.parampred2=parampred2;
    minparams.parampred5=parampred5;


%no noise
p=reshape(parampred0,length(parampred0(:)),1);
t=reshape(paramtheor,length(paramtheor(:)),1);
ta=find(t>0);
tar0=sum(p(ta)>0)/sum(t>0);
tr=find(t<0);
trr0=sum(p(tr)<0)/sum(t<0);
sum0=(tar0+trr0)/2;

%1:10 noise
p=reshape(parampred1,length(parampred1(:)),1);
t=reshape(paramtheor,length(paramtheor(:)),1);
ta=find(t>0);
tar1=sum(p(ta)>0)/sum(t>0);
tr=find(t<0);
trr1=sum(p(tr)<0)/sum(t<0);
sum1=(tar1+trr1)/2;

%1:5 noise
p=reshape(parampred2,length(parampred2(:)),1);
t=reshape(paramtheor,length(paramtheor(:)),1);
ta=find(t>0);
tar2=sum(p(ta)>0)/sum(t>0);
tr=find(t<0);
trr2=sum(p(tr)<0)/sum(t<0);
sum2=(tar2+trr2)/2;

%1:2 noise
p=reshape(parampred5,length(parampred5(:)),1);
t=reshape(paramtheor,length(paramtheor(:)),1);
ta=find(t>0);
tar5=sum(p(ta)>0)/sum(t>0);
tr=find(t<0);
trr5=sum(p(tr)<0)/sum(t<0);
sum5=(tar5+trr5)/2;
end
```
