## Supplementary Code for "Network Inference from Perturbation Time Course Data": Makefigures_2F.m

```
%% Generate Data for 50% perturbation
% skip to next section to use generated data
load("param3wrapper.mat");
count=50;
y0=[0;0;0];
pert1=0.5;
pert2=0.5;
pert3=0.5;
for i = 1:count
param = threenoderesults.actual_param{1,i};
clc;
display(i);  
display(param);  
actual_param{1,i}=param;
timeint=1;
tmax=100;
        data_nostim=eval_data_3node_polyval_nostim(timeint,tmax,param,y0,y0,y0,y0,pert1,pert2,pert3,0);
   
        y_0=[data_nostim.Y(tmax+1,1),data_nostim.Y(tmax+1,2),data_nostim.Y(tmax+1,3)];
        y_p1=[data_nostim.YP1(tmax+1,1),data_nostim.YP1(tmax+1,2),data_nostim.YP1(tmax+1,3)];
        y_p2=[data_nostim.YP2(tmax+1,1),data_nostim.YP2(tmax+1,2),data_nostim.YP2(tmax+1,3)];
        y_p3=[data_nostim.YP3(tmax+1,1),data_nostim.YP3(tmax+1,2),data_nostim.YP3(tmax+1,3)];
tmax=10;
timeint=5;
data0_3tp=eval_data_3node_polyval(timeint,tmax,param,y_0,y_p1,y_p2,y_p3,pert1,pert2,pert3,0);
data10_3tp=eval_data_3node_polyval(timeint,tmax,param,y_0,y_p1,y_p2,y_p3,pert1,pert2,pert3,0.10);
data20_3tp=eval_data_3node_polyval(timeint,tmax,param,y_0,y_p1,y_p2,y_p3,pert1,pert2,pert3,0.20);
data50_3tp=eval_data_3node_polyval(timeint,tmax,param,y_0,y_p1,y_p2,y_p3,pert1,pert2,pert3,0.50);

timeint=5/3;
data0_7tp=eval_data_3node_polyval(timeint,tmax,param,y_0,y_p1,y_p2,y_p3,pert1,pert2,pert3,0);
data10_7tp=eval_data_3node_polyval(timeint,tmax,param,y_0,y_p1,y_p2,y_p3,pert1,pert2,pert3,0.10);
data20_7tp=eval_data_3node_polyval(timeint,tmax,param,y_0,y_p1,y_p2,y_p3,pert1,pert2,pert3,0.20);
data50_7tp=eval_data_3node_polyval(timeint,tmax,param,y_0,y_p1,y_p2,y_p3,pert1,pert2,pert3,0.50);

timeint=1;
data0_11tp=eval_data_3node_polyval(timeint,tmax,param,y_0,y_p1,y_p2,y_p3,pert1,pert2,pert3,0);
data10_11tp=eval_data_3node_polyval(timeint,tmax,param,y_0,y_p1,y_p2,y_p3,pert1,pert2,pert3,0.10);
data20_11tp=eval_data_3node_polyval(timeint,tmax,param,y_0,y_p1,y_p2,y_p3,pert1,pert2,pert3,0.20);
data50_11tp=eval_data_3node_polyval(timeint,tmax,param,y_0,y_p1,y_p2,y_p3,pert1,pert2,pert3,0.50);

timeint=0.5;
data0_21tp=eval_data_3node_polyval(timeint,tmax,param,y_0,y_p1,y_p2,y_p3,pert1,pert2,pert3,0);
data10_21tp=eval_data_3node_polyval(timeint,tmax,param,y_0,y_p1,y_p2,y_p3,pert1,pert2,pert3,0.10);
data20_21tp=eval_data_3node_polyval(timeint,tmax,param,y_0,y_p1,y_p2,y_p3,pert1,pert2,pert3,0.20);
data50_21tp=eval_data_3node_polyval(timeint,tmax,param,y_0,y_p1,y_p2,y_p3,pert1,pert2,pert3,0.50);

datas_3tp{1,i}=data0_3tp;
datas_3tp{2,i}=data10_3tp;
datas_3tp{3,i}=data20_3tp;
datas_3tp{4,i}=data50_3tp;

datas_7tp{1,i}=data0_7tp;
datas_7tp{2,i}=data10_7tp;
datas_7tp{3,i}=data20_7tp;
datas_7tp{4,i}=data50_7tp;

datas_11tp{1,i}=data0_11tp;
datas_11tp{2,i}=data10_11tp;
datas_11tp{3,i}=data20_11tp;
datas_11tp{4,i}=data50_11tp;

datas_21tp{1,i}=data0_21tp;
datas_21tp{2,i}=data10_21tp;
datas_21tp{3,i}=data20_21tp;
datas_21tp{4,i}=data50_21tp;

end
threenoderesults_50pert.actual_param=actual_param;

threenoderesults_50pert.datas_3tp=datas_3tp;
threenoderesults_50pert.datas_7tp=datas_7tp;
threenoderesults_50pert.datas_11tp=datas_11tp;
threenoderesults_50pert.datas_21tp=datas_21tp;

save ('50pertdata.mat', 'threenoderesults_50pert')

%% Generate and plot 2F
load("param3wrapper_positive.mat");
load("50pertdata_positive.mat");
our3tpdmraout = cell(4,50);
for i = 1:4
for j = 1:50
tempstruct = threenoderesults.datas_3tp(i,j);
tempstruct2 = threenoderesults_50pert.datas_3tp(i,j);
[theorthreejacdata,obsthreejacdata,ssjac] = dmrathreeanalyzer(1,1,5,tempstruct{1},tempstruct2{1});
tempobs = [median(obsthreejacdata(1,1,:)),median(obsthreejacdata(1,2,:)),median(obsthreejacdata(1,3,:)),median(obsthreejacdata(2,1,:)),median(obsthreejacdata(2,2,:)),median(obsthreejacdata(2,3,:)),median(obsthreejacdata(3,1,:)),median(obsthreejacdata(3,2,:)),median(obsthreejacdata(3,3,:))];
our3tpdmraout(i,j) = {[tempobs;tempobs;tempobs;tempobs;tempobs;tempobs;tempobs;tempobs;tempobs;tempobs]
};
end
end
our7tpdmraout = cell(4,50);
for i = 1:4
for j = 1:50
tempstruct = threenoderesults.datas_7tp(i,j);
tempstruct2 = threenoderesults_50pert.datas_7tp(i,j);
[theorthreejacdata,obsthreejacdata,ssjac] = dmrathreeanalyzer(1,1,5/3,tempstruct{1},tempstruct2{1});
tempobs = [median(obsthreejacdata(1,1,:)),median(obsthreejacdata(1,2,:)),median(obsthreejacdata(1,3,:)),median(obsthreejacdata(2,1,:)),median(obsthreejacdata(2,2,:)),median(obsthreejacdata(2,3,:)),median(obsthreejacdata(3,1,:)),median(obsthreejacdata(3,2,:)),median(obsthreejacdata(3,3,:))];
our7tpdmraout(i,j) = {[tempobs;tempobs;tempobs;tempobs;tempobs;tempobs;tempobs;tempobs;tempobs;tempobs]
};
end
end
our11tpdmraout = cell(4,50);
for i = 1:4
for j = 1:50
tempstruct = threenoderesults.datas_11tp(i,j);
tempstruct2 = threenoderesults_50pert.datas_11tp(i,j);
[theorthreejacdata,obsthreejacdata,ssjac] = dmrathreeanalyzer(1,1,1,tempstruct{1},tempstruct2{1});
tempobs = [median(obsthreejacdata(1,1,:)),median(obsthreejacdata(1,2,:)),median(obsthreejacdata(1,3,:)),median(obsthreejacdata(2,1,:)),median(obsthreejacdata(2,2,:)),median(obsthreejacdata(2,3,:)),median(obsthreejacdata(3,1,:)),median(obsthreejacdata(3,2,:)),median(obsthreejacdata(3,3,:))];
our11tpdmraout(i,j) = {[tempobs;tempobs;tempobs;tempobs;tempobs;tempobs;tempobs;tempobs;tempobs;tempobs]
};
end
end
our21tpdmraout = cell(4,50);
for i = 1:4
for j = 1:50
tempstruct = threenoderesults.datas_21tp(i,j);
tempstruct2 = threenoderesults_50pert.datas_21tp(i,j);
[theorthreejacdata,obsthreejacdata,ssjac] = dmrathreeanalyzer(1,1,0.5,tempstruct{1},tempstruct2{1});
tempobs = [median(obsthreejacdata(1,1,:)),median(obsthreejacdata(1,2,:)),median(obsthreejacdata(1,3,:)),median(obsthreejacdata(2,1,:)),median(obsthreejacdata(2,2,:)),median(obsthreejacdata(2,3,:)),median(obsthreejacdata(3,1,:)),median(obsthreejacdata(3,2,:)),median(obsthreejacdata(3,3,:))];
our21tpdmraout(i,j) = {[tempobs;tempobs;tempobs;tempobs;tempobs;tempobs;tempobs;tempobs;tempobs;tempobs]
};
end
end

jacparam = cell(1,50);
for i=1:50
jacparam(1,i) = {threenoderesults.actual_param{1,i}([3,4,5,8,9,10,13,14,15])};
end

SSEcell = cell(4,50);
for i = 1:4
for j = 1:50
SSEcell(i,j) = {[0,0,0,0,0,0,0,0,0,0]};
end
end

count=50;
ax_x=[-2,2];
ax_y=[-4,4];
figure;
subplot(1,4,1)
hold on;
area([0,2],[4,4],'FaceColor','g');
alpha(0.2)
area([0,-2],[-4,-4],'FaceColor','g');
alpha(0.2)
area([0,2],[-4,-4],'FaceColor','r');
alpha(0.2)
area([0,-2],[4,4],'FaceColor','r');
alpha(0.2)
[min3,sum0_3,sum1_3,sum2_3,sum5_3]=fig3func(jacparam,our3tpdmraout,SSEcell,count);
for i=1:count
hold on;
plot(min3.paramtheor(i,:),min3.parampred5(i,:),"r.",'MarkerSize',20);
plot(min3.paramtheor(i,:),min3.parampred2(i,:),"b.",'MarkerSize',20);
plot(min3.paramtheor(i,:),min3.parampred1(i,:),"g.",'MarkerSize',20);
plot(min3.paramtheor(i,:),min3.parampred0(i,:),"k.",'MarkerSize',20);
xlim([-2 2]);
ylim([-4 4]);
end
plot([0,0],ax_y,'k-','LineWidth',1.5)
plot(ax_y,[0,0],'k-','LineWidth',1.5)
title("3 timepoints",'FontSize',12);
subplot(1,4,2)
hold on;
area([0,2],[4,4],'FaceColor','g');
alpha(0.2)
area([0,-2],[-4,-4],'FaceColor','g');
alpha(0.2)
area([0,2],[-4,-4],'FaceColor','r');
alpha(0.2)
area([0,-2],[4,4],'FaceColor','r');
alpha(0.2)
[min7,sum0_7,sum1_7,sum2_7,sum5_7]=fig3func(jacparam,our7tpdmraout,SSEcell,count);
for i=1:count
hold on;
plot(min7.paramtheor(i,:),min7.parampred5(i,:),"r.",'MarkerSize',20);
plot(min7.paramtheor(i,:),min7.parampred2(i,:),"b.",'MarkerSize',20);
plot(min7.paramtheor(i,:),min7.parampred1(i,:),"g.",'MarkerSize',20);
plot(min7.paramtheor(i,:),min7.parampred0(i,:),"k.",'MarkerSize',20);
xlim([-2 2]);
ylim([-4 4]);
end
plot([0,0],ax_y,'k-','LineWidth',1.5)
plot(ax_y,[0,0],'k-','LineWidth',1.5)
title("7 timepoints",'FontSize',12);
subplot(1,4,3)
hold on;
area([0,2],[4,4],'FaceColor','g');
alpha(0.2)
area([0,-2],[-4,-4],'FaceColor','g');
alpha(0.2)
area([0,2],[-4,-4],'FaceColor','r');
alpha(0.2)
area([0,-2],[4,4],'FaceColor','r');
alpha(0.2)
[min11,sum0_11,sum1_11,sum2_11,sum5_11]=fig3func(jacparam,our11tpdmraout,SSEcell,count);
for i=1:count
hold on;
plot(min11.paramtheor(i,:),min11.parampred5(i,:),"r.",'MarkerSize',20);
plot(min11.paramtheor(i,:),min11.parampred2(i,:),"b.",'MarkerSize',20);
plot(min11.paramtheor(i,:),min11.parampred1(i,:),"g.",'MarkerSize',20);
plot(min11.paramtheor(i,:),min11.parampred0(i,:),"k.",'MarkerSize',20);
xlim([-2 2]);
ylim([-4 4]);
end
plot([0,0],ax_y,'k-','LineWidth',1.5)
plot(ax_y,[0,0],'k-','LineWidth',1.5)
title("11 timepoints",'FontSize',12);
subplot(1,4,4)
hold on;
area([0,2],[4,4],'FaceColor','g');
alpha(0.2)
area([0,-2],[-4,-4],'FaceColor','g');
alpha(0.2)
area([0,2],[-4,-4],'FaceColor','r');
alpha(0.2)
area([0,-2],[4,4],'FaceColor','r');
alpha(0.2)
[min21,sum0_21,sum1_21,sum2_21,sum5_21]=fig3func(jacparam,our21tpdmraout,SSEcell,count);
for i=1:count
hold on;
plot(min21.paramtheor(i,:),min21.parampred5(i,:),"r.",'MarkerSize',20);
plot(min21.paramtheor(i,:),min21.parampred2(i,:),"b.",'MarkerSize',20);
plot(min21.paramtheor(i,:),min21.parampred1(i,:),"g.",'MarkerSize',20);
plot(min21.paramtheor(i,:),min21.parampred0(i,:),"k.",'MarkerSize',20);
xlim([-2 2]);
ylim([-4 4]);
end
plot([0,0],ax_y,'k-','LineWidth',1.5)
plot(ax_y,[0,0],'k-','LineWidth',1.5)
title("21 timepoints",'FontSize',12);
size = [8.5 2.125];
res = 300;
set(gcf,'paperunits','inches','paperposition',[0 0 size]);
print('2F.tiff','-dtiff',['-r' num2str(res)]);


% count=50;
% actual_param=jacparam;
% [min3,sum0_3,sum1_3,sum2_3,sum5_3]=fig3func(jacparam,our3tpdmraout,SSEcell,count);
% [min7,sum0_7,sum1_7,sum2_7,sum5_7]=fig3func(jacparam,our7tpdmraout,SSEcell,count);
% [min11,sum0_11,sum1_11,sum2_11,sum5_11]=fig3func(jacparam,our11tpdmraout,SSEcell,count);
% [min21,sum0_21,sum1_21,sum2_21,sum5_21]=fig3func(jacparam,our21tpdmraout,SSEcell,count);
% d=figure;
% in=roundn([sum0_3,sum1_3,sum2_3,sum5_3;sum0_7,sum1_7,sum2_7,sum5_7;sum0_11,sum1_11,sum2_11,sum5_11;sum0_21,sum1_21,sum2_21,sum5_21],-2);
% hm=heatmap(in');
% caxis([0.5 1])
% hm.Colormap=hot;
% hm.YData = ["No Noise" "10:1 S:N" "5:1 S:N" "2:1 S:N"];
% hm.XData = ["3 Timepoints" "7 Timepoints" "11 Timepoints" "21 Timepoints"];
% hm.Title="Classification Accuracy";
% hm.FontSize=11;
% res = 300;
% set(gcf,'paperunits','inches','PaperPosition',[0 0 3.5 3]);
% print('New2F.tiff','-dtiff',['-r' num2str(res)]);
```
