## Supplementary Code for "Network Inference from Perturbation Time Course Data": threeODE.m

```
function dy = threeODE(t,y)

dy1 = 1/(1 + 3*y(4)) - y(1);
dy2 = 1/(1 + 3*y(5)) - y(2);
dy3 = 1/(1 + 3*y(6)) - y(3);
dy4 = (-5)*y(4);
dy5 = (-5)*y(5);
dy6 = (-5)*y(6);

dy = [dy1; dy2; dy3; dy4; dy5; dy6];
```
