## Supplementary Code for "Network Inference from Perturbation Time Course Data": Celltransitionwrapper_parfor.m

```
function [cell_transitionresults] = Celltransitionwrapper_parfor(count,pert,multistart)

tic
for i = 1:count
    
    counter = 1;
    disp(i);
    good = 0;
    param = zeros(1,9);
    while(~good)
             
        param(2) = rand(1);
        param(3) = rand(1);
        param(4) = rand(1);
        param(6) = (1-param(3))*rand(1);
        param(7) = (1-param(4))*rand(1);
        param(8) = (1-param(2))*rand(1);
       
        param(1) = -param(4)-param(7);
        param(5) = -param(2)-param(8);
        param(9) = -param(3)-param(6);
     
        data=eval_data_celltransition(1,10,param,pert,0);
        topval = max([max(abs(data.Y)) max(abs(data.YP1)) max(abs(data.YP2)) max(abs(data.YP3))]);
        minval = min([min((data.Y)) min((data.YP1)) min((data.YP2)) min((data.YP3))]);
      
        if topval <= 1 && minval >=0
            good = 1;
        end           
        
        counter = counter+1;
        
    end 

display(param);  
actual_param{1,i}=param;
tmax=5;
timeint=5/2;
data0_3tp=eval_data_celltransition(timeint,tmax,param,pert,0);
data10_3tp=eval_data_celltransition(timeint,tmax,param,pert,0.10);
data20_3tp=eval_data_celltransition(timeint,tmax,param,pert,0.20);
data50_3tp=eval_data_celltransition(timeint,tmax,param,pert,0.50);

timeint=5/6;
data0_7tp=eval_data_celltransition(timeint,tmax,param,pert,0);
data10_7tp=eval_data_celltransition(timeint,tmax,param,pert,0.10);
data20_7tp=eval_data_celltransition(timeint,tmax,param,pert,0.20);
data50_7tp=eval_data_celltransition(timeint,tmax,param,pert,0.50);

timeint=1/2;
data0_11tp=eval_data_celltransition(timeint,tmax,param,pert,0);
data10_11tp=eval_data_celltransition(timeint,tmax,param,pert,0.10);
data20_11tp=eval_data_celltransition(timeint,tmax,param,pert,0.20);
data50_11tp=eval_data_celltransition(timeint,tmax,param,pert,0.50);

timeint=1/4;
data0_21tp=eval_data_celltransition(timeint,tmax,param,pert,0);
data10_21tp=eval_data_celltransition(timeint,tmax,param,pert,0.10);
data20_21tp=eval_data_celltransition(timeint,tmax,param,pert,0.20);
data50_21tp=eval_data_celltransition(timeint,tmax,param,pert,0.50);

datas_3tp{1,i}=data0_3tp;
datas_3tp{2,i}=data10_3tp;
datas_3tp{3,i}=data20_3tp;
datas_3tp{4,i}=data50_3tp;

datas_7tp{1,i}=data0_7tp;
datas_7tp{2,i}=data10_7tp;
datas_7tp{3,i}=data20_7tp;
datas_7tp{4,i}=data50_7tp;

datas_11tp{1,i}=data0_11tp;
datas_11tp{2,i}=data10_11tp;
datas_11tp{3,i}=data20_11tp;
datas_11tp{4,i}=data50_11tp;

datas_21tp{1,i}=data0_21tp;
datas_21tp{2,i}=data10_21tp;
datas_21tp{3,i}=data20_21tp;
datas_21tp{4,i}=data50_21tp;

end

for i=1:count
% 3 timepoints
for j=1:4
        for k=1:multistart
            [predicted_param_3tp{j,i}(k,:),SSEs_3tp{j,i}(k,1)]=estimateCT(datas_3tp{j,i});
 
% 7 timepoints
    
            [predicted_param_7tp{j,i}(k,:),SSEs_7tp{j,i}(k,1)]=estimateCT(datas_7tp{j,i});
     
% 11 timepoints

            [predicted_param_11tp{j,i}(k,:),SSEs_11tp{j,i}(k,1)]=estimateCT(datas_11tp{j,i});
            
% 21 timepoints
            [predicted_param_21tp{j,i}(k,:),SSEs_21tp{j,i}(k,1)]=estimateCT(datas_21tp{j,i});
        end
    end
    
end

cell_transitionresults.actual_param=actual_param;

cell_transitionresults.datas_3tp=datas_3tp;
cell_transitionresults.predicted_param_3tp=predicted_param_3tp;
cell_transitionresults.SSEs_3tp=SSEs_3tp;

cell_transitionresults.datas_7tp=datas_7tp;
cell_transitionresults.predicted_param_7tp=predicted_param_7tp;
cell_transitionresults.SSEs_7tp=SSEs_7tp;

cell_transitionresults.datas_11tp=datas_11tp;
cell_transitionresults.predicted_param_11tp=predicted_param_11tp;
cell_transitionresults.SSEs_11tp=SSEs_11tp;

cell_transitionresults.datas_21tp=datas_21tp;
cell_transitionresults.predicted_param_21tp=predicted_param_21tp;
cell_transitionresults.SSEs_21tp=SSEs_21tp;


    save ('celltransitionwrapper.mat', 'cell_transitionresults')
    toc
end
```
