## Supplementary Code for "Network Inference from Perturbation Time Course Data": estimateCT.m

```
function [fmin,finSSE]=estimateCT(data)
global Y YP1 YP2 YP3
tic
Y=data.Y;
YP1=data.YP1;
YP2=data.YP2;
YP3=data.YP3;
t=data.T;
timeint=t(2)-t(1);
% Decide the order of polynomial function to fit the data
poly_order=4;
% if length(t)<5
% poly_order=2;
% end

poly_YP1=polyfit(t,YP1(:,1),poly_order);
poly_YP2=polyfit(t,YP2(:,2),poly_order);
poly_YP3=polyfit(t,YP3(:,3),poly_order);
        conoptions = optimoptions('fmincon','MaxFunctionEvaluations',10000);
        % Set upper and lower bounds for edge weights

% lb = [-1 -1 -1 -1 -1 -1 -1 -1 -1];
% ub = [1 1 1 1 1 1 1 1 1];

lb = [0 0 0 0 0 0 0 0 0];
ub = [1 1 1 1 1 1 1 1 1];
% Obtain LSE solution through fmincon starting from random initial guess. 
% Try/catch is used-if there is an error, it will restart from a different
% random initial guess
good=0;
try
    while (~good)
        initial=[rand(1),rand(1),rand(1),rand(1),rand(1),rand(1),rand(1),rand(1),rand(1)];
        [t,estY] = ode15s(@odeqns,0:timeint:t(length(t)),[Y(1,1);Y(1,2);Y(1,3)],odeset('RelTol',1e-6),initial);
        [t,estYP1] = ode15s(@odep1,0:timeint:t(length(t)),[YP1(1,2);YP1(1,3)],odeset('RelTol',1e-6),initial);
        [t,estYP2] = ode15s(@odep2,0:timeint:t(length(t)),[YP2(1,1);YP2(1,3)],odeset('RelTol',1e-6),initial);
        [t,estYP3] = ode15s(@odep3,0:timeint:t(length(t)),[YP3(1,1);YP3(1,2)],odeset('RelTol',1e-6),initial);
        topval = max([max(abs(estY)) max(abs(estYP1)) max(abs(estYP2)) max(abs(estYP3))]);
%         minval = min([min((estY)) min((estYP1)) min((estYP2)) min((estYP3))]);
        if topval<100
        good=1;
        end
    end
    [fmin,finSSE]=fmincon(@nestedfun,initial,[],[],[],[],lb,ub,[],conoptions);
catch
    try
        good=0;
        while (~good)
        initial=[rand(1),rand(1),rand(1),rand(1),rand(1),rand(1),rand(1),rand(1),rand(1)];
            [t,estY] = ode15s(@odeqns,0:timeint:t(length(t)),[Y(1,1);Y(1,2);Y(1,3)],odeset('RelTol',1e-6),initial);
            [t,estYP1] = ode15s(@odep1,0:timeint:t(length(t)),[YP1(1,2);YP1(1,3)],odeset('RelTol',1e-6),initial);
            [t,estYP2] = ode15s(@odep2,0:timeint:t(length(t)),[YP2(1,1);YP2(1,3)],odeset('RelTol',1e-6),initial);
            [t,estYP3] = ode15s(@odep3,0:timeint:t(length(t)),[YP3(1,1);YP3(1,2)],odeset('RelTol',1e-6),initial);
            topval = max([max(abs(estY)) max(abs(estYP1)) max(abs(estYP2)) max(abs(estYP3))]);
            minval = min([min((estY)) min((estYP1)) min((estYP2)) min((estYP3))]);
            if topval<10 
                good=1;
            end
        end
        [fmin,finSSE]=fmincon(@nestedfun,initial,[],[],[],[],lb,ub,[],conoptions);
    catch
        try 
        fmin=ones(9);
        finSSE=10000;
        end
        
        end
    end

fmin(1) = -fmin(4)-fmin(7);
fmin(5) = -fmin(2)-fmin(8);
fmin(9) = -fmin(3)-fmin(6);

display(fmin);
toc
        
function SSE = nestedfun(r2)
    [t,guessY] = ode15s(@odeqns,0:timeint:t(length(t)),[Y(1,1);Y(1,2);Y(1,3)],odeset('RelTol',1e-6),r2);
    [t,guessYP1] = ode15s(@odep1,0:timeint:t(length(t)),[YP1(1,2);YP1(1,3)],odeset('RelTol',1e-6),r2);
    [t,guessYP2] = ode15s(@odep2,0:timeint:t(length(t)),[YP2(1,1);YP2(1,3)],odeset('RelTol',1e-6),r2);
    [t,guessYP3] = ode15s(@odep3,0:timeint:t(length(t)),[YP3(1,1);YP3(1,2)],odeset('RelTol',1e-6),r2);

    % normalize by all elements of actual data
%     SSE = (sum(sum(((Y(2:11,:)-guessY(2:11,:))./(Y(2:11,:))).^2)...
%         +sum(((YP1(2:11,2)-guessYP1(2:11,1))./(YP1(2:11,2))).^2)...
%         +sum(((YP1(2:11,3)-guessYP1(2:11,2))./(YP1(2:11,3))).^2)...
%         +sum(((YP2(2:11,1)-guessYP2(2:11,1))./(YP2(2:11,1))).^2)...
%         +sum(((YP2(2:11,3)-guessYP2(2:11,2))./(YP2(2:11,3))).^2)...
%         +sum(((YP3(2:11,1)-guessYP3(2:11,1))./(YP3(2:11,1))).^2)...
%         +sum(((YP3(2:11,2)-guessYP3(2:11,2))./(YP3(2:11,2))).^2))).^0.5;
% % % normalize by median elements of actual data
%     SSE = (sum(sum(((Y(2:11,:)-guessY(2:11,:))./median(Y,1)).^2)...
%         +sum(((YP1(2:11,2)-guessYP1(2:11,1))./median(YP1(:,2),1)).^2)...
%         +sum(((YP1(2:11,3)-guessYP1(2:11,2))./median(YP1(:,3),1)).^2)...
%         +sum(((YP2(2:11,1)-guessYP2(2:11,1))./median(YP2(:,1),1)).^2)...
%         +sum(((YP2(2:11,3)-guessYP2(2:11,2))./median(YP2(:,3),1)).^2)...
%         +sum(((YP3(2:11,1)-guessYP3(2:11,1))./median(YP3(:,1),1)).^2)...
%         +sum(((YP3(2:11,2)-guessYP3(2:11,2))./median(YP3(:,2),1)).^2))).^0.5;
% % no normalization 
    SSE = (sum(sum(((Y(1:length(t),:)-guessY(1:length(t),:))).^2)...
        +sum(((YP1(1:length(t),2)-guessYP1(1:length(t),1))).^2)...
        +sum(((YP1(1:length(t),3)-guessYP1(1:length(t),2))).^2)...
        +sum(((YP2(1:length(t),1)-guessYP2(1:length(t),1))).^2)...
        +sum(((YP2(1:length(t),3)-guessYP2(1:length(t),2))).^2)...
        +sum(((YP3(1:length(t),1)-guessYP3(1:length(t),1))).^2)...
        +sum(((YP3(1:length(t),2)-guessYP3(1:length(t),2))).^2))).^0.5;
end
    function dzdt1 = odeqns(t,z,r)
        dzdt1 = [
            ((-r(4)-r(7))*z(1)+r(2)*z(2)+r(3)*z(3))
            (r(4)*z(1)+(-r(2)-r(8))*z(2)+r(6)*z(3))
            (r(7)*z(1)+r(8)*z(2)+(-r(3)-r(6))*z(3))
        ];
    end
    function dzdt1 = odep1(t,z,r)
        Y_pert=polyval(poly_YP1,t);   
        dzdt1 = [
            (r(4)*Y_pert+(-r(2)-r(8))*z(1)+r(6)*z(2))
            (r(7)*Y_pert+r(8)*z(1)+(-r(3)-r(6))*z(2))
                 ];
    end
       function dzdt1 = odep2(t,z,r)
        Y_pert=polyval(poly_YP2,t);
        dzdt1 = [
            ((-r(4)-r(7))*z(1)+r(2)*Y_pert+r(3)*z(2))
            (r(7)*z(1)+r(8)*Y_pert+(-r(3)-r(6))*z(2))
        ];
       end
       function dzdt1 = odep3(t,z,r)
        Y_pert=polyval(poly_YP3,t);
        dzdt1 = [
            ((-r(4)-r(7))*z(1)+r(2)*z(2)+r(3)*Y_pert)
            (r(4)*z(1)+(-r(2)-r(8))*z(2)+r(6)*Y_pert)
        ];
       end
        
end
```
