## Supplementary Code for "Network Inference from Perturbation Time Course Data": estimateGuptadata.m

```
function guessGupta= estimateGuptadata(data_3_node,preds0)
global poly_YP1 poly_YP2 poly_YP3
Y=data_3_node.Y;
YP1=data_3_node.YP1;
YP2=data_3_node.YP2;
YP3=data_3_node.YP3;
t=data_3_node.T;
timeint=t(2)-t(1);
poly_order=8;
poly_YP1=polyfit(t,YP1(:,1),poly_order);
poly_YP2=polyfit(t,YP2(:,2),poly_order);
poly_YP3=polyfit(t,YP3(:,3),poly_order);

r2=preds0{1,1}(1:9);

[t,guessY] = ode15s(@odeqns,0:timeint:t(length(t)),[Y(1,1);Y(1,2);Y(1,3)],odeset('RelTol',1e-6),r2);
[t,guessYP1] = ode15s(@odep1,0:timeint:t(length(t)),[YP1(1,2);YP1(1,3)],odeset('RelTol',1e-6),r2);
[t,guessYP2] = ode15s(@odep2,0:timeint:t(length(t)),[YP2(1,1);YP2(1,3)],odeset('RelTol',1e-6),r2);
[t,guessYP3] = ode15s(@odep3,0:timeint:t(length(t)),[YP3(1,1);YP3(1,2)],odeset('RelTol',1e-6),r2);

guessGupta.Y=guessY; 
guessGupta.YP1=guessYP1; 
guessGupta.YP2=guessYP2; 
guessGupta.YP3=guessYP3; 
guessGupta.T=t; 
   
 function dzdt1 = odeqns(t,z,r)
        dzdt1 = [
            ((-r(4)-r(7))*z(1)+r(2)*z(2)+r(3)*z(3))
            (r(4)*z(1)+(-r(2)-r(8))*z(2)+r(6)*z(3))
            (r(7)*z(1)+r(8)*z(2)+(-r(3)-r(6))*z(3))
        ];
    end
    function dzdt1 = odep1(t,z,r)
%     global poly_YP1 poly_YP2 poly_YP3
        Y_pert=polyval(poly_YP1,t);   
        dzdt1 = [
            (r(4)*Y_pert+(-r(2)-r(8))*z(1)+r(6)*z(2))
            (r(7)*Y_pert+r(8)*z(1)+(-r(3)-r(6))*z(2))
                 ];
    end
       function dzdt1 = odep2(t,z,r)
%        global poly_YP1 poly_YP2 poly_YP3
        Y_pert=polyval(poly_YP2,t);
        dzdt1 = [
            ((-r(4)-r(7))*z(1)+r(2)*Y_pert+r(3)*z(2))
            (r(7)*z(1)+r(8)*Y_pert+(-r(3)-r(6))*z(2))
        ];
       end
       function dzdt1 = odep3(t,z,r)
%        global poly_YP1 poly_YP2 poly_YP3
        Y_pert=polyval(poly_YP3,t);
        dzdt1 = [
            ((-r(4)-r(7))*z(1)+r(2)*z(2)+r(3)*Y_pert)
            (r(4)*z(1)+(-r(2)-r(8))*z(2)+r(6)*Y_pert)
        ];
       end
        
end
```
