## Supplementary Code for "Network Inference from Perturbation Time Course Data": estimateMarkov.m

```
function [fmin,finSSE]=estimateMarkov(data)
global Y YP1 YP2 YP3
tic
Y=data.Y;
YP1=data.YP1;
YP2=data.YP2;
YP3=data.YP3;
t=data.T;
timeint=t(2)-t(1);
        conoptions = optimoptions('fmincon','MaxFunctionEvaluations',10000);
        % Set upper and lower bounds for edge weights

lb = [0 0 0 0 0 0 0 0 0];
ub = [1 1 1 1 1 1 1 1 1];
% Obtain LSE solution through fmincon starting from random initial guess. 
% Try/catch is used-if there is an error, it will restart from a different
% random initial guess
good=0;
steps=length(Y);
% try
    initial=[2*rand(1)-1,2*rand(1)-1,2*rand(1)-1,2*rand(1)-1,2*rand(1)-1,2*rand(1)-1,2*rand(1)-1,2*rand(1)-1,2*rand(1)-1];
    [fmin,finSSE]=fmincon(@nestedfun,initial,[],[],[],[],lb,ub,[],conoptions);
% catch
% end
display(fmin);
toc
        
function SSE = nestedfun(prob)
    guessY(1,1:3)=Y(1,1:3);
    guessYP1(1,1:3)=YP1(1,1:3);
    guessYP2(1,1:3)=YP2(1,1:3);
    guessYP3(1,1:3)=YP3(1,1:3);

for i=2:steps  
    guessY(i,1)=guessY(i-1,1)*prob(1)+guessY(i-1,2)*prob(2)+guessY(i-1,3)*prob(3);
    guessY(i,2)=guessY(i-1,1)*prob(4)+guessY(i-1,2)*prob(5)+guessY(i-1,3)*prob(6);
    guessY(i,3)=guessY(i-1,1)*prob(7)+guessY(i-1,2)*prob(8)+guessY(i-1,3)*prob(9);
    
    guessYP1(i,1)=guessYP1(i-1,1)*prob(1)+guessYP1(i-1,2)*prob(2)+guessYP1(i-1,3)*prob(3);
    guessYP1(i,2)=guessYP1(i-1,1)*prob(4)+guessYP1(i-1,2)*prob(5)+guessYP1(i-1,3)*prob(6);
    guessYP1(i,3)=guessYP1(i-1,1)*prob(7)+guessYP1(i-1,2)*prob(8)+guessYP1(i-1,3)*prob(9);
    
    guessYP2(i,1)=guessYP2(i-1,1)*prob(1)+guessYP2(i-1,2)*prob(2)+guessYP2(i-1,3)*prob(3);
    guessYP2(i,2)=guessYP2(i-1,1)*prob(4)+guessYP2(i-1,2)*prob(5)+guessYP2(i-1,3)*prob(6);
    guessYP2(i,3)=guessYP2(i-1,1)*prob(7)+guessYP2(i-1,2)*prob(8)+guessYP2(i-1,3)*prob(9);
    
    guessYP3(i,1)=guessYP3(i-1,1)*prob(1)+guessYP3(i-1,2)*prob(2)+guessYP3(i-1,3)*prob(3);
    guessYP3(i,2)=guessYP3(i-1,1)*prob(4)+guessYP3(i-1,2)*prob(5)+guessYP3(i-1,3)*prob(6);
    guessYP3(i,3)=guessYP3(i-1,1)*prob(7)+guessYP3(i-1,2)*prob(8)+guessYP3(i-1,3)*prob(9);
end
% % no normalization 
    SSE = (sum(sum(((Y(1:11,:)-guessY)).^2)...
        +sum(((YP1-guessYP1)).^2)...
        +sum(((YP2-guessYP2)).^2)...
        +sum(((YP3-guessYP3)).^2))).^0.5;

end
 
end
```
