## Supplementary Code for "Network Inference from Perturbation Time Course Data": eval_data_celltransition.m

```
function data= eval_data_celltransition(timeint,tmax,params,pert,noise)

% Generate data without perturbation
[t,Y] = ode15s(@odeqns,0:timeint:tmax,pert(1,:),odeset('RelTol',1e-6),params);
[t,YP1] = ode15s(@odeqns,0:timeint:tmax,pert(2,:),odeset('RelTol',1e-6),params);
[t,YP2] = ode15s(@odeqns,0:timeint:tmax,pert(3,:),odeset('RelTol',1e-6),params);
[t,YP3] = ode15s(@odeqns,0:timeint:tmax,pert(4,:),odeset('RelTol',1e-6),params);

% Add random noise from normal distribution
D=zeros(1,3);
for i = 1:length(t)
    for j = 1:3
        for k = 1:3
        randomdata = normrnd(0,noise*(abs(Y(i,j))),1,1);
        D(1,k) = Y(i,j) + randomdata;
        end
        Y(i,j)=median(D(1,:));
    end
end

for i = 1:length(t)
    for j = 1:3
        for k = 1:3
        randomdata = normrnd(0,noise*(abs(YP1(i,j))),1,1);
        D(1,k) = YP1(i,j) + randomdata;
        end
        YP1(i,j)=median(D(1,:));
    end
end
        

for i = 1:length(t)
    for j = 1:3
        for k = 1:3
        randomdata = normrnd(0,noise*(abs(YP2(i,j))),1,1);
        D(1,k) = YP2(i,j) + randomdata;
        end
        YP2(i,j)=median(D(1,:));
    end
end


for i = 1:length(t)
    for j = 1:3
        for k = 1:3
        randomdata = normrnd(0,noise*(abs(YP3(i,j))),1,1);
        D(1,k) = YP3(i,j) + randomdata;
        end
        YP3(i,j)=median(D(1,:));
    end
end

data.Y=Y;
data.YP1=YP1;
data.YP2=YP2;
data.YP3=YP3;
data.T=t;
    
function dzdt1 = odeqns(t,z,r)
        dzdt1 = [
            (r(1)*z(1)+r(2)*z(2)+r(3)*z(3))
            (r(4)*z(1)+r(5)*z(2)+r(6)*z(3))
            (r(7)*z(1)+r(8)*z(2)+r(9)*z(3))
        ];
    end
 
end
```
