## Supplementary Code for "Network Inference from Perturbation Time Course Data": Makefigures_3.m

```
%% Create Gupta Data for SUM159 cells
pert=[1/3,1/3,1/3;1,0,0;0,1,0;0,0,1];
markov_prob=[0.58,0.01,0.04,0.35,0.99,0.49,0.07,0,0.47];
steps=11;
f=1;
noise=0;
data_3_node=Markovtransition (pert,markov_prob,steps,f,noise);
save ("Gupta_data.mat", 'data_3_node');

%% Estimate Gupta Data through Discrete Markov
%Look at preds0_SSE values to see the accuracy of fit of estimated model
load("Gupta_data.mat");
multistart=1;
for n=1:multistart
    [preds0_markov{1,1}(n,:),preds0_SSE_markov{1,1}(n,1)]= estimateMarkov(data_3_node);
n
end
save("Gupta_data.mat","data_3_node","preds0_markov");

%% Estimate Gupta data through Continuous Differential Parameters
%Look at preds0_SSE_2 values to see the accuracy of fit of estimated model
load("Gupta_data.mat");
multistart=1;
for n=1:multistart
    [preds0{1,1}(n,:),preds0_SSE{1,1}(n,1)]= estimateCT_no_noise(data_3_node);
n
end
save("Gupta_data.mat","data_3_node","preds0_markov","preds0");

%% Generate fit data from estimates
load("Gupta_data.mat")
guessGupta=estimateGuptadata(data_3_node,preds0);
save ("guessGuptadata.mat", "guessGupta");
%% Plot 3B
load("Gupta_data.mat");
load('guessGuptadata.mat')

figure;
subplot(1,4,1);
hold on

plot(data_3_node.T,data_3_node.Y(:,1),'r.', 'MarkerSize', 15);
plot(data_3_node.T,data_3_node.Y(:,2),'b.', 'MarkerSize', 15);
plot(data_3_node.T,data_3_node.Y(:,3),'y.', 'MarkerSize', 15);
plot(guessGupta.T,guessGupta.Y(:,1),'r--', 'MarkerSize', 15);
plot(guessGupta.T,guessGupta.Y(:,2),'b--', 'MarkerSize', 15);
plot(guessGupta.T,guessGupta.Y(:,3),'y--', 'MarkerSize', 15);

ylim([0 1])

subplot(1,4,2);
hold on

plot(data_3_node.T,data_3_node.YP1(:,1),'r.', 'MarkerSize', 15);
plot(data_3_node.T,data_3_node.YP1(:,2),'b.', 'MarkerSize', 15);
plot(data_3_node.T,data_3_node.YP1(:,3),'y.', 'MarkerSize', 15);
plot(data_3_node.T,data_3_node.YP1(:,1),'r--', 'MarkerSize', 15);
plot(guessGupta.T,guessGupta.YP1(:,1),'b--', 'MarkerSize', 15);
plot(guessGupta.T,guessGupta.YP1(:,2),'y--', 'MarkerSize', 15);
ylim([0 1])

subplot(1,4,3);
hold on

plot(data_3_node.T,data_3_node.YP2(:,1),'r.', 'MarkerSize', 15);
plot(data_3_node.T,data_3_node.YP2(:,2),'b.', 'MarkerSize', 15);
plot(data_3_node.T,data_3_node.YP2(:,3),'y.', 'MarkerSize', 15);
plot(guessGupta.T,guessGupta.YP2(:,1),'r--', 'MarkerSize', 15);
plot(data_3_node.T,data_3_node.YP2(:,2),'b--', 'MarkerSize', 15);
plot(guessGupta.T,guessGupta.YP2(:,2),'y--', 'MarkerSize', 15);
ylim([0 1])

subplot(1,4,4);
hold on

plot(data_3_node.T,data_3_node.YP3(:,1),'r.', 'MarkerSize', 15);
plot(data_3_node.T,data_3_node.YP3(:,2),'b.', 'MarkerSize', 15);
plot(data_3_node.T,data_3_node.YP3(:,3),'y.', 'MarkerSize', 15);
plot(guessGupta.T,guessGupta.YP3(:,1),'r--', 'MarkerSize', 15);
plot(guessGupta.T,guessGupta.YP3(:,2),'b--', 'MarkerSize', 15);
plot(data_3_node.T,data_3_node.YP3(:,3),'y--', 'MarkerSize', 15);
ylim([0 1])

res = 300;
set(gcf,'paperunits','inches','PaperPosition',[0 0 6 2]);

print('3B.tiff','-dtiff',['-r' num2str(res)]);
 %% Plot 3C Markov vs DLMRA prediction
clear;
outmap=load("Gupta_data.mat");
for i=1:9
    if abs(outmap.preds0{1,1}(1,i))<=0.001
        outmap.preds0{1,1}(1,i)=0;
    end
    
        if abs(outmap.preds0_markov{1,1}(1,i))<=0.001
        outmap.preds0_markov{1,1}(1,i)=0;
        end
end
    for i=1:3
        outmap.preds0_markov{1,1}(1,(i-1)*3+i)=outmap.preds0_markov{1,1}(1,(i-1)*3+1)-1;
    end
gt=reshape(outmap.preds0_markov{1,1}./abs(outmap.preds0_markov{1,1}),[3,3])';
p1=reshape(outmap.preds0{1,1}./abs(outmap.preds0{1,1}),[3,3])';
gt(isnan(gt))=0;
p1(isnan(p1))=0;
    
b=figure;
subplot(1,2,1);
h=heatmap(gt,'CellLabelColor','none'); colorbar off; %h.XDisplayLabels={'0','1','2','3'}; h.Colormap=redgreencmap;
h.Colormap=redgreencmap;
Ax = gca;
Ax.XDisplayLabels = nan(size(Ax.XDisplayData));
Ax.YDisplayLabels = nan(size(Ax.YDisplayData));

% title('Ground Truth')
subplot(1,2,2);
h=heatmap(p1,'CellLabelColor','none'); colorbar off; %h.XDisplayLabels={'0','1','2','3'}; h.Colormap=redgreencmap;
h.Colormap=redgreencmap;
Ax = gca;
Ax.XDisplayLabels = nan(size(Ax.XDisplayData));
Ax.YDisplayLabels = nan(size(Ax.YDisplayData));

res = 300;
set(gcf,'paperunits','inches','PaperPosition',[0 0 5.0 1.5]);
print('3C.tiff','-dtiff',['-r' num2str(res)]);
                
%% Generate data and estimates for 50 random cell transition models
% Skip to the next section to plot existing data
count=50;
multistart=10;
pert=[1/3,1/3,1/3;0,0.5,0.5;0.5,0,0.5;0.5,0.5,0];
cell_transitionresults = Celltransitionwrapper_parfor(count,pert,multistart);
%% plot Fig 3D
clear all;
load('celltransitionwrapper.mat')
count=50;
ax_x=[-1,1];
ax_y=[-2,2];
figure;
subplot(1,4,1)
hold on;
area([0,1],[2,2],'FaceColor','g');
alpha(0.2)
area([0,-1],[-2,-2],'FaceColor','g');
alpha(0.2)
area([0,1],[-2,-2],'FaceColor','r');
alpha(0.2)
area([0,-1],[2,2],'FaceColor','r');
alpha(0.2)

[min3,sum0_3,sum1_3,sum2_3,sum5_3]=fig3func(cell_transitionresults.actual_param,cell_transitionresults.predicted_param_3tp,cell_transitionresults.SSEs_3tp,count);
for i=1:count
hold on;
plot(min3.paramtheor(i,:),min3.parampred5(i,:),"r.",'MarkerSize',20);
plot(min3.paramtheor(i,:),min3.parampred2(i,:),"b.",'MarkerSize',20);
plot(min3.paramtheor(i,:),min3.parampred1(i,:),"g.",'MarkerSize',20);
plot(min3.paramtheor(i,:),min3.parampred0(i,:),"k.",'MarkerSize',20);
xlim([-1 1]);
ylim([-2 2]);
xlim([0 1]);
ylim([0 1]);
end
  plot([0,0],ax_y,'k-','LineWidth',1.5)
  plot(ax_y,[0,0],'k-','LineWidth',1.5)
title("3 timepoints",'FontSize',12);

subplot(1,4,2)
hold on;
area([0,1],[2,2],'FaceColor','g');
alpha(0.2)
area([0,-1],[-2,-2],'FaceColor','g');
alpha(0.2)
area([0,1],[-2,-2],'FaceColor','r');
alpha(0.2)
area([0,-1],[2,2],'FaceColor','r');
alpha(0.2)

[min7,sum0_7,sum1_7,sum2_7,sum5_7]=fig3func(cell_transitionresults.actual_param,cell_transitionresults.predicted_param_7tp,cell_transitionresults.SSEs_7tp,count);
for i=1:count
hold on;
plot(min7.paramtheor(i,:),min7.parampred5(i,:),"r.",'MarkerSize',20);
plot(min7.paramtheor(i,:),min7.parampred2(i,:),"b.",'MarkerSize',20);
plot(min7.paramtheor(i,:),min7.parampred1(i,:),"g.",'MarkerSize',20);
plot(min7.paramtheor(i,:),min7.parampred0(i,:),"k.",'MarkerSize',20);
xlim([-1 1]);
ylim([-2 2]);
xlim([0 1]);
ylim([0 1]);
end
  plot([0,0],ax_y,'k-','LineWidth',1.5)
  plot(ax_y,[0,0],'k-','LineWidth',1.5)
title("7 timepoints",'FontSize',12);

subplot(1,4,3)
hold on;
area([0,1],[2,2],'FaceColor','g');
alpha(0.2)
area([0,-1],[-2,-2],'FaceColor','g');
alpha(0.2)
area([0,1],[-2,-2],'FaceColor','r');
alpha(0.2)
area([0,-1],[2,2],'FaceColor','r');
alpha(0.2)

[min11,sum0_11,sum1_11,sum2_11,sum5_11]=fig3func(cell_transitionresults.actual_param,cell_transitionresults.predicted_param_11tp,cell_transitionresults.SSEs_11tp,count);
for i=1:count
hold on;
plot(min11.paramtheor(i,:),min11.parampred5(i,:),"r.",'MarkerSize',20);
plot(min11.paramtheor(i,:),min11.parampred2(i,:),"b.",'MarkerSize',20);
plot(min11.paramtheor(i,:),min11.parampred1(i,:),"g.",'MarkerSize',20);
plot(min11.paramtheor(i,:),min11.parampred0(i,:),"k.",'MarkerSize',20);
xlim([-1 1]);
ylim([-2 2]);
xlim([0 1]);
ylim([0 1]);
end
  plot([0,0],ax_y,'k-','LineWidth',1.5)
  plot(ax_y,[0,0],'k-','LineWidth',1.5)
title("11 timepoints",'FontSize',12);

subplot(1,4,4)
hold on;
area([0,1],[2,2],'FaceColor','g');
alpha(0.2)
area([0,-1],[-2,-2],'FaceColor','g');
alpha(0.2)
area([0,1],[-2,-2],'FaceColor','r');
alpha(0.2)
area([0,-1],[2,2],'FaceColor','r');
alpha(0.2)

[min21,sum0_21,sum1_21,sum2_21,sum5_21]=fig3func(cell_transitionresults.actual_param,cell_transitionresults.predicted_param_21tp,cell_transitionresults.SSEs_21tp,count);
for i=1:count
hold on;
plot(min21.paramtheor(i,:),min21.parampred5(i,:),"r.",'MarkerSize',20);
plot(min21.paramtheor(i,:),min21.parampred2(i,:),"b.",'MarkerSize',20);
plot(min21.paramtheor(i,:),min21.parampred1(i,:),"g.",'MarkerSize',20);
plot(min21.paramtheor(i,:),min21.parampred0(i,:),"k.",'MarkerSize',20);
xlim([-1 1]);
ylim([-2 2]);
xlim([0 1]);
ylim([0 1]);
end
  plot([0,0],ax_y,'k-','LineWidth',1.5)
  plot(ax_y,[0,0],'k-','LineWidth',1.5)
title("21 timepoints",'FontSize',12);

size = [8.5 2.125];
res = 300;
set(gcf,'paperunits','inches','paperposition',[0 0 size]);
print('3D.tiff','-dtiff',['-r' num2str(res)]);
%% Classification Accuracy
% Classification Accuracy is always 1 for cell transition model
% clear all;
% load('celltransitionwrapper.mat')
% %Uncomment the next line and run if you generated the full data. Otherwise
% %just run the section to generate paper figures
% %load('param3wrapper.mat')
% count=50;
% actual_param=cell_transitionresults.actual_param;
% [min3,sum0_3,sum1_3,sum2_3,sum5_3]=fig3func(cell_transitionresults.actual_param,cell_transitionresults.predicted_param_3tp,cell_transitionresults.SSEs_3tp,count);
% [min7,sum0_7,sum1_7,sum2_7,sum5_7]=fig3func(cell_transitionresults.actual_param,cell_transitionresults.predicted_param_7tp,cell_transitionresults.SSEs_7tp,count);
% [min11,sum0_11,sum1_11,sum2_11,sum5_11]=fig3func(cell_transitionresults.actual_param,cell_transitionresults.predicted_param_11tp,cell_transitionresults.SSEs_11tp,count);
% [min21,sum0_21,sum1_21,sum2_21,sum5_21]=fig3func(cell_transitionresults.actual_param,cell_transitionresults.predicted_param_21tp,cell_transitionresults.SSEs_21tp,count);
% 
% d=figure; 
% in=roundn([sum0_3,sum1_3,sum2_3,sum5_3;sum0_7,sum1_7,sum2_7,sum5_7;sum0_11,sum1_11,sum2_11,sum5_11;sum0_21,sum1_21,sum2_21,sum5_21],-2);
% hm=heatmap(in');
% caxis([0.5 1])
% hm.Colormap=hot;
% hm.YData = ["No Noise" "10:1 S:N" "5:1 S:N" "2:1 S:N"];
% hm.XData = ["3 Timepoints" "7 Timepoints" "11 Timepoints" "21 Timepoints"];
% hm.Title="Classification Accuracy (3 Node)";
% hm.FontSize=11;
% res = 300;
% set(gcf,'paperunits','inches','PaperPosition',[0 0 3.5 3]);
% print('2F.tiff','-dtiff',['-r' num2str(res)]);
```
