## Supplementary Code for "Network Inference from Perturbation Time Course Data": Markovtransition.m

```
function data=Markovtransition (pert,prob,steps,f,noise)


Y(1,1:3)=pert(1,1:3);
YP1(1,1:3)=pert(2,1:3);
YP2(1,1:3)=pert(3,1:3);
YP3(1,1:3)=pert(4,1:3);
T(1)=0;

for i=2:steps  
    Y(i,1)=f*(Y(i-1,1)*prob(1)+Y(i-1,2)*prob(2)+Y(i-1,3)*prob(3));
    Y(i,2)=f*(Y(i-1,1)*prob(4)+Y(i-1,2)*prob(5)+Y(i-1,3)*prob(6));
    Y(i,3)=f*(Y(i-1,1)*prob(7)+Y(i-1,2)*prob(8)+Y(i-1,3)*prob(9));
    
    YP1(i,1)=f*(YP1(i-1,1)*prob(1)+YP1(i-1,2)*prob(2)+YP1(i-1,3)*prob(3));
    YP1(i,2)=f*(YP1(i-1,1)*prob(4)+YP1(i-1,2)*prob(5)+YP1(i-1,3)*prob(6));
    YP1(i,3)=f*(YP1(i-1,1)*prob(7)+YP1(i-1,2)*prob(8)+YP1(i-1,3)*prob(9));
    
    YP2(i,1)=f*(YP2(i-1,1)*prob(1)+YP2(i-1,2)*prob(2)+YP2(i-1,3)*prob(3));
    YP2(i,2)=f*(YP2(i-1,1)*prob(4)+YP2(i-1,2)*prob(5)+YP2(i-1,3)*prob(6));
    YP2(i,3)=f*(YP2(i-1,1)*prob(7)+YP2(i-1,2)*prob(8)+YP2(i-1,3)*prob(9));
    
    YP3(i,1)=f*(YP3(i-1,1)*prob(1)+YP3(i-1,2)*prob(2)+YP3(i-1,3)*prob(3));
    YP3(i,2)=f*(YP3(i-1,1)*prob(4)+YP3(i-1,2)*prob(5)+YP3(i-1,3)*prob(6));
    YP3(i,3)=f*(YP3(i-1,1)*prob(7)+YP3(i-1,2)*prob(8)+YP3(i-1,3)*prob(9));
    
    T(i)=i-1;
end

% Add random noise from normal distribution
D=zeros(1,3);
for i = 1:length(T)
    for j = 1:3
        for k = 1:3
        randomdata = normrnd(0,noise*(abs(Y(i,j))),1,1);
        D(1,k) = Y(i,j) + randomdata;
        end
        Y(i,j)=median(D(1,:));
    end
end

for i = 1:length(T)
    for j = 1:3
        for k = 1:3
        randomdata = normrnd(0,noise*(abs(YP1(i,j))),1,1);
        D(1,k) = YP1(i,j) + randomdata;
        end
        YP1(i,j)=median(D(1,:));
    end
end
        

for i = 1:length(T)
    for j = 1:3
        for k = 1:3
        randomdata = normrnd(0,noise*(abs(YP2(i,j))),1,1);
        D(1,k) = YP2(i,j) + randomdata;
        end
        YP2(i,j)=median(D(1,:));
    end
end


for i = 1:length(T)
    for j = 1:3
        for k = 1:3
        randomdata = normrnd(0,noise*(abs(YP3(i,j))),1,1);
        D(1,k) = YP3(i,j) + randomdata;
        end
        YP3(i,j)=median(D(1,:));
    end
end

data.Y=Y;
data.YP1=YP1;
data.YP2=YP2;
data.YP3=YP3;
data.T=T;
```
