## Supplementary Code for "Network Inference from Perturbation Time Course Data": estimate2node.m

```
function [fmin,finSSE]=estimate2node (data)
tic
global Y YP1 YP2
tic
Y=data.Y;
YP1=data.YP1;
YP2=data.YP2;
t=data.T;
timeint=t(2)-t(1);

% Decide the order of polynomial function to fit the data
poly_order=length(t)-1;

poly_YP1=polyfit(t,YP1(:,1),poly_order);
poly_YP2=polyfit(t,YP2(:,2),poly_order);
         conoptions = optimoptions('fmincon','MaxFunctionEvaluations',10000);
% Set upper and lower bounds for edge weights
%          lb = [-10 -10 -10 -10 -10 -10 -10 -10];
%          ub = [10 10 10 10 10 10 10 10];
         lb = [-1 -1 -1 -1 -1 -1];
         ub = [1 1 1 1 1 1];
%          lb = [-10 -10 -10 0 -10 -10];
%          ub = [10 10 10 0 10 10];

% Obtain LSE solution through fmincon starting from random initial guess. 
% Try/catch is used-if there is an error, it will restart from a different
% random initial guess
try
        good=0;
        while (~good)
        initial=[rand(1),2*rand(1)-1,2*rand(1)-1,rand(1),2*rand(1)-1,2*rand(1)-1];
        [t,estY] = ode15s(@odeqns,0:timeint:t(length(t)),[Y(1,1);Y(1,2)],odeset('RelTol',1e-6),initial);
        [t,estYP1] = ode15s(@odep1,0:timeint:t(length(t)),[YP1(1,2)],odeset('RelTol',1e-6),initial);
        [t,estYP2] = ode15s(@odep2,0:timeint:t(length(t)),[YP2(1,1)],odeset('RelTol',1e-6),initial);
        topval = max([max(abs(estY)) max(abs(estYP1)) max(abs(estYP2))]);
        minval = min([min((estY)) min((estYP1)) min((estYP2))]);
        if topval<10 && minval>=0
        good=1;
        end
        end
            [fmin,finSSE]=fmincon(@nestedfun,initial,[],[],[],[],lb,ub,[],conoptions);
catch
    try
        good=0;
        while (~good)
        initial=[rand(1),2*rand(1)-1,2*rand(1)-1,rand(1),2*rand(1)-1,2*rand(1)-1];
        [t,estY] = ode15s(@odeqns,0:timeint:t(length(t)),[Y(1,1);Y(1,2)],odeset('RelTol',1e-6),initial);
        [t,estYP1] = ode15s(@odep1,0:timeint:t(length(t)),[YP1(1,2)],odeset('RelTol',1e-6),initial);
        [t,estYP2] = ode15s(@odep2,0:timeint:t(length(t)),[YP2(1,1)],odeset('RelTol',1e-6),initial);
        topval = max([max(abs(estY)) max(abs(estYP1)) max(abs(estYP2))]);
        minval = min([min((estY)) min((estYP1)) min((estYP2))]);
        if topval<10 && minval>=0
        good=1;
        end
        end
            [fmin,finSSE]=fmincon(@nestedfun,initial,[],[],[],[],lb,ub,[],conoptions);
    catch
    try
        good=0;
        while (~good)
        initial=[rand(1),2*rand(1)-1,2*rand(1)-1,rand(1),2*rand(1)-1,2*rand(1)-1];
        [t,estY] = ode15s(@odeqns,0:timeint:t(length(t)),[Y(1,1);Y(1,2)],odeset('RelTol',1e-6),initial);
        [t,estYP1] = ode15s(@odep1,0:timeint:t(length(t)),[YP1(1,2)],odeset('RelTol',1e-6),initial);
        [t,estYP2] = ode15s(@odep2,0:timeint:t(length(t)),[YP2(1,1)],odeset('RelTol',1e-6),initial);
        topval = max([max(abs(estY)) max(abs(estYP1)) max(abs(estYP2))]);
        minval = min([min((estY)) min((estYP1)) min((estYP2))]);
        if topval<10 && minval>=0
        good=1;
        end
        end
            [fmin,finSSE]=fmincon(@nestedfun,initial,[],[],[],[],lb,ub,[],conoptions);
    catch
    end
    end
end
        display(fmin);
        toc
        
function SSE = nestedfun(r2)
         
[t,guessY] = ode15s(@odeqns,0:timeint:t(length(t)),[Y(1,1);Y(1,2)],odeset('RelTol',1e-6),r2);
[t,guessYP1] = ode15s(@odep1,0:timeint:t(length(t)),[YP1(1,2)],odeset('RelTol',1e-6),r2);
[t,guessYP2] = ode15s(@odep2,0:timeint:t(length(t)),[YP2(1,1)],odeset('RelTol',1e-6),r2);

     SSE = (sum(sum((Y-guessY).^2)...
         +sum((YP1(:,2)-guessYP1).^2)...
         +sum((YP2(:,1)-guessYP2).^2))).^0.5;
    %WHOLE NORMALIZED
% SSE = (sum(sum(((Y(2:10,:)-guessY(2:10,:))./(Y(2:10,:))).^2)...
%         +sum(((YP1(2:10,2)-guessYP1(2:10,1))./(YP1(2:10,2))).^2)...
%         +sum(((YP2(2:10,1)-guessYP2(2:10,1))./(YP2(2:10,1))).^2))).^0.5;
% MEDIAN NORMALIZED
%     SSE = (sum(sum(((Y(2:10,1)-guessY(2:10,1))./median(Y(:,1),1)).^2)...
%         +sum(((Y(2:10,2)-guessY(2:10,1))./median(Y(:,2),1)).^2)...
%         +sum(((YP1(2:10,2)-guessYP1(2:10,1))./median(YP1(:,2),1)).^2)...
%         +sum(((YP2(2:10,1)-guessYP2(2:10,1))./median(YP2(:,1),1)).^2))).^0.5;


end   
    function dzdt1 = odeqns(t,z,r)
        dzdt1 = [
            (r(1)+r(2)*z(1)+r(3)*z(2))
            (r(4)+r(5)*z(1)+r(6)*z(2))
        ];
    end
    function dzdt1 = odep1(t,z,r2)
         Y_pert=polyval(poly_YP1,t);

        dzdt1 = [
            (r2(4)+r2(5)*Y_pert+r2(6)*z)
                 ];
    end
       function dzdt1 = odep2(t,z,r2)
        Y_pert=polyval(poly_YP2,t);

        dzdt1 = [
            (r2(1)+r2(2)*z+r2(3)*Y_pert)
        ];
       end
        
end
```
