## Supplementary Code for "Network Inference from Perturbation Time Course Data": GenerateHFData.m

```
function datas= GenerateHFData(Stim,pert,dt,TSPAN,noise)

kref=[1000,150,150,1000,150,150,1000,150,150,1000,150,150,1000,150,150,1000,150,150,1000,150,150,1000,150,150,1000,150,150,1000,150,150;];

t=linspace(0,TSPAN,floor(TSPAN/dt)+1);

Co=GetHFICs;

% Inputs=logspace(-6,-5,25);
options=[];
Inputs=[0.0030,1.2,1.2];
% Inputs=[1,1,1];

for i=1:4
    Co(1)=Stim;
%     Co(3)=Inputs(1)*pert(i,1);
%     Co(5)=Inputs(2)*pert(i,2);
%     Co(8)=Inputs(3)*pert(i,3);
kref=[1000,150,150,1000,150,150,1000,150,150,1000,150,150,1000,150,150,1000,150,150,1000,150,150,1000,150,150,1000,150,150,1000,150,150;];

    kref(3)=kref(3)*pert(i,1);
     
    kref(15)=kref(15)*pert(i,2);

    kref(27)=kref(27)*pert(i,3);
    
    [t,y]=ode15s(@HuangFerrellMAPK_S,t,Co,options,kref);
    
    % Observables divided by total protein
    Data{i}.Y(1:11,1)=(y(1:11,25).*1)./(Inputs(1));
    Data{i}.Y(1:11,2)=(y(1:11,24).*1)./(Inputs(2));
    Data{i}.Y(1:11,3)=(y(1:11,23).*1)./(Inputs(3));
%     Data{i}.Y(1:11,1)=(y(1:11,4).*1)./(Inputs(1));
%     Data{i}.Y(1:11,2)=(y(1:11,7).*1)./(Inputs(2));
%     Data{i}.Y(1:11,3)=(y(1:11,10).*1)./(Inputs(3));
    
%     Data{i}.Y(:,1)=y(:,25);
%     Data{i}.Y(:,2)=y(:,24);
%     Data{i}.Y(:,3)=y(:,23);
    Y_all{i}=y;
end

datas.Y=Data{1}.Y;
datas.YP1=Data{2}.Y;
datas.YP2=Data{3}.Y;
datas.YP3=Data{4}.Y;
datas.T=t;

D=zeros(1,3);
for a = 1:length(t)
    for b = 1:3
        for c = 1:3
        randomdata = normrnd(0,noise*(abs(datas.Y(a,b))),1,1);
        D(1,c) = datas.Y(a,b) + randomdata;
        end
        datas.Y(a,b)=median(D(1,:));
    end
end

for a = 1:length(t)
    for b = 1:3
        for c = 1:3
        randomdata = normrnd(0,noise*(abs(datas.YP1(a,b))),1,1);
        D(1,c) = datas.YP1(a,b) + randomdata;
        end
        datas.YP1(a,b)=median(D(1,:));
    end
end
        
for a = 1:length(t)
    for b = 1:3
        for c = 1:3
        randomdata = normrnd(0,noise*(abs(datas.YP2(a,b))),1,1);
        D(1,c) = datas.YP2(a,b) + randomdata;
        end
        datas.YP2(a,b)=median(D(1,:));
    end
end


for a = 1:length(t)
    for b = 1:3
        for c = 1:3
        randomdata = normrnd(0,noise*(abs(datas.YP3(a,b))),1,1);
        D(1,c) = datas.YP3(a,b) + randomdata;
        end
        datas.YP3(a,b)=median(D(1,:));
    end
end
```
