## Supplementary Code for "Network Inference from Perturbation Time Course Data": GetHFICs.m

```
function x0=GetHFICs

%Initial conditions vector
x0=zeros(25,1);
x0(1) = 3.0E-5;
x0(2) = 3.0E-4;
x0(3) = 0.0030;
x0(4) = 0.0;
x0(5) = 1.2;
x0(6) = 0.0;
x0(7) = 0.0;
x0(8) = 1.2;
x0(9) = 0.0;
x0(10) = 0.0;
x0(11) = 0.12;
x0(12) = 3.0E-4;
x0(13) = 0.0;
x0(14) = 0.0;
x0(15) = 0.0;
x0(16) = 0.0;
x0(17) = 0.0;
x0(18) = 0.0;
x0(19) = 0.0;
x0(20) = 0.0;
x0(21) = 0.0;
x0(22) = 0.0;
x0(23) = 0;
x0(24) = 0;
x0(25) = 0;
%x0(26) = 0;
```
