## Supplementary Code for "Network Inference from Perturbation Time Course Data": GetHFS.m

```
function S=GetHFS

S=[-1	1	1	0	0	0	0	0	0	0	0	0	0	0	0	0	0	0	0	0	0	0	0	0	0	0	0	0	0	0
0	0	0	-1	1	1	0	0	0	0	0	0	0	0	0	0	0	0	0	0	0	0	0	0	0	0	0	0	0	0
-1	1	0	0	0	1	0	0	0	0	0	0	0	0	0	0	0	0	0	0	0	0	0	0	0	0	0	0	0	0
0	0	1	-1	1	0	-1	1	1	0	0	0	-1	1	1	0	0	0	0	0	0	0	0	0	0	0	0	0	0	0
0	0	0	0	0	0	-1	1	0	0	0	1	0	0	0	0	0	0	0	0	0	0	0	0	0	0	0	0	0	0
0	0	0	0	0	0	0	0	1	-1	1	0	-1	1	0	0	0	1	0	0	0	0	0	0	0	0	0	0	0	0
0	0	0	0	0	0	0	0	0	0	0	0	0	0	1	-1	1	0	-1	1	1	0	0	0	-1	1	1	0	0	0
0	0	0	0	0	0	0	0	0	0	0	0	0	0	0	0	0	0	-1	1	0	0	0	1	0	0	0	0	0	0
0	0	0	0	0	0	0	0	0	0	0	0	0	0	0	0	0	0	0	0	1	-1	1	0	-1	1	0	0	0	1
0	0	0	0	0	0	0	0	0	0	0	0	0	0	0	0	0	0	0	0	0	0	0	0	0	0	1	-1	1	0
0	0	0	0	0	0	0	0	0	0	0	0	0	0	0	0	0	0	0	0	0	-1	1	1	0	0	0	-1	1	1
0	0	0	0	0	0	0	0	0	-1	1	1	0	0	0	-1	1	1	0	0	0	0	0	0	0	0	0	0	0	0
1	-1	-1	0	0	0	0	0	0	0	0	0	0	0	0	0	0	0	0	0	0	0	0	0	0	0	0	0	0	0
0	0	0	1	-1	-1	0	0	0	0	0	0	0	0	0	0	0	0	0	0	0	0	0	0	0	0	0	0	0	0
0	0	0	0	0	0	1	-1	-1	0	0	0	0	0	0	0	0	0	0	0	0	0	0	0	0	0	0	0	0	0
0	0	0	0	0	0	0	0	0	0	0	0	1	-1	-1	0	0	0	0	0	0	0	0	0	0	0	0	0	0	0
0	0	0	0	0	0	0	0	0	0	0	0	0	0	0	0	0	0	1	-1	-1	0	0	0	0	0	0	0	0	0
0	0	0	0	0	0	0	0	0	0	0	0	0	0	0	0	0	0	0	0	0	0	0	0	1	-1	-1	0	0	0
0	0	0	0	0	0	0	0	0	0	0	0	0	0	0	1	-1	-1	0	0	0	0	0	0	0	0	0	0	0	0
0	0	0	0	0	0	0	0	0	1	-1	-1	0	0	0	0	0	0	0	0	0	0	0	0	0	0	0	0	0	0
0	0	0	0	0	0	0	0	0	0	0	0	0	0	0	0	0	0	0	0	0	0	0	0	0	0	0	1	-1	-1
0	0	0	0	0	0	0	0	0	0	0	0	0	0	0	0	0	0	0	0	0	1	-1	-1	0	0	0	0	0	0
];

S(23,:)=S(10,:)+S(21,:);
S(24,:)=S(7,:)+S(17,:)+S(18,:)+S(19,:);
S(25,:)=S(4,:)+S(14,:)+S(15,:)+S(16,:);
```
