## Supplementary Code for "Network Inference from Perturbation Time Course Data": HuangFerrellMAPK_S.m

```
function xdot=HuangFerrellMAPK_S(t,x,k)


% %
% % This file is automatically generated with 
% % the System Biology Format Converter (http://sbfc.sourceforge.net/)
% % from an SBML file.
% %
% % The conversion system has the following limitations:
% %  - You may have to re order some reactions and Assignment Rules definition
% %  - Delays are not taken into account
% %  - You should change the lsode parameters (start, end, steps) to get better results
% %
% 
% %
% % The following line is there to be sure that Octave think that this file 
% % is a script and not function file
% %
% 1;
% 
% %
% % Model name = Huang1996_MAPK_ultrasens
% %
% % is urn:miriam:biomodels.db:BIOMD0000000009
% % is urn:miriam:biomodels.db:MODEL6615048798
% % isDescribedBy urn:miriam:pubmed:8816754
% %

%% Compartment: id = compartment, name = compartment, constant
	compartment_compartment=1;
%% Parameter:   id =  K_PP_norm_max, name = K_PP_norm_max
	global_par_K_PP_norm_max=0.900049;
%% assignmentRule: variable = K_PP_norm
	%x(23)=(x(10)+x(21))/(x(10)+x(9)+x(8)+x(17)+x(18)+x(21)+x(22));
%% assignmentRule: variable = rel_K_PP_max
	%x(26)=x(23)/global_par_K_PP_norm_max;
%% assignmentRule: variable = KK_PP_norm
	%x(24)=(x(7)+x(17)+x(18)+x(19))/(x(7)+x(6)+x(5)+x(17)+x(18)+x(15)+x(16)+x(19)+x(20));
%% assignmentRule: variable = KKK_P_norm
	%x(25)=(x(4)+x(15)+x(16))/(x(3)+x(4)+x(15)+x(16));
    
    %Stoichiometric matrix S pasted from Excel file
    S=GetHFS;
    
    
%% Reaction: id = r1a, name = binding of MAPKKK activator
    %v1:association and v2:dissociation
	% Local Parameter:   id =  a1, name = a1
	reaction_r1a_a1=k(1);%1000.0;
	% Local Parameter:   id =  d1, name = d1
	reaction_r1a_d1=k(2);%150.0;

	v(1)=compartment_compartment*reaction_r1a_a1*x(1)*x(3);
    v(2)=compartment_compartment*reaction_r1a_d1*x(13);
% Reaction: id = r1b, name = MAPKKK activation
    %v3
	% Local Parameter:   id =  k2, name = k2
	reaction_r1b_k2=k(3);%150.0;

	v(3)=compartment_compartment*reaction_r1b_k2*x(13);
% Reaction: id = r2a, name = binding of MAPKKK inactivator
    %v4:association and v5:dissociation
	% Local Parameter:   id =  a2, name = a2
	reaction_r2a_a2=k(4);%1000.0;
	% Local Parameter:   id =  d2, name = d2
	reaction_r2a_d2=k(5);%150.0;

	v(4)=compartment_compartment*reaction_r2a_a2*x(2)*x(4);
    v(5)=compartment_compartment*reaction_r2a_d2*x(14);
% Reaction: id = r2b, name = MAPKKK inactivation
    %v6
	% Local Parameter:   id =  k2, name = k2
	reaction_r2b_k2=k(6);%150.0;

	v(6)=compartment_compartment*reaction_r2b_k2*x(14);
% Reaction: id = r3a, name = binding P-MAPKKK and MAPKK
    %v7:association and v8:dissociation
	% Local Parameter:   id =  a3, name = a3
	reaction_r3a_a3=k(7);%1000.0;
	% Local Parameter:   id =  d3, name = d3
	reaction_r3a_d3=k(8);%150.0;

	v(7)=compartment_compartment*reaction_r3a_a3*x(5)*x(4);
    v(8)=compartment_compartment*reaction_r3a_d3*x(15);
% Reaction: id = r3b, name = phosphorylation of MAPKK
    %v9
	% Local Parameter:   id =  k3, name = k3
	reaction_r3b_k3=k(9);%150.0;

	v(9)=compartment_compartment*reaction_r3b_k3*x(15);
% Reaction: id = r4a, name = binding MAPKK-Pase and P-MAPKK
    %v10:association and v11:dissociation
	% Local Parameter:   id =  a4, name = a4
	reaction_r4a_a4=k(10);%1000.0;
	% Local Parameter:   id =  d4, name = d4
	reaction_r4a_d4=k(11);%150.0;

	v(10)=compartment_compartment*reaction_r4a_a4*x(6)*x(12);
    v(11)=compartment_compartment*reaction_r4a_d4*x(20);
% Reaction: id = r4b, name = dephosphorylation of P-MAPKK
    %v12
	% Local Parameter:   id =  k4, name = k4
	reaction_r4b_k4=k(12);%150.0;

	v(12)=compartment_compartment*reaction_r4b_k4*x(20);
% Reaction: id = r5a, name = binding P-MAPKKK and P-MAPKK
    %v13:association and v14:dissociation
	% Local Parameter:   id =  a5, name = a5
	reaction_r5a_a5=k(13);%1000.0;
	% Local Parameter:   id =  d5, name = d5
	reaction_r5a_d5=k(14);%150.0;

	v(13)=compartment_compartment*reaction_r5a_a5*x(6)*x(4);
    v(14)=compartment_compartment*reaction_r5a_d5*x(16);
% Reaction: id = r5b, name = phosphorylation of P-MAPKK
    %v15
	% Local Parameter:   id =  k5, name = k5
	reaction_r5b_k5=k(15);%150.0;

	v(15)=compartment_compartment*reaction_r5b_k5*x(16);
% Reaction: id = r6a, name = binding MAPKK-Pase and PP-MAPKK
    %v16:association and v17:dissociation
	% Local Parameter:   id =  a6, name = a6
	reaction_r6a_a6=k(16);%1000.0;
	% Local Parameter:   id =  d6, name = d6
	reaction_r6a_d6=k(17);%150.0;

	v(16)=compartment_compartment*reaction_r6a_a6*x(7)*x(12);
    v(17)=compartment_compartment*reaction_r6a_d6*x(19);
% Reaction: id = r6b, name = dephosphorylation of PP-MAPKK
    %v18
	% Local Parameter:   id =  k6, name = k6
	reaction_r6b_k6=k(18);%150.0;

	v(18)=compartment_compartment*reaction_r6b_k6*x(19);
% Reaction: id = r7a, name = binding MAPK and PP-MAPKK
    %v19:association and v20:dissociation
	% Local Parameter:   id =  a7, name = a7
	reaction_r7a_a7=k(19);%1000.0;
	% Local Parameter:   id =  d7, name = d7
	reaction_r7a_d7=k(20);%150.0;

	v(19)=compartment_compartment*reaction_r7a_a7*x(8)*x(7);
    v(20)=compartment_compartment*reaction_r7a_d7*x(17);
% Reaction: id = r7b, name = phosphorylation of MAPK
    %v21
	% Local Parameter:   id =  k7, name = k7
	reaction_r7b_k7=k(21);%150.0;

	v(21)=compartment_compartment*reaction_r7b_k7*x(17);
% Reaction: id = r8a, name = binding MAPK-Pase and P-MAPK
    %v22:association and v23:dissociation
	% Local Parameter:   id =  a8, name = a8
	reaction_r8a_a8=k(22);%1000.0;
	% Local Parameter:   id =  d8, name = d8
	reaction_r8a_d8=k(23);%150.0;

	v(22)=compartment_compartment*reaction_r8a_a8*x(9)*x(11);
    v(23)=compartment_compartment*reaction_r8a_d8*x(22);
% Reaction: id = r8b, name = dephosphorylation of P-MAPK
    %v24
	% Local Parameter:   id =  k8, name = k8
	reaction_r8b_k8=k(24);%150.0;

	v(24)=compartment_compartment*reaction_r8b_k8*x(22);
% Reaction: id = r9a, name = binding PP-MAPKK and P-MAPK
    %v25:association and v26:dissociation
	% Local Parameter:   id =  a9, name = a9
	reaction_r9a_a9=k(25);%1000.0;
	% Local Parameter:   id =  d9, name = d9
	reaction_r9a_d9=k(26);%150.0;

	v(25)=compartment_compartment*reaction_r9a_a9*x(9)*x(7);
    v(26)=compartment_compartment*reaction_r9a_d9*x(18);
% Reaction: id = r9b, name = phosphorylation of P-MAPK
    %v27
	% Local Parameter:   id =  k9, name = k9
	reaction_r9b_k9=k(27);%150.0;

	v(27)=compartment_compartment*reaction_r9b_k9*x(18);
% Reaction: id = r10a, name = binding MAPK-Pase and PP-MAPK
    %v28:association and v29:dissociation
	% Local Parameter:   id =  a10, name = a10
	reaction_r10a_a10=k(28);%1000.0;
	% Local Parameter:   id =  d10, name = d10
	reaction_r10a_d10=k(29);%150.0;

	v(28)=compartment_compartment*reaction_r10a_a10*x(10)*x(11);
    v(29)=compartment_compartment*reaction_r10a_d10*x(21);
% Reaction: id = r10b, name = dephosphorylation of PP-MAPK
    %v30
	% Local Parameter:   id =  k10, name = k10
	reaction_r10b_k10=k(30);%150.0;

	v(30)=compartment_compartment*reaction_r10b_k10*x(21);
	xdot=S*v';
    
%     zeros(26,1);
% 	% Species:   id = E1, name = MAPKKK activator, affected by kineticLaw
% 
% 	xdot(1) = (1/(compartment_compartment))*((-1.0 * reaction_r1a) + ( 1.0 * reaction_r1b));
% 
% 	% Species:   id = E2, name = MAPKKK inactivator, affected by kineticLaw
% 
% 	xdot(2) = (1/(compartment_compartment))*((-1.0 * reaction_r2a) + ( 1.0 * reaction_r2b));
% 
% 	% Species:   id = KKK, name = MAPKKK, affected by kineticLaw
% 
% 	xdot(3) = (1/(compartment_compartment))*((-1.0 * reaction_r1a) + ( 1.0 * reaction_r2b));
% 
% 	% Species:   id = P_KKK, name = P-MAPKKK, affected by kineticLaw
% 
% 	xdot(4) = (1/(compartment_compartment))*(( 1.0 * reaction_r1b) + (-1.0 * reaction_r2a) + (-1.0 * reaction_r3a) + ( 1.0 * reaction_r3b) + (-1.0 * reaction_r5a) + ( 1.0 * reaction_r5b));
% 
% 	% Species:   id = KK, name = MAPKK, affected by kineticLaw
% 
% 	xdot(5) = (1/(compartment_compartment))*((-1.0 * reaction_r3a) + ( 1.0 * reaction_r4b));
% 
% 	% Species:   id = P_KK, name = P-MAPKK, affected by kineticLaw
% 
% 	xdot(6) = (1/(compartment_compartment))*(( 1.0 * reaction_r3b) + (-1.0 * reaction_r4a) + (-1.0 * reaction_r5a) + ( 1.0 * reaction_r6b));
% 
% 	% Species:   id = PP_KK, name = PP-MAPKK, affected by kineticLaw
% 
% 	xdot(7) = (1/(compartment_compartment))*(( 1.0 * reaction_r5b) + (-1.0 * reaction_r6a) + (-1.0 * reaction_r7a) + ( 1.0 * reaction_r7b) + (-1.0 * reaction_r9a) + ( 1.0 * reaction_r9b));
% 
% 	% Species:   id = K, name = MAPK, affected by kineticLaw
% 
% 	xdot(8) = (1/(compartment_compartment))*((-1.0 * reaction_r7a) + ( 1.0 * reaction_r8b));
% 
% 	% Species:   id = P_K, name = P-MAPK, affected by kineticLaw
% 
% 	xdot(9) = (1/(compartment_compartment))*(( 1.0 * reaction_r7b) + (-1.0 * reaction_r8a) + (-1.0 * reaction_r9a) + ( 1.0 * reaction_r10b));
% 
% 	% Species:   id = PP_K, name = PP-MAPK, affected by kineticLaw
% 
% 	xdot(10) = (1/(compartment_compartment))*(( 1.0 * reaction_r9b) + (-1.0 * reaction_r10a));
% 
% 	% Species:   id = KPase, name = MAPK-Pase, affected by kineticLaw
% 
% 	xdot(11) = (1/(compartment_compartment))*((-1.0 * reaction_r8a) + ( 1.0 * reaction_r8b) + (-1.0 * reaction_r10a) + ( 1.0 * reaction_r10b));
% 
% 	% Species:   id = KKPase, name = MAPKK-Pase, affected by kineticLaw
% 
% 	xdot(12) = (1/(compartment_compartment))*((-1.0 * reaction_r4a) + ( 1.0 * reaction_r4b) + (-1.0 * reaction_r6a) + ( 1.0 * reaction_r6b));
% 
% 	% Species:   id = E1_KKK, name = activ_MAPKKK, affected by kineticLaw
% 
% 	xdot(13) = (1/(compartment_compartment))*(( 1.0 * reaction_r1a) + (-1.0 * reaction_r1b));
% 
% 	% Species:   id = E2_P_KKK, name = inact_P-MAPKKK, affected by kineticLaw
% 
% 	xdot(14) = (1/(compartment_compartment))*(( 1.0 * reaction_r2a) + (-1.0 * reaction_r2b));
% 
% 	% Species:   id = P_KKK_KK, name = P-MAPKKK_MAPKK, affected by kineticLaw
% 
% 	xdot(15) = (1/(compartment_compartment))*(( 1.0 * reaction_r3a) + (-1.0 * reaction_r3b));
% 
% 	% Species:   id = P_KKK_P_KK, name = P-MAPKKK_P-MAPKK, affected by kineticLaw
% 
% 	xdot(16) = (1/(compartment_compartment))*(( 1.0 * reaction_r5a) + (-1.0 * reaction_r5b));
% 
% 	% Species:   id = PP_KK_K, name = PP-MAPKK_MAPK, affected by kineticLaw
% 
% 	xdot(17) = (1/(compartment_compartment))*(( 1.0 * reaction_r7a) + (-1.0 * reaction_r7b));
% 
% 	% Species:   id = PP_KK_P_K, name = PP-MAPKK_P-MAPK, affected by kineticLaw
% 
% 	xdot(18) = (1/(compartment_compartment))*(( 1.0 * reaction_r9a) + (-1.0 * reaction_r9b));
% 
% 	% Species:   id = KKPase_PP_KK, name = MAPKK-Pase_PP-MAPKK, affected by kineticLaw
% 
% 	xdot(19) = (1/(compartment_compartment))*(( 1.0 * reaction_r6a) + (-1.0 * reaction_r6b));
% 
% 	% Species:   id = KKPase_P_KK, name = MAPKK-Pase_P-MAPKK, affected by kineticLaw
% 
% 	xdot(20) = (1/(compartment_compartment))*(( 1.0 * reaction_r4a) + (-1.0 * reaction_r4b));
% 
% 	% Species:   id = KPase_PP_K, name = MAPK-Pase_PP-MAPK, affected by kineticLaw
% 
% 	xdot(21) = (1/(compartment_compartment))*(( 1.0 * reaction_r10a) + (-1.0 * reaction_r10b));
% 
% 	% Species:   id = KPase_P_K, name = MAPK-Pase_P-MAPK, affected by kineticLaw
% 
% 	xdot(22) = (1/(compartment_compartment))*(( 1.0 * reaction_r8a) + (-1.0 * reaction_r8b));
% 
% 	% Species:   id = K_PP_norm, name = K_PP_norm, defined in a rule 
% 	xdot(23) = xdot(10)+xdot(21);
% 
% 	% Species:   id = KK_PP_norm, name = KK_PP_norm, defined in a rule 
% 	xdot(24) = xdot(7)+xdot(17)+xdot(18)+xdot(19);
% 
% 	% Species:   id = KKK_P_norm, name = KKK_P_norm, defined in a rule 
% 	xdot(25) = xdot(4)+xdot(15)+xdot(16);
% 
% 	% Species:   id = rel_K_PP_max, name = relative maximal K_PP, defined in a rule 
	%xdot(26) = 1;

end
% function z=pow(x,y)
%     z=x^y;
% end
% function z=root(x,y)
%     z=y^(1/x)
% end
% function z = piecewise(varargin)
% 	numArgs = nargin;
% 	result = 0;
% 	foundResult = 0;
% 	for k=1:2: numArgs-1
% 		if varargin{k+1} == 1
% 			result = varargin{k};
% 			foundResult = 1;
% 			break;
%         end
% 	end
% 	if foundResult == 0
% 		result = varargin{numArgs};
%     end 
%     z = result;
% end 


%Creating linespace
%t=linspace(0,90,100);

%Solving equations
%x=lsode("f",x0,t);

%ploting the results
%plot(t,x);
```
