## Supplementary Code for "Network Inference from Perturbation Time Course Data": Make_figure_4.m

```
%% Generate HF data and estimates 
Stim=2.5E-6;
pert=[1,1,1;0,1,1;1,0,1;1,1,0];
dt=25;
TSPAN=250;
data_3_node=GenerateHFData(Stim,pert,dt,TSPAN,0);
% Obtain estimates for HF data 
multistart=1;
for n=1:multistart
    [preds0{1,1}(n,:),preds0_SSE{1,1}(n,1)]= estimateHF(data_3_node);
n
end
save ("HFdata.mat", "data_3_node", "preds0")
%% Generate fit data from estimates
load('HFdata.mat')
guessHF=estimateHFdata(data_3_node,preds0);
save ("guessHFdata.mat", "guessHF");
%% Plot 4C
load('HFdata.mat')
load('guessHFdata.mat')

figure;
subplot(1,4,1);
hold on

plot(data_3_node.T,data_3_node.Y(:,1),'b.', 'MarkerSize', 15);
plot(data_3_node.T,data_3_node.Y(:,2),'r.', 'MarkerSize', 15);
plot(data_3_node.T,data_3_node.Y(:,3),'y.', 'MarkerSize', 15);
plot(guessHF.T,guessHF.Y(:,1),'b--', 'MarkerSize', 15);
plot(guessHF.T,guessHF.Y(:,2),'r--', 'MarkerSize', 15);
plot(guessHF.T,guessHF.Y(:,3),'y--', 'MarkerSize', 15);

title("Vehicle",'FontSize',9);
ylim([0 0.06])

subplot(1,4,2);
hold on

plot(data_3_node.T,data_3_node.YP1(:,1),'b.', 'MarkerSize', 15);
plot(data_3_node.T,data_3_node.YP1(:,2),'r.', 'MarkerSize', 15);
plot(data_3_node.T,data_3_node.YP1(:,3),'y.', 'MarkerSize', 15);
plot(data_3_node.T,data_3_node.YP1(:,1),'b--', 'MarkerSize', 15);
plot(guessHF.T,guessHF.YP1(:,1),'r--', 'MarkerSize', 15);
plot(guessHF.T,guessHF.YP1(:,2),'y--', 'MarkerSize', 15);
title("Perturb MAPKKK", 'FontSize',9);
ylim([0 0.06])

subplot(1,4,3);
hold on

plot(data_3_node.T,data_3_node.YP2(:,1),'b.', 'MarkerSize', 15);
plot(data_3_node.T,data_3_node.YP2(:,2),'r.', 'MarkerSize', 15);
plot(data_3_node.T,data_3_node.YP2(:,3),'y.', 'MarkerSize', 15);
plot(guessHF.T,guessHF.YP2(:,1),'b--', 'MarkerSize', 15);
plot(data_3_node.T,data_3_node.YP2(:,2),'r--', 'MarkerSize', 15);
plot(guessHF.T,guessHF.YP2(:,2),'y--', 'MarkerSize', 15);
title("Perturb MAPKK", 'FontSize',9);
ylim([0 0.06])

subplot(1,4,4);
hold on

plot(data_3_node.T,data_3_node.YP3(:,1),'b.', 'MarkerSize', 15);
plot(data_3_node.T,data_3_node.YP3(:,2),'r.', 'MarkerSize', 15);
plot(data_3_node.T,data_3_node.YP3(:,3),'y.', 'MarkerSize', 15);
plot(guessHF.T,guessHF.YP3(:,1),'b--', 'MarkerSize', 15);
plot(guessHF.T,guessHF.YP3(:,2),'r--', 'MarkerSize', 15);
plot(data_3_node.T,data_3_node.YP3(:,3),'y--', 'MarkerSize', 15);
title("Perturb MAPK", 'FontSize',9);
ylim([0 0.06])

res = 300;
set(gcf,'paperunits','inches','PaperPosition',[0 0 6 2]);
% set(gcf,'paperunits','inches','PaperPosition',[0 0 3.5 1.5]);
print('4C.tiff','-dtiff',['-r' num2str(res)]);
```
