## Supplementary Code for "Network Inference from Perturbation Time Course Data": addgtvals.m

```
function St=addgtvals(St)
qpred=zeros(16,12);
% Add ground truth values to outmap
St.best_est.qpred0(1,:) = [1 -1 0 0 0 1 -1 0 0 1 1 -1];
St.best_est.qpred0(2,:) = [1 -1 0 0 0 1 -1 0 0 1 -1 -1];
St.best_est.qpred0(3,:) = [1 -1 0 0 0 1 -1 0 0 -1 1 -1];
St.best_est.qpred0(4,:) = [1 -1 0 0 0 1 -1 0 1 -1 -1 -1];
St.best_est.qpred0(5,:) = [1 -1 0 0 0 1 -1 0 0 1 1 -1];
St.best_est.qpred0(6,:) = [1 -1 0 0 0 1 -1 0 1 1 -1 -1];
St.best_est.qpred0(7,:) = [1 -1 0 0 0 1 -1 0 1 -1 1 -1];
St.best_est.qpred0(8,:) = [1 -1 0 0 0 1 -1 0 1 -1 -1 -1];
St.best_est.qpred0(9,:) = [1 -1 0 0 1 -1 -1 0 -1 1 1 -1];
St.best_est.qpred0(10,:) = [1 -1 0 0 1 -1 -1 0 1 1 -1 -1];
St.best_est.qpred0(11,:) = [1 -1 0 0 1 -1 -1 0 1 -1 1 -1];
St.best_est.qpred0(12,:) = [1 -1 0 0 1 -1 -1 0 1 -1 -1 -1];
St.best_est.qpred0(13,:) = [1 -1 0 0 1 -1 -1 0 1 1 1 -1];
St.best_est.qpred0(14,:) = [1 -1 0 0 1 -1 -1 0 1 1 -1 -1];
St.best_est.qpred0(15,:) = [1 -1 0 0 1 -1 -1 0 1 -1 1 -1];
St.best_est.qpred0(16,:) = [1 -1 0 0 1 -1 -1 0 1 -1 -1 -1];
end
```
