## Supplementary Code for "Network Inference from Perturbation Time Course Data": calcffljacobian.m

```
%Parameter Values
alphay = 1;
alphaz = 1;
betay = 1;
betaz = 1;
H = 2;
tmax=5;
timeint=tmax/10;
Ks=dlmread('Rich_Kout_K25.txt');
Bs=dlmread('Rich_Bout.txt');
pert1=0.5;
pert2=0.5;
pert3=0.5;
m=1;
% fmin=zeros(count,10);
% finSSE=zeros(count,1);
y0=zeros(16,3);
yp1=zeros(16,3);
yp2=zeros(16,3);
yp3=zeros(16,3);

data_nostim=feedforwarddatagenerator_nostim(timeint,100,Ks,Bs,pert1,pert2,pert3,0);
% jac=calcjacob(data_nostim,Ks,Bs);

for i = 1:2
    for j = 1:2
        for k = 1:2
            for l = 1:2
                key = i*1000+j*100+k*10+l;
                nostimdata=data_nostim(key);              
                y0(m,:)=nostimdata.Y(100,:);
                yp1(m,:)=nostimdata.YP1(100,:);
                yp2(m,:)=nostimdata.YP2(100,:);
                yp3(m,:)=nostimdata.YP3(100,:);
                m=m+1;
            end
        end
    end
end
                

data0=feedforwarddatagenerator_stim(timeint,tmax,y0,yp1,yp2,yp3,Ks,Bs,pert1,pert2,pert3,0);
op=1;
for i = 1:2
    for j = 1:2
        for k =1:2
            for l =1:2
                
                key = i*1000+j*100+k*10+l;
                K=Ks(op,:);
                
                Kxy = K(1);%.1;
                Kyz = K(2);%.1;
                Kxz = K(3);%.1;

                ijklstruct = data0(key);
%                 ijklstruct = data_nostim(key);
                T = ijklstruct.T;
                Y0 = ijklstruct.Y;
%                  Y0 = ijklstruct.YP1;
%                  Y0 = ijklstruct.YP2;
                 
%                  Y0 = ijklstruct.YP3;
                
%                 display(length(T))
                
                x = Y0(:,1);
                y = Y0(:,2);
                z = Y0(:,3);
                
                jac = zeros(9,length(T));
                
                jac(1,:) = -1;
                jac(2,:) = 0;
                jac(3,:) = 0;
                jac(5,:) = -1;
                jac(6,:) = 0;
                jac(9,:) = -1;
                
                if i == 1       %% X -> Y Activator
                   
                    for t = 1:length(T)
                        
                        jac(4,t) = (H*x(t)^(H-1)/(Kxy^H))/(1+(x(t)/Kxy)^H)^2;
%                         jac(4,t) = (H*x(t)^(H-1)/((Kxy^H)*(1+(x(t)/Kxy)^H)^2));

                                            
                    end
                                        
                else            %% X -> Y Repressor
                                        
                    for t = 1:length(T)
                        
                        jac(4,t) = (-H*x(t)^(H-1)/(Kxy^H))/(1+(x(t)/Kxy)^H)^2;
                                            
                    end
                    
                end
                
                if j == 1       %% AND Gate
                    
                    if k == 1 && l == 1         %% %% X -> Z Activator  Y -> Z Activator
                    
                        for t = 1:length(T)
                        
                            jac(7,t) = (H*x(t)^(H-1)/Kxz^H)*(y(t)^H/Kyz^H)/(((1+x(t)^H/Kxz^H)^2)*((1+y(t)^H/Kyz^H)));
                            
                            jac(8,t) = (x(t)^H/Kxz^H)*(H*y(t)^(H-1)/Kyz^H)/(((1+x(t)^H/Kxz^H))*((1+y(t)^H/Kyz^H)^2));                            
                            
                        end
                                      
                    elseif k == 1 && l == 2     %% X -> Z Activator  Y -| Z Repressor
                        
                        for t = 1:length(T)
                        
                            jac(7,t) = (H*x(t)^(H-1)/Kxz^H)/(((1+x(t)^H/Kxz^H)^2)*((1+y(t)^H/Kyz^H)));
                            
                            jac(8,t) = (-1)*(x(t)^H/Kxz^H)*(H*y(t)^(H-1)/Kyz^H)/(((1+x(t)^H/Kxz^H))*((1+y(t)^H/Kyz^H)^2));                            
                            
                        end
                      
                    elseif k == 2 && l == 1     %% X -| Z Repressor  Y -> Z Activator

                        for t = 1:length(T)
                        
                            jac(7,t) = (-1)*(H*x(t)^(H-1)/Kxz^H)*(y(t)^H/Kyz^H)/(((1+x(t)^H/Kxz^H)^2)*((1+y(t)^H/Kyz^H)));
                            
                            jac(8,t) = (H*y(t)^(H-1)/Kyz^H)/(((1+x(t)^H/Kxz^H))*((1+y(t)^H/Kyz^H)^2));                            
                            
                        end  
                        
                    else                        %% X -| Z Repressor  Y -| Z Repressor
                        
                        for t = 1:length(T)
                        
                            jac(7,t) = (-1)*(H*x(t)^(H-1)/Kxz^H)/(((1+x(t)^H/Kxz^H)^2)*((1+y(t)^H/Kyz^H)));
                            
                            jac(8,t) = (-1)*(H*y(t)^(H-1)/Kyz^H)/(((1+x(t)^H/Kxz^H))*((1+y(t)^H/Kyz^H)^2));                            
                            
                        end
                        
                    end
                    
                else            %% OR Gate
                    
                    if k == 1 && l == 1         %% %% X -> Z Activator  Y -> Z Activator
                    
                        for t = 1:length(T)
                        
                            jac(7,t) = (H*x(t)^(H-1)/Kxz^H)/((1+x(t)^H/Kxz^H+y(t)^H/Kyz^H)^2);

                            jac(8,t) = (H*y(t)^(H-1)/Kyz^H)/((1+x(t)^H/Kxz^H+y(t)^H/Kyz^H)^2);
                            
                        end
                                      
                    elseif k == 1 && l == 2     %% X -> Z Activator  Y -| Z Repressor
                        
                        for t = 1:length(T)
                        
                            jac(7,t) = (H*x(t)^(H-1)*y(t)^H/(Kxz^H*Kyz^H))/((1+x(t)^H/Kxz^H+y(t)^H/Kyz^H)^2);

                            jac(8,t) = (-H*x(t)^H*y(t)^(H-1)/(Kxz^H*Kyz^H)-H*y(t)^(H-1)/Kyz^H)/((1+x(t)^H/Kxz^H+y(t)^H/Kyz^H)^2);
                            
                        end
                      
                    elseif k == 2 && l == 1     %% X -| Z Repressor  Y -> Z Activator

                        for t = 1:length(T)
                        
                            jac(7,t) = (-H*x(t)^H*y(t)^(H-1)/(Kxz^H*Kyz^H)-H*x(t)^(H-1)/Kxz^H)/((1+x(t)^H/Kxz^H+y(t)^H/Kyz^H)^2);
                            
                            jac(8,t) = (H*x(t)^H*y(t)^(H-1)/(Kxz^H*Kyz^H))/((1+x(t)^H/Kxz^H+y(t)^H/Kyz^H)^2);
                            
                        end  
                        
                    else                        %% X -| Z Repressor  Y -| Z Repressor
                        
                        for t = 1:length(T)
                        
                            jac(7,t) = (-2*H*x(t)^(H-1)/Kxz^H)/((1+x(t)^H/Kxz^H+y(t)^H/Kyz^H)^2);
                            
                            jac(8,t) = (-2*H*y(t)^(H-1)/Kyz^H)/((1+x(t)^H/Kxz^H+y(t)^H/Kyz^H)^2);
                            
                        end
                        
                    end                    
                    
                end
                
                Stim1= ones(11,1);
                Stim2= zeros(11,1);
                Stim3= zeros(11,1);
                jacinv = jac';
                Jacob=[Stim1 jacinv(:,1:9)];
                Jacob_Ian{op,1}=Jacob;
                meanjac21(op,1)=mean(jac(4,:));
                meanjac31(op,1)=mean(jac(7,:));
                meanjac32(op,1)=mean(jac(8,:));
figure;
                Tslid(length(fmin_data{1,1}(:,1)),1)=zeros();
                Tslid(1,1)=((length(T)-(length(fmin_data{1,1}(:,1))))/2)*timeint;

                for p=2:length(fmin_data{1,1}(:,1))
                Tslid(p,1)=Tslid(p-1,1)+timeint;
                end
                for p=1:10    
                subplot(3,4,p)
                plot(T,Jacob(:,p));
                hold on
                plot(Tslid,fmin_data{op,1}(:,p),".", "Markersize", 15);
                hold off
                xlim([0,tmax])
                ylim([-2,2])
                end
                
                
                op=op+1;

%                 suptitle(strcat('FFL Case ',num2str(key)));
                set(gcf, 'Units', 'Normalized', 'OuterPosition', [0, 0.04, 10, 0.96]);
%                 saveas(gcf,strcat('FFLCase',num2str(key),'.tif'));
            end
        end
    end
end
% disp(meanjac31(12,1));
% disp(meanjac32(12,1));
```
