## Supplementary Code for "Network Inference from Perturbation Time Course Data": estimateffl.m

```
function [fmin,finSSE]=estimateffl (data,timeint,tmax,y0,yp1,yp2,yp3)

% S2=y0(1,2);
% S3=y0(1,3);
tic
finSSE=zeros(1,100);
Y=data.Y;
YP1=data.YP1;
YP2=data.YP2;
YP3=data.YP3;
t=data.T;

    poly_order=length(t)-1;

poly_YP1=polyfit(t,YP1(:,1),poly_order);
poly_YP2=polyfit(t,YP2(:,2),poly_order);
poly_YP3=polyfit(t,YP3(:,3),poly_order);
        
lasttimepoint = length(t);
ss(1) = Y(lasttimepoint,1);
ss(2) = Y(lasttimepoint,2);
ss(3) = Y(lasttimepoint,3);

fc = zeros(3,3);
fc(:,1) = YP1(lasttimepoint,1:3)';
fc(:,2) = YP2(lasttimepoint,1:3)';
fc(:,3) = YP3(lasttimepoint,1:3)';


Rp = zeros(3,3);

for i = 1:3
    for j = 1:3

        frac = fc(i,j)/ss(i);

        Rp(i,j) = 2*(frac-1)/(frac+1);

    end
end

Rpinv = inv(Rp);
dgRpinv = diag(diag(Rpinv));

ssjac = ((-1)*dgRpinv\Rpinv);
ssjacvec = reshape(ssjac',[1,9]);

        
%         options = optimoptions('fmincon','MaxFunEval',6000);
        conoptions = optimoptions('fmincon','MaxFunctionEvaluations',10000);

lb = [-10 -10 -10 -10 -10 -10 -10 -10 -10 -10 -10 10];
ub = [10 10 10 10 10 10 10 10 10 10 10 10];


for i = 1:9
   
    newindex = i + floor((i-1)/3)+1;
    
    if abs(ssjacvec(i)) < 0.01
        lb(newindex) = 0;
        ub(newindex) = 0;
    end
    
end


try
        initial=[2*rand(1),4*rand(1)-2,4*rand(1)-2,4*rand(1)-2,2*rand(1),4*rand(1)-2,4*rand(1)-2,4*rand(1)-2,2*rand(1),4*rand(1)-2,4*rand(1)-2,4*rand(1)-2];

          [fmin,finSSE] =  fmincon(@nestedfun,initial,[],[],[],[],lb,ub,[],conoptions);

catch
    try
        initial=[2*rand(1),4*rand(1)-2,4*rand(1)-2,4*rand(1)-2,2*rand(1),4*rand(1)-2,4*rand(1)-2,4*rand(1)-2,2*rand(1),4*rand(1)-2,4*rand(1)-2,4*rand(1)-2];
%         [fmin,finSSE] = fminunc(@nestedfun,initial,options);
          [fmin,finSSE] =  fmincon(@nestedfun,initial,[],[],[],[],lb,ub,[],conoptions);

    catch
        try
        initial=[2*rand(1),4*rand(1)-2,4*rand(1)-2,4*rand(1)-2,2*rand(1),4*rand(1)-2,4*rand(1)-2,4*rand(1)-2,2*rand(1),4*rand(1)-2,4*rand(1)-2,4*rand(1)-2];
%         [fmin,finSSE] = fminunc(@nestedfun,initial,options);
          [fmin,finSSE] =  fmincon(@nestedfun,initial,[],[],[],[],lb,ub,[],conoptions);

        catch
        end
    end
end
fmin(5)=-(fmin(6)*Y(1,1)+fmin(7)*Y(1,2)+fmin(8)*Y(1,3));
fmin(9)=-(fmin(10)*Y(1,1)+fmin(11)*Y(1,2)+fmin(12)*Y(1,3));

        
function SSE = nestedfun(r2)
        
    % Solve the ODEs
                S2=-(r2(6)*Y(1,1)+r2(7)*Y(1,2)+r2(8)*Y(1,3));
               
                S3=-(r2(10)*Y(1,1)+r2(11)*Y(1,2)+r2(12)*Y(1,3));

                
[t,guessY] = ode15s(@odeqns,0:timeint:10,[Y(1,1);Y(1,2);Y(1,3)],odeset('RelTol',1e-6),r2);
[t,guessYP1] = ode15s(@odep1,0:timeint:10,[YP1(1,2);YP1(1,3)],odeset('RelTol',1e-6),r2);
[t,guessYP2] = ode15s(@odep2,0:timeint:10,[YP2(1,1);YP2(1,3)],odeset('RelTol',1e-6),r2);
[t,guessYP3] = ode15s(@odep3,0:timeint:10,[YP3(1,1);YP3(1,2)],odeset('RelTol',1e-6),r2);

% [t,guessY] = ode15s(@odeqns,0:timeint:tmax,[0;0;0],odeset('RelTol',1e-6),r2);
% [t,guessYP1] = ode15s(@odep1,0:timeint:tmax,[0;0],odeset('RelTol',1e-6),r2);
% [t,guessYP2] = ode15s(@odep2,0:timeint:tmax,[0;0],odeset('RelTol',1e-6),r2);
% [t,guessYP3] = ode15s(@odep3,0:timeint:tmax,[0;0],odeset('RelTol',1e-6),r2);

 SSE = (sum(sum((Y-guessY).^2)+sum((YP1(:,2)-guessYP1(:,1)).^2)+sum((YP1(:,3)-guessYP1(:,2)).^2)+sum((YP2(:,1)-guessYP2(:,1)).^2)+sum((YP2(:,3)-guessYP2(:,2)).^2)+sum((YP3(:,1)-guessYP3(:,1)).^2)+sum((YP3(:,2)-guessYP3(:,2)).^2))).^0.5;
 %   SSE = (sum(sum(((Y(2:11,:)-guessY(2:11,:))./Y(2:11,:))).^2)+sum(((YP1(2:11,2)-guessYP1(2:11,1))./YP1(2:11,2)).^2)+sum(((YP1(2:11,3)-guessYP1(2:11,2))./YP1(2:11,3)).^2)+sum(((YP2(2:11,1)-guessYP2(2:11,1))./YP2(2:11,1)).^2)+sum(((YP2(2:11,3)-guessYP2(2:11,2))./YP2(2:11,3)).^2)+sum(((YP3(2:11,1)-guessYP3(2:11,1))./YP3(2:11,1)).^2)+sum(((YP3(2:11,2)-guessYP3(2:11,2))./YP3(2:11,2)).^2)).^0.5;
   
    function dzdt1 = odeqns(t,z,r)
        dzdt1 = [
            (r(1)+r(2)*z(1)+r(3)*z(2)+r(4)*z(3))
            (S2+r(6)*z(1)+r(7)*z(2)+r(8)*z(3))
            (S3+r(10)*z(1)+r(11)*z(2)+r(12)*z(3))
        ];
    end
    function dzdt1 = odep1(t,z,r)
        Y_pert=polyval(poly_YP1,t);   
        dzdt1 = [
            (S2+r(6)*Y_pert+r(7)*z(1)+r(8)*z(2))
            (S3+r(10)*Y_pert+r(11)*z(1)+r(12)*z(2))
                 ];
    end
       function dzdt1 = odep2(t,z,r)
        Y_pert=polyval(poly_YP2,t);
        dzdt1 = [
            (r(1)+r(2)*z(1)+r(3)*Y_pert+r(4)*z(2))
            (S3+r(10)*z(1)+r(11)*Y_pert+r(12)*z(2))
        ];
       end
       function dzdt1 = odep3(t,z,r)
        Y_pert=polyval(poly_YP3,t);
        dzdt1 = [
            (r(1)+r(2)*z(1)+r(3)*z(2)+r(4)*Y_pert)
            (S2+r(6)*z(1)+r(7)*z(2)+r(8)*Y_pert)
        ];
       end
        
end
endend
```
