## Supplementary Code for "Network Inference from Perturbation Time Course Data": feedforwarddatagenerator_MB5.m

```
function out = feedforwarddatagenerator_MB5(timeint,tmax,y0,yp1,yp2,yp3,Ks,Bs,pert1,pert2,pert3,noise)

options = odeset('RelTol',1e-6);

ffdataMapObj = containers.Map('KeyType','double','ValueType','any');
op=1;

for i = 1:2
    for j = 1:2
        for k = 1:2
            for l = 1:2
                
                key = i*1000+j*100+k*10+l;
                K=Ks(op,:);
                B=Bs(op,:);
                                                 
                [Y,YP1,YP2,YP3,T] = feedforwardloop_nostim(timeint,tmax,y0(op,:),yp1(op,:),yp2(op,:),yp3(op,:),[i j k l],K,B,pert1,pert2,pert3);
                op=op+1;

                
% D=zeros(1,3);
% for a = 1:length(T)
%     for b = 1:3
%         for c = 1:3
%         randomdata = normrnd(0,noise*(abs(Y(a,b))),1,1);
%         D(1,c) = Y(a,b) + randomdata;
%         end
%         Y(a,b)=median(D(1,:));
%     end
% end
% 
% for a = 1:length(T)
%     for b = 1:3
%         for c = 1:3
%         randomdata = normrnd(0,noise*(abs(YP1(a,b))),1,1);
%         D(1,c) = YP1(a,b) + randomdata;
%         end
%         YP1(a,b)=median(D(1,:));
%     end
% end
%         
% for a = 1:length(T)
%     for b = 1:3
%         for c = 1:3
%         randomdata = normrnd(0,noise*(abs(YP2(a,b))),1,1);
%         D(1,c) = YP2(a,b) + randomdata;
%         end
%         YP2(a,b)=median(D(1,:));
%     end
% end
% 
% 
% for a = 1:length(T)
%     for b = 1:3
%         for c = 1:3
%         randomdata = normrnd(0,noise*(abs(YP3(a,b))),1,1);
%         D(1,c) = YP3(a,b) + randomdata;
%         end
%         YP3(i,j)=median(D(1,:));
%     end
% end
                
                casestruct.Y = Y;
                casestruct.YP1 = YP1;
                casestruct.YP2 = YP2;
                casestruct.YP3 = YP3;
                casestruct.T = T;

                ffdataMapObj(key) = casestruct;
                
            end
        end
    end
end

out = ffdataMapObj;
end
```
