## Supplementary Code for "Network Inference from Perturbation Time Course Data": feedforwardloop.m

```
function [Y,YP1,YP2,YP3,t]= feedforwardloop(timeint,tmax,y0,flags,K,B,pert1,pert2,pert3)
options = odeset('RelTol',1e-6);
[t,Y] = ode15s(@ffl_nopert,0:timeint:tmax,y0,options,flags,K,B);

if length(t)>10
    poly_order=9;
else
    poly_order=length(t)-1;
end

YP1=zeros((1+(tmax/timeint)),3);
YP2=zeros((1+(tmax/timeint)),3);
YP3=zeros((1+(tmax/timeint)),3);

YP1(:,1)=pert1*Y(:,1);
YP2(:,2)=pert2*Y(:,2);
YP3(:,3)=pert3*Y(:,3);

poly_YP1=polyfit(t,YP1(:,1),poly_order);
% % % % % % % fit_YP1=polyval(poly_YP1,t2);
poly_YP2=polyfit(t,YP2(:,2),poly_order);
% % % % % % % fit_YP2=polyval(poly_YP2,t2);
poly_YP3=polyfit(t,YP3(:,3),poly_order);

[t,YP1_data] = ode15s(@ffl_pert1,0:timeint:tmax,[y0(2);y0(3)],options,flags,K,B);
[t,YP2_data] = ode15s(@ffl_pert2,0:timeint:tmax,[y0(1);y0(3)],options,flags,K,B);
[t,YP3_data] = ode15s(@ffl_pert3,0:timeint:tmax,[y0(1);y0(2)],options,flags,K,B);

YP1(:,2)=YP1_data(:,1);
YP1(:,3)=YP1_data(:,2);
YP2(:,1)=YP2_data(:,1);
YP2(:,3)=YP2_data(:,2);
YP3(:,1)=YP3_data(:,1);
YP3(:,2)=YP3_data(:,2);

function dm = ffl_nopert(t,m,flags,K,B)

x = m(1);
y = m(2);
z = m(3);

%Parameter Values
Bx = B(1);
By = B(2);%1;%0
Bz = B(3);%1;%0
alphay = 1;
alphaz = 1;
betay = 1;
betaz = 1;
H = 2;
Kxy = K(1);%.1;
Kyz = K(2);%.1;
Kxz = K(3);%.1;

%dx = 0;
dx = -x;
% dy = By + betay*f(x,Kxy,H,flags(1) - alphay*y);
dy = (betay*f(x,Kxy,H,flags(1))) - alphay*y;
% dz = Bz + betaz*G(x,Kxz,Kyz,y,H,flags) - alphaz*z;
dz = (betaz*G(x,Kxz,Kyz,y,H,flags)) - alphaz*z;
dm = [dx; dy; dz];

end

function dm = ffl_pert1(t,m,flags,K,B)

y = m(1);
z = m(2);

%Parameter Values
Bx = B(1);
By = B(2);%1;%0
Bz = B(3);%1;%0
alphay = 1;
alphaz = 1;
betay = 1;
betaz = 1;
H = 2;
Kxy = K(1);%.1;
Kyz = K(2);%.1;
Kxz = K(3);%.1;

%dx = 0;
Y_pert= polyval(poly_YP1,t);
dy = (betay*f(Y_pert,Kxy,H,flags(1))) - alphay*y;
dz = betaz*G(Y_pert,Kxz,Kyz,y,H,flags) - alphaz*z;

dm = [dy; dz];

end

function dm = ffl_pert2(t,m,flags,K,B)

x = m(1);
z = m(2);

%Parameter Values
Bx = B(1);
By = B(2);%1;%0
Bz = B(3);%1;%0
alphay = 1;
alphaz = 1;
betay = 1;
betaz = 1;
H = 2;
Kxy = K(1);%.1;
Kyz = K(2);%.1;
Kxz = K(3);%.1;

%dx = 0;
Y_pert= polyval(poly_YP2,t);

dx = -x;
dz = betaz*G(x,Kxz,Kyz,Y_pert,H,flags) - alphaz*z;
dm = [dx; dz];

end

function dm = ffl_pert3(t,m,flags,K,B)

x = m(1);
y = m(2);

%Parameter Values
Bx = B(1);
By = B(2);%1;%0
Bz = B(3);%1;%0
alphay = 1;
alphaz = 1;
betay = 1;
betaz = 1;
H = 2;
Kxy = K(1);%.1;
Kyz = K(2);%.1;
Kxz = K(3);%.1;

%dx = 0;
Y_pert= polyval(poly_YP3,t);

dx = -x;
dy = (betay*f(x,Kxy,H,flags(1))) - alphay*y;
dm = [dx; dy];

end

function out = G(u,Ku,Kv,v,H,flags)
    if flags(2) == 1    % AND Gate
        out = f(u,Ku,H,flags(3))*f(v,Kv,H,flags(4));
    else                % OR Gate        
        out = fc(u,Ku,Kv,v,H,flags(3))+fc(v,Kv,Ku,u,H,flags(4));
        
    end
end

function out = f(u,K,H,flag)
    if flag == 1    % Activator
        out = ((u/K)^H)/(1+((u/K)^H));
    else            % Repressor                
        out = (1/(1+((u/K)^H))); 
    end
end


function out = fc(u,Ku,Kv,v,H,flag)
    if flag == 1       % Activator     
        out = ((u/Ku)^H)/(1+((u/Ku)^H)+((v/Kv)^H));       
    else               % Repressor
        out = (1/(1+((u/Ku)^H)+((v/Kv)^H)));   
    end
end

end
```
