## Supplementary Code for "Network Inference from Perturbation Time Course Data": fflwrapper.m

```
function [ffldata] = fflwrapper(count,timeint,tmax,Ks,Bs,pert1,pert2,pert3,multistart)
fmin=cell(16,50);
finSSE=cell(16,50);
for i=1:16
    for j=1:50
    fmin{i,j}=zeros(multistart,10);
    finSSE{i,j}=zeros(multistart,1);
    end
end

y0=zeros(16,3);
yp1=zeros(16,3);
yp2=zeros(16,3);
yp3=zeros(16,3);

data_nostim=feedforwarddatagenerator_nostim(timeint,100,Ks,Bs,pert1,pert2,pert3,0);
% jac=calcjacob(data_nostim,Ks,Bs);
m=1;
for i = 1:2
    for j = 1:2
        for k = 1:2
            for l = 1:2
                key = i*1000+j*100+k*10+l;
                nostimdata=data_nostim(key);              
                y0(m,:)=nostimdata.Y(100,:);
                yp1(m,:)=nostimdata.YP1(100,:);
                yp2(m,:)=nostimdata.YP2(100,:);
                yp3(m,:)=nostimdata.YP3(100,:);
                m=m+1;
            end
        end
    end
end
                
data0=feedforwarddatagenerator_stim(timeint,tmax,y0,yp1,yp2,yp3,Ks,Bs,pert1,pert2,pert3,0);
for h=1:count
data10{h}=feedforwarddatagenerator_stim(timeint,tmax,y0,yp1,yp2,yp3,Ks,Bs,pert1,pert2,pert3,0.1);
data20{h}=feedforwarddatagenerator_stim(timeint,tmax,y0,yp1,yp2,yp3,Ks,Bs,pert1,pert2,pert3,0.2);
data50{h}=feedforwarddatagenerator_stim(timeint,tmax,y0,yp1,yp2,yp3,Ks,Bs,pert1,pert2,pert3,0.5);
end

for h=1:count
    m=1;
for i = 1:2
    for j = 1:2
        for k = 1:2
            for l = 1:2
                key = i*1000+j*100+k*10+l;
                data_10{m,h}=data10{h}(key);
                data_20{m,h}=data20{h}(key);
                data_50{m,h}=data50{h}(key);
                m=m+1;
            end
        end
    end
end
end

            
parfor h=1:count
for m = 1:16
%                 data_10_{m}=data_10{m,h};
%                 data_20_{m}=data_20{m,h};
%                 data_50_{m}=data_50{m,h};

                for n=1:multistart
                [fmin{m,h}(n,:),finSSE{m,h}(n,:)]= estimateffl(data_10{m,h},timeint,tmax,y0(m,:),yp1(m,:),yp2(m,:),yp3(m,:));
                end
                fmin_data10{m,h}=fmin{m,h};
                fmin_data10_SSE{m,h}=finSSE{m,h};
                for n=1:multistart
                [fmin{m,h}(n,:),finSSE{m,h}(n,1)]= estimateffl(data_20{h},timeint,tmax,y0(m,:),yp1(m,:),yp2(m,:),yp3(m,:));
                end
                fmin_data20{m,h}=fmin{m,h};
                fmin_data20_SSE{m,h}=finSSE{m,h};
                for n=1:multistart
                [fmin{m,h}(n,:),finSSE{m,h}(n,1)]= estimateffl(data_50{h},timeint,tmax,y0(m,:),yp1(m,:),yp2(m,:),yp3(m,:));
                end
                fmin_data50{m,h}=fmin{m,h};
                fmin_data50_SSE{m,h}=finSSE{m,h};
                disp(m);
%                 m=m+1;
end
end

datas{1,1}=data0;
datas{2,1}=data10;
datas{3,1}=data20;
datas{4,1}=data50;

ffldata.datas=datas;
% ffldata.preds0=fmin_data0;
ffldata.preds10=fmin_data10;
ffldata.preds20=fmin_data20;
ffldata.preds50=fmin_data50;
% ffldata.preds0_SSE=fmin_data0_SSE;
ffldata.preds10_SSE=fmin_data10_SSE;
ffldata.preds20_SSE=fmin_data20_SSE;
ffldata.preds50_SSE=fmin_data50_SSE;
% end
%     save ('ffldata.mat','ffldata')
end
```
