## Supplementary Code for "Network Inference from Perturbation Time Course Data": fflwrapper_NoNoise.m

```
function [ffldata] = fflwrapper_NoNoise(timeint,tmax,Ks,Bs,pert1,pert2,pert3,multistart)
fmin=cell(1,1);
finSSE=cell(1,1);
for i=1:16
    for j=1:1
    fmin{i,j}=zeros(multistart,10);
    finSSE{i,j}=zeros(multistart,1);
    end
end

y0=zeros(16,3);
yp1=zeros(16,3);
yp2=zeros(16,3);
yp3=zeros(16,3);

data_nostim=feedforwarddatagenerator_nostim(timeint,100,Ks,Bs,pert1,pert2,pert3,0);
% jac=calcjacob(data_nostim,Ks,Bs);
m=1;
for i = 1:2
    for j = 1:2
        for k = 1:2
            for l = 1:2
                key = i*1000+j*100+k*10+l;
                nostimdata=data_nostim(key);              
                y0(m,:)=nostimdata.Y(100,:);
                yp1(m,:)=nostimdata.YP1(100,:);
                yp2(m,:)=nostimdata.YP2(100,:);
                yp3(m,:)=nostimdata.YP3(100,:);
                m=m+1;
            end
        end
    end
end
                

data0=feedforwarddatagenerator_stim(timeint,tmax,y0,yp1,yp2,yp3,Ks,Bs,pert1,pert2,pert3,0);
    m=1;
for i = 1:2
    for j = 1:2
        for k = 1:2
            for l = 1:2
                key = i*1000+j*100+k*10+l;
                data_0{m,1}=data0(key);
                m=m+1;
            end
        end
    end
end
        
for m = 1:16
                for n=1:multistart
                [fmin_data0{m,1}(n,:),fmin_data0_SSE{m,1}(n,1)]= estimateffl(data_0{m,1},timeint,tmax,y0(m,:),yp1(m,:),yp2(m,:),yp3(m,:));
                end
end

datas{1,1}=data0;
ffldata.datas=datas;
ffldata.preds0=fmin_data0;
ffldata.preds0_SSE=fmin_data0_SSE;
% end
%    save ('ffldata_NoNoise_fullpert.mat','ffldata')
end
```
