## Supplementary Code for "Network Inference from Perturbation Time Course Data": full vs half.m

```
%%
tmax=10;
timeint=1;
pert1=0;
pert2=0;
pert3=0;
Bs=dlmread('Rich_Bout.txt');
Ks=dlmread('Rich_Kout_K24.txt');
data_nostim=feedforwarddatagenerator_nostim(timeint,100,Ks,Bs,pert1,pert2,pert3,0);
% jac=calcjacob(data_nostim,Ks,Bs);
m=1;
for i = 1:2
    for j = 1:2
        for k = 1:2
            for l = 1:2
                key = i*1000+j*100+k*10+l;
                nostimdata=data_nostim(key);              
                y0(m,:)=nostimdata.Y(100,:);
                yp1(m,:)=nostimdata.YP1(100,:);
                yp2(m,:)=nostimdata.YP2(100,:);
                yp3(m,:)=nostimdata.YP3(100,:);
                m=m+1;
            end
        end
    end
end


load("ffldata_MRA_pert0_k24_randstart.mat")
for j=1:16
    [M,I] = min(ffldata.preds{j,2});
    bestest.preds0{j,1}(1,:)=ffldata.preds{j,1}(I,:);
end
for j=1:16
best_est.preds0{j,1}(1,1:4)=bestest.preds0{j,1}(1,1:4);
best_est.preds0{j,1}(1,6:8)=bestest.preds0{j,1}(1,5:7);
best_est.preds0{j,1}(1,10:12)=bestest.preds0{j,1}(1,8:10);
best_est.preds0{j,1}(1,5)=-(y0(j,1)*bestest.preds0{j,1}(1,5)+y0(j,2)*bestest.preds0{j,1}(1,6)+y0(j,3)*bestest.preds0{j,1}(1,7));
best_est.preds0{j,1}(1,9)=-(y0(j,1)*bestest.preds0{j,1}(1,8)+y0(j,2)*bestest.preds0{j,1}(1,9)+y0(j,3)*bestest.preds0{j,1}(1,10));;
end


%% makefig
St_half=load('best_est_multistart10.mat');
St_full=load('best_est_full_inhibition.mat');
%St=Allsum;
% Add ground truth values to outmap
outmap_half=addgtvals(St_half);
outmap_full=addgtvals(St_full);

inds_edges=1:12; %including all edges
p=0.5:0.01:1;
% inds=[1111,1112,1121,1122,1211,1212,1221,1222,2111,2112,2121,2122,2211,2212,2221,2222];
% inds=inds(1:length(outmap));
    gt=[];

% Get No Noise Case
n0s_half=[];
n0s_full=[];

for i=1:16
    % No Noise
    q=outmap_half.best_est.preds0{i,1}(1,inds_edges);
    q(q>0)=1;
    q(q<0)=-1;
    q(q==0)=0;
    n0s_half=[n0s_half;q'];
    
    r=outmap_full.best_est.preds0{i,1}(1,inds_edges);
    r(r>0)=1;
    r(r<0)=-1;
    r(r==0)=0;
    n0s_full=[n0s_full;r'];
    
    gt=[gt,outmap_half.best_est.qpred0(i,inds_edges)];

end
c0_half=(n0s_half==gt');
a0_half=sum(reshape(c0_half,[length(inds_edges),16])',2)/length(inds_edges);
c0_full=(n0s_full==gt');
a0_full=sum(reshape(c0_full,[length(inds_edges),16])',2)/length(inds_edges);
e=figure; 
hm=heatmap([a0_full,a0_half]');
caxis([0.8 1])
hm.Colormap=hot;
hm.ColorData=roundn(hm.ColorData,-2);
set(gcf,'position',[0,0,550,80])
hm.YData = ["100% Inhibition","50% Inhibition"];
hm.FontSize=8;
```
