## Supplementary Code for "Network Inference from Perturbation Time Course Data": Make_Figures_5_6.m

```
% This function reproduces all the MATLAB figures in the manuscript. It is
% intendend to be run in sections. The functions needed to generate the 
% simulation data as well as plot the figures are included. Sometimes,
% the code to generate the data for the models are provided but they take 
% a long time to run. Parallelization is recommended. Alternatively, those 
% sections can be skipped for pre-run mat files which are also provided. 
% Also smaller versions of the data are provided as well which can be run.

%% Generate data for Figure 5&6 (Non Linear Models)
% Run this section to generate Non Linear Model Data and Estimates.
%
% Already generated data is also available in the next section. Skip this 
% section to make figures from saved data
tmax=10;
timeint=1;
multistart=10;
Ks=dlmread('Rich_Kout_K25.txt');
Bs=dlmread('Rich_Bout.txt');

pert1=0;
pert2=0;
pert3=0;
ffldata=fflwrapper_NoNoise(timeint,tmax,Ks,Bs,pert1,pert2,pert3,multistart);
save ('ffldata_NoNoise_fullpert.mat','ffldata')

pert1=0.5;
pert2=0.5;
pert3=0.5;
ffldata=fflwrapper_NoNoise(timeint,tmax,Ks,Bs,pert1,pert2,pert3,multistart);
save ('ffldata_NoNoise_halfpert.mat','ffldata')

count=50;
ffldata=fflwrapper(count,timeint,tmax,Ks,Bs,pert1,pert2,pert3,multistart);
save ('ffldata_Noise_halfpert.mat','ffldata')
%% Plot 5B
clear;
St_full=load("ffldata_noNoise_fullpert.mat");
for j=1:16
    [M,I]=min(St_full.ffldata.preds0_SSE{j,1});
    best_est.preds0(j,1:12)=St_full.ffldata.preds0{j,1}(I,1:12);  
end
outmap=addgtvals(St_full);
% Add ground truth values to outmap
gt=reshape(outmap.best_est.qpred0(1,1:12),[4,3])';
p1=reshape(best_est.preds0(1,1:12)./abs(best_est.preds0(1,1:12)),[4,3])';
gt(isnan(gt))=0;
p1(isnan(p1))=0;
    
b=figure;
subplot(2,1,1);
h=heatmap(gt,'CellLabelColor','none'); colorbar off; %h.XDisplayLabels={'0','1','2','3'}; h.Colormap=redgreencmap;
h.Colormap=redgreencmap;
Ax = gca;
Ax.XDisplayLabels = nan(size(Ax.XDisplayData));
Ax.YDisplayLabels = nan(size(Ax.YDisplayData));

% title('Ground Truth')
subplot(2,1,2);
h=heatmap(p1,'CellLabelColor','none'); colorbar off; %h.XDisplayLabels={'0','1','2','3'}; h.Colormap=redgreencmap;
h.Colormap=redgreencmap;
Ax = gca;
Ax.XDisplayLabels = nan(size(Ax.XDisplayData));
Ax.YDisplayLabels = nan(size(Ax.YDisplayData));

res = 300;
set(gcf,'paperunits','inches','PaperPosition',[0 0 1.2 1.5]);
print('5B.tiff','-dtiff',['-r' num2str(res)]);


%% Figure 5D
                load('ffldata_NoNoise_fullpert.mat')
                ans=ffldata.datas{1,1}(1111);
                subplot(1,4,1);
                hold on
                plot(ans.T,ans.Y(:,1),'b.--', 'MarkerSize', 15);
                plot(ans.T,ans.Y(:,2),'r.--', 'MarkerSize', 15);
                plot(ans.T,ans.Y(:,3),'y.--', 'MarkerSize', 15);

%                 title("Vehicle",'FontSize',10);
                ylim([0 1])

                subplot(1,4,2);
                hold on
                plot(ans.T,ans.YP1(:,1),'b.--', 'MarkerSize', 15);
                plot(ans.T,ans.YP1(:,2),'r.--', 'MarkerSize', 15);
                plot(ans.T,ans.YP1(:,3),'y.--', 'MarkerSize', 15);
%                 title("Perturb x_1", 'FontSize',10);
                ylim([0 1])
                
                subplot(1,4,3);
                hold on
                plot(ans.T,ans.YP2(:,1),'b.--', 'MarkerSize', 15);
                plot(ans.T,ans.YP2(:,2),'r.--', 'MarkerSize', 15);
                plot(ans.T,ans.YP2(:,3),'y.--', 'MarkerSize', 15);
%                 title("Perturb x_2", 'FontSize',10);
                ylim([0 1])
                
                subplot(1,4,4);
                hold on
                plot(ans.T,ans.YP3(:,1),'b.--', 'MarkerSize', 15);
                plot(ans.T,ans.YP3(:,2),'r.--', 'MarkerSize', 15);
                plot(ans.T,ans.YP3(:,3),'y.--', 'MarkerSize', 15);
%                 title("Perturb x_3", 'FontSize',10);
                ylim([0 1])

                res = 300;
                set(gcf,'paperunits','inches','PaperPosition',[0 0 3.5 1.5]);
                print('5D.tiff','-dtiff',['-r' num2str(res)]);

%% Figure 5E
                load('ffldata_NoNoise_halfpert.mat')
                ans=ffldata.datas{1,1}(1111);
                subplot(1,4,1);
                hold on
                plot(ans.T,ans.Y(:,1),'b.--', 'MarkerSize', 15);
                plot(ans.T,ans.Y(:,2),'r.--', 'MarkerSize', 15);
                plot(ans.T,ans.Y(:,3),'y.--', 'MarkerSize', 15);

%                 title("Vehicle",'FontSize',10);
                ylim([0 1])

                subplot(1,4,2);
                hold on
                plot(ans.T,ans.YP1(:,1),'b.--', 'MarkerSize', 15);
                plot(ans.T,ans.YP1(:,2),'r.--', 'MarkerSize', 15);
                plot(ans.T,ans.YP1(:,3),'y.--', 'MarkerSize', 15);
%                 title("Perturb x_1", 'FontSize',10);
                ylim([0 1])
                
                subplot(1,4,3);
                hold on
                plot(ans.T,ans.YP2(:,1),'b.--', 'MarkerSize', 15);
                plot(ans.T,ans.YP2(:,2),'r.--', 'MarkerSize', 15);
                plot(ans.T,ans.YP2(:,3),'y.--', 'MarkerSize', 15);
%                 title("Perturb x_2", 'FontSize',10);
                ylim([0 1])
                
                subplot(1,4,4);
                hold on
                plot(ans.T,ans.YP3(:,1),'b.--', 'MarkerSize', 15);
                plot(ans.T,ans.YP3(:,2),'r.--', 'MarkerSize', 15);
                plot(ans.T,ans.YP3(:,3),'y.--', 'MarkerSize', 15);
%                 title("Perturb x_3", 'FontSize',10);
                ylim([0 1])

                res = 300;
                set(gcf,'paperunits','inches','PaperPosition',[0 0 3.5 1.5]);
                print('5E.tiff','-dtiff',['-r' num2str(res)]);

%% %% Figure 5F
  % Percentile Method
St=load('ffldata_NoNoise_fullpert.mat');
% Add ground truth values to outmap
outmap=addgtvals(St);
inds_edges=1:12; %including all edges
p=0.5:0.01:1;

% Get No Noise Case
n_full=[];
for i=1:16
    % No Noise
    [M,I] = min(outmap.ffldata.preds0_SSE{i,1});
    q=outmap.ffldata.preds0{i,1}(I,inds_edges);
    q(q>0)=1;
    q(q<0)=-1;
    q(q==0)=0;
    n_full=[n_full;q'];
end

St=load('ffldata_NoNoise_halfpert.mat');
% Add ground truth values to outmap
outmap=addgtvals(St);
inds_edges=1:12; %including all edges
p=0.5:0.01:1;
gt=[];
% Get No Noise Case
n_half=[];
for i=1:16
    % No Noise
    [M,I] = min(outmap.ffldata.preds0_SSE{i,1});
    q=outmap.ffldata.preds0{i,1}(I,inds_edges);
    q(q>0)=1;
    q(q<0)=-1;
    q(q==0)=0;
    n_half=[n_half;q'];
    gt=[gt,outmap.best_est.qpred0(i,inds_edges)];
end

c_full=(n_full==gt');
a_full=sum(reshape(c_full,[length(inds_edges),16])',2)/length(inds_edges);

c_half=(n_half==gt');
a_half=sum(reshape(c_half,[length(inds_edges),16])',2)/length(inds_edges);

b_full=sum(reshape(c_full,[length(inds_edges),16]),2)/16;
b_half=sum(reshape(c_half,[length(inds_edges),16]),2)/16;

% e=figure; 
hm=heatmap([a_full,a_half]');
caxis([0 1])
hm.Colormap=hot;
hm.ColorData=roundn(hm.ColorData,-2);
set(gcf,'position',[0,0,600,200])
hm.YData = ["100% Inhibition" "50% Inhibition"];
res = 300;
set(gcf,'paperunits','inches','PaperPosition',[0 0 3.5 3]);
print('5F.tiff','-dtiff',['-r' num2str(res)]);
%% Make Values to plot in 6a
for i=1:50
Values(i,1)=1+normrnd(0,1,1,1);
Values(i,2)=0+normrnd(0,1,1,1);
Values(i,3)=-1+normrnd(0,1,1,1);
end
save("Values.mat","Values")
%% Figure 6a
% Save Figures as tiff
clear;
load("Values.mat")
% h1=boxplot(Values,'Notch','on','Labels',{'Node Edge 1','Node Edge 2','Node Edge 3'},'Whisker',10);
lines = findobj(gcf, 'type', 'line', 'Tag', 'Median');
set(lines, 'Color', 'g');
q = quantile(Values(:,1),[0.5 0.4 0.6 0.1 0.9 ]);
qa50 = q(1);  qa40 = q(2);  qa60 = q(3); qa10 = q(4); qa90 = q(5);  
q = quantile(Values(:,2),[0.5 0.4 0.6 0.1 0.9 ]);
qb50 = q(1);  qb40 = q(2);  qb60 = q(3); qb10 = q(4); qb90 = q(5);  
q = quantile(Values(:,3),[0.5 0.4 0.6 0.1 0.9 ]);
qc50 = q(1);  qc40 = q(2);  qc60 = q(3); qc10 = q(4); qc90 = q(5);  
res=300;
x=ones(length(Values)).*(1+(rand(length(Values))-0.5)/5);
x1=ones(length(Values)).*(1+(rand(length(Values))-0.5)/10);
x2=ones(length(Values)).*(1+(rand(length(Values))-0.5)/15);
f1=scatter(x(:,1),Values(:,1),'k','filled');f1.MarkerFaceAlpha = 0.4;hold on 
f2=scatter(x1(:,2).*2,Values(:,2),'k','filled');f2.MarkerFaceAlpha = f1.MarkerFaceAlpha;hold on
f3=scatter(x2(:,3).*3,Values(:,3),'k','filled');f3.MarkerFaceAlpha = f1.MarkerFaceAlpha;hold on
line(xlim(), [0,0], 'LineWidth', 1, 'Color', 'b');

h1=boxplot(Values,'Labels',{'Edge 1','Edge 2','Edge 3'},'Whisker',10);
set(h1,{'linew'},{2})
% xlabel('Representative Node Edges','FontSize',10)
% ylabel('Estimated Node Edges','FontSize',10)
title({"Classification of Node Edges"; "at 50th percentile (median)"},'FontSize',10)

set(h1(5,1), 'YData', [qa50 qa50 qa50 qa50]);% blue box  
upWhisker = get(h1(1,1), 'YData');  
set(h1(1,1), 'YData', [qa50 upWhisker(2)])  
dwWhisker = get(h1(2,1), 'YData');  
set(h1(2,1), 'YData', [dwWhisker(1) qa50])

set(h1(5,2), 'YData', [qb50 qb50 qb50 qb50 qb50]);% blue box  
upWhisker = get(h1(1,2), 'YData');  
set(h1(1,2), 'YData', [qb50 upWhisker(2)])  
dwWhisker = get(h1(2,2), 'YData');  
set(h1(2,2), 'YData', [ dwWhisker(1) qb50])

set(h1(5,3), 'YData', [qc50 qc50 qc50 qc50 qc50]);% blue box  
upWhisker = get(h1(1,3), 'YData');  
set(h1(1,3), 'YData', [qc50 upWhisker(2)])  
dwWhisker = get(h1(2,3), 'YData');  
set(h1(2,3), 'YData', [ dwWhisker(1) qc50])
set(gcf,'paperunits','inches','paperposition',[0 0 2.3 2.5]);
print('6A1.tiff','-dtiff',['-r' num2str(res)]);

figure;
f1=scatter(x(:,1),Values(:,1),'k','filled');f1.MarkerFaceAlpha = 0.4;hold on 
f2=scatter(x1(:,2).*2,Values(:,2),'k','filled');f2.MarkerFaceAlpha = f1.MarkerFaceAlpha;hold on
f3=scatter(x2(:,3).*3,Values(:,3),'k','filled');f3.MarkerFaceAlpha = f1.MarkerFaceAlpha;hold on
line(xlim(), [0,0], 'LineWidth', 0.5, 'Color', 'b');

h2=boxplot(Values,'Labels',{'Edge 1','Edge 2','Edge 3'},'Whisker',10);
set(h2,{'linew'},{2})
title({"Classification of Node Edges"; "between 40th-60th percentile"},'FontSize',10)

set(h2(5,1), 'YData', [qa40 qa60 qa60 qa40 qa40]);% blue box  
upWhisker = get(h2(1,1), 'YData');  
set(h2(1,1), 'YData', [qa60 upWhisker(2)])  
dwWhisker = get(h2(2,1), 'YData');  
set(h2(2,1), 'YData', [ dwWhisker(1) qa40])

set(h2(5,2), 'YData', [qb40 qb60 qb60 qb40 qb40]);% blue box  
upWhisker = get(h2(1,2), 'YData');  
set(h2(1,2), 'YData', [qb60 upWhisker(2)])  
dwWhisker = get(h2(2,2), 'YData');  
set(h2(2,2), 'YData', [ dwWhisker(1) qb40])

set(h2(5,3), 'YData', [qc40 qc60 qc60 qc40 qc40]);% blue box  
upWhisker = get(h2(1,3), 'YData');  
set(h2(1,3), 'YData', [qc60 upWhisker(2)])  
dwWhisker = get(h2(2,3), 'YData');  
set(h2(2,3), 'YData', [ dwWhisker(1) qc40])
set(gcf,'paperunits','inches','paperposition',[0 0 2.3 2.5]);
print('6A2.tiff','-dtiff',['-r' num2str(res)]);

figure;
f1=scatter(x(:,1),Values(:,1),'k','filled');f1.MarkerFaceAlpha = 0.4;hold on 
f2=scatter(x1(:,2).*2,Values(:,2),'k','filled');f2.MarkerFaceAlpha = f1.MarkerFaceAlpha;hold on
f3=scatter(x2(:,3).*3,Values(:,3),'k','filled');f3.MarkerFaceAlpha = f1.MarkerFaceAlpha;hold on
line(xlim(), [0,0], 'LineWidth', 1, 'Color', 'b');

h3=boxplot(Values,'Labels',{'Edge 1','Edge 2','Edge 3'},'Whisker',10);
set(h3,{'linew'},{2})
% xlabel('Representative Node Edges','FontSize',10)
% ylabel('Estimated value of Node Edges','FontSize',10)
title({"Classification of Node Edges"; "between 10th-90th percentile"},'FontSize',10)

set(h3(5,1), 'YData', [qa10 qa90 qa90 qa10 qa10]);% blue box  
upWhisker = get(h3(1,1), 'YData');  
set(h3(1,1), 'YData', [qa90 upWhisker(2)])  
dwWhisker = get(h3(2,1), 'YData');  
set(h3(2,1), 'YData', [ dwWhisker(1) qa10])

set(h3(5,2), 'YData', [qb10 qb90 qb90 qb10 qb10]);% blue box  
upWhisker = get(h3(1,2), 'YData');  
set(h3(1,2), 'YData', [qb90 upWhisker(2)])  
dwWhisker = get(h3(2,2), 'YData');  
set(h3(2,2), 'YData', [ dwWhisker(1) qb10])

set(h3(5,3), 'YData', [qc10 qc90 qc90 qc10 qc10]);% blue box  
upWhisker = get(h3(1,3), 'YData');  
set(h3(1,3), 'YData', [qc90 upWhisker(2)])  
dwWhisker = get(h3(2,3), 'YData');  
set(h3(2,3), 'YData', [ dwWhisker(1) qc10])

set(gcf,'paperunits','inches','paperposition',[0 0 2.3 2.5]);
print('6A3.tiff','-dtiff',['-r' num2str(res)]);

%% Select best estimates from multistart
clear all;
load("ffldata_Noise_halfpert.mat")
for j=1:16
    for i=1:50
    [M,I]=min(ffldata.preds10_SSE{j,i});
    best_est.preds10{j,1}(i,1:12)=ffldata.preds10{j,i}(I,1:12);
    
    [M,I]=min(ffldata.preds20_SSE{1,i});
    best_est.preds20{j,1}(i,1:12)=ffldata.preds20{j,i}(I,1:12);
    
    [M,I]=min(ffldata.preds50_SSE{1,i});
    best_est.preds50{j,1}(i,1:12)=ffldata.preds50{j,i}(I,1:12);
end
end
load("ffldata_noNoise_halfpert.mat")
for j=1:16
    [M,I]=min(ffldata.preds0_SSE{j,1});
    best_est.preds0{j,1}(1,1:12)=ffldata.preds0{j,1}(I,1:12);  
end
save ('best_est.mat','best_est')

%% Figure 6(b,c,d) S7
  % Percentile Method
St=load('best_est.mat');
% Add ground truth values to outmap
outmap=addgtvals(St);
inds_edges=1:12; %including all edges
p=0.5:0.01:1;

% Get No Noise Case
n0s=[];
for i=1:16
    % No Noise
    q=outmap.best_est.preds0{i,1}(1,inds_edges);
    q(q>0)=1;
    q(q<0)=-1;
    q(q==0)=0;
    n0s=[n0s;q'];
end

% Get Noise Cases, with percentile aproach
for k=1:length(p)
    n1=[];
    n2=[];
    n5=[];
    gt=[];
    
    for i=1:16
       
        % 1:10
        q1u=quantile(outmap.best_est.preds10{i,1}(:,inds_edges),p(k));
        q1d=quantile(outmap.best_est.preds10{i,1}(:,inds_edges),1-p(k));
        mat1=zeros(1,length(inds_edges));
        lp=(double(q1u>0) .* double(q1d>0)); mat1(find(lp))=1;
        ln=(double(q1u<0) .* double(q1d<0)); mat1(find(ln))=-1;
        ll=(double(q1u>0) .* double(q1d<0)); mat1(find(ll))=0;
        n1=[n1,mat1];
        
        % 1:5
        q2u=quantile(outmap.best_est.preds20{i,1}(:,inds_edges),p(k));
        q2d=quantile(outmap.best_est.preds20{i,1}(:,inds_edges),1-p(k));
        mat2=zeros(1,length(inds_edges));
        lp=(double(q2u>0) .* double(q2d>0)); mat2(find(lp))=1;
        ln=(double(q2u<0) .* double(q2d<0)); mat2(find(ln))=-1;
        ll=(double(q2u>0) .* double(q2d<0)); mat2(find(ll))=0;
        n2=[n2,mat2];
        
        % 1:2
        q5u=quantile(outmap.best_est.preds50{i,1}(:,inds_edges),p(k));
        q5d=quantile(outmap.best_est.preds50{i,1}(:,inds_edges),1-p(k));
        mat5=zeros(1,length(inds_edges));
        lp=(double(q5u>0) .* double(q5d>0)); mat5(find(lp))=1;
        ln=(double(q5u<0) .* double(q5d<0)); mat5(find(ln))=-1;
        ll=(double(q5u>0) .* double(q5d<0)); mat5(find(ll))=0;
        n5=[n5,mat5];
        
        gt=[gt,outmap.best_est.qpred0(i,inds_edges)];
        
    end
    
    n1s(:,k)=n1';
    n2s(:,k)=n2';
    n5s(:,k)=n5';
    gts=gt;
    
    % Regulation vs no regulation.
    n1t=abs(n1);
    n2t=abs(n2);
    n5t=abs(n5);
    gtt=abs(gt);
    
    TPR1(k)=sum((double(n1t==1) .* double(gtt==1)))/sum(gtt);
    TPR2(k)=sum((double(n2t==1) .* double(gtt==1)))/sum(gtt);
    TPR5(k)=sum((double(n5t==1) .* double(gtt==1)))/sum(gtt);
    
    FPR1(k)=sum((double(n1t==1) .* double(gtt==0)))/sum(gtt==0);
    FPR2(k)=sum((double(n2t==1) .* double(gtt==0)))/sum(gtt==0);
    FPR5(k)=sum((double(n5t==1) .* double(gtt==0)))/sum(gtt==0);
    
end

% Plot ROC curves
d=figure;
plot([1,FPR1,0],[1,TPR1,0],'g','LineWidth',1)
disp(['AUC1=',num2str(trapz([1,FPR1,0]',[1,TPR1,0]'))])
hold on

plot([1,FPR2,0],[1,TPR2,0],'b--','LineWidth',2)
disp(['AUC2=',num2str(trapz(flipud([1,FPR2,0]'),flipud([1,TPR2,0]')))])

plot([1,FPR5,0],[1,TPR5,0],'r','LineWidth',2)
disp(['AUC5=',num2str(trapz(flipud([1,FPR5,0]'),flipud([1,TPR5,0]')))])
xlabel('False Positive Rate', 'FontSize',10)
ylabel('True Positive Rate', 'FontSize',10)
legend('10:1 Signal:Noise','5:1 Signal:Noise','2:1 Signal:Noise','Location','southeast','FontSize',9)
size = [2.25 2.25];
res = 300;
set(gcf,'paperunits','inches','paperposition',[0 0 size]);
print('6B.tiff','-dtiff',['-r' num2str(res)]);

ind1=min(find(FPR1<0.05));
ind2=min(find(FPR2<0.05));
ind5=min(find(FPR5<0.05));

c0=(n0s==gt');
a0=sum(reshape(c0,[length(inds_edges),16])',2)/length(inds_edges);

c1=(n1s(:,ind1)==gt');
a1=sum(reshape(c1,[length(inds_edges),16])',2)/length(inds_edges);

c2=(n2s(:,ind2)==gt');
a2=sum(reshape(c2,[length(inds_edges),16])',2)/length(inds_edges);

c5=(n5s(:,ind5)==gt');
a5=sum(reshape(c5,[length(inds_edges),16])',2)/length(inds_edges);

b0=sum(reshape(c0,[length(inds_edges),16]),2)/16;
b1=sum(reshape(c1,[length(inds_edges),16]),2)/16;
b2=sum(reshape(c2,[length(inds_edges),16]),2)/16;
b5=sum(reshape(c5,[length(inds_edges),16]),2)/16;

e=figure; 
hm=heatmap([a0,a1,a2,a5]','FontSize',9);
caxis([0.5 1])
hm.Colormap=hot;
hm.ColorData=roundn(hm.ColorData,-2);
hm.YData = ["No Noise" "10:1 S:N" "5:1 S:N" "2:1 S:N"];
size = [5.75 2.25];
res = 300;
set(gcf,'paperunits','inches','paperposition',[0 0 size]);
print('6C.tiff','-dtiff',['-r' num2str(res)]);

figure;
h2=heatmap([b0,b1,b2,b5]','FontSize',8);
caxis([0 1])
h2.Colormap=hot;
h2.ColorData=roundn(h2.ColorData,-2);
h2.YData = ["No Noise" "10:1 S:N" "5:1 S:N" "2:1 S:N"];
h2.XData = ["S_1_,_e_x" "F_1_1" "F_1_2" "F_1_3" "S_2_,_b" "F_2_1" "F_2_2" "F_2_3" "S_3_,_b" "F_3_1" "F_3_2" "F_3_3"];
size = [5.75 2.25];
res = 300;
set(gcf,'paperunits','inches','paperposition',[0 0 size]);
print('6D.tiff','-dtiff',['-r' num2str(res)]);

% Figure S7
s1=figure;
subplot(2,2,1); h_1=heatmap(double(reshape(c0,[length(inds_edges),16]))); caxis([0.5 1]); colorbar off; title('No Noise');
h_1.YData = ["S_1_,_e_x" "F_1_1" "F_1_2" "F_1_3" "S_2_,_b" "F_2_1" "F_2_2" "F_2_3" "S_3_,_b" "F_3_1" "F_3_2" "F_3_3"];
subplot(2,2,2); h_2=heatmap(double(reshape(c1,[length(inds_edges),16]))); caxis([0.5 1]); colorbar off; title('10:1 Signal:Noise');
h_2.YData = ["S_1_,_e_x" "F_1_1" "F_1_2" "F_1_3" "S_2_,_b" "F_2_1" "F_2_2" "F_2_3" "S_3_,_b" "F_3_1" "F_3_2" "F_3_3"];
subplot(2,2,3); h_3=heatmap(double(reshape(c2,[length(inds_edges),16]))); caxis([0.5 1]); colorbar off; title('5:1 Signal:Noise');
h_3.YData = ["S_1_,_e_x" "F_1_1" "F_1_2" "F_1_3" "S_2_,_b" "F_2_1" "F_2_2" "F_2_3" "S_3_,_b" "F_3_1" "F_3_2" "F_3_3"];
subplot(2,2,4); h_4=heatmap(double(reshape(c5,[length(inds_edges),16]))); caxis([0.5 1]); colorbar off; title('2:1 Signal:Noise');
h_4.YData = ["S_1_,_e_x" "F_1_1" "F_1_2" "F_1_3" "S_2_,_b" "F_2_1" "F_2_2" "F_2_3" "S_3_,_b" "F_3_1" "F_3_2" "F_3_3"];

% set(gcf,'position',[0,0,600,600])
size = [6.75 8.75];
res = 300;
set(gcf,'paperunits','inches','paperposition',[0 0 size]);
print('S7.tiff','-dtiff',['-r' num2str(res)]);

%% Supplement 6: Data vs fit
load('ffldata_NoNoise_halfpert.mat');
load('best_est.mat');
param=zeros(1,15);
m=1;
for i = 1:2
    for j = 1:2
        for k = 1:2
            for l = 1:2
                key = i*1000+j*100+k*10+l;
                data_a=ffldata.datas{1,1}(key);
                q=best_est.preds0{m,1}(1,1:12);
                param=[0,q(1),q(2),q(3),q(4),q(5),0,q(6),q(7),q(8),q(9),0,q(10),q(11),q(12)];
                data_e=eval_data_3node_polyval(1,10,param,data_a.Y(1,:),data_a.YP1(1,:),data_a.YP2(1,:),data_a.YP3(1,:),0.5,0.5,0.5,0);

                subplot(16,4,4*m-3);
                hold on
                plot(data_e.T,data_e.Y(:,1),'b-', 'LineWidth', 1);
                plot(data_e.T,data_e.Y(:,2),'r-', 'LineWidth', 1);
                plot(data_e.T,data_e.Y(:,3),'y-', 'LineWidth', 1);
                plot(data_a.T,data_a.Y(:,1),'b.', 'MarkerSize', 10);
                plot(data_a.T,data_a.Y(:,2),'r.', 'MarkerSize', 10);
                plot(data_a.T,data_a.Y(:,3),'y.', 'MarkerSize', 10);
%                 title("Vehicle",'FontSize',10);
%                 ylim([0 1])

                subplot(16,4,4*m-2);
                hold on
                plot(data_e.T,data_e.YP1(:,1),'b-', 'LineWidth', 1);
                plot(data_e.T,data_e.YP1(:,2),'r-', 'LineWidth', 1);
                plot(data_e.T,data_e.YP1(:,3),'y-', 'LineWidth', 1);
                plot(data_a.T,data_a.YP1(:,1),'b.', 'MarkerSize', 10);
                plot(data_a.T,data_a.YP1(:,2),'r.', 'MarkerSize', 10);
                plot(data_a.T,data_a.YP1(:,3),'y.', 'MarkerSize', 10);
%                 title("Perturb x_1", 'FontSize',10);
%                 ylim([0 1])
                
                subplot(16,4,4*m-1);
                hold on
                plot(data_e.T,data_e.YP2(:,1),'b-', 'LineWidth', 1);
                plot(data_e.T,data_e.YP2(:,2),'r-', 'LineWidth', 1);
                plot(data_e.T,data_e.YP2(:,3),'y-', 'LineWidth', 1);
                plot(data_a.T,data_a.YP2(:,1),'b.', 'MarkerSize', 10);
                plot(data_a.T,data_a.YP2(:,2),'r.', 'MarkerSize', 10);
                plot(data_a.T,data_a.YP2(:,3),'y.', 'MarkerSize', 10);
                %                 title("Perturb x_2", 'FontSize',10);
%                 ylim([0 1])
                
                subplot(16,4,4*m);
                hold on
                plot(data_e.T,data_e.YP3(:,1),'b-', 'LineWidth', 1);
                plot(data_e.T,data_e.YP3(:,2),'r-', 'LineWidth', 1);
                plot(data_e.T,data_e.YP3(:,3),'y-', 'LineWidth', 1);
                plot(data_a.T,data_a.YP3(:,1),'b.', 'MarkerSize', 10);
                plot(data_a.T,data_a.YP3(:,2),'r.', 'MarkerSize', 10);
                plot(data_a.T,data_a.YP3(:,3),'y.', 'MarkerSize', 10);
                %                 title("Perturb x_3", 'FontSize',10);
%                 ylim([0 1])
                m=m+1;
            end
        end
    end
end

                res = 300;
                set(gcf,'paperunits','inches','PaperPosition',[0 0 6.5 12.5]);
                print('S6.tiff','-dtiff',['-r' num2str(res)]);
                
```
