## Supplementary figures and images for "Network Inference from Perturbation Time Course Data"

### 1C.tiff

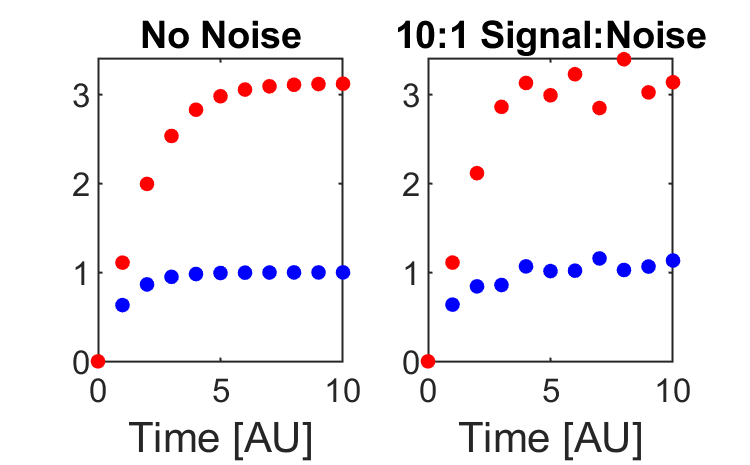

### 1E.tiff

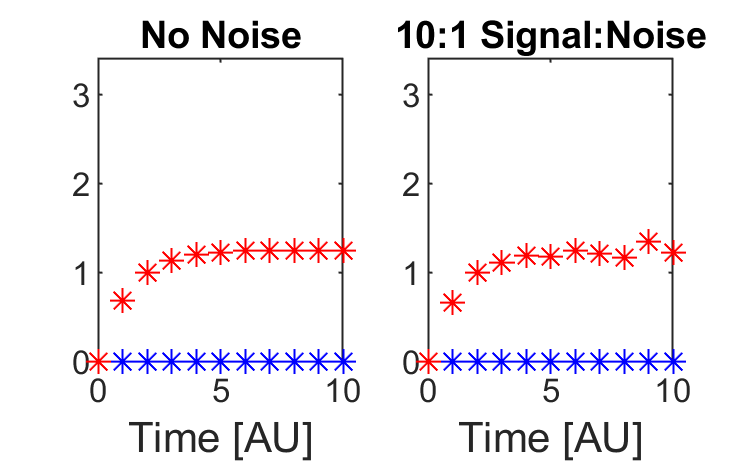

### 1G.tiff

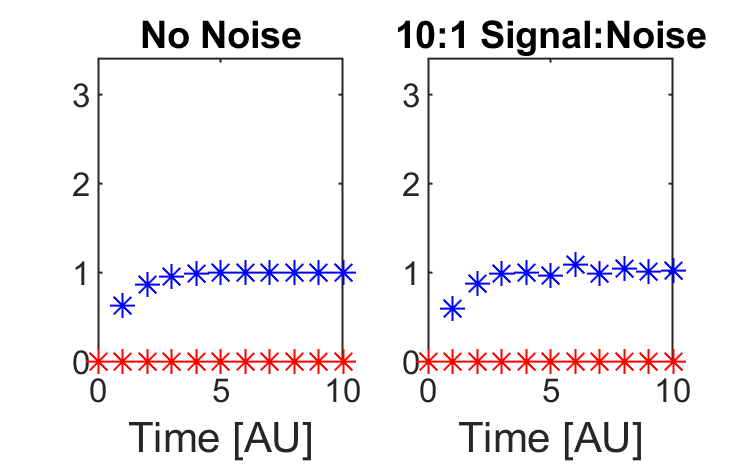

### 1H.tiff

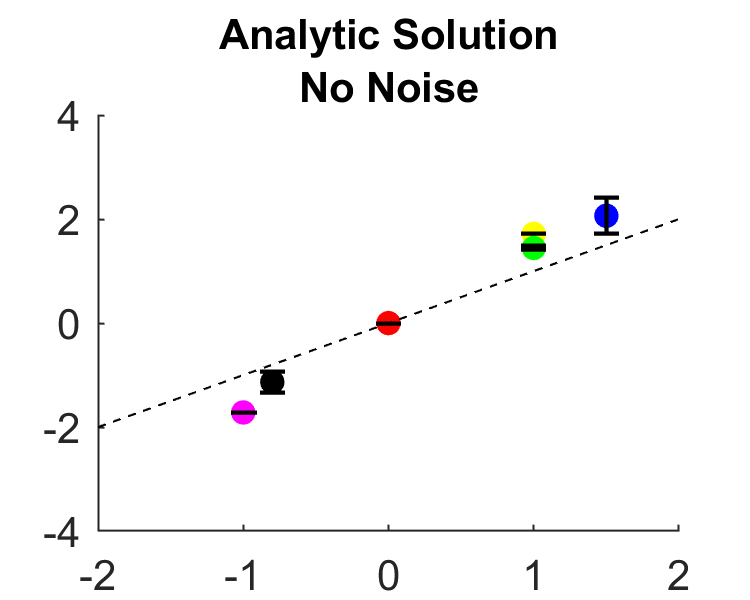

### 1I.tiff

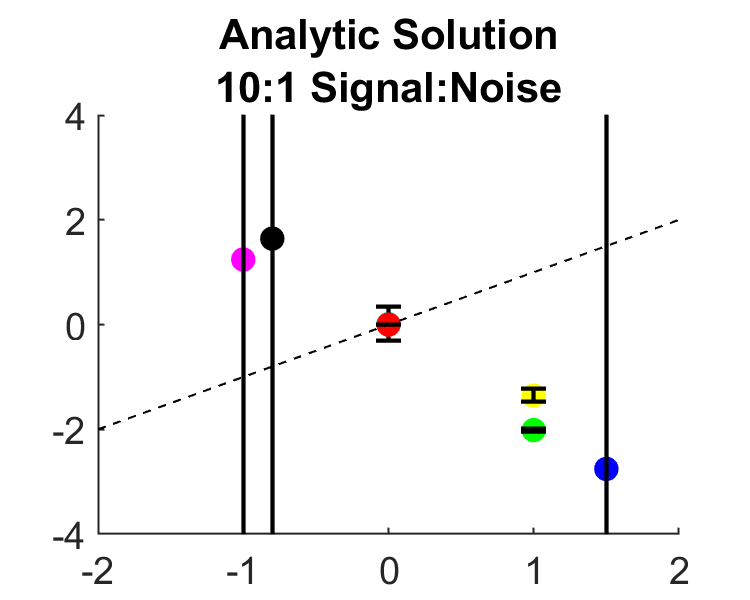

### 1I_inset.tiff

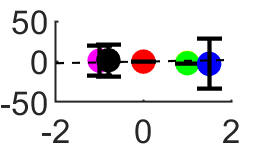

### 1J.tif

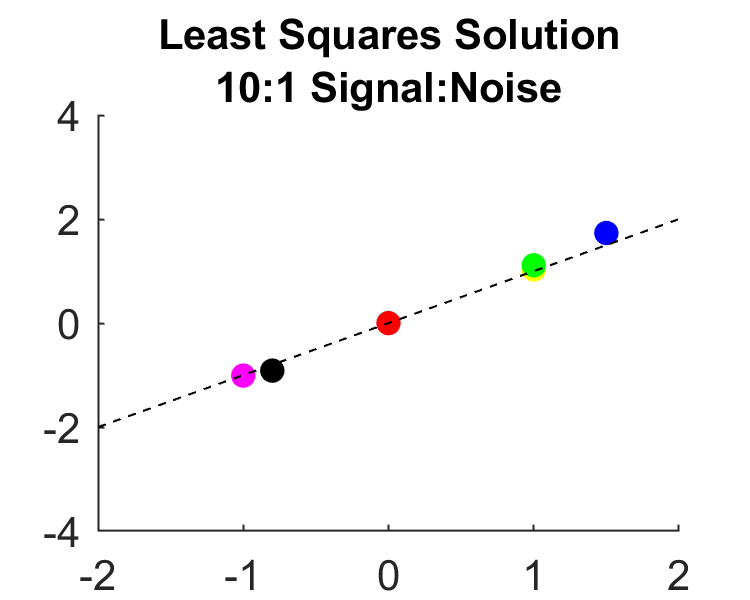

### 2C.tiff

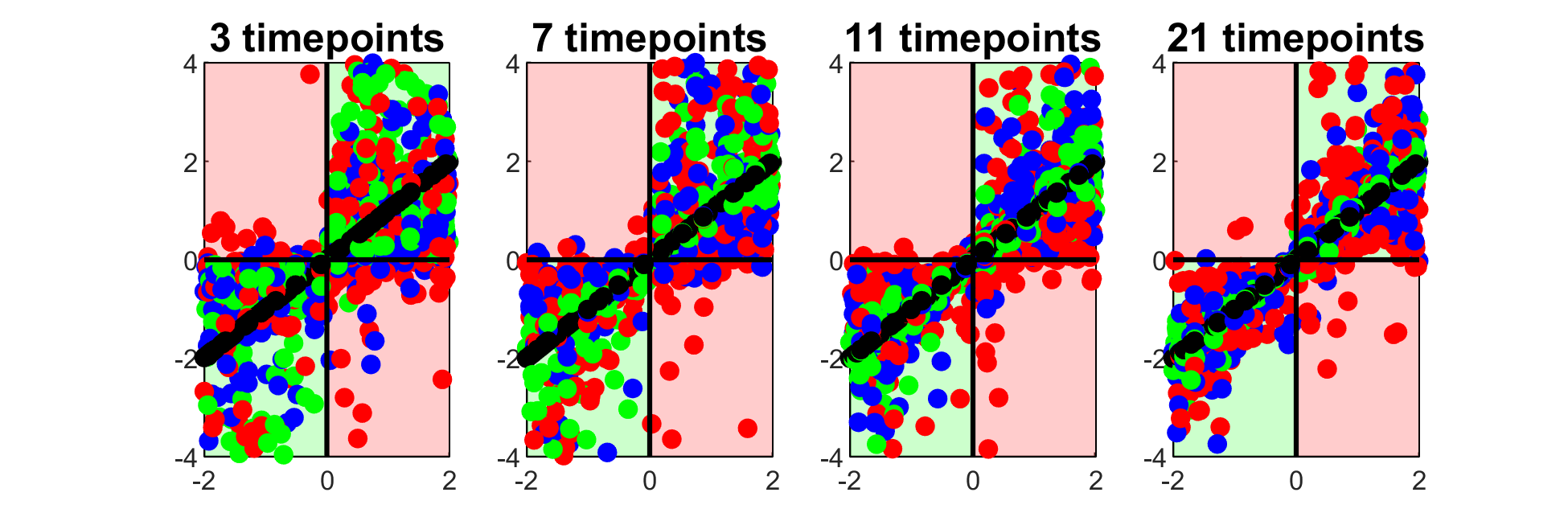

### 2D.tiff

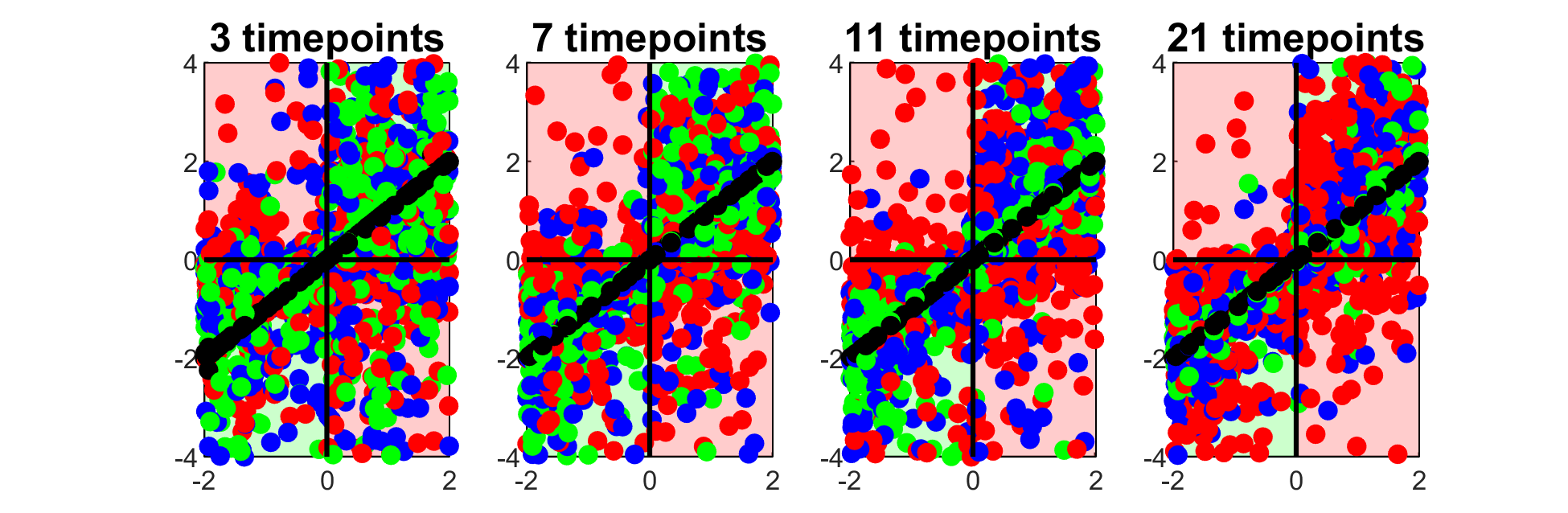

### 2D.tiff

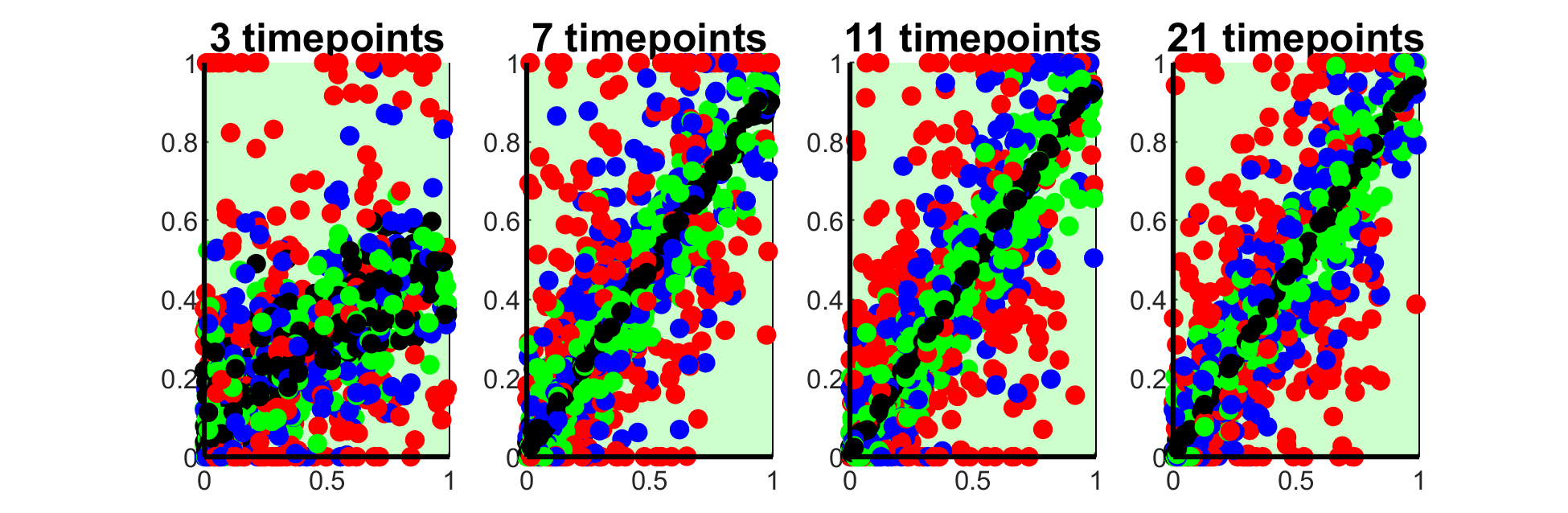

### 2E.tiff

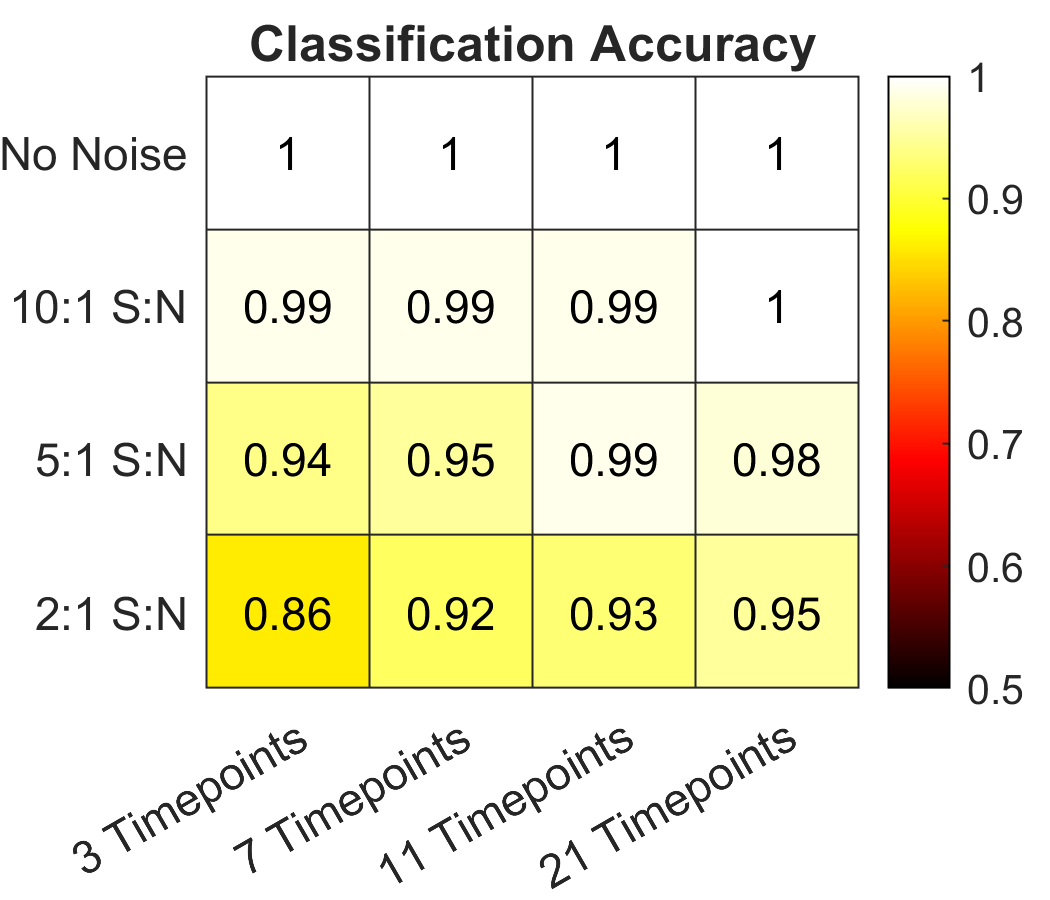

### 2F.tiff

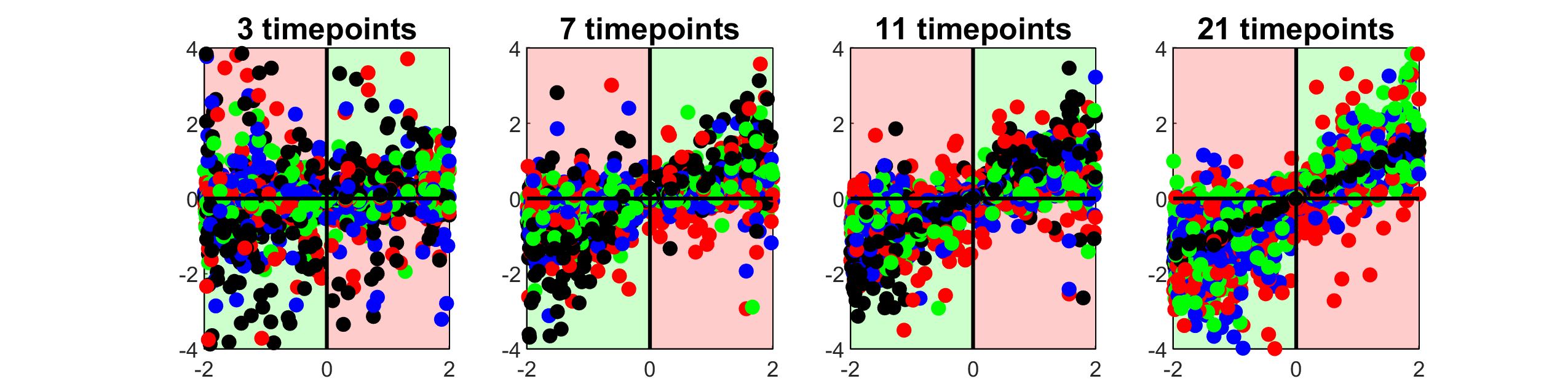

### 2F.tiff

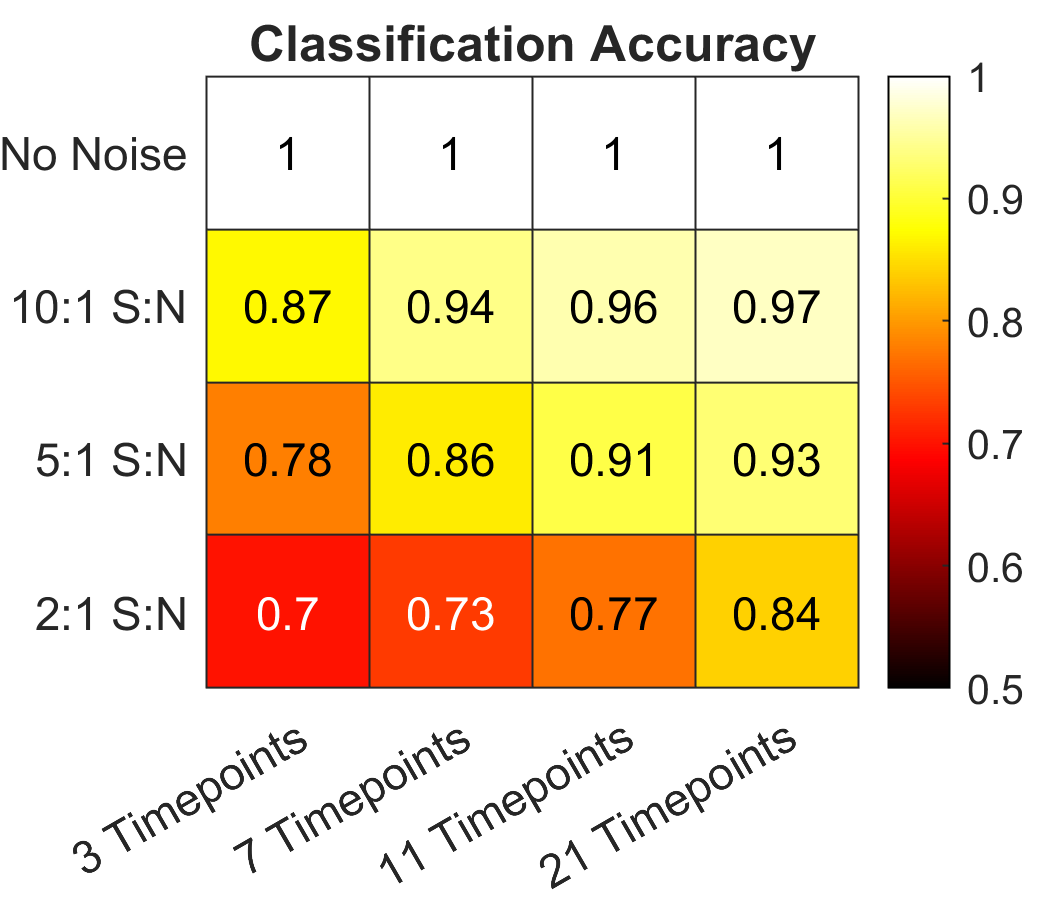

### 3B.tiff

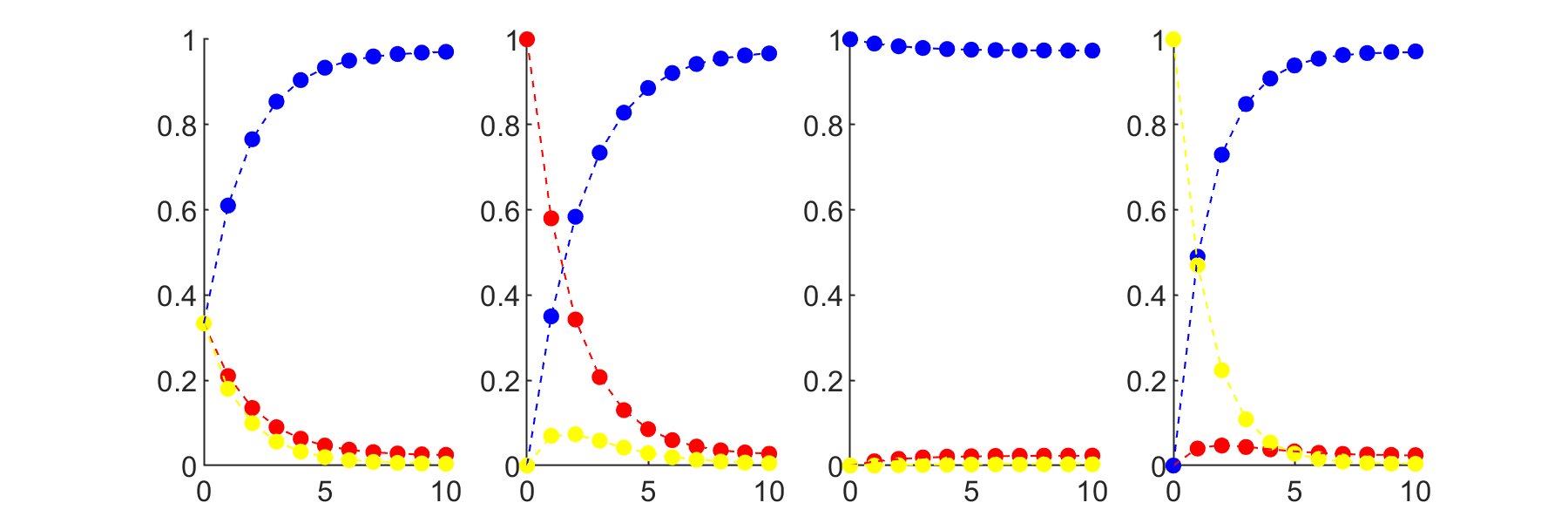

### 3C.tiff

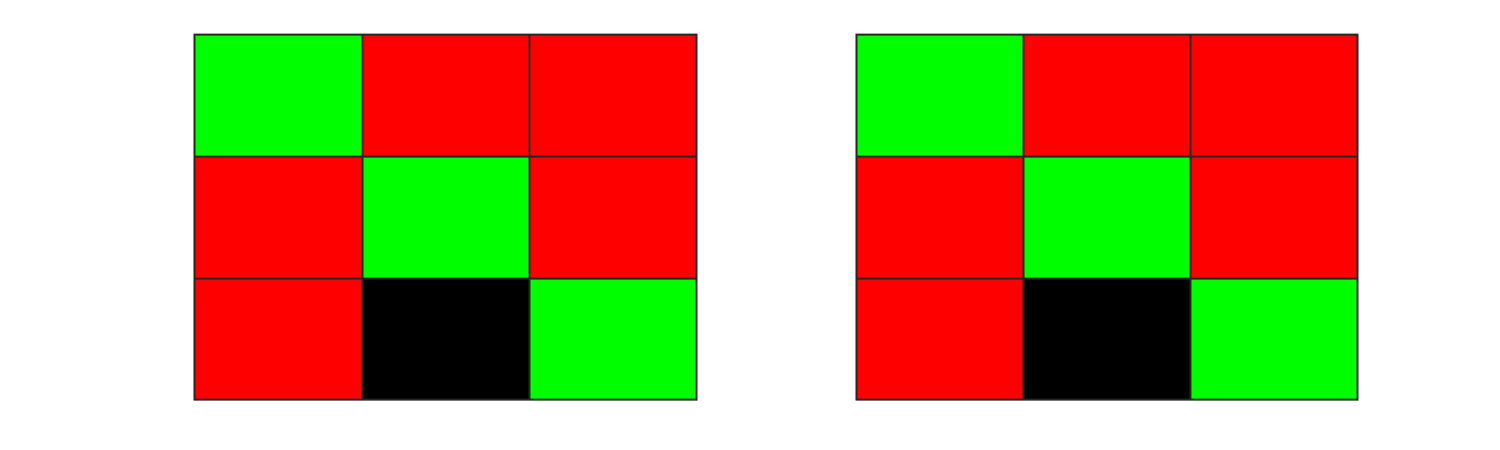

### 3D.tiff

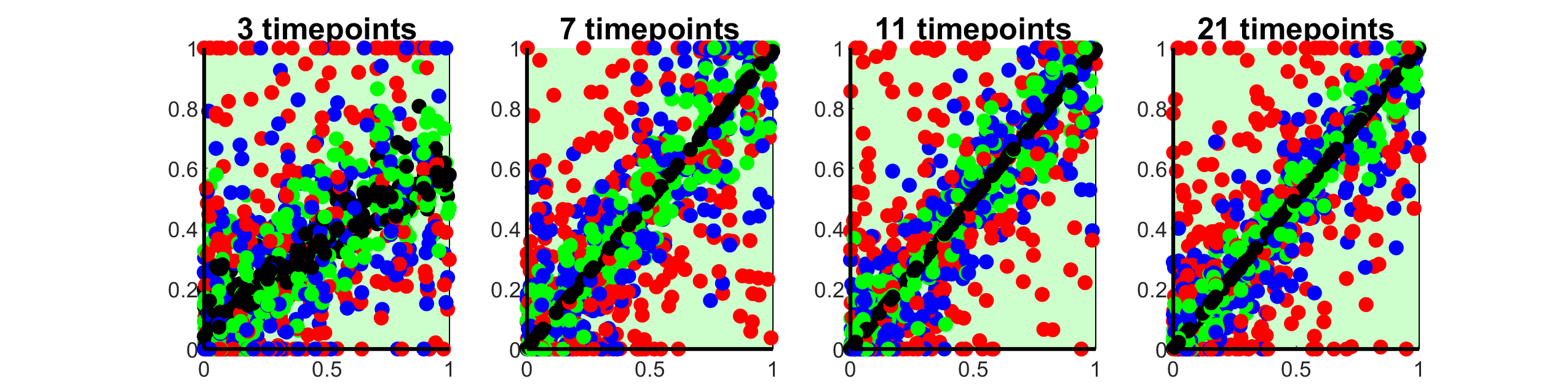

### 4C.tiff

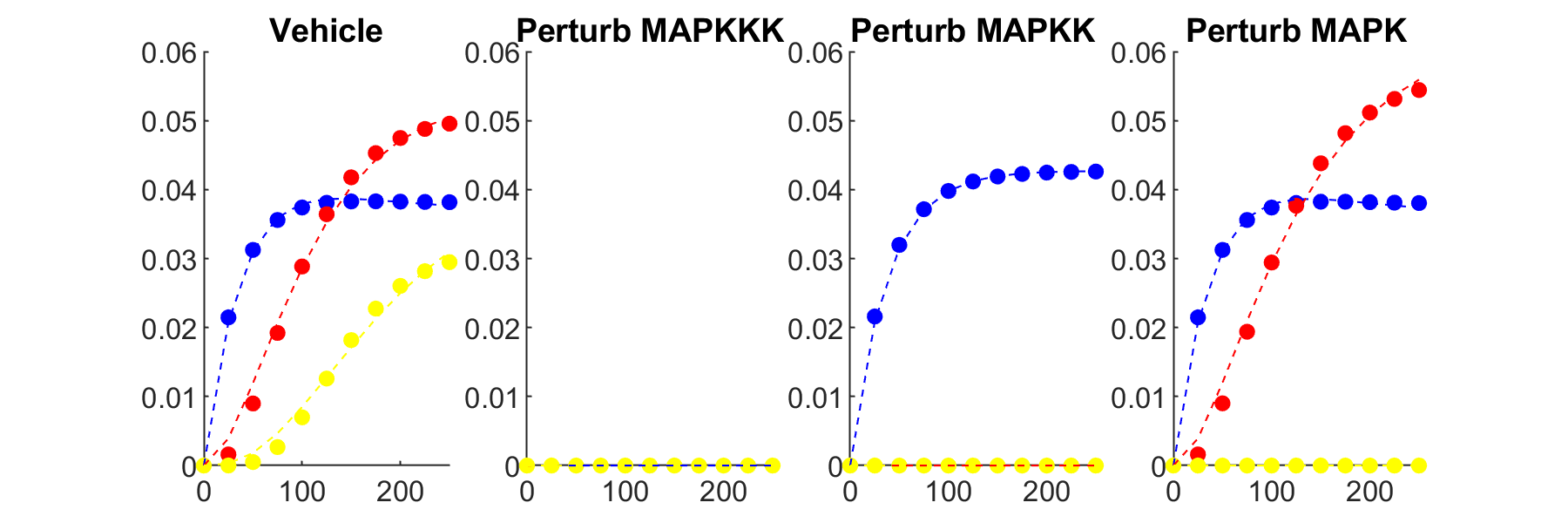

### 5B.tiff

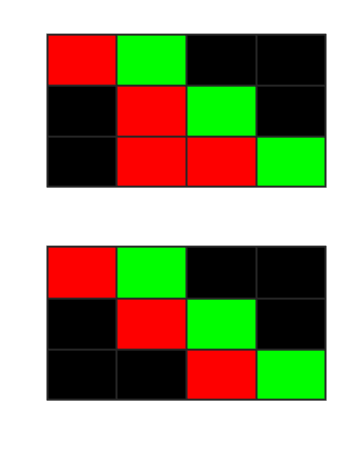

### 5D.tiff

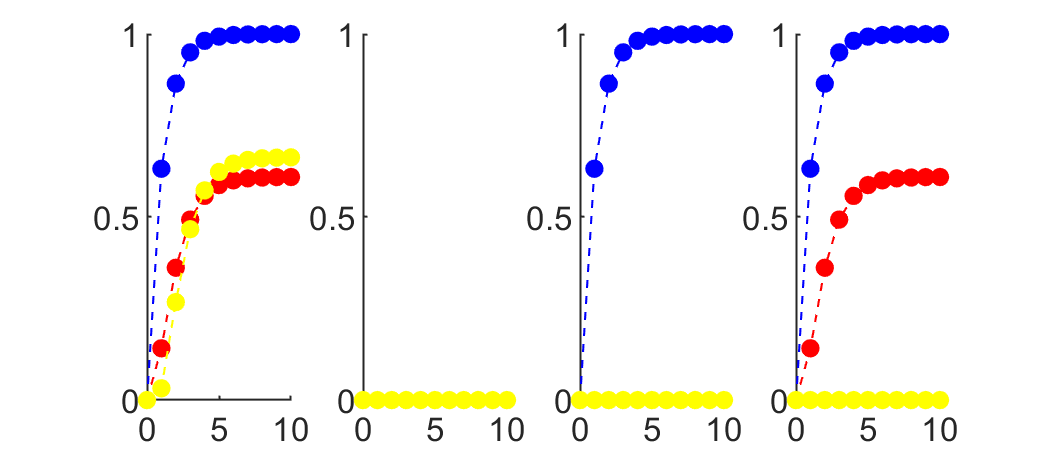

### 5E.tiff

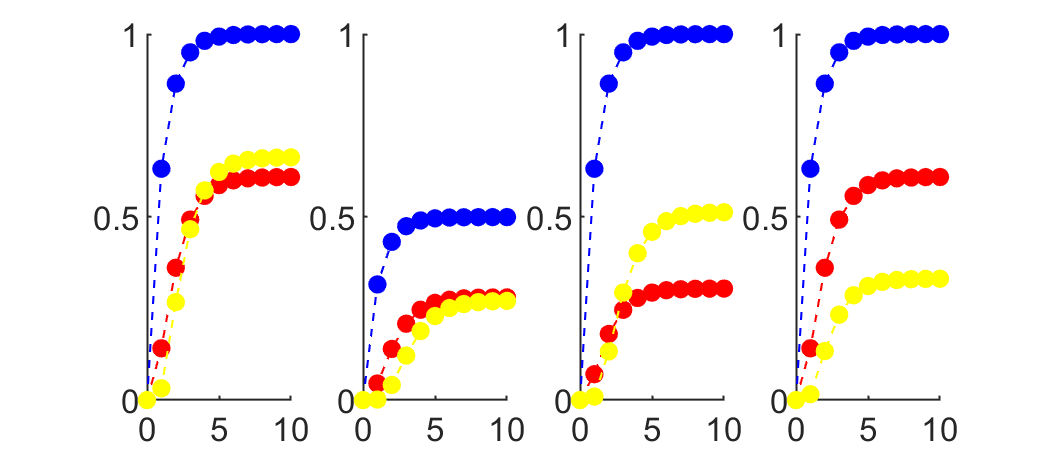

### 5F.tiff

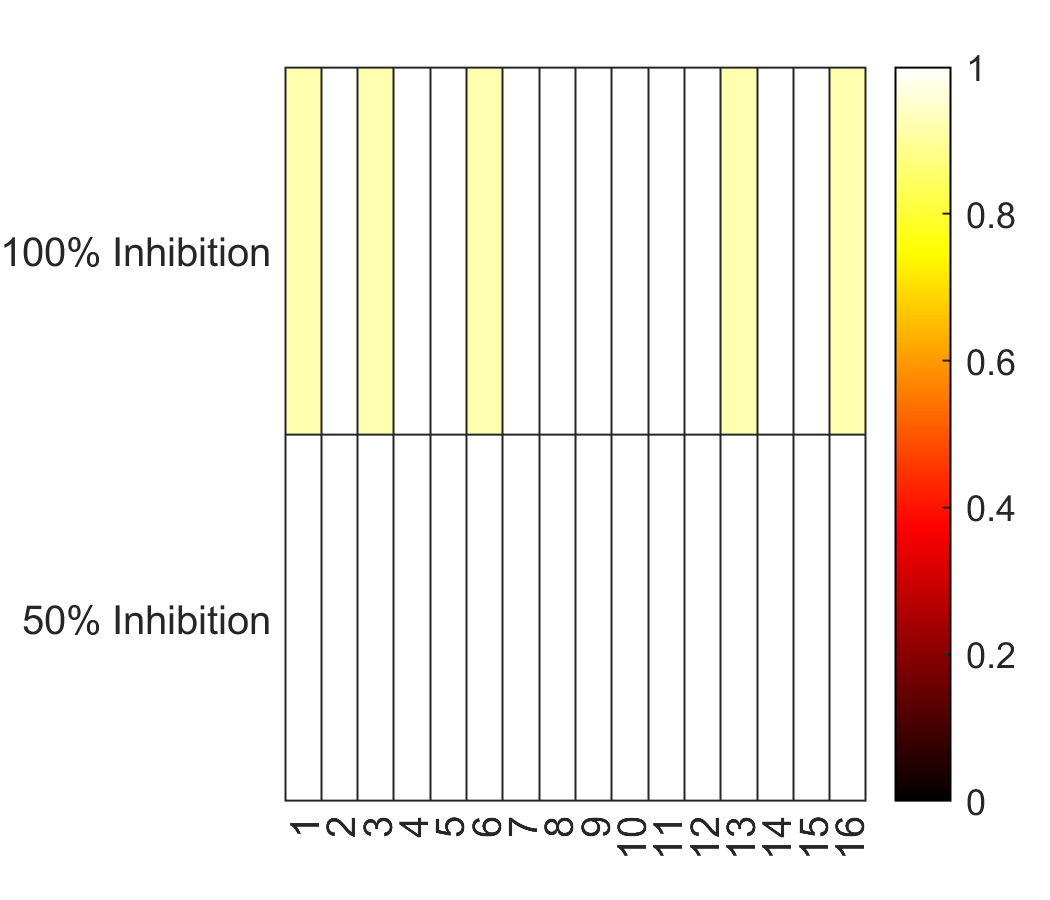

### 6A1.tiff

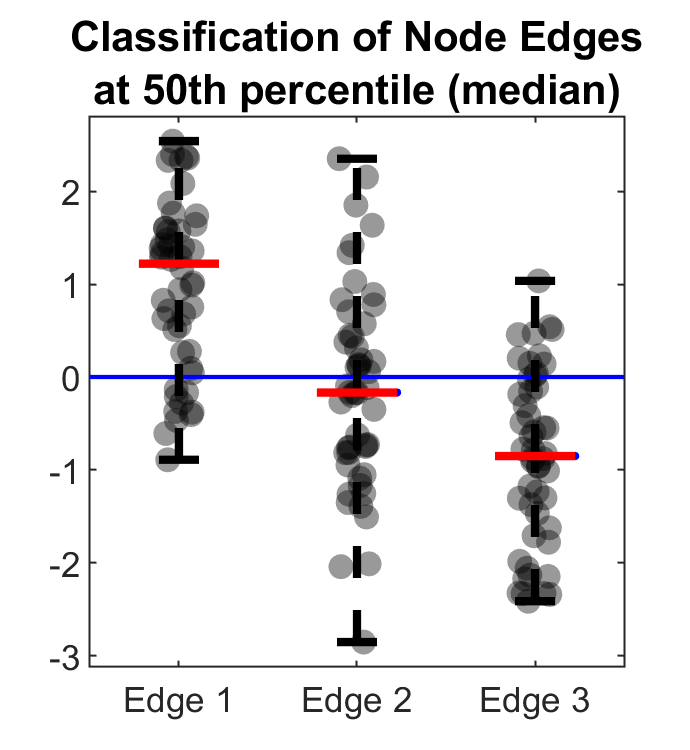

### 6A2.tiff

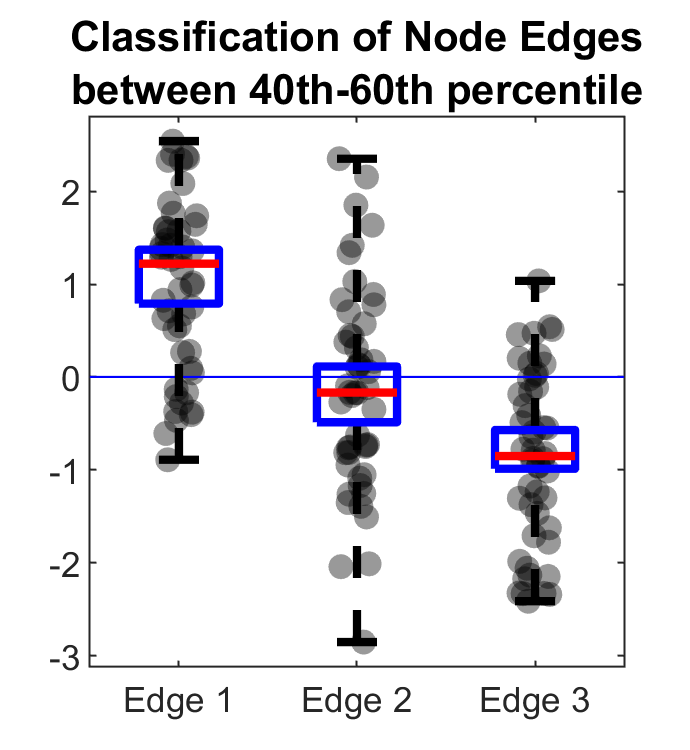

### 6A3.tiff

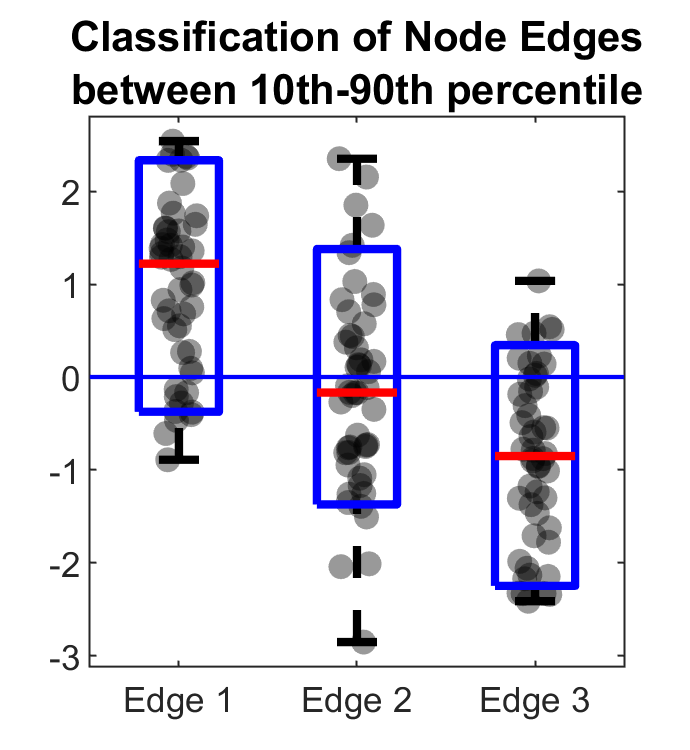

### 6B.tiff

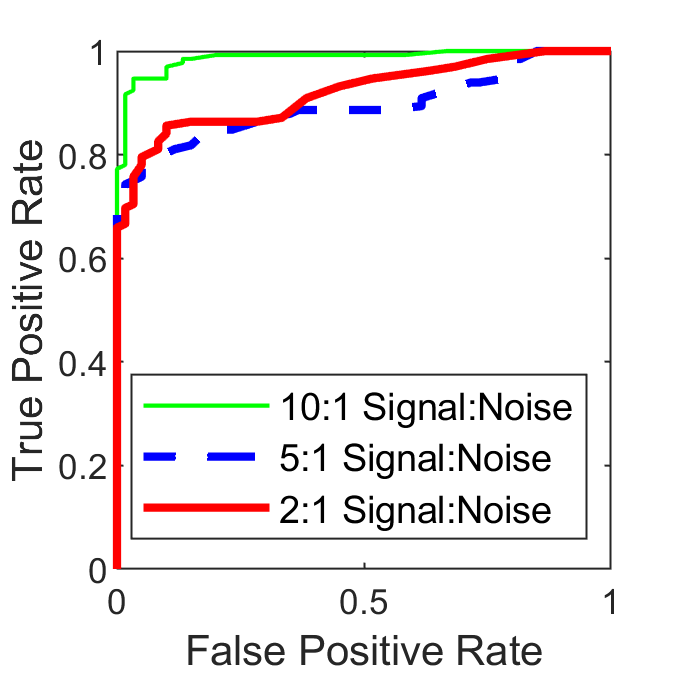

### 6C.tiff

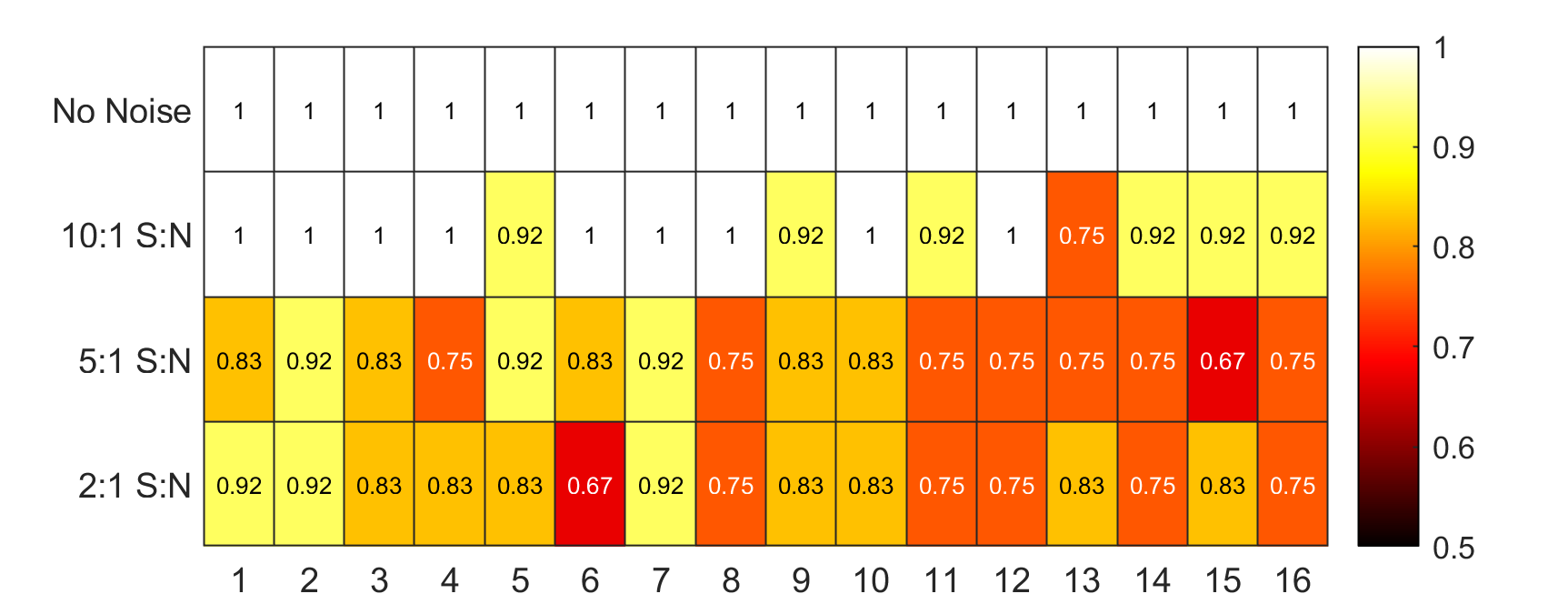

### 6D.tiff

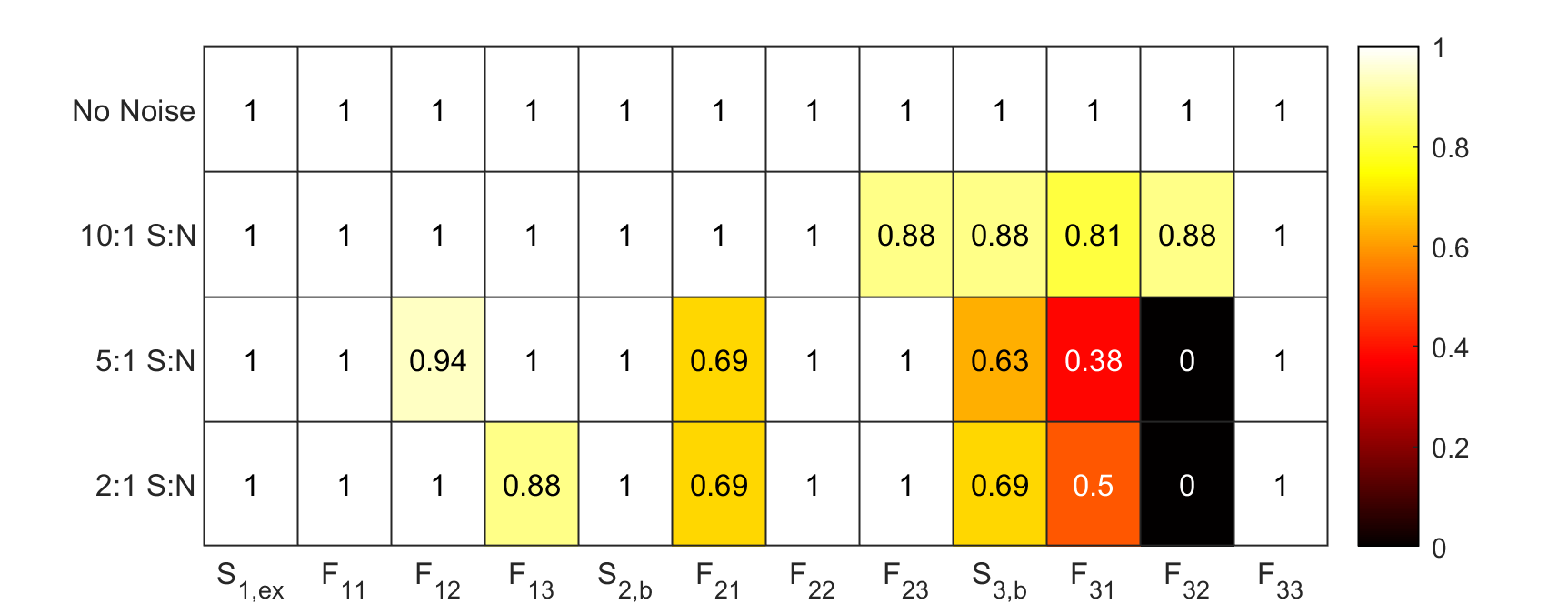

### New2D.tiff

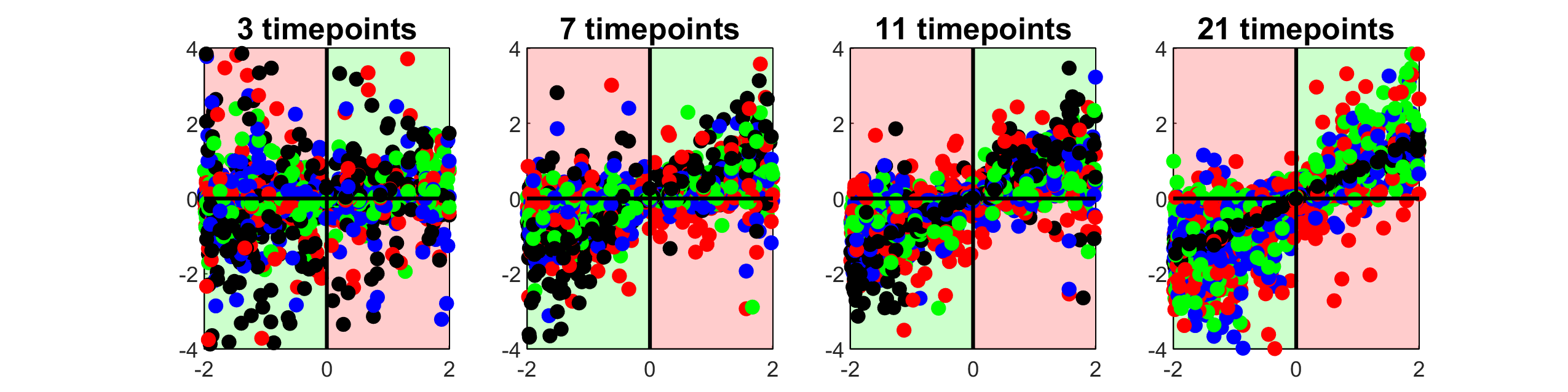

### New2F.tiff

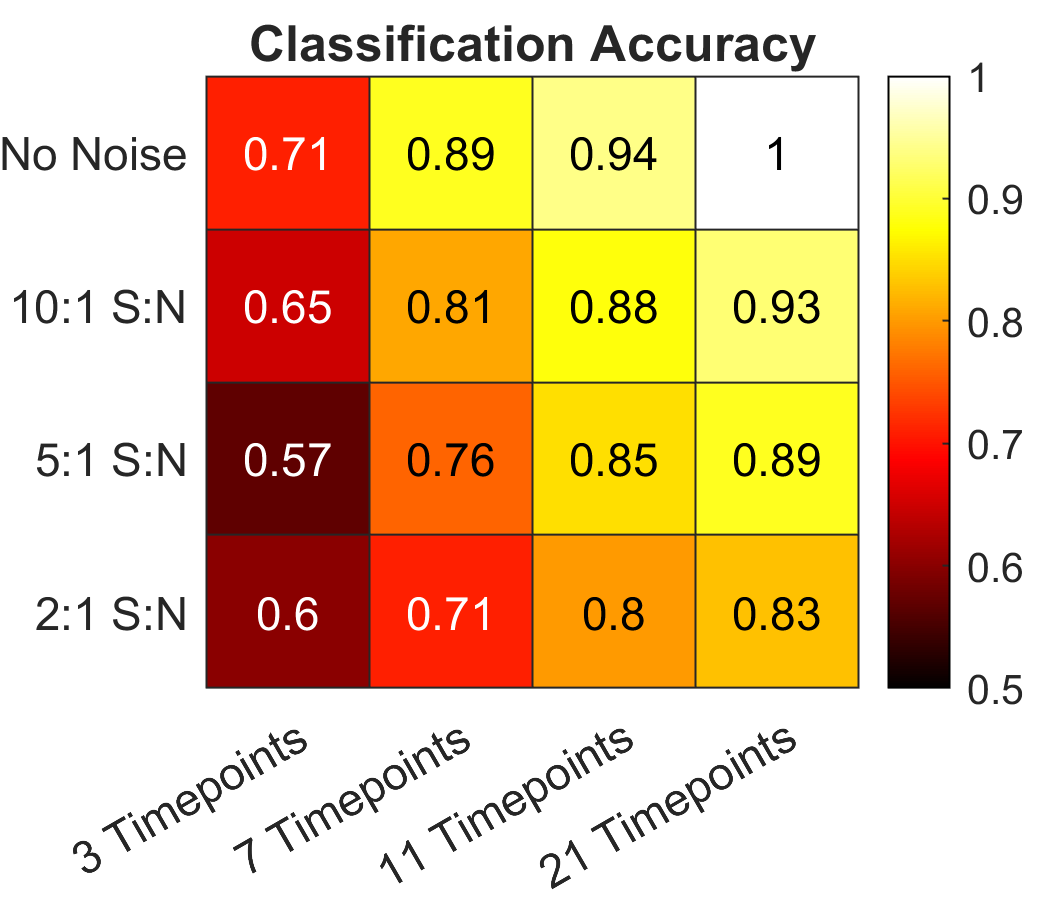

### S1B.tiff

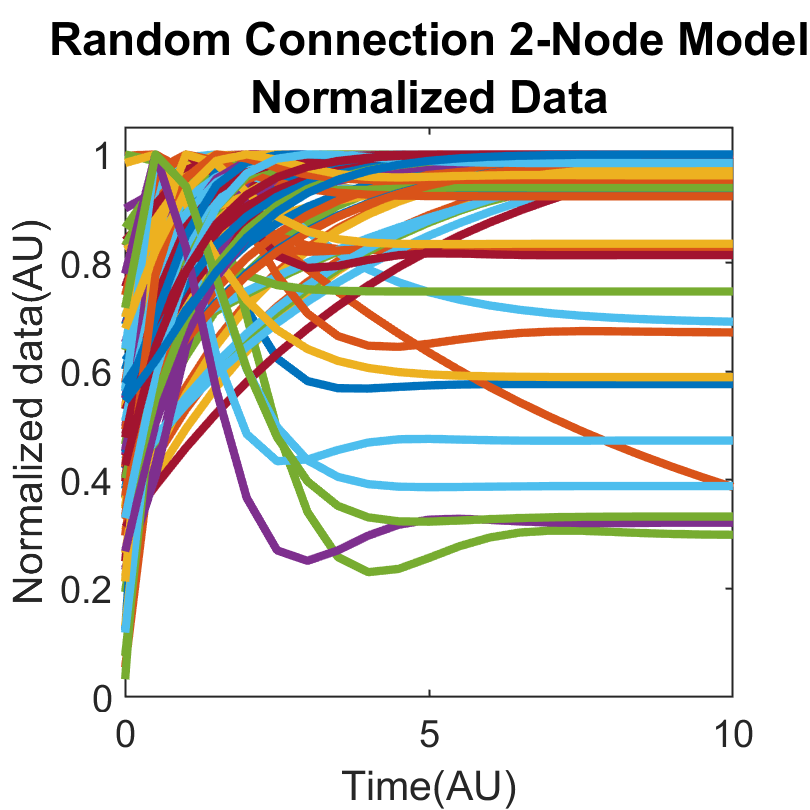

### S1C.tiff

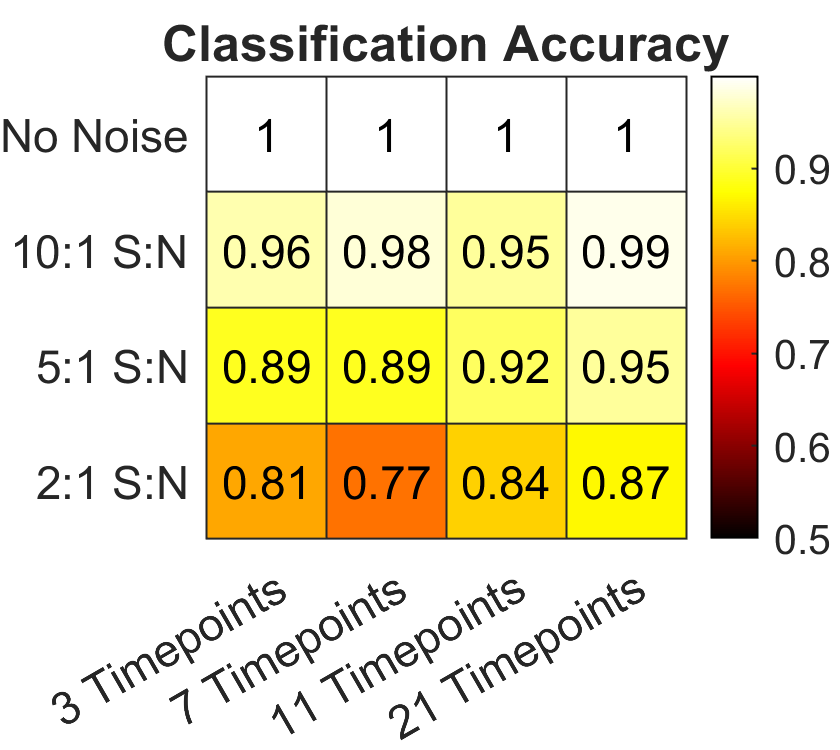

### S1D.tiff

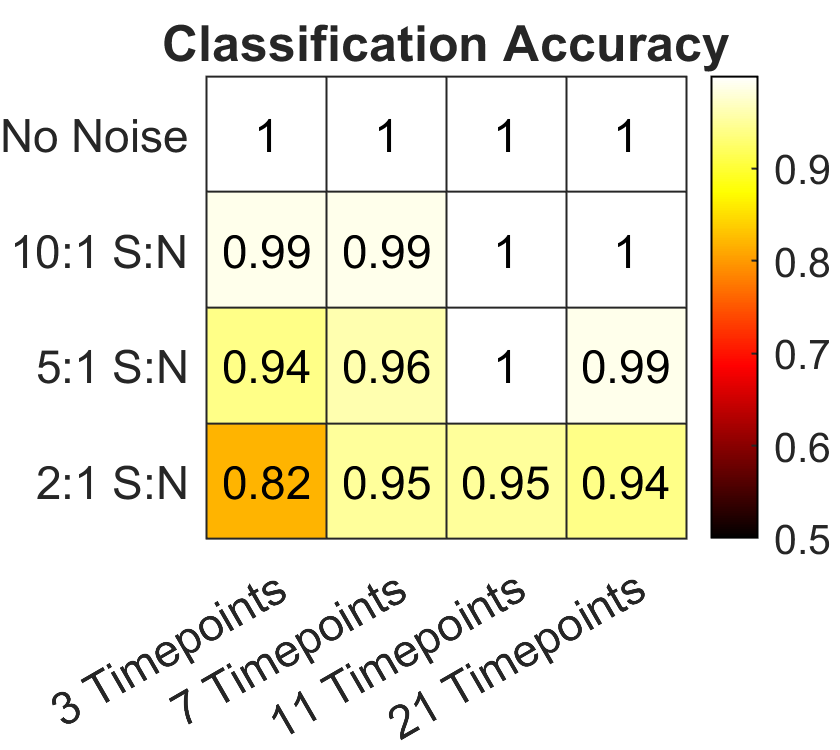

### S1E.tiff

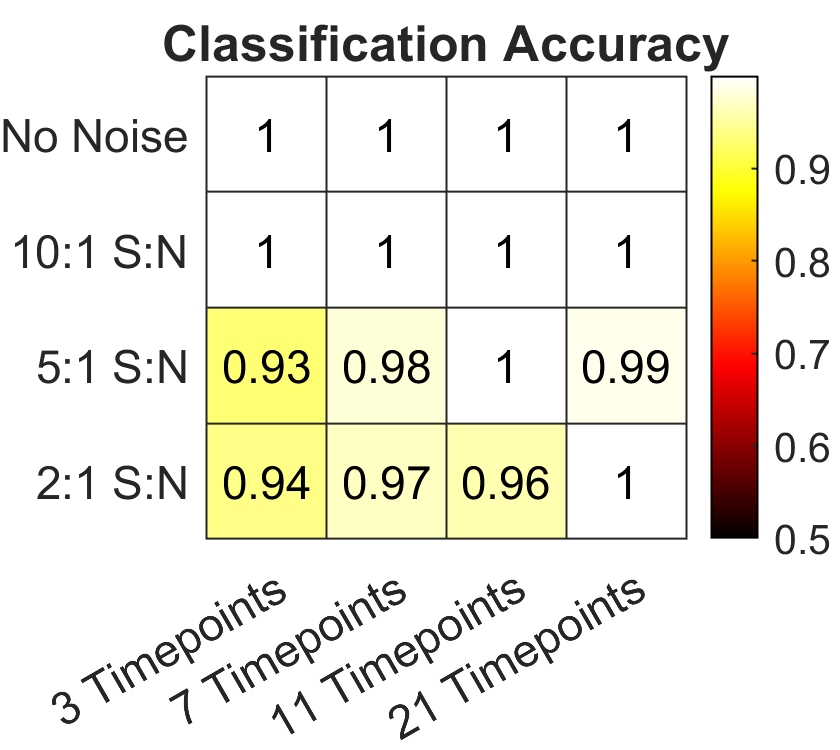

### S1F.tiff

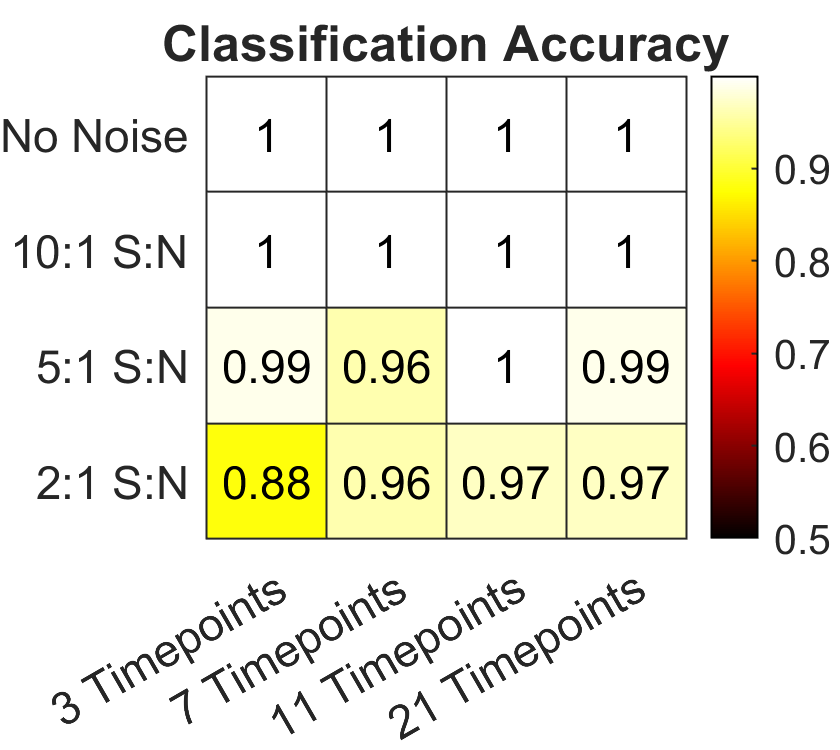

### S2B.tiff

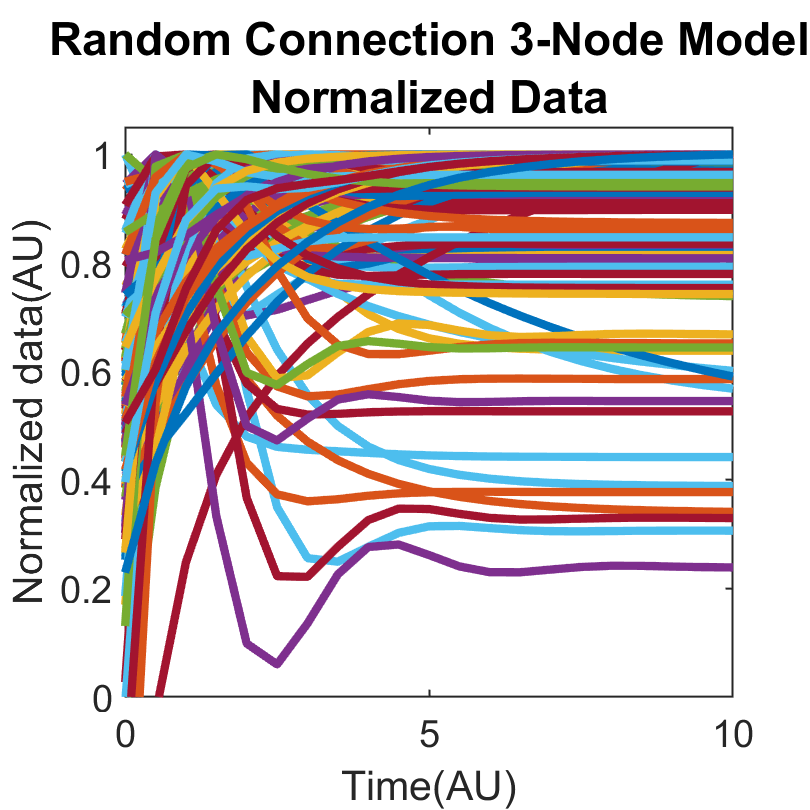

### S2C.tiff

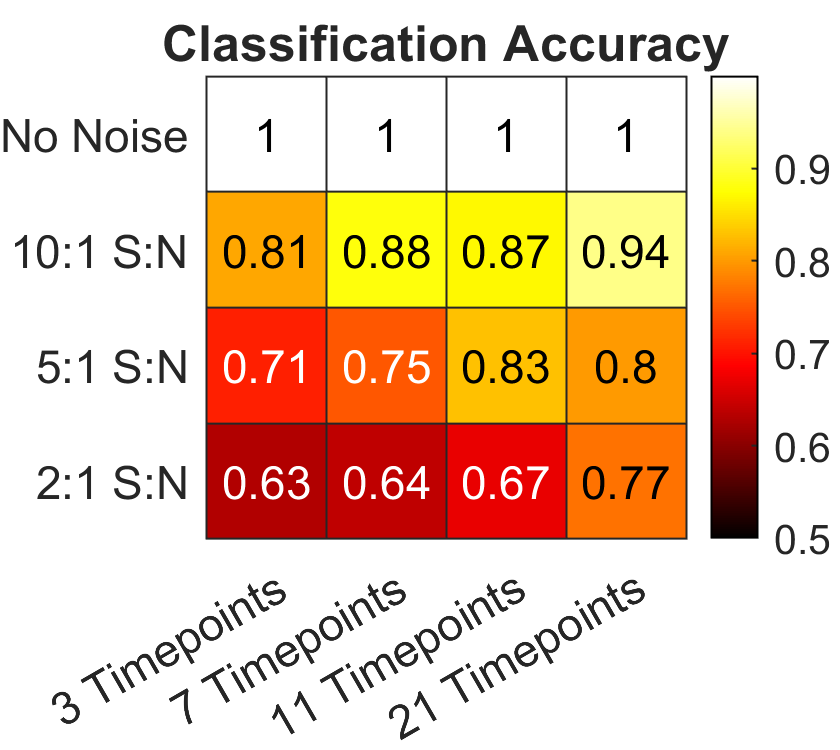

### S2D.tiff

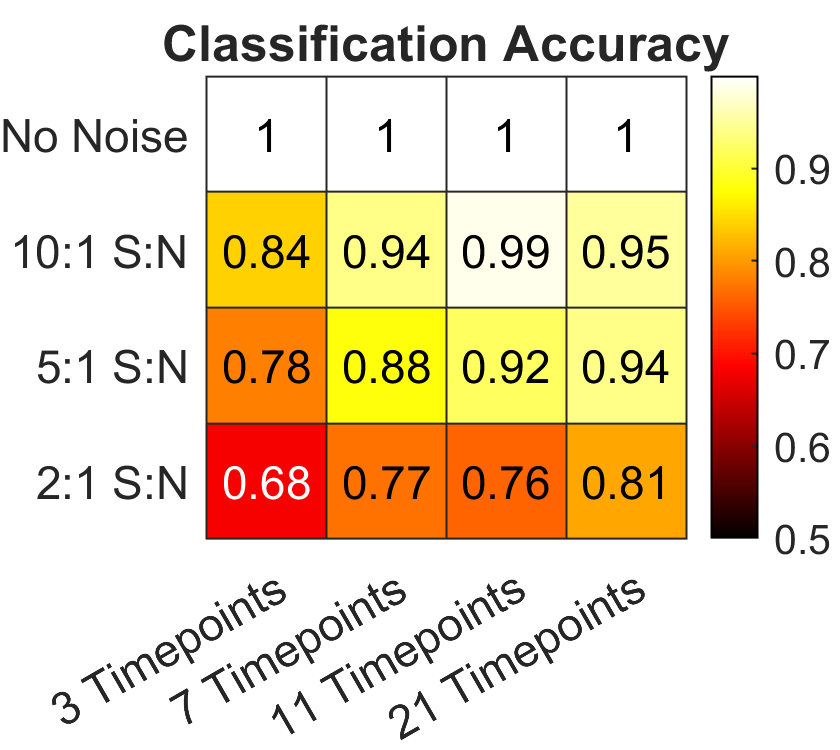

### S2E.tiff

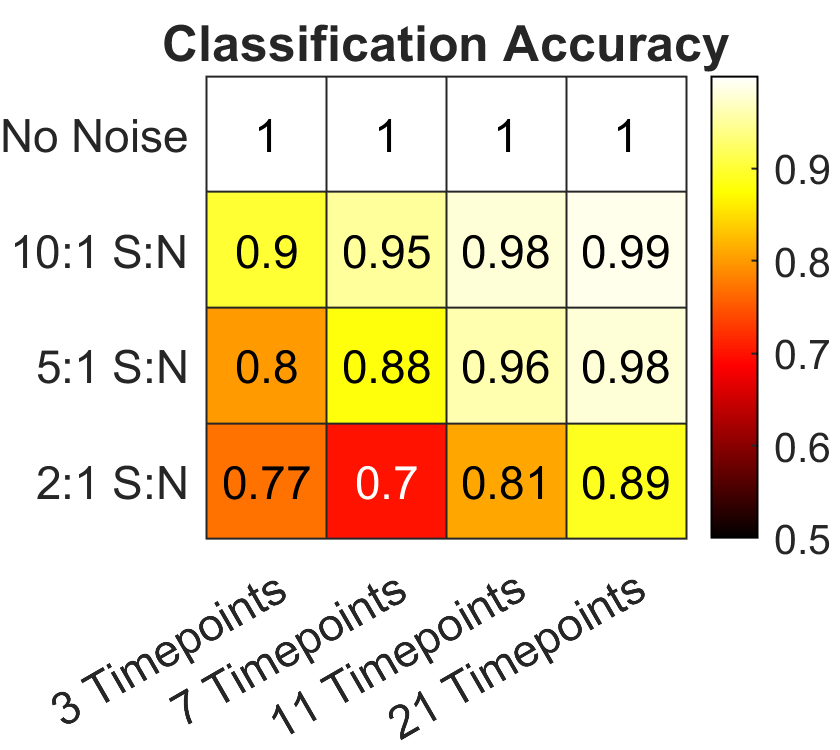

### S2F.tiff

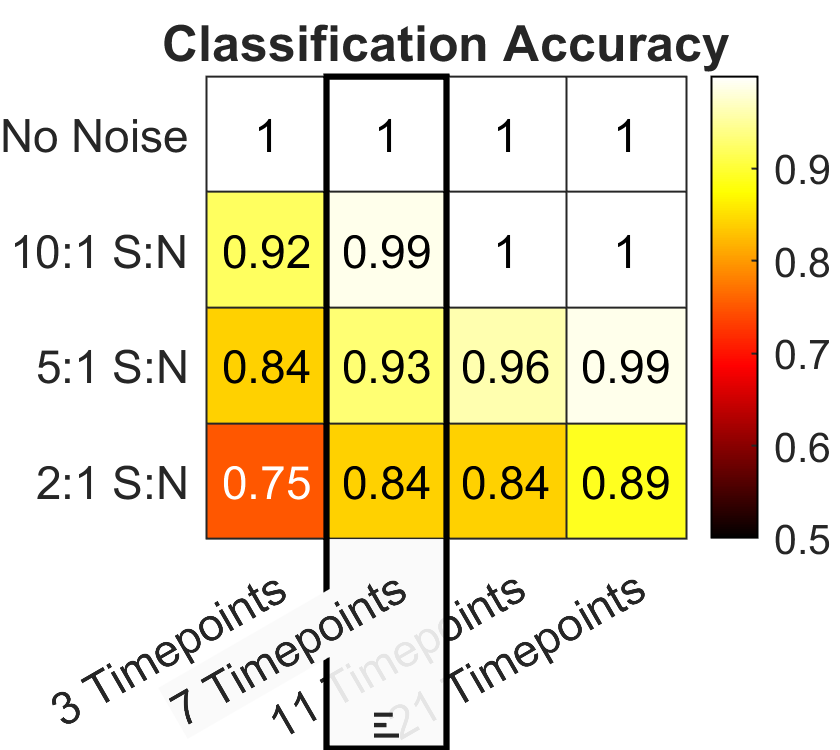

### S3A.tiff

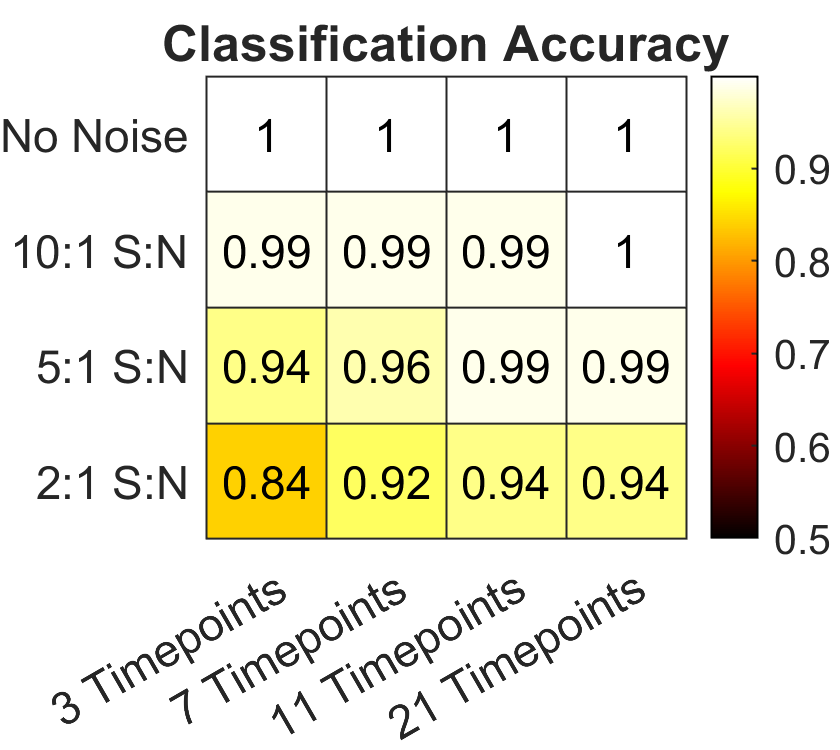

### S3B.tiff

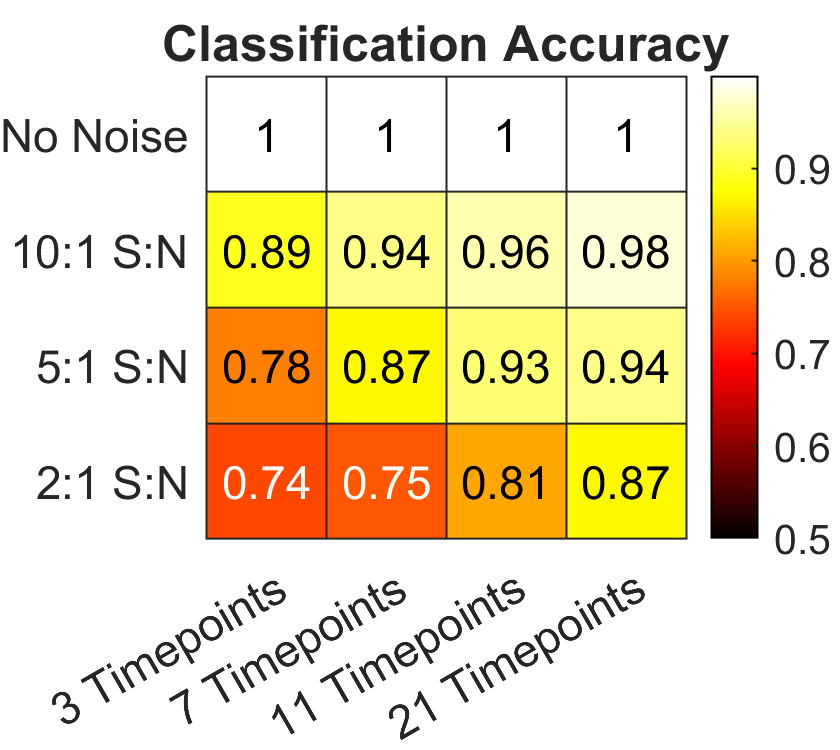

### S4.tiff

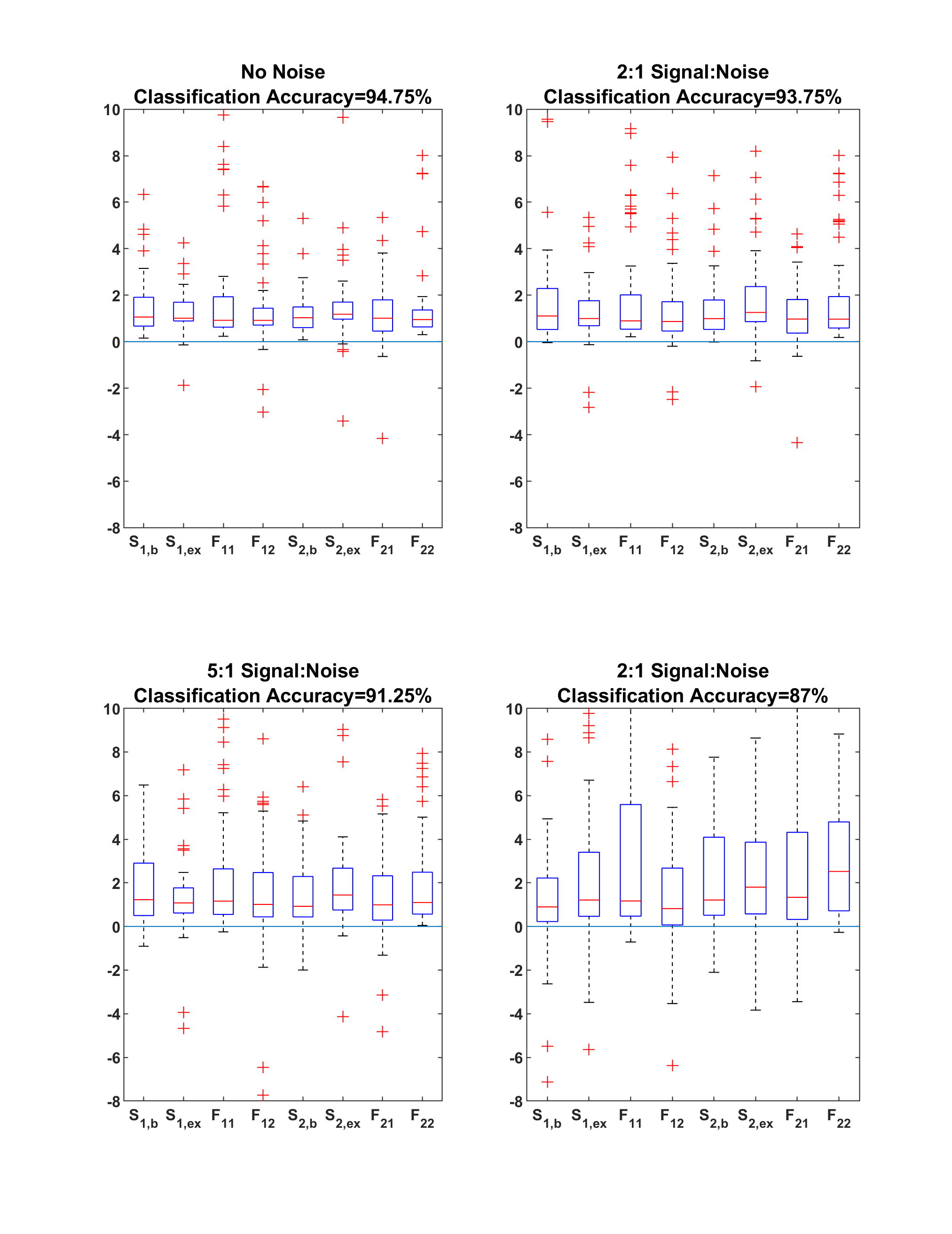

### S5.tiff

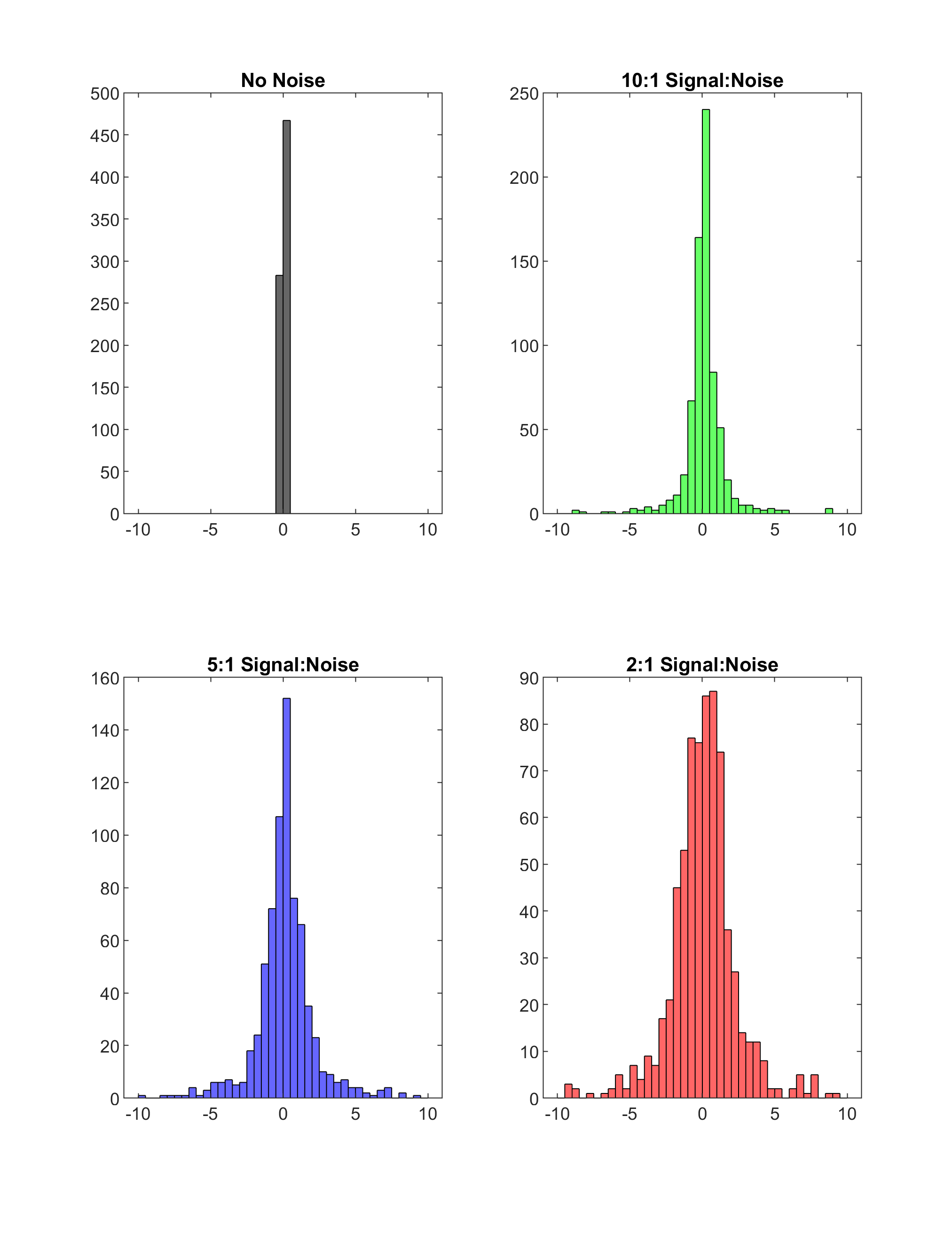

### S6.tiff

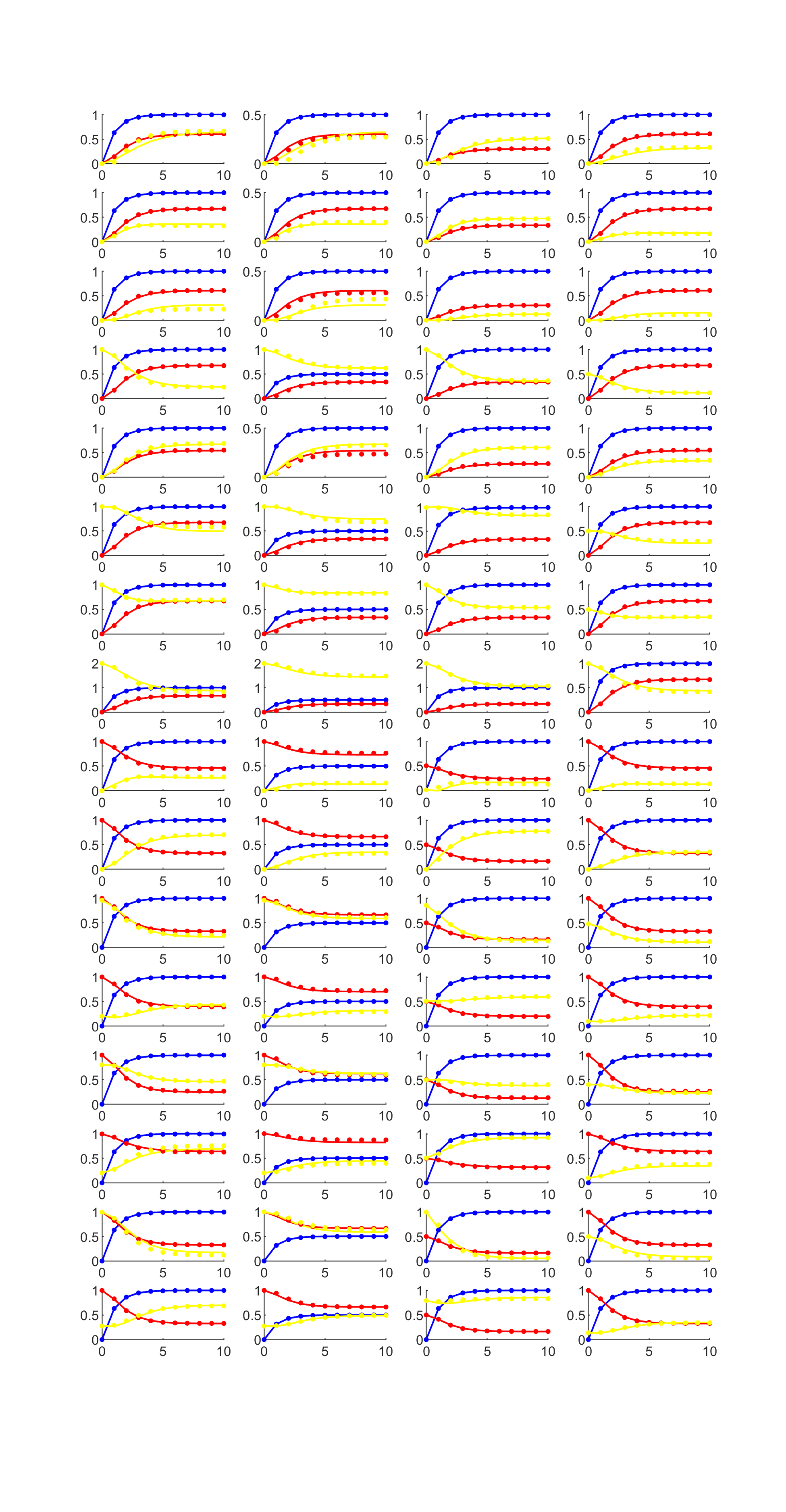

### S7.tiff

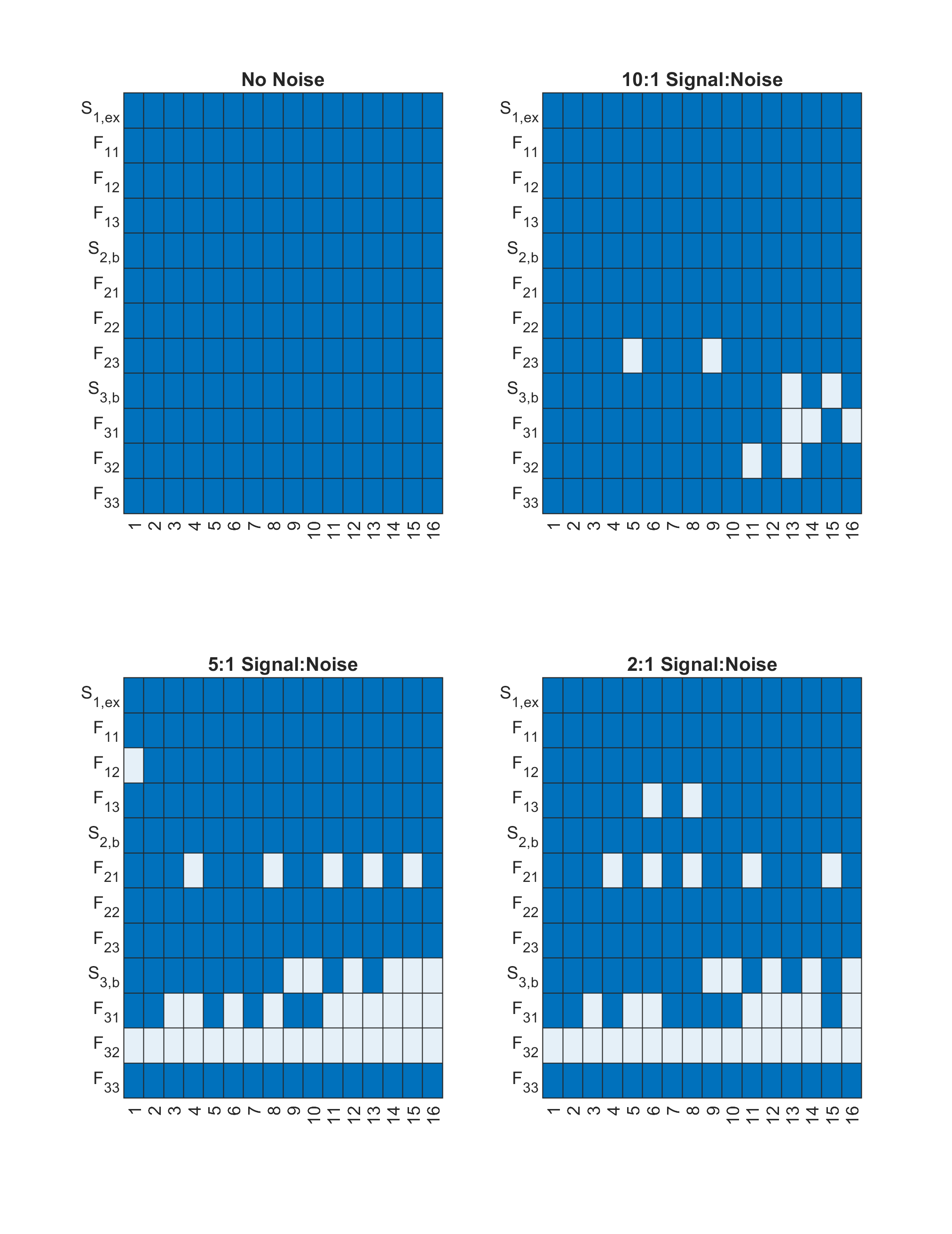
